# Whole chloroplast genomes reveal a complex genetic legacy of lost lineages, past radiations and secondary contacts in the dominant temperate deciduous tree genus *Fagus*

**DOI:** 10.1101/2025.06.03.653586

**Authors:** James R.P. Worth, Guido W. Grimm, Tokuko Ihara-Udino, Pan Li, Aristotelis C. Papageorgiou, Marco C. Simeone, Parvin Salehi Shanjani, José A. López-Sáez, Yu-Chung Chiang, Keiko Kitamura, Nobuhiro Tomaru, Thomas Denk

## Abstract

**Background—:** Major northern temperate tree genera emerged in the Eocene and now have vast ranges across the Northern Hemisphere. Here we undertake a multi-discipline study to provide novel insights into the formation and biogeographic history of a dominant, yet understudied temperate tree genus, *Fagus* L.

**Data and methodology—:** A whole chloroplast genome phylogeny (eighty-two chloroplast genomes) with multi-accessions of most species was determined and contrasted with the timing of lineage diversification inferred from three different nuclear and chloroplast gene time-calibrated phylogenies using 101 fossils as age priors.

**Main results—:** Five deeply diverged chloroplast lineages were revealed that, with the exception of a Eurasian lineage, underwent divergence decoupled from speciation predating modern species lineages by up to 28 million years (Ma). The geographic distribution of chloroplast lineages reflects a complex history of genetic admixture during past contact between extinct and modern species including between different subgenera in East Asia and long-term persistence particularly in Japan which is the hotspot of plastome diversity.

**Conclusion—:** Modern *Fagus* forests are the consequence of 62 million years of evolution, migration, genetic exchange and species extinction with most modern species harbouring a mosaic of genetic material, a legacy of multiple admixture events in deep time.

## Introduction

Temperate deciduous forests are extensively distributed across the Northern Hemisphere occupying mid-latitudes between conifer dominated boreal forests to the north and evergreen subtropical forests to the south (Olson et al. 2001). Despite their vast and fragmented distribution across different continents, temperate deciduous forests are similar in structure and physiognomy sharing many tree genera including *Acer*, *Betula*, *Carpinus*, *Castanea, Corylus*, *Fagus*, *Fraxinus*, *Quercus*, *Tilia* and *Ulmus*. Similar assemblages can be traced back until the Eocene (56 to 33.9 Ma; Dillhoff et al. 2005), while some extant genera were already present by the Late Cretaceous and Paleocene (∼80–60 Ma; Grímsson et al. 2016). Fossils of dominant temperate deciduous tree species have been extensively preserved in the fossil record, providing a broad picture of the processes that have shaped trans-continental distribution of modern temperate deciduous forests, and their constituent genera (Tiffney 1985, Manchester 1999). The fossil record provides direct information about (*i*) the timing of evolution and extinction of lineages, (*ii*) intra-continental range shifts during glacial-interglacial cycles and (*iii*) the role of ephemeral land bridges, the Bering land bridge (BLB; connecting East Asia and Western North America) and the North Atlantic land bridge (NALB; linking Western Eurasia and Eastern North America), for inter-continental dispersal during the past 65 or more million years (see Milne 2006).

Molecular-phylogenetic analyses of dominant temperate deciduous genera can offer powerful insights into the biogeographical history of these forests (Petit et al. 1993, 2002, Okaura & Harada, 2002; Magni et al. 2005) and may not only help to test hypotheses based on fossil evidence such as the direction and timing of past dispersal but could also reveal information that is difficult or impossible to decipher from the fossil record alone: *(i)* past overlapping ranges of congeneric species as evidenced by exchange of genetic material; *(ii)* the number of dispersal events during migration; and *(iii)* the geographical boundaries of glacial refugia. Due to their unique characteristics, nuclear and organellular genomes can provide different insights into the past. The maternal inheritance of organelle genomes in many angiosperms generally results in their poor dispersal capacity. This, coupled with their high potential for “capture” via asymmetric introgression and hybridisation, means that organellular lineage distributions can be strongly correlated with geography and dispersal history rather than species boundaries (Zhou *et al*. 2022; McLay *et al*. 2023). This is especially true for wind-pollinated woody angiosperms such as Fagaceae and Nothofagaceae as well as some insect-pollinated genera such as *Eucalyptus*, in which restricted seed dispersal has led to a (pronounced) intraspecific subdivision of their chloroplast genomes (Petit et al. 2002, McKinnon et al. 2004, Acosta & Premoli 2010, Li et al. 2025), thereby increasing effective population size (Hoelzer, 1997). In contrast, nuclear DNA variation that can be transmitted by highly dispersed pollen as well best reflects species history due to the usually high gene flow between constituent populations (Petit & Excoffier 2009).

This paper utilises DNA variation from chloroplast genomes and (up to) 31 nuclear genetic markers coupled with fossil-calibrated molecular dating to examine the speciation and migration history of a key Northern Hemisphere temperate deciduous forest genus–*Fagus*, the beech. *Fagus* is a small genus with 14 species subdivided in two subgenera: *Fagus* subgenus *Fagus,* with disjunct distribution areas in Western Eurasia (4 spp.), East Asia (5 spp. + probably extinct *F. chienii* Cheng) and (eastern) North America (2 spp.) and *Fagus* subgenus *Englerianae* Denk & G.W.Grimm with three species in East Asia (Denk et al. 2024). Despite their low number of species, beeches are the dominant trees of temperate forests across large parts of the Northern Hemisphere in three continents (Peters 1997; **Fig. 1**).

**Figure 1.**
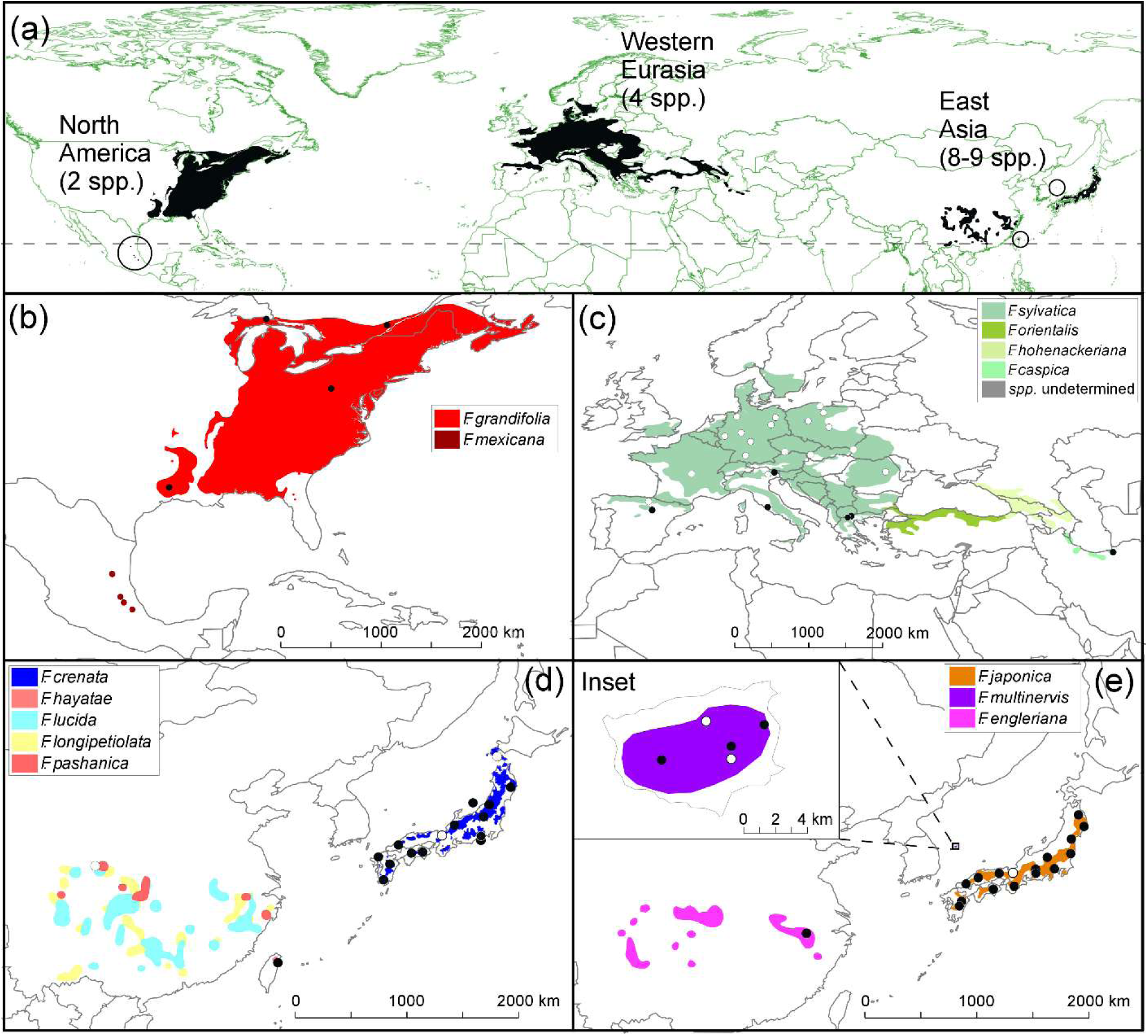
| **Distribution of *Fagus* species across the Northern Hemisphere** (black area) with small areas of occurrences indicated by circles (a). The source of each sample (black dots for those sampled for this study and white circles for those sourced from NCBI GenBank) where geographical reference information was available is shown for North America (b), Western Eurasia (c), East Asian species of *Fagus* subgenus *Fagus* (d) and East Asian *Fagus* subgenus *Englerianae* (e).

Early evidence from Sanger-based sequencing studies showed that the chloroplast signatures of *Fagus* do not follow species or even subgeneric boundaries (Stanford 1998, Manos & Stanford 2001, Okaura & Harada 2002, Oh et al. 2016). In line with its short-distance seed dispersal (Martínez & González-Taboada 2009, Oddou-Muratorio et al. 2010), phylogeographic studies of *Fagus* species have revealed strong geographic structuring of chloroplast variation consistent with locations of glacial refugia and migration routes (Demesure et al. 1996, Magri et al. 2006, Ying et al. 2016).

The fossil record of beech is extensive and allowed timing of intrageneric divergences (Grímsson et al. 2016, Renner et al. 2016), placing the first radiations into the Eocene. Recent studies using nuclear data show that its history, particularly within Eurasia, involved past genetic exchange between species’ lineages (Cardoni et al. 2022, data generated by Jiang et al. 2022) that cannot be modelled using a single species tree.

Here, we utilise a near genus-wide sampling of chloroplast genomes, including all traditionally recognised species (1–2 in Western Eurasia, single species in North America; Shen 1992; Govaerts & Frodin 1998), to elucidate past migration and mixing patterns of *Fagus*, establish a spatio-temporal framework for “chloroplast capture”, and date the formation of extant species. With the exception of Chinese species, geographically spread multiple accessions of each species are included with a particular focus on the Japanese Archipelago. Japan is key to resolving phylogenetic relationships in *Fagus*, including ancient reticulation, due to the remarkable sharing of chloroplast variation in *F. crenata* Blume and *F. japonica* Maxim. (Okaura & Harada 2002) despite belonging to long-established distinct evolutionary lineages (Denk et al. 2005, Denk & Grimm 2009, Renner et al. 2016) and the nesting of Western Eurasian species (*F. sylvatica* L. s.l. according to Govaerts & Frodin 1998) within this variation (Okaura & Harada 2002). In addition, the Japanese Archipelago is a major suture zone in East Asia for a diverse range of organisms (Aoki et al. 2011, Iwasaki et al. 2012, Tojo et al. 2017), and a major cross-road for ancient and more recent *Fagus* migration in the North-East Asia region, the biodiversity centre of *Fagus* during the Miocene (Denk & Grimm 2009). To overcome possible problems of low genetic variability of chloroplast markers (Stanford 1998, Manos & Stanford 2001, McLachlan et al. 2005, Paffetti et al. 2007, Oh et al. 2016) we employed whole chloroplast genome sequencing to maximise the potential to discover genetic variation. In addition, we sequenced two single-copy nuclear gene regions (Crabs Claw Gene [*CRC*], Oh & Manos 2008; second LEAFY intron [LFYi2], Renner et al. 2016) for a representative subset of samples used for chloroplast analyses to assess and confirm their systematic-phylogenetic placement. We use the newly generated and published nuclear data (Renner et al. 2016, Jiang et al. 2022) to establish species consensus data for testing and dating several topological alternatives in a fossilized birth-death (FBD) dating framework (Heath et al. 2014) building on Renner et al. (2016). In addition, we applied traditional node dating (ND) using a selected set of fossil priors to establish absolute minimum estimates for plastome and nucleome divergences. Our set-up recognizes that the modern species are likely genetic mosaics and that their formation involved multiple reticulation events (Cardoni et al. 2022); the modern species form a species network (**Fig. 2**).

**Figure 2.**
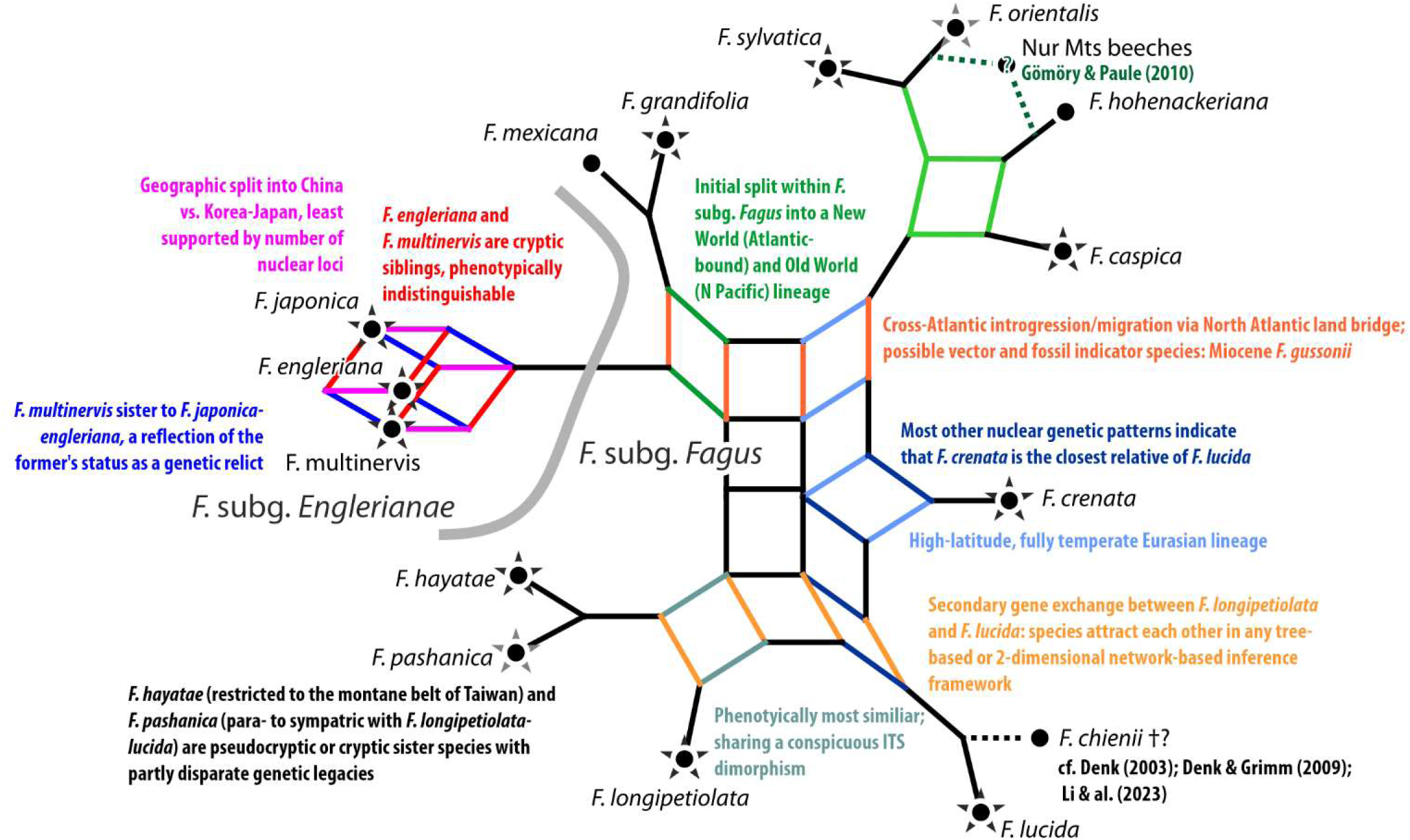
| Consensus network summarising all known inter-species relationships in beeches based on various nuclear and morphological studies. (Denk 2003, Denk et al. 2005, Grimm et al. 2007, Denk & Grimm, 2009, Grímsson et al. 2015, Renner et al. 2016, Cardoni et al. 2022, Jiang et al. 2022). *Fagus chienii* is probably extinct: a recent recollection in the original area only retrieved individuals morphologically (and genetically) indistinguishable from *F. pashanica* (Li et al. 2023). Stars highlight species covered in this study including newly generated data (black stars) and data harvested from gene banks (grey stars). After Denk et al. (2024).

By comparing chloroplast and conflicting nuclear dated phylogenetic trees on the background of past *Fagus* distribution patterns, we put forward a comprehensive history of *Fagus* and explain the processes underlying plastid hyperdiversity characterizing the three surviving species in modern-day north-eastern Asia, *F. crenata* (*F.* subg. *Fagus), F. japonica* and *F*. *multinervis* Nakai (*F.* subg. *Englerianae*).

## Materials and Methods

### Sampling

A total of 48 *Fagus* samples (**Fig. 1**; **Table SM3-1** in SupplM&M.pdf) were collected representing all three species of *Fagus* subgenus *Englerianae* and eight (out of 11) species of *F.* subgenus *Fagus* (Denk et al. 2024). In Japan, 28 representative samples (11 with determined *crenata-*haplo-types; **SupplGenetics_cpDNA.xlsx**, sheet *New and used plastomes*) were selected based on previous range-wide genetic studies of *F. crenata* (Fujii et al. 2002) and *F. japonica* (Hiraoka & Tomaru 2009).

### Chloroplast genome assembly

DNA extractions were done using a modified CTAB protocol (Doyle 1990). DNA concentration and quality were assessed by agarose gel electrophoresis and a Qubit 2.0 fluorometer (Life Technologies). DNA was sent to the Beijing Genomic Institute where short-size Truseq DNA libraries were constructed and paired-end sequencing was performed on an Illumina HiSeq2000 Genome Analyser. The assemble of the whole chloroplast genomes was undertaken using default settings in GETORGANELLE (Jin et al. 2020). Where samples did not produce a circular genome, assembly was achieved by mapping contigs to the *F. crenata* whole genome (GenBank accession MH171101.2) in GENEIOUS v. 9.0.5 (Biomatters, Auckland, New Zealand).

### Sequencing of nuclear genes

A subset of the samples used for whole chloroplast genome sequencing (**SupplGenetics_ cpDNA.xlsx**, sheet *New and used plastomes*) were sequenced for the single-copy nuclear loci second intron of *LEAFY* (LFYi2) and *CRABS CLAW* (*CRC*) gene. The former has been well characterized in *Fagus* (Oh et al. 2016, Renner et al. 2016) while the latter has been useful in resolving Fagaceae phylogenetic relationships (Oh & Manos 2008), although only two sequences of the American *F. grandifolia* Ehrh. (s.str.) and *F. mexicana* Martinez have been produced in *Fagus*. Assumedly single-copy, both nuclear markers show intra-individual sequence polymorphism. To increase LFYi2 amplification success, two primer pairs were designed (see **Table SM2-2** in SupplM&M.pdf); PCR and cloning protocols are detailed in **SupplM&M.pdf**, *section 2.8.1*.

### Phylogenetic and sequence diversity analyses

The 48 new chloroplast genomes were combined with 34 from GenBank representing eleven species (**Table SM3-1** in SupplM&M.pdf). All 82 chloroplast genomes were aligned using MAFFT v. 1.3.6 (Katoh et al. 2002) in GENEIOUS v. 9.0.5. Phylogenetic relationships were investigated using two methods after deleting the first inverted repeat (IRa) and the *ycf1* pseudogene. Firstly, a neighbour-net (Bryant & Moulton 2004) based on uncorrected *p*-distances (excluding indels) was constructed in SPLITSTREE v. 4.14.4 (Huson & Bryant 2006). Secondly, phylogenetic trees were inferred using maximum likelihood with RAxML v. 8.2.10 (Stamatakis 2014; see **SupplM&M.pdf**, *section 4*, for details); (competing) branch support was estimated via non-parametric bootstrapping (Felsenstein 1985) and visualised using consensus networks (Holland & Moulton 2003, Schliep et al. 2017). Three partition schemes were used: (1) unpartitioned; (2) full-partitioned based on the annotation of *F. crenata* chloroplast genome with separate partitions for every intergenic spacer, intron, tRNA and rRNA, 1^st^, 2^nd^ and 3^rd^ codon position of each protein-coding gene; and (3) a “logical” partition analysis with five partitions: (a) intergenic spacers (non-coding, neutrally evolving); (b) introns (non-coding, typically low-divergent), (c) 1^st^ + 2^nd^ codon positions (determining amino-acid, conserved); (d) 3^rd^ codon positions (high level of synonymous mutations); and (e) tRNAs + rRNAs (structurally constrained, highly conserved). The resulting best tree was then rooted in FIGTREE v. 1.4.3 (Rambaut, 2014) at the branch leading to the *F. grandifolia* samples.

In addition to the 101 *CRC* sequences from 20 individuals and 161 LFYi2 sequences from 28 individuals obtained in this study, available *CRC* (2 samples) and LFYi2 (77 samples) GenBank accessions of *Fagus* were harvested and included in the analyses (**SupplM&M.pdf**, *section 2*). For final ML tree inference (with **RAxML-NG**, Kozlov et al. 2019; GTR+Γ+I substitution model) and subsequent dating, we used individual-level strict and modal consensus sequences to ensure direct comparability with the chloroplast tip set; neighbour-nets were based on “phylogenetic Bray-Curtis” distances (Göker & Grimm 2008; **SupplM&M.pdf**, *section 4.2*).

Sequence diversity of chloroplast and nuclear alignments was calculated in DNASP v. 6.12.03 (Rozas & Rozas 1999) including the number of single nucleotide polymorphisms, parsimony informative sites and indels and nucleotide diversity. For in-depth analyses of mutational patterns, we inferred (full) Median networks (Bandelt & Forster 1999, Bandelt et al. 2000) for selected plastid sequence regions.

All input (matrices) and inference files (trees, networks) are included in the Online Supporting Archive (OSA).

### Molecular dating

Using tree-based models to date divergence is problematic in *Fagus* because of probable repeated reticulation events during the past. Ideally, one would infer and date a coalescent multi-species coalescent network (MSCN; Yu & Nakhleh 2015; Allman et al. 2019) but this is not feasible. Explicit MSCN approaches require sufficiently signal-rich multigene data sets and cannot consider the entirety of the fossil record during dating, a core element of our study design. Moreover, they are designed to infer (few) anastomoses in groups of species that, in general, evolve dichotomously and are well sorted. In beech, we look at a highly reticulate evolution of genetic mosaics (Cardoni et al. 2022; this study).

For establishing species divergence and timing of chloroplast genome decoupling, we hence used a tree-based workaround (**Fig. 3**, detailed in **SupplM&M.pdf**, *section 5*). In brief, we used a multi-step protocol using data obtained in this study combined with nuclear data from earlier studies (Denk et al. 2005; Renner et al. 2016; Cardoni et al. 2022; Jiang et al. 2022) to infer species divergence times based on the nuclear data and to correlate them to major phases of plastome divergence: the formation of the five main plastome lineages and their diversification into ‘species-level plastid types’ (SLPTs). The resultant four topological scenarios (**Fig. 4**) and data sets build up a comprehensive species network for *Fagus* (cf. **Fig. 2**) based on the following general assumptions:

1. **Nuclear data capture primary speciation events**—Nuclear divergences represent speciation events, i.e. the formation of new species that are reproductively isolated to a degree that they undergo subsequent homogenisation (accumulation of specific sequence patterns such as SNPs and diagnostic oligonucleotide motifs) and/or genotype stabilisation (accumulation of private, specific polymorphism) because inbreeding outcompetes outbreeding.
2. **Nuclear data capture also secondary reticulation**—Secondary contact of species (hybridisation and introgression) will lead to genetic exchange of evolved, specific nuclear signatures. Corresponding divergence estimates represent the final isolation of two inter-mixing species. That is, the nuclear data divergences may reflect both deep, primary and more recent, secondary speciation events.
3. **Plastid data captures geographic sorting events**—These include allopatric speciation events in the course of major vicariance and dispersal processes *and* intra-species differentiation within a widespread species, defined as a group of populations characterised by shared nuclear genotype(s) and phenotypes. Genetic drift in *Fagus* plastomes is primarily a function of distance, and only secondarily (if at all; e.g. in narrow-endemics with small extant populations, *F. multinervis* and *F. hayatae* Palibin) of effective population size. Thus, plastid divergences can only be established when the (primordial) mothers, the ‘last common mothers’, were already geographically isolated. Two *Fagus* trees growing at the same location and sharing the same ancestral area, will – irrespectively of their species – have (relatively) similar plastomes. In addition, the closer a modern-day *Fagus* population (or species) is to the ‘last common mother’, the less evolved will be its plastome.

**Figure 3.**
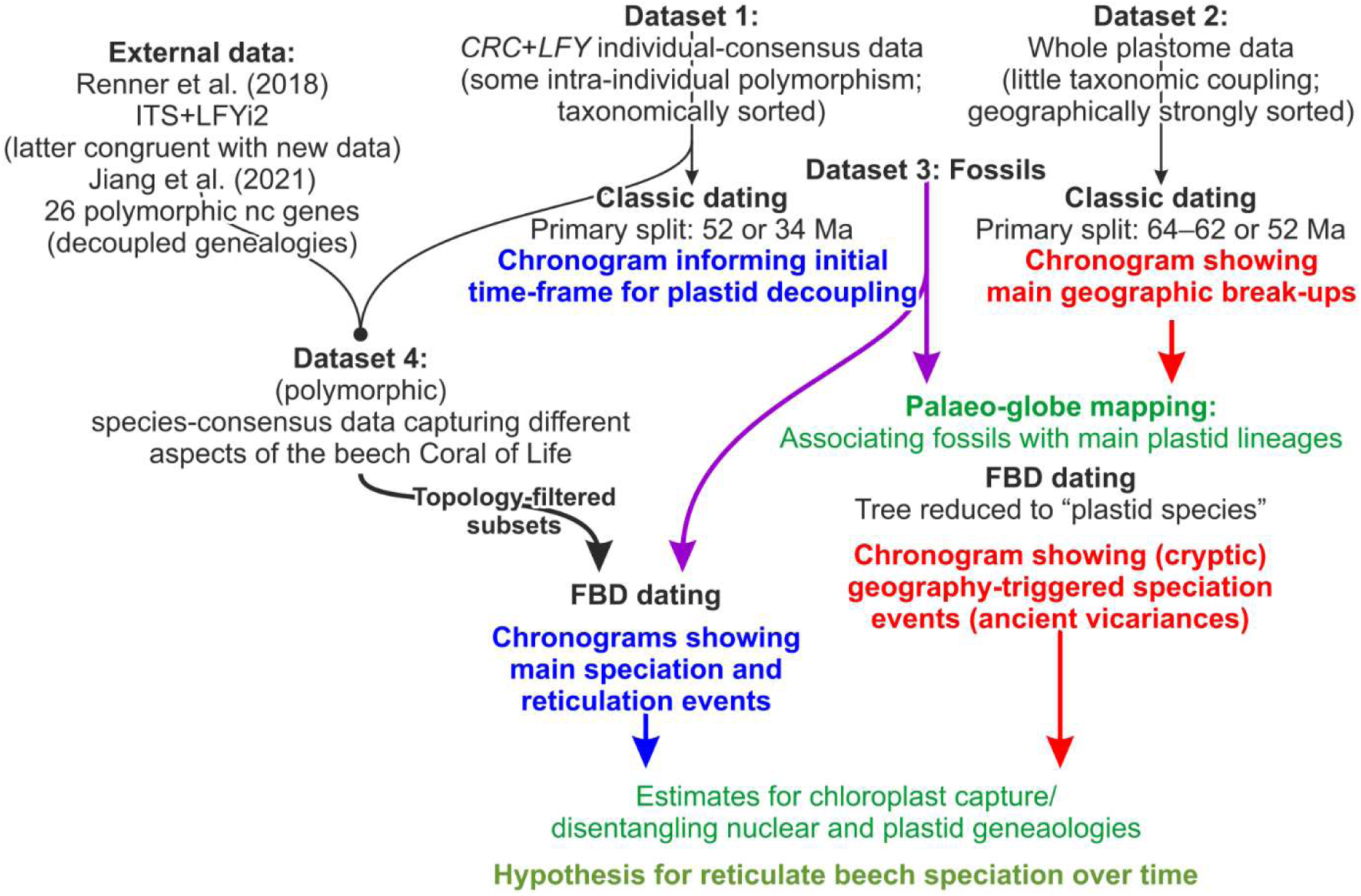
| Dating experiments flowchart. Datasets and approaches in black text; blue text denotes dated nuclear trees; red text denotes dated plastid trees (see **SupplM&M.pdf**, *section 5*, for details). Green text denotes concluding hypotheses and results (detailed in **SupplR&D.pdf**, *section 3*).

**Figure 4.**
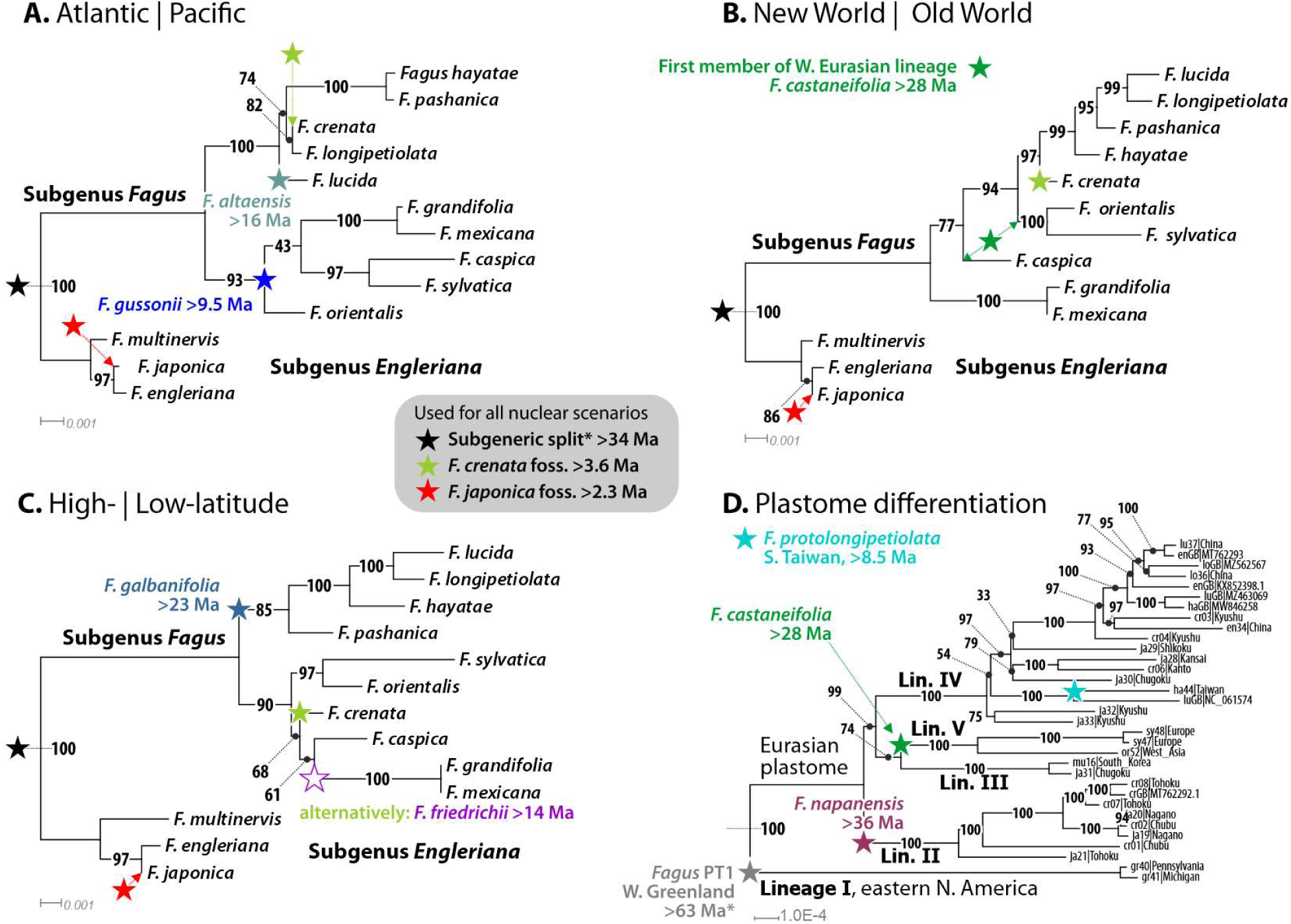
**| Age priors used for establishing minimum estimates for final speciation events and last contacts between species lineages**. **A. Atlantic | Pacific scenario**—primary split within *Fagus* subgenus *Fagus* between Atlantic-(W. Eurasian, E. North American) and Pacific-bound (E. Asian) species. **B**. **New World | Old World** scenario—between E. North American spp. and Eurasian spp. **C. High-| Low-latitude scenario**—between (originally) high-latitude, circum-arctic and (ancient) mid-latitude, continental-montane E. Asian species. **D.** For the plastome tree. The shown topologies are unconstrained, i.e. inferred directly from the congruence-filtered gene sample.

We thus used four principal datasets (**Fig. 3**): *(i)* the concatenated nuclear *CRC*-LFYi2 individual (strict) consensus data (gaps treated as missing data) and *(ii)* complete chloroplast genome (plastome) data sets pruned to the 20 individuals for which *CRC* and LFYi2 were sequenced and representing all major plastid haplotypes and *(iii)* a set of 101 fossil occurrences to generate meta-calibrated FBD-dated trees (**SupplFossilTable.xlsx**, sheets *cpFBD*, *ncFBD*) and select (conservative) age priors for node dating (**Fig. 4**). In addition (*iv*), we compiled a species-level strict consensus data set combining our new *CRC-*LFYi2 data with nuclear data from earlier studies (Denk et al. 2005, Renner et al. 2016, Cardoni et al. 2022, Jiang et al. 2022). For further details on used data sets, rationale and dating experiments see **SupplM&M.pdf**, *section 5*, and **SupplR&D.pdf**, *section 3*.

## Results

### Chloroplast genome size and overall genetic diversity patterns

Circular genomes were obtained for all but four *F. crenata* samples whereby 2–3 chloroplast contigs were assembled into a whole chloroplast genome using the *F. crenata* reference in GENEIOUS. The MAFFT alignment of all 82 whole chloroplast genomes was 159,840 bp in length and 132,787 bp when excluding the IRa region and the *ycf1* pseudogene.

A total of 57 distinct chloroplast haplotypes were observed when solely SNPs were considered or 77 when indels and SNPs were used. Six haplotypes were found in more than one sample of the same species, up to c. 955 km air-distance apart in case of Italian and Greek *F. sylvatica* (**Table 1**). Overall, *Fagus* chloroplast genomes are highly conserved (96.8% invariable sites; 1,638 SNPs of which 1,172 are parsimony informative), with an average of 1.26 variable sites per 100 bp and generally length-homogenous. The two nuclear markers were by a factor of 10 more variable with 169 (LFYi2) and 177 (*CRC*) SNPs, respectively, and on average 11.2 (*CRC*) and 14.8 (LFYi2) variable sites per 100 bp (**Table 2**).

**Table 1.**
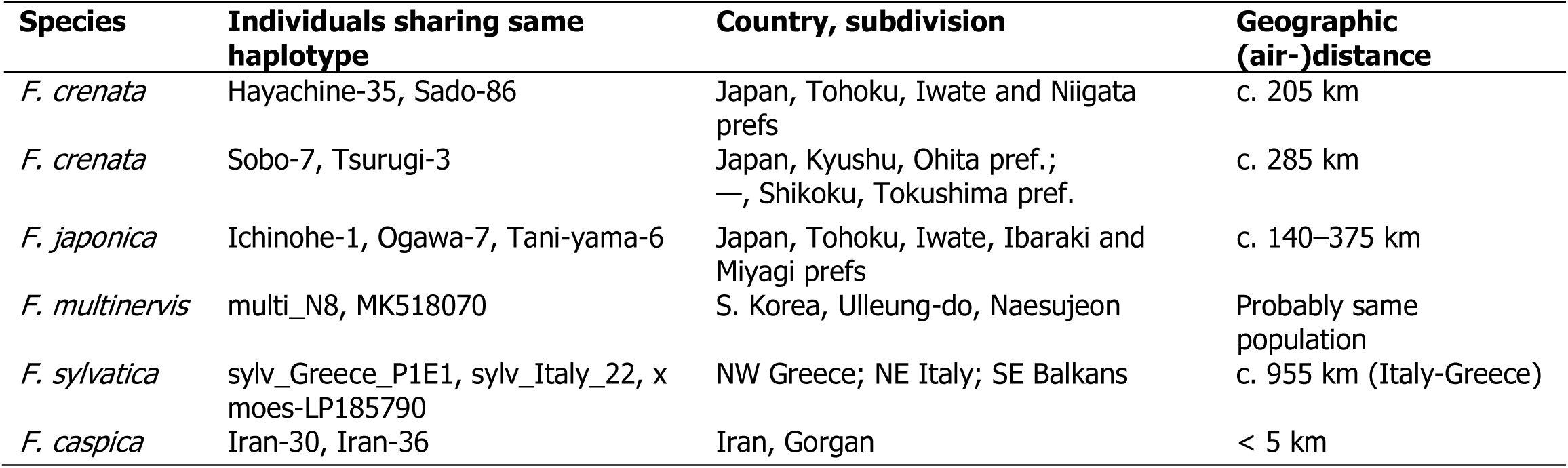
| List of samples that shared the same whole chloroplast-based haplotype. , their geographic origins and air distance between them.

**Table 2.** | Summary of single nucleotide polymorphism (SNP) and length-polymorphism (InDel) sites per data set. Abbrev.: FDS, number of fully determined sites, i.e. ungapped and sites excluding missing data; InvS; invariable sites; PIS, parsimony-informative sites.

| **Data set** | **No. Taxa** | **No. bp** | **No. FDS** | **No. InvS** | **SNPs** | **SNPs/100bp** | **Nucleotide diversity** | **No. PIS** | **Total number of InDel sites** | **Av. InDel length** | **InDel Diversity per site** |
| --- | --- | --- | --- | --- | --- | --- | --- | --- | --- | --- | --- |
| cpDNA | 77 | 132787 | 130166 | 128528 | 1638 | 1.26 | 0.00176 | 1172 | 2621 | 4.84 | 0.00058 |
| CRC | 103 | 1702 | 1580 | 1306 | 177 | 11.20 | 0.01196 | 97 | 122 | 2.22 | 0.00229 |
| LFYi2 | 231 | 1297 | 1142 | 821 | 169 | 14.80 | 0.01942 | 152 | 148 | 3.64 | 0.00257 |

We found major differences between the plastid and nuclear data with regard to the coherence of species, and subgenera in the East Asian species. Maximum intra-species plastid variation is about 3-times the minimum inter-species divergence. The subgeneric affiliation of a species is irrelevant: plastomes of *F. engleriana* Seemen ex Diels are most similar to those of certain *F. crenata* and *F. lucida* Rehder & E.H.Wilson individuals; plastomes of *F. japonica* and *F. crenata* can be more similar to each other than to their siblings of the same subgenus. In the nuclear data, the species of the two subgenera are clearly distinct, and, in the case of *Fagus* subgenus *Fagus*, intra-specific variation is generally lower than inter-specific divergence (**Table 3**). Only the geographical extremities of the genus’ range are genetically highly-coherent in both nuclear and plastid data: European *F. sylvatica* (24 complete chloroplast genomes available, ≤ 23 SNPs difference) and eastern North American *F. grandifolia* (4 plastomes, with 5–13 SNPs difference).

**Table 3.** | Coherence in phylogenetic signal across the three datasets in terms of intra- and inter-taxon diversity based on SNPs. Taxa in bold (green backgrounds) show coherent patterns where intra-taxonomic diversity < inter-taxonomic divergence, while non-bold (orange backgrounds) are not coherent, that is, intra-taxonomic diversity >> inter-taxonomic divergence. The closest tip (species, individual, accession, # = clone) for each taxon is also shown.

| Taxon | Max. intra-taxon diversity |  |  | Min. inter-taxon divergence |  |  | Closest taxon/ individual |  |  |
| --- | --- | --- | --- | --- | --- | --- | --- | --- | --- |
|  | cpDNA | CRC | LFY | cpDNA | CRC | LFY | cpDNA | CRC | LFY |
| *Fagus* subgenus *Englerianae* | 311 | 29 | 27 | 15 | 37 | 92 | *F. lucida* (lu37) | *F. longipetiolata* (lo36#5) | *F. hayatae* (ha43#6);<br>*F. longipetiolata* (KU565425) |
| *F. engleriana* | 99 | 11 | 6 | 15 | 3 | 1 | *F. lucida* (lu37) | *F. japonica* (ja32#4) | *F. japonica* (ja21#6; KU565443;<br>KU565444) |
| ***F. multinervis*** | **5** | **11** | **7** | **18** | **9** | **4** | *F. japonica* (ja31) | *F. japonica* (ja26#2) | *F. japonica* (ja21#6; ja26#5;<br>KU565443; KU565444) |
| *F. japonica* | 287 | 26 | 27 | 18 | 3 | 1 | *F. multinervis* (MN894556) | *F. engleriana* (en34#8) | *F. engleriana* (en34#1; en34#2;<br>en34#3; KT352991; KU565441) |
| *F.* subgenus *Fagus* | 485 | 92 | 71 | 15 | 37 | 92 | *F. engleriana* (MT762293) | *F. japonica* (ja21#10) | *F. japonica* (KU565459;<br>KU565460) |
| ***F. grandifolia*** | **12** | **59** | **22** | **373** | **9** | **29** | *F. japonica* (ja21; ja24; ja26) | *F. mexicana* (EU189865) | *F. caspica* (or52#2; or52#4; or52#6);<br>*F. hayatae* (ha43#6);<br>*F. longipetiolata* (KU565425) |
| *F. mexicana* | N/D | N/A | N/D | N/D | N/A | N/D | – | – | – |
| *F. crenata* | 322 | 36 | 40 | 73 | 17 | 28 | *F. engleriana* (KX852398) | *F. longipetiolata* (lo36#7) | *F. longipetiolata* (KU565421) |
| *F. longipetiolata* | N/A | 47 | 30 | 37 | 16 | 9 | *F. engleriana* (MT762293) | *F. hayatae* (ha43#2;<br>ha43#3; ha43#4) | *F. hayatae* (ha43#4) |
| *F. lucida* | N/A | N/D | 15 | 15 | N/D | 14 | *F. engleriana* (MT762293) | – | *F. hayatae* (ha43#4; ha43#6);<br>*F. longipetiolata* (KU565424) |
| ***F. hayatae*** | **2** | **10** | **8** | **192** | **16** | **9** | *F. japonica* (ja33) | *F. longipetiolata* (lo36#8) | *F. longipetiolata* (KU565425) |
| *F. pashanica* | N/A | N/D | N/A | N/A | N/D | N/A | – | – | – |
| ***F. caspica*** | **0** | **25** | **13** | **171** | **21** | **29** | *F. sylvatica* (sy48) | *F. sylvatica* (sy49#27; sy49#28) | *F. grandifolia* (gr38#3) |
| *F. orientalis* | N/D | N/D | N/A | N/D | N/D |  | – | – | – |
| ***F. sylvatica*** | **23** | **10** | **24** | **171** | **21** | **44** | *F. caspica* (or52) | *F. caspica* (or52#4; or52#20;<br>or52#21) | *F. crenata* (KU565411; cr08#4;<br>cr08#5; cr08#6) |

Note that shared haplotypes of *F. crenata* (individual cr02) viz *F. japonica* ja20 (1 SNP difference)/ ja19 (14 SNPs) are not considered for pairwise comparison. N/A indicates calculation was not possible because the taxon was represented by only one individual, while N/D indicates that no data was available. No data available for *F. hohenackeriana*.

The plastome differentiation shows a strong geographic correlation (**Fig. 5**), with the plastomes of *F. grandifolia* being most distinct from all others (357–440 SNPs difference vs. max. 315 SNPs across Eurasia). When using the intra-species diversity observed in *F. sylvatica* as guideline for species-level haplotype groups, the newly assembled and published plastomes would comprise at least 23 species-level plastid types (SLPT) with highest diversity in the Japanese Archipelago (13 SLPTs vs. ∼6 in China and Taiwan, 2 in Western Eurasia and 1 in North America, **Table 4**).

**Figure 5.**
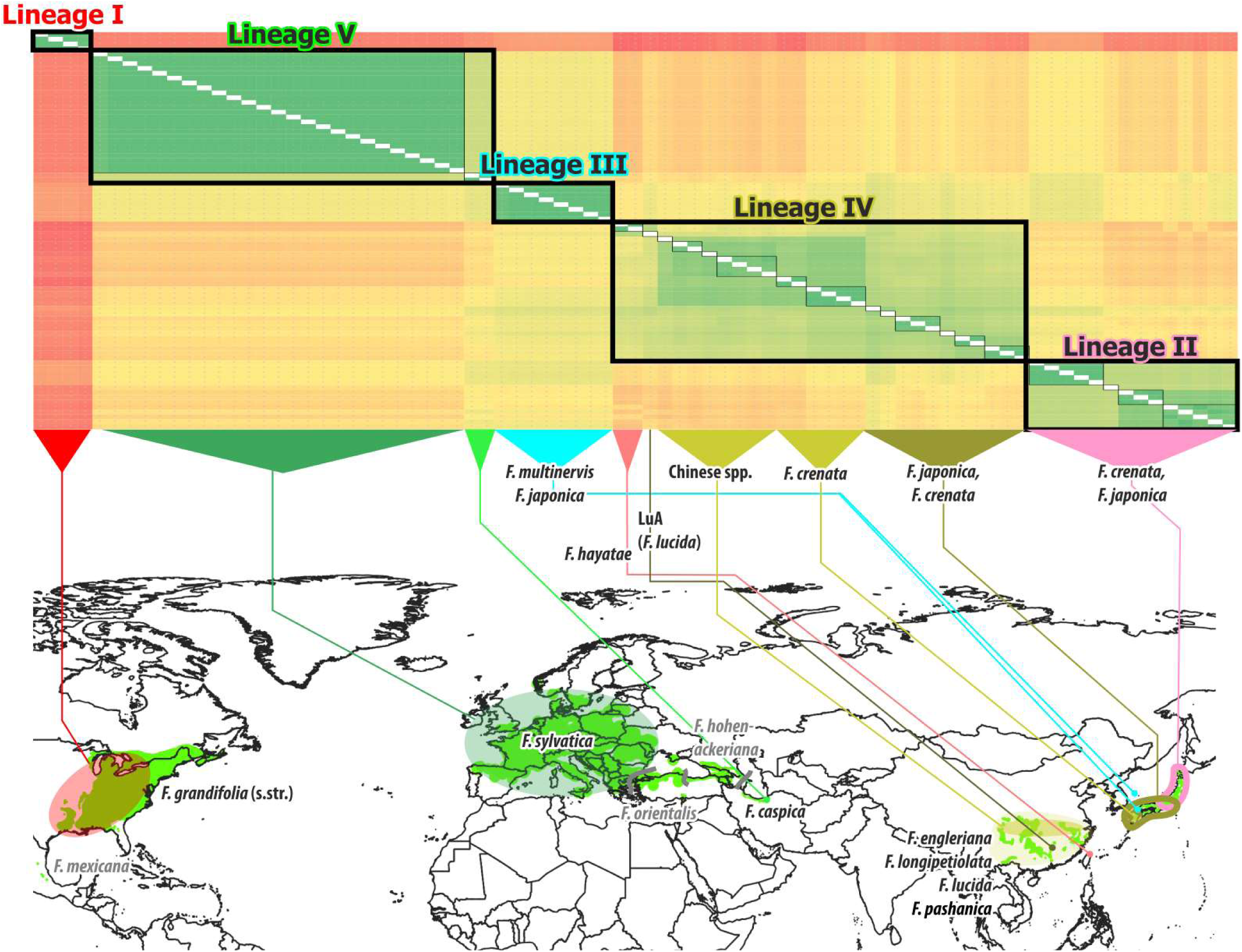
| SNP dissimilarity matrix (heat-map) put in a geographic context. Colour gradient in the heat map (top) ranges from ≤ 4 SNPs difference (dark green) to >400 SNPs difference (dark red). Thick black boxes: main plastome lineages, thin black boxes: ‘species-level plastid types’ (see text). No complete plastome data are available for *F. orientalis*, the Caucasian *F. hohenackeriana* and *F. mexicana.* The exact provenance of most Chinese plastomes (‘Chinese spp.’ group, part of the Lin. IV core clade) is unknown/not reported in the original papers (cf. **SupplGenet-ics_cpDNA.xlsx**, sheet *New and used plastomes*).

**Table 4.** | Putative past speciation events inferred from chloroplast genomes diversity patterns. #CCP = Number of complete chloroplast genomes (= sequenced individuals); #SLPT = Number of species-level plastid types using plastome differentiation within *F. sylvatica* as baseline and representative for a widespread but genetically coherent (regarding both its nucleome and plastome) species (see also **SupplGenetics_cpDNA.xlsx**, sheet *PlstmDissim*). Abbrev.: *cr* = *F. crenata*, *ja* = *F. japonica*

| Region<sup>a</sup> | Plastid Lineage | Species, population | Taxonomic status | #CCP (new) | #SLPT | Max. intra-SLPT | Min. inter-SLPT | Vs. *F. grandifolia* |
| --- | --- | --- | --- | --- | --- | --- | --- | --- |
| WEA | V | *F. sylvatica* | Exclusive | 24+1 (6) | 1 | 23 | 170 | 400–409 |
| WEA | V | *F. caspica* | Exclusive | 2 (all) | 1 | 0 | 170 | 384–390 |
| NEA | II | NE *cr* + *ja* | Shared | 14 (12) | 4 | 3–50 | 62–146 | 357–413 |
| NEA | III | *mu* + *ja* | Exclusive | 8 (4) | 1 | 20 | 193 | 360–372 |
| NEA | IV | SW *cr* + *ja* | Shared | 17 (15) | 8 | 1–26 | 68–116 | 369–434 |
| SEA | IV | *F. hayatae* | Exclusive | 2 (all) | 1 | 2 | 135 | 432–440 |
| SEA | IV | Chinese spp.<sup>b</sup> | Shared | 9 (2) | 5(+)<sup>b</sup> | 30–50 | 55–135 | 407–434 |
| ENA | I | *F. grandifolia* | Exclusive | 4 (all) | 1 | 13 | 357 | N/A |

<sup>a</sup> Denotes the following regions: Western Eurasia (WEA), North-East Asia (NEA), southern East Asia (SEA) and eastern North America (ENA).
<sup>b</sup> *F. engleriana* of *Fagus* subg. *Englerianae* with 3 SLPT, one shared with Chinese spp. of *F.* subg. *Fagus*; *F. longipetiolata* (1 SLPT), *F. lucida* (3 SLPT) and *F. pashanica* (single accession) of *F.* subg. *Fagus*. More broadly sampled data may increase the total number of (closely related) SLPTs or the number of shared SLPTs. Only one SLPT (LuA) is substantially distinct; all others from the core clade of Lineage IV plastomes (cf. **Figs 5, 6; Fig. SR2-2** in SupplR&D.pdf)

### Decoupled chloroplast and nuclear genealogies

For the chloroplast data set, distance-based (neighbour net) and ML phylogenetic analyses identified five main lineages (**Figs 5, 6**; Fig. SR2-2 in SupplR&D.pdf). Most haplotypes of the two endemic Japanese species, *F. crenata* and *F. japonica*, were scattered across two of the five main lineages. One, ‘Lineage II’, is exclusive to *F. crenata* and *F. japonica*; the other, ‘Lineage IV’, is shared with the remaining East Asian species. The haplotypes of *F. multinervis,* endemic to Ulleung Island, South Korea and a single *F. japonica* sample closest to Ulleung Island, are equally distinct to those of the Western Eurasian and East Asian species forming a third continental Eurasian lineage, high-latitude ‘Lineage III’ and is either sister to the Western Eurasian haplotypes, ‘Lineage V’ (ML-BS ≤ 87), or to Lineage IV (ML-BS ≤ 99, depending on the used model and partitioning scheme; **Table SR2-1** in SupplR&D.pdf).

Assuming that the first plastome divergence occurred between the New and Old World (see following section), the second divergence was between ‘Lineage II’ and the dominant continental Eurasian plastid lineages: ‘Lineage IV’ in East Asia and ‘Lineage V’ in Western Eurasia, with subsequent radiation of the latter two lineages; in case of Lineage IV two major bursts, reflected by the large Lineage IV fan and the high-coherent small subfan including two *F. crenata* SLPTs and SLPTs of (today’s) southern East Asian species (**Fig. 6**, preceding page).

**Figure 6.**
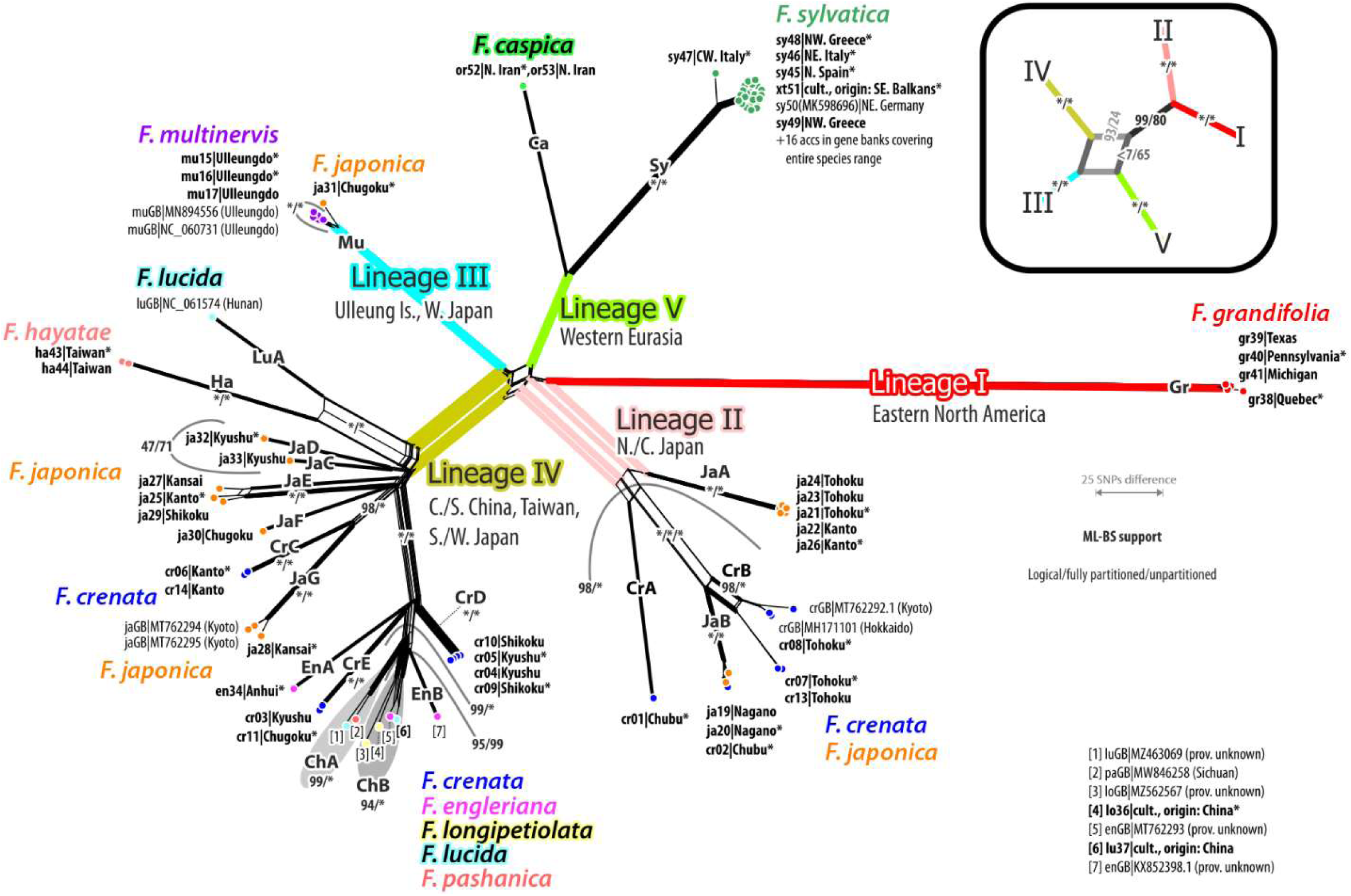
| Neighbour-net of the whole chloroplast genome data set based on uncorrected *p*-distances (excluding indels) New samples (sequenced individuals) in bold font; * included as tip in dating analyses. Species-level plastid types (SLPT) determined based on the divergence within *F. sylvatica* (V-**Sy**) are indicated. Edge labels give ranges of non-parametric bootstrap support (* = 100) for the corresponding branch under maximum likelihood (ML)(see also **SupplGenetics_cpDNA.xlsx**, sheet *cp splits*; for a correspondingly annotated ML tree see **Fig. SR2-2** in SupplR&D.pdf).

In ‘Lineage V’, the European *Fagus* (*F. sylvatica*, SLPT V-Sy) are clearly distinct from their Iranian sisters (*F. caspica* Denk & G.W.Grimm; V-Ca; **Figs 5, 6**) reflecting the geographic distance between the sampled populations (min. 2800 km air-distance; cf. **Fig. 1**).

In contrast, the newly added nuclear sequence regions showed high taxonomic coherence (cloned sequences, individual consensus sequences). The most prominent split is between the two sub-genera, *Fagus* subgenus *Englerianae* including *F. japonica* and *Fagus* subgenus *Fagus* including *F. crenata*, irrespective of geography and plastid lineage (**Fig. 7**). *CRC* data produced higher support along the backbone (BS ≥ 98 for all but one branch, when using strict individual consensus sequences); LFYi2 data showed a higher capacity to sort the leaves. For instance, the *CRC* data did not resolve relationships between the East Asian species (no data available for *F. lucida* and *F. pashanica* C.C.Yang), while LFYi2 resolved a *F. crenata* clade with moderate branch support (BS = 67), high character support (long root branch) and outside the high-supported (BS = 94) clade comprising the other three East Asian species of *Fagus* subgenus *Fagus*. *CRC* and LFYi2 support partially different inter-species relationships (**Fig. 7**); *CRC* provides strong support for an ‘Euamerican’ clade, a sister relationship between *F. grandifolia-F. mexicana* and *F. caspica-F. sylvatica* (BS = 100 when using species consensus data) while LFYi2 prefers the alternative of a New World | Old World split within *Fagus* subgenus *Fagus* (BS = 39, alternative BS < 10, when using species consensi). Within the East Asian species complex, LFYi2 supports a closer relationship between *F. hayatae-F. pashanica* and *F. longipetiolata* or *F. lucida* (BS = 50 and 49) but rejects a sister relationship between the latter (BS < 1 when using species consensus data). The lack of resolution in the individual consensus *CRC* phylogram (**Fig. 7**) relates to a dimorphism (*CRC* heterozygosity) in *F. longipetiolata,* including one allelic lineage (‘Lo1’ in **Fig. SR1-3** in SupplR&D.pdf) shared with *F. crenata* (‘Cr0’ and ‘Cr1’ alleles), while the other (‘Lo2’) represents a sister type of the *F. hayatae CRC*.

**Figure 7.**
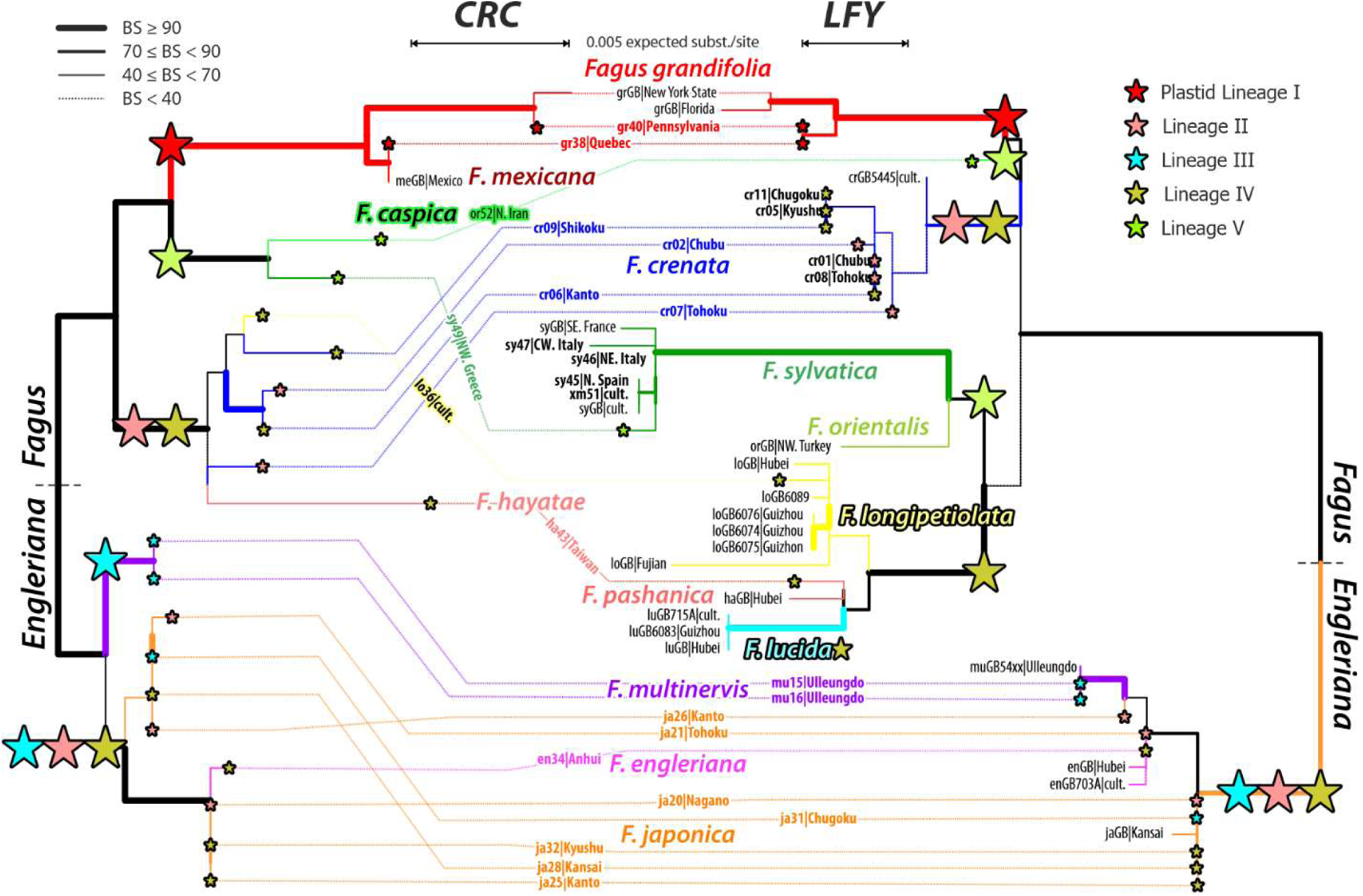
| Comparison of *CRC* and LFYi2 Maximum Likelihood phylogenetic trees (strict individual consensi) estimated from a subset of the samples used for the whole chloroplast genome analyses (bold font) plus individuals covered by earlier studies (normal font). The chloroplast-based lineages of each tip and major branch within the trees are indicated by stars. Bootstrap support values are indicated by the thickness of the branches.

However, individual nuclear gene regions – in addition to *CRC* and LFYi2, data are available for the ITS region and 28 low-copy nuclear genes – support or reject different topological scenarios (**Table 5**; complete information on locus-wise support from species consensus sequences for topological alternatives captured in the nuclear data is included in **SupplDating.xlsx**, sheet *pre-FBD topology test*, and **SupplR&D.pdf**, *section 1*).

**Table 5.** | Topological aspects of the *Fagus* species network and their support from nuclear species consensus data. #T = total number of available markers, #S = number of (strongly) supporting nuclear markers, iBS = individual BS support from supporting markers, #R = number of rejecting or partly conflicting nuclear markers, #U = number of uninformative or indifferent nuclear markers.

| Aspect | #T | #S | iBS | #R | #U |
| --- | --- | --- | --- | --- | --- |
| Primary subgeneric split (*F. subg. Englerianae* \| *F. subg. Fagus*) | 31 | 27 | ≥ 57<sup>a</sup> | 2<sup>b</sup> | 2 |
| New World \| Old World split in *F. subg. Fagus* | 31 | 4 | 39–99.9 | 12 | 8 |
| Euamerican clade joining N. American + W. Eurasian spp. | 31 | 4 | 65/96–100 | 10 | 11 |
| East Asian clade within *Fagus* subgenus *Fagus* | 31 | 4 | ≥ 92 | 9 | 16 |
| High-latitude lineage, *F. crenata* closest to W. Eurasian spp. | 31 | 1(7)<sup>c</sup> | 41 (20–98) | 10 | 10 |
| *F. crenata* + *F. longipetiolata* (CRC-preferred) | 31 | 5 | ≥ 59 | 12 | 16 |
| *F. crenata* + *F. lucida* (ITS-preferred) | 30 | 2 | 30 and 95 | 12 | 17 |
| *F. crenata* + *F. hayatae-pashanica* | 31 | 1 | 100 | 16 | 10 |
| Southern E. Asian clade (LFY-preferred)<sup>d</sup> | 30 | 4 | 50–99 | 19 | 7 |
| *F. longipetiolata* + *F. hayatae-pashanica* | 31 | 1 | 50 | 14 | 15 |
| *F. longipetiolata* + *F. lucida* | 30 | 4 | 76–99 | 7 | 12 |

<sup>a</sup> BS ≥ 83 except for two markers (P38, P49 of Jiang et al. 2022)
<sup>b</sup> Signal artefact in Jiang et al.'s (2022) markers P28 and P34 (cf. Cardoni et al. 2022)
<sup>c</sup> Little direct support for a high-latitude clade but support for (semi-)compatible bipartitions linking (part of) the western Eurasian species to *F. crenata*, *F. lucida* and the North American species.
<sup>d</sup> Including the Chinese and Taiwanese spp. of *F. subg. Fagus*: *F. hayatae*, *F. longipetiolata*, *F. lucida* and *F. pashanica*

### Spatio-temporal framework for Fagus and plastome evolution

The oldest *Fagus* fossils are dispersed pollen from western Greenland (Danian, 64–62 Ma; 60° N paleo-latitude; northern North Atlantic region, NAT); contemporaneous or younger pollen (66–52 Ma) occur in north-eastern Siberia (Beringia, BER; ≥ 70° N; **Fig. 8a**; for details see **SupplR&D**, *section 4*). About 10–15 Ma later, *Fagus* thrived in western North America (WNA), Kamchatka (BER), northern Siberia (northern Asia: NAS) and the Russian Far East (North-East Asia, NEA) while maintaining stable populations in western Greenland and on Axel Heiberg Island (NAT; **Fig. 8a**), i.e. covering about max. 8,000 km air-distance and an area of at least 10 Mkm². The distinctness of the New World plastomes (Lineage I) is the modern legacy of the geographic remoteness of the *first* common mothers of today’s eastern North American (ENA) *Fagus*, with FBD dating modelling a latest Cretaceous / Paleocene stem (66 [73–64] Ma) and crown age (∼63 Ma; directly informed by western Greenland pollen) for plastome Lineage I (details provided in **SupplDating.xlsx**, sheet *final dating*; **SupplR&D.pdf**, *section 3.3*) The first global radiation reflected in the plastome signatures predates the manifestation (morphology and nuclear genes) of the two main modern-day lineages, *Fagus* subgenus *Englerianae* and *F.* subgenus *Fagus,* by up to ∼25 Ma (**Fig. 9:[0–1]**; **SupplR&D.pdf**, *section 3.3.2*).

**Figure 8.**
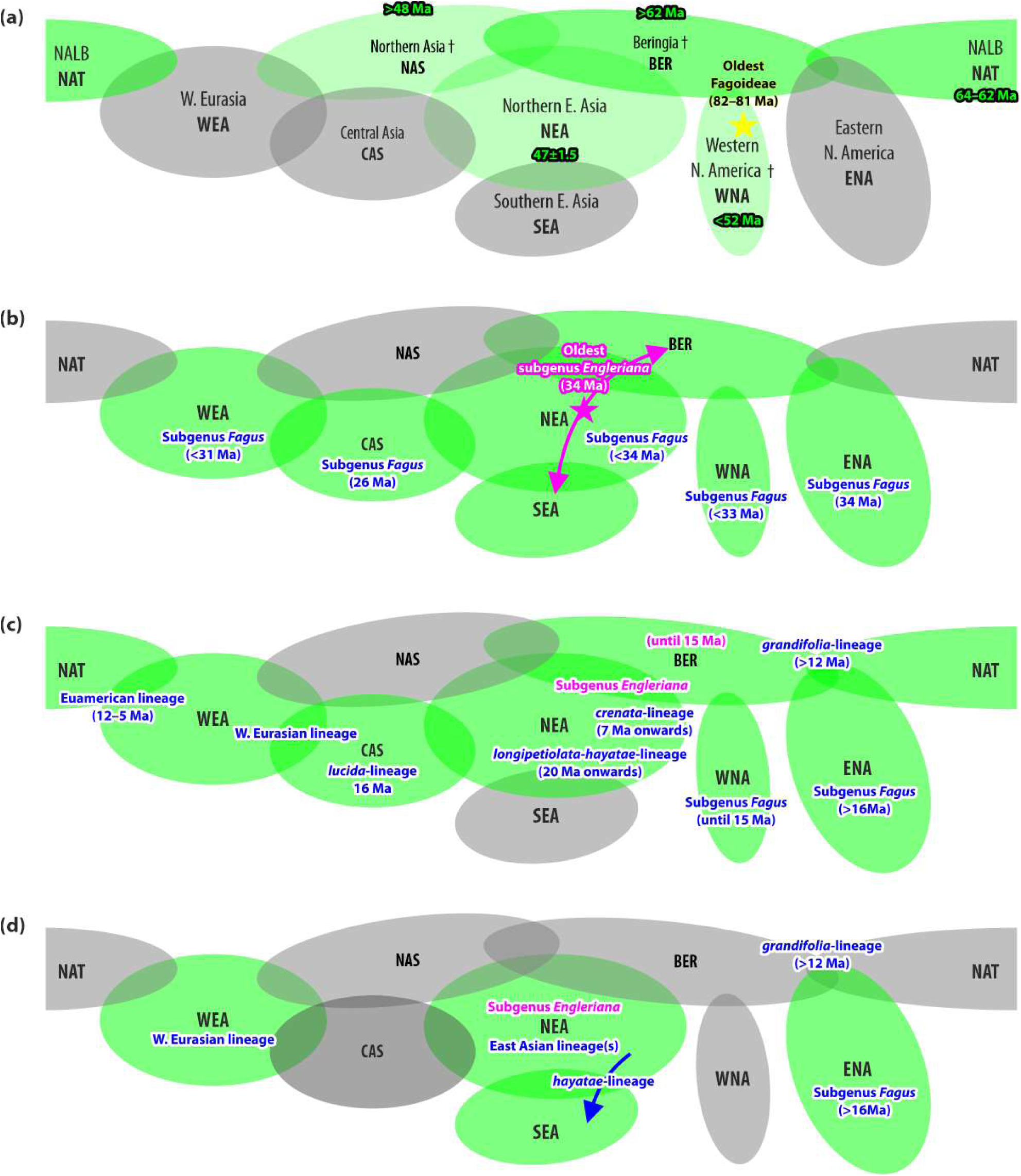
| Spatio-temporal framework for *Fagus* and *Fagus* plastome evolution based on the fossil record of beech. revisited for this study. Grey shaded areas not occupied by *Fagus* fossils at the time, green coloured areas host *Fagus* fossils. Biogeographic regions, which in their extent change over time but remained stable regarding their relative position to each other (cf. **SupplR&D.pdf**, *section 4*), abbreviated by three letters. (a) Situation in the (pre-)Eocene (>34 Ma); (b) at the Oligocene-Eocene boundary (∼34 Ma) and onwards; (c) during the Miocene (23–5 Ma), the ‘high-time’ of beech evolution; and (d) after the Miocene.

**Figure 9.**
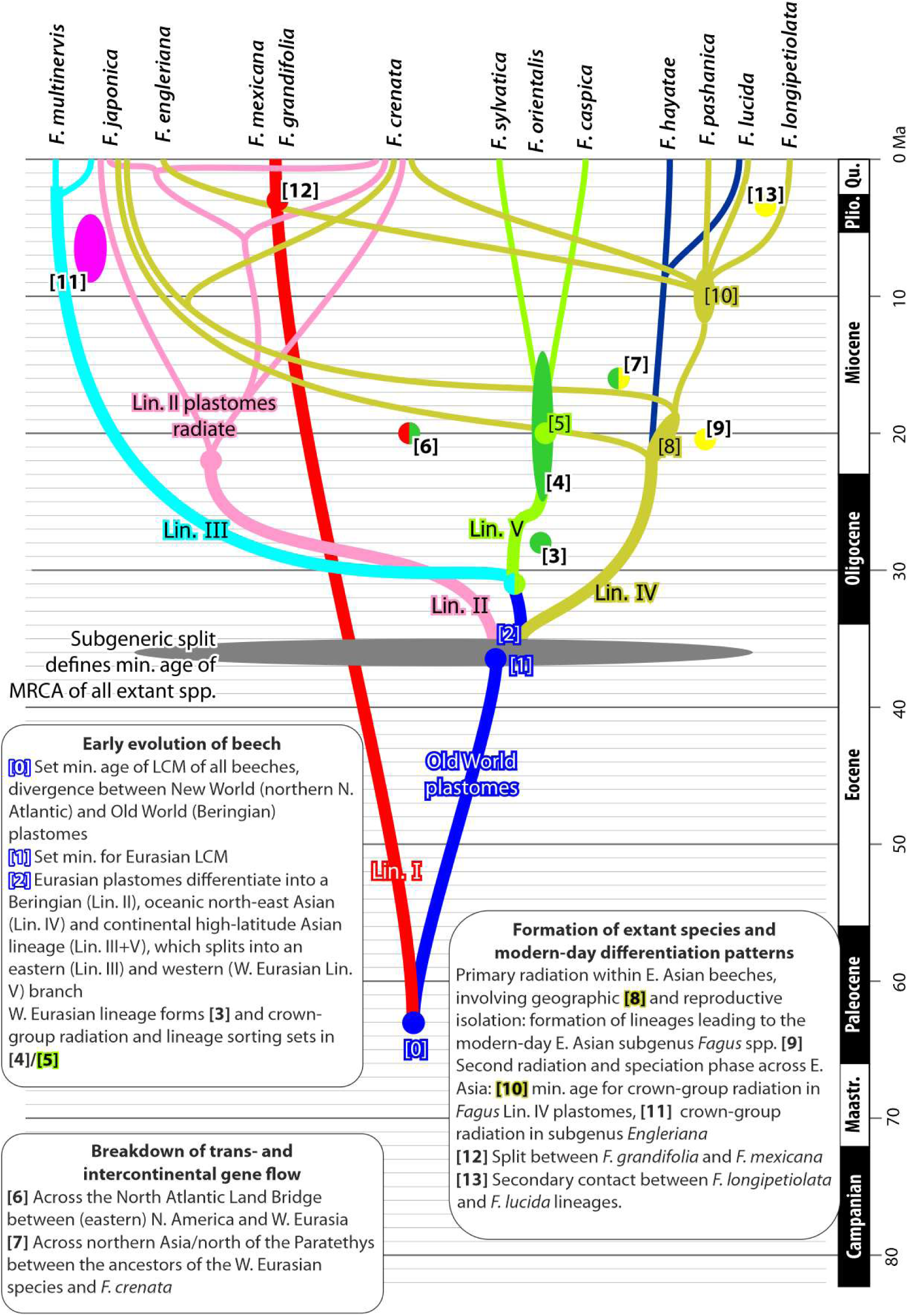
| Summary of the timing of divergence of major chloroplast lineages and their spread to the gene pool of modern species via lineage sorting and ancient reticulation. Major events over time are described in the text boxes.

During the late Eocene to Oligocene cooling, *Fagus* moved into lower latitudes (**Fig. 8b**). The Oligocene-Eocene boundary is marked by the phenotypical sorting into the two modern subgeneric lineages; the 34 Ma Kraskino paleoflora (NEA; Pavlyutkin et al. 2014) includes fossils that can be unambiguously assigned to either *Fagus* subgenus *Englerianae* or *F.* subgenus *Fagus*. Thus, the last common ancestor of all modern-day *Fagus* species and the onset of earliest nuclear divergences can be assumed to be older than 34 Ma (**Fig. 9**). The Eurasian plastome pool had started to diversify at the Eocene-Oligocene boundary as well (**Fig. 9 [1–2]**), with FBD dating estimating a Lineage II stem age (modelled as equalling the Eurasian plastid MRCA) of 44 (49– 40) Ma. Today, Lineage II plastomes are restricted to the central and northern part of the Japanese Archipelago and comprise four species-level plastid types (SLPTs; **Fig. 6; Fig. SR2-6** in SupplR&D.pdf).

The differentiation into three equally distinct continental Eurasian plastome lineages III, IV and V before 31 Ma (**Fig. 9; Table SR3-1** in SupplR&D.pdf) resulted from large-scale range expansion followed by fragmentation in Eurasian *Fagus* (**Fig. 8b**). With the closure of the Turgai Strait (in modern-day Kazakhstan), *Fagus* migrated into Western Eurasia (WEA) while maintaining a large distribution area covering northern (NAS) and central Asia (CAS). North-East Asian and Beringian populations were up to 1,600 km apart; while the fossil-species †*F. castaneifolia* spanned at least 5,000 km across the continent (CAS to WEA). *Fagus* subgenus *Englerianae* migrated into lower latitudes (modern-day Guizhou Province, 26° N paleolatitude). The combination of geographic distance and emerging barriers in Central Asia (Qinghai-Tibet Plateau; continental steppes; cf. **SupplR&D**, *section 4*) led to a triple vicariance (formation of lineages III, IV and V) followed by further and rapid differentiation of the plastid gene pools. Two sublineages of Lineage V (European *F. sylvatica* vs. Hyrcanian *F. caspica*; **Fig. 9 [5]**) diverged 3–9 Ma before the radiation of the East Asian Lineage II and IV plastomes during the Early Miocene (**Fig. 9**; SupplR&D.pdf, *section 3.3.4*). Simultaneously or slightly earlier, the Western Eurasian *Fagus* lineage diverged as a nuclear-genetically and morphologically distinct coherent entity (**Fig. 9 [3,4]; Table 6; SupplR&D.pdf**, *section 3.3.3*).

**Table 6.** | Stem and crown age estimates (in Ma) for Western Eurasian *Fagus* lineage based on node dating (ND) and a fossilized birth-death (FBD) dating framework. Given are the medians, with 95% HPD intervals in brackets (see SupplDating,xlsx, sheet ‘final dating’, for the full results).

| Data, topology | ND |  | FBD |  |
| --- | --- | --- | --- | --- |
|  | Stem | Crown<sup>a</sup> | Stem | Crown<sup>b</sup> |
| Nuclear, Atlantic \| Pacific | 20 (28–10) | 14 (22–6) | 29 (38–20) | 25 (33–18)<sup>a</sup> |
| —, High \| Low Latitude | 17 (22–14) | 16 (21–10) | 35 (38–34) | 33 (38–27)<sup>a</sup> |
| —, New \| Old World | 28 (30–28) | 25 (29–16) | 34 (44–31) | 30 (43–20) |
| Plastid (Lin. V) | 31 (33–31) | 20 (26–14) | 34<sup>c</sup> | 31<sup>c</sup> |

<sup>a</sup> Equals the MRCA of extant tips
<sup>b</sup> Modelled crown age, i.e. first modelled divergence within the according clade in the maximum clade credibility tree.
<sup>c</sup> Directly informed by fossil tip within the modelled speciation process.

### By the Pliocene all modern-day species of both subgenera were established as genetically

±coherent entities (**Fig. 8d; Table 7**). In *Fagus* subgenus *Englerianae*, *F. multinervis* diverged ∼5 Ma before the split between the continental *F. engleriana* and *F. japonica* (**Fig. 9:[11]**) and after the crown group radiation in the Lineage IV haplotypes found in *F. engleriana* and the southwestern populations of *F. japonica.* The latter speciation event coincides with the estimated divergence of the Lineage III plastomes on either side of the Sea of Japan (*F. multinervis* ↔ *F. japonica*; see also **Table SR3-9** in SupplR&D). In eastern North America, the southernmost populations (*F. mexicana*) had been largely isolated from those of the Appalachians (*F. grandifolia*; **Fig. 9:[12]**). Intra-continental migration led to the secondary contact and bidirectional gene exchange between the lineages of *F. longipetiolata* and *F. lucida* in southern East Asia (SEA, **Fig. 8c,d; Fig. 9:[13]**; detailed in **Fig. SR2-7** in SupplR&D.pdf); species lineages that go back to at least the Middle Miocene (**Table 7**). The young estimates for the MRCA of *F. longipetiolata* and *F. lucida* in the node-dated High | Low Latitude and New World | Old World scenarios reflect the sharing of evolved and specific nuclear alleles between the two species and accordingly similar, albeit polymorphic, species consensus sequences. The miscellaneous affinity of other nuclear markers of *F. lucida* as well as the three completely sequenced chloroplast genomes indicate that this contact was only the last of repeated reticulation events involved in the formation of modern-day *F. lucida*.

**Table 7.** | Youngest stem age estimates (in Ma) for modern-day species lineages in nuclear dated trees. (across all three tested scenarios, cf. SupplDating.xlsx, sheet ‘final dating’). Given are the medians, with 95% HPD intervals in brackets.

| Species lineage, split | Last contact with | ND | FBD |
| --- | --- | --- | --- |
| *F. crenata* | W. Eurasian or N. American lineage | 14 (16–14) | 35 (38–34) |
| – | Southern East Asian spp. | 19 (22–16) | 26 (35–28) |
| *F. lucida* | *F. longipetiolata* | 3 (7–1) | 30 (34–26) |
| – | Remainder of East Asian spp. | 16 (17–16) | 31 (35–28) |
| *F. hayatae-pashanica* | Other East Asian spp. | 16 (17–16) | 25 (27–24) |
| *F. longipetiolata* | Other than *F. lucida* | 17 (20–16) | 26 (35–28) |
| *F. grandifolia* \| *F. mexicana* | – | 1 ($\leq 3$) | 1 ($\leq 4$) |
| *F. sylvatica* \| *F. caspica* | – | 9 (15–3) | 17 (21–15) |
| *F. caspica* | *F. sylvatica-caspica* | 14 (22–6) | 21 (23–19) |

## Discussion

### Evolutionary signal in Fagus plastome phylogeny

Plastome genealogies reflect (bio)geographic rather than taxonomic patterns (see **Fig. 6** vs **Fig. 7**; Zhou et al. 2022 for Fagaceae in general; Li et al. 2025 for East Asian species of *Quercus* sect. *Cerris*). In *Fagus* and other dominant extra-tropical wind-pollinated trees, plastid differentiation requires geographic isolation and large distribution areas to develop genetic gradients eventually leading to distinct plastid haplotypes. The example of the North American *F. grandifolia* (four near identical plastomes sequenced covering ∼1.4 Mkm²) and the European *F. sylvatica* (24 near identical plastomes, ∼2 Mkm², most of the area recolonised after the Last Glacial Maximum; cf. Magri et al. 2006) demonstrate that in *Fagus* the plastid gene pool shows no significant geographic differentiation within species with large and continuous distributions. Hence, we consider the high regional diversity in *Fagus* in East Asia and particularly in the Japanese Archipelago (**Figs 5, 6**; **Figs SR2-6, SR2-8** in SupplR&D.pdf) a legacy of past speciation processes that involved range expansions, allopatric speciation and (re-)migration. The most compelling result of our study is that two species, today confined to the Japanese islands and belonging to two different, at least 34 Ma old subgenera, carry cousin (*F. crenata* and *F. japonica*) and sister (*F. crenata*) species-level plastid types (SLPTs) of all other extant East Asian *Fagus* species, growing today at far distances (**Fig. 1c,d; Figs 5, 6, 9**). Our data demonstrate that “ghost species”, only persisting as distinct SLPTs and representing ancient, extinct as well as modern species lineages must have come into contact at various times in the geological past especially in North-East Asia, leading to the decoupling in the plastome lineages from taxonomic boundaries and eventually to the hyperdiverse chloroplast genome pool in Japanese species.

### Ghost lineages of Fagus – deep time introgression and hybridisation

The mutual holophyly of *Fagus* subgenus *Englerianae* and *F.* subgenus *Fagus* is long and well established (Denk 2003, Denk et al. 2005, Denk & Grimm 2009, Renner et al. 2016, Cardoni et al. 2022, Jiang et al. 2022, this study). The fossil record and our dating experiments place the last common ancestor of *Fagus* subgenus *Fagus* and the first common ancestors of the North American lineage into the late Eocene/Oligocene (**Figs 8, 9**; see also Renner et al. 2016), long after the main plastid differentiation (**Figs 8–10**). The Lineage I plastome of today’s North American species is the legacy of an extinct Atlantic-Arctic *Fagus* lineage that was assimilated when early members of *Fagus* subgenus *Fagus* from Beringia came into contact with their ancient cousins thriving in the western part of the NALB since the early Paleocene (Denk & Grimm 2009, Grímsson et al. 2016; **Fig. 10a**). Nucleome data do not record this legacy because of intra-specific homogenisation when the eastern North American populations of *Fagus* subgenus *Fagus,* the introgressor of the Arctic-Atlantic *Fagus* lineage, evolved into the modern-day species. Rare exceptions may be puzzling patterns of allelic variation (heterozygosity or paralogy) retained in *F. mexicana* but lost in *F. grandifolia* that reflect ancient links across the Beringian land bridge and the subgeneric boundaries (Jiang et al.’s 2022 locus P49; cf. Cardoni et al. 2022, data S5). The original plastomes of the introgressor (plastomes more similar to those of the Eurasian lineages) were lost when its western North American populations went extinct during the later Miocene (**Fig. 8**).

**Figure 10.**
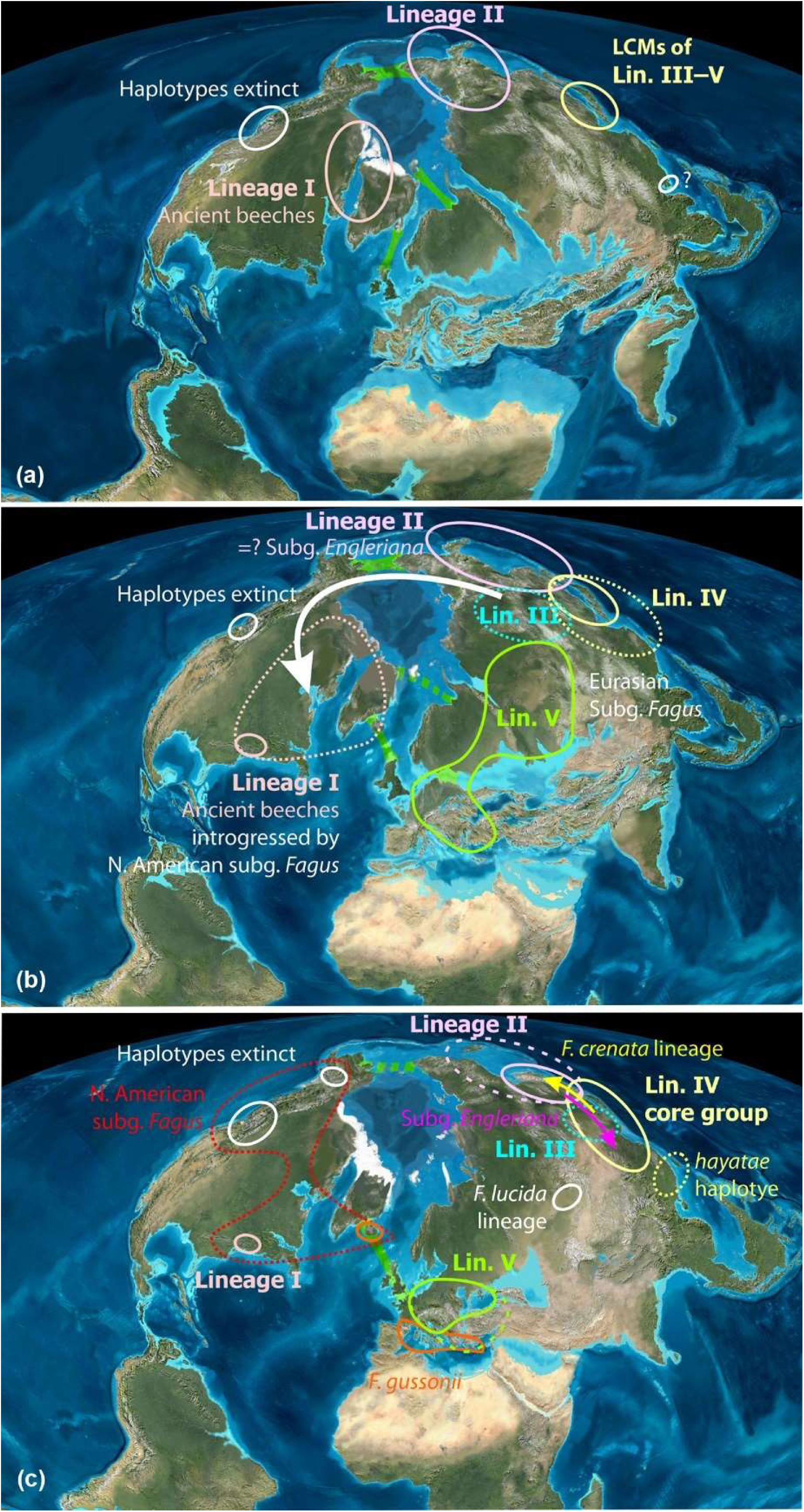
[Following page] **| The fossil record with major plastid lineages distribution overlayed.** The figure summarises our interpretation of all collected direct evidence (molecular differentiation patterns in modern-day beeches; revised total fossil record) and inferences performed for this study as reported in the text (**Figs 5–9; Tables 3–7**) and further detailed in **SupplR&D.pdf**, *sections 3*, *4*. Palaeomaps by R. Blakey for and from Denk & Grimm (2009): (a) (Pre-)Eocene; b) Oligocene; (c) Miocene. A kmz-file for GoogleEarth showing the individual occurrences mapped on the palaoeglobes of Scotese (2014) is included in the OSA. Abbrev.: LCM = last common mother, i.e. the ancestral taxon/population from which (part of) the modern-day beeches inherited their plastome.

The Lineage II plastomes are restricted to the northeastern half of the Japanese Archipelago, roughly corresponding to the ancient North-East Asian-Beringian range of *Fagus* subgenus *Englerianae* (**Fig. 8c,d; Fig. SR2-6** in SupplR&D.pdf). Their distinctness from the remaining Eurasian plastomes makes it unlikely that they represent the original plastomes of *F. crenata* or (one of) its precursor(s). According to our dating estimates, Lineage II plastomes radiated in parallel to those of Lineage IV during the Oligocene and Miocene. Lineage IV is found in all East Asian members of *Fagus* subgenus *Fagus*. Such parallel diversification suggests that the populations carrying those plastid lineages were genetically near-incompatible or did not come into contact; for example, because of being restricted to different bioclimatic niches. Alternatively, the Lineage II plastome may represent one of the donors of a hybrid (allopolyploid, cf. Cardoni et al. 2022) first common ancestor of *Fagus* subgenus *Englerianae.* Species of *Fagus* subgenus *Englerianae* also show a conspicuous polymorphism in the nuclear-encoded multicopy ribosomal spacers, the ITS1 and ITS2 of the 35S rDNA cistron (Denk et al. 2005, Grimm et al. 2007), while, according to the data compiled by Jiang et al. (2022) and for this study, they are genetically homogenised in their single-/low-copy loci (but not all resolve the subgeneric split: P49, F114 of Jiang et al. 2022). This could point to a stabilised allopolyploid (see e.g. Volkov et al. 2007) in which the genetic signature of one donor is largely lost in the single-/low copy coding genes but retained in the multicopy, non-coding genome parts and, in this case, the plastid gene pool. The presence of three different chloroplast lineages (Lin. II, III, and IV) in individuals of *F. japonica* would be the legacy of their allopolyploid ancestor and reflecting its capacity to asymmetrically introgress and replace other *Fagus* at different times and in different places, i.e. in North-East Asia and Beringia during the Oligocene and Miocene, and China from the Miocene onwards.

Finally, the Lineage III plastomes are another possible ghost lineage leaving only unique plastid haplotypes as a legacy, carried by two species that only recently diverged (cf. **Table SR3-9** in SupplR&D.pdf). Lineage III plastomes could be linked to *Fagus* subgenus *Englerianae* and/or represent the other donor of an allopolyploid first common subgenus-*Engleriana-*ancestor. They are genetically the least modified (evolved) plastomes of Eurasia (**Figs 5, 6; Table 3**; exemplified in **Fig. SR2-10** in SupplR&D.pdf; see also **ODA**, file *00_bits.xlsx* tabulating most-divergent plastid gene regions and their mutational patterns). Hence, they did not undergo considerable genetic drift, despite their small extant population size and strongly restricted distribution (37 km^2^ on Ulleung Island, Cai et al. 2021; single population in Japan). Their modern-day area lies in the centre of the Oligocene-Miocene diversity hotspot (Tanai 1995, Pavlyutkin et al. 2014; Momohara & Ito 2023) and the lack of genetic drift could be explained by their ancestors persisting in suitable niches since the Miocene. However, Lineage III is phylogenetically intermediate between the Western Eurasian Lineage V plastomes and the East Asian Lineage IV (**Fig. 6; Figs SR2-2, SR2-3** and **Table SR2-1** in SupplR&D.pdf), both associated with *Fagus* subgenus *Fagus*. In view of the fossil record (**SupplFossilTable.xlsx; Fig. 8**), the tectonic-climatic history of the Northern Hemisphere (Westerhold et al. 2020, Scotese et al. 2021) and our dating experiments, the Lineage III plastomes appear to reflect a vicariant speciation event in the nascent *Fagus* subgenus *Fagus*, which had quickly expanded across northern Eurasia (**Figs 8, 10**). A northern Asian lineage of *Fagus* subgenus *Fagus* would have retreated stepwise to lower latitudes and the Pacific coast during the early Oligocene cooling, where it would have come in contact with the precursor or another donor of *Fagus* subgenus *Englerianae*.

Although there is good evidence for an allopolyploid origin of *Fagus* subgenus *Englerianae* (Cardoni et al. 2022), to further confirm or reject the hypothesis of an allopolyploid origin of *Fagus* subgenus *Fagus* will require an in-depth analysis of the entire nucleome (as done by Ding et al. 2023 for Juglandaceae) and further exploration of intra-genomic nuclear heterozygosity in beeches (data by Jiang et al. 2022, Denk et al. 2024; **SupplR&D.pdf**, *section 1*). Such an in-depth analysis may also reveal genetic legacies of extinct lineages in the nucleome paralleling those in the beech plastome. That asymmetric and bidirectional introgression and hybridisation played a major role in the formation of all modern species of *Fagus* is, however, the only explanation for the puzzling taxonomic mixing of plastomes of the Japanese *Fagus* and intra-genomic nuclear (allelic) variation across the entire genus.

### The Japanese Archipelago - crossroad and storehouse

In order to retain the plastid hyperdiversity (**Figs 5, 6; Tables 3, 4**), the Japanese Archipelago must have played a crucial role in the biogeographic history of *Fagus* as crossroad and storehouse. Three main factors underlie this diversity. Firstly, the paleobotanical record (see *Results* section; **SupplFossilTable.xlsx**, sheet *main list*) shows that Japan and adjacent continental regions have had *Fagus* continually present from the Eocene onwards (**Fig. 10**). Terrestrial volcanism in the Pacific Island arcs since the Late Cretaceous would have provided similar physical conditions as today in the Japanese Archipelago, favouring forest formations with *Fagus* during the Paleogene and Neogene (Bindeman et al. 2010). More than 20 fossil-species of *Fagus* have been reported from the Far East including the Japanese Archipelago (**SupplFossilTable.xlsx**, sheet *main list*), among them the earliest unambiguous records of both subgenera (34 Ma old Kraskino palaeoflora; Pavlyutkin et al. 2014). Secondly, the wide latitudinal range of the Japanese Archipelago leading to multiple past connections with mainland East Asia during lower sea levels would have enabled the migration of divergent lineages of *Fagus* from both cool and warm climates. The first massive invasion wave would have involved the precursors and ancestral members of modern *Fagus* subgenus *Englerianae*. The precursor of *F. crenata* likely invaded after the mid-Miocene when it took over the native Japanese species of *Fagus* subgenus *Fagus*; all of which are now extinct whereas their sister taxa survived in China and Taiwan (Lineage IV plastomes), but also a few ancient *Fagus* subgenus *Englerianae* haplotypes (early diverging Lineage II + IV plastomes; Pliocene: SLPT II-JaB).

Thirdly, the persistent topographic and climatic heterogeneity of the Japanese Archipelago along with the fact that it has been relatively sheltered from extreme climates and landscape changes during glacials (Ono 1984) would have enabled the survival of *Fagus* and its plastid diversity in multiple refugia. Indeed, during the Last Glacial Maximum, temperate tree species including *Fagus* likely persisted across a wide latitude in small refugial populations in coastal areas from southern Kyushu to northeastern Honshu (Tsukada 1982, Worth et al. 2013, Kimura et al. 2014). From our analyses, we deduce that the two parapatric (rarely sympatric) modern-day Japanese species *F. crenata* and *F. japonica* have been shaped by repeated phases of ± ancient hybridization and asymmetric introgression involving extinct species and lineages (**Figs 8, 9**); see also **SupplR&D.pdf**, *section 3.3.4*): the plastomes of the two modern Japanese species encapsulate five independent hybridization events across three major plastome lineages covering a time span from ca. 34 to 1 Ma. In contrast, their nucleomes are markedly distinct showing no sign of recent gene exchange because they have been repeatedly homogenised in the aftermath of secondary contacts. Further studies are required to understand the factors that allowed weakened species barriers since these two species are currently ecologically diverged and are not known to hybridise (Okaura & Harada 2002, Suzuki et al. 2024).

### Episodic migration into lower latitudes of East Asia

*Fagus* originated at high latitudes (Denk & Grimm 2009, Grímsson et al. 2016) and migrated to lower latitudes during the late Eocene-Oligocene cooling (this study). A legacy of a (first) southward expansion are the Lineage IV plastomes carried by *F. hayatae* (SLPT IV-Ha) and one individual of *F. lucida* (IV-LuA). This expansion is dated to the Early Miocene by both FBD- and ND-estimates and ≥ 8 myrs younger than FBD-estimates for the stem age of the *F. hayatae-pashanica* lineage (**Fig. 9; Table 7**; see also **Fig. SR3-12** and **Table SR3-6** in SupplR&D.pdf). At the time the island populations (*F. hayatae*) got completely isolated from the continental populations (*F. pashanica*), their maternal lineages had been long diverged and sorted.

With respect to the geographic history of the *F. lucida* lineage, characterised by a unique leaf and cupule phenotype shared with fossils from the 34 Ma old Kraskino paleoflora (Pavlyutkin et al. 2014) and the Early Miocene of the Altai Mountains (Central Asia, ∼16 Ma; Iljinskaya 1982; Denk & Grimm 2009; this study), it is conceivable that the precursors of *F. lucida* locally introgressed ancestors of *F. hayatae-pashanica,* during migration into their current range. The *F. lucida* species lineage may represent an early continental lineage within *Fagus* subgenus *Fagus* that originally thrived outside the range and niche of the precursors and extinct sisters (donors of Lin. IV plastomes in *F. crenata* and *F. japonica*) of *F. longipetiolata* or *F. hayatae-pashanica*. The only known Miocene candidate precursor is the Central Asian †*F. altaensis* (Denk & Grimm 2009), older fossils with affinity to *F. lucida* but not to any other E. Asian species are present in the north-eastern Asian 34 Ma old Kraskino flora and coincide with maximum possible stem ages of the *F. lucida* species lineage (**Table 7**). Extensive mixing with (an)other, genetically already isolated, *Fagus* subgenus *Fagus* lineage(s) reflects its migratory history further: Jiang et al. (2022) encountered increased levels of heterozygosity at (strongly) evolved alleles shared between *F. lucida* and *F. longipetiolata* (cf. Cardoni et al. 2022, data S5) despite mostly low levels of interspecific gene flow occurring in the middle Pleistocene (Li et al. 2024).

The *F. longipetiolata* species lineage originated in northern North-East Asia during the Eocene-Oligocene transition and underwent several phases of past reticulation related to its southward migration. The modern-day species is characterised by markedly distinct nuclear alleles and ITS variants (Denk et al. 2005, Cardoni et al. 2022; see also Renner et al. 2016). Thus, among the *longipetiolata-*like fossils and prior to the Late Miocene of North-East Asia are the ancestors of the extant species. This lineage migrated into southern East Asia during the Miocene when it came into secondary contact with and was introgressed by the ancestors of *F. lucida* (shared nuclear alleles + SLPT IV-ChB), and eventually evolved into the *F. longipetiolata* last common ancestor (LCA). The last common mothers (LCM) of *F. longipetiolata* and the SLPT IV-ChB are estimated to be of latest Miocene age (**Fig. 9; Table 7; Table SR3-7** in SupplR&D.pdf). The *longipetiolata-* LCM, with an estimated Early Miocene stem and Middle Miocene crown age (**Table SR3-7** in SupplR&D.pdf), may have been exclusive to the precursor of the extant species. It came from the same lineage that also acted as a maternal donor of some *F. lucida* and *F. engleriana*; a lineage formed during the latest radiations within the Lineage IV core clade (**Figs 6, 9**). All available nucleome data (Denk et al. 2005, Renner et al. 2016, Jiang et al. 2022; this study) on *F. longipetiolata* indicates that while its ancestors underwent reticulate evolution, the modern-day species is likely holophyletic: the *longipetiolata-*LCA included not only the *longipetiolata-*LCM but also the last common father of the extant populations.

### Plastid and nuclear concordance in the western Eurasian lineage

The Western Eurasian *Fagus* lineage is the only lineage within *Fagus* that has fully congruent plastid and nuclear differentiation patterns. This suggests that all extant Western Eurasian *Fagus* and their precursors shared an exclusive last common ancestor (LCA). Represented by the oldest records of the fossil-species †*F. castaneifolia* (early Oligocene of Germany and Greece; Denk et al. 2012, Velitzelos et al. 2014), the West-Eurasian-LCA was geographically and genetically isolated from its East Asian sisters since the Oligocene (**Figs 9, 10b,c; Table 6**). Crown-group radiation within the Western Eurasian lineage must have started simultaneously or shortly after this isolation (**Fig. 9; Table 6**).

After reaching Europe (**Fig. 10b**), the Western Eurasian lineage came into contact with a different evolutionary lineage, the eastern populations of a once trans-Atlantic *Fagus* lineage and extinct siblings of the (eastern) North American *Fagus* (**Fig. 10c**). The age of an Euamerican MRCA (Atlantic | Pacific topology) is ± coeval with the stem age of the Western Eurasian lineage in the New World | Old World topology supported by the majority of nuclear gene regions (**Fig. 9; Table 5**). The fossil-species †*F. gussonii* found in the Miocene of the proto-Mediterranean region and the Late Miocene of Iceland is a likely candidate for a late representative of this extinct lineage acting as another (paternal?) donor for the modern-day beeches of Western Eurasia. The lack of plastomes with North American affinity in the beeches of Europe and the Euxinian-Hyrcanian region would indicate that the introgression was unilateral (pollen-mediated), from the Atlantic lineage (†*F. gussonii*) into the West-Eurasian lineage (coeval fossil-species †*F. haidingeri*). However, †*F. haidingeri* is a polymorphic species including precursors of the modern-day species in Western Eurasia (Denk 2004, Denk & Grimm 2009). Part of this variability could have been due to Atlantic-West-Eurasian hybrids, or introgressed Atlantic-lineage *Fagus* such as †*F. gussonii*, still carrying New World-type plastomes (sister lineage or part of the today eastern N. American Lin. I plastomes), and coexisting with the eastern immigrants during the Miocene into the Pliocene. Similar cases are known both from molecular phylogenies and the fossil record woody angiosperm taxa such as *Smilax* (Zhao et al. 2013, Denk et al. 2015), *Platanus* (Vargas et al. 2014; Danika et al. 2024) and others (Kadereit & Baldwin 2012; Denk et al. 2023). During the Pleistocene, remnants of this Atlantic-European beech lineage died out, with its niche occupied by modern-day *F. sylvatica*, the youngest of all Western Eurasian species. Recent work suggests that the European *F. sylvatica* is genetically the most homogenous of all Western Eurasian species and only recently diverged from its eastern sister(s) (Gömöry & Paule 2010, Gömöry et al. 2018, Cardoni et al. 2022).

### Conclusions

Sequencing the whole chloroplast genome of samples representing nearly the entire range of *Fagus*, excluding only a few of the recently described Eurasian species, revealed the existence of five major chloroplast lineages whose geographic distribution and association with species reflects inter- and intra-continental migration, divergence and widespread ancient hybridization between both extinct and extant species lineages. This study provides the most detailed insight yet into the depth of chloroplast divergence in *Fagus* and the complex patterns of sharing of lineages across species boundaries; puzzling differentiation patterns that can be tied to the dynamic biogeographic history of beech as inferred from its exemplary fossil record. Future studies should include samples from under sampled regions including Turkey and Georgia and the western and southern outposts of Chinese species to ascertain whether further lineage diversity occurs in these areas. Regarding future phylogenomic studies on the nucleome, the main challenge will not be to infer a coalescent but to sort and document the conflicting signals accumulated in course of more than 50 myrs of reticulate evolution.

## Acknowledgements

This project was funded by the Japanese Society for the Promotion of Science Grant-in-Aid for Young Scientists A (16H06197) and Japanese Society for the Promotion of Science Grants in Aid for Scientific Research (B) (19H02980). We would like to thank C. Furusawa and H. Kanehara for their invaluable assistance with laboratory work.

## Competing Interests

The authors declare no competing interests.

## Author contributions

JRPW, GWG and TD conceived and designed the experiments, JRPW, PL, ACP, MCS, PSS, JAL-S, Y-CC, KK and NT conducted the fieldwork, JRPW and TU-I undertook the laboratory analyses, JRPW, GWG and TD analysed the data and wrote the manuscript. All authors contributed to the interpretation of results and helped revise the manuscript. JRPW and GWG contributed equally to this work.

## Data Availability and Supplementary Information

The whole chloroplast genome and newly generated cloned nuclear sequences are available on GenBank (*CRC*: accession nos OM322828–OM322928; LFYi2: OM322929–OM323089; **SupplGenetics_ncDNA.xlsx**, sheet *New LFYi2 and CRC accs*); for accession numbers of complete plastome sequences see **Table SM3-1** in SupplM&M). Used molecular matrices, xml-files (for dating analyses), and inference files are included in the Online Data Archive at LINK_TBD.

All supplementary spread-sheet (xlsx) and pdf files can be accessed via the *figshare* project “Supplementary information to Worth et al. (2025)” under the following URL: https://figshare.com/projects/Supplement_to_Worth_et_al_2025_Reticulate_history_of_beech/251480

## Notes

https://figshare.com/projects/Supplement_to_Worth_et_al_2025_Reticulate_history_of_beech/251480

### Competing Interest Statement

The authors have declared no competing interest.

