## Supplementary material for "Whole chloroplast genomes reveal a complex genetic legacy of lost lineages, past radiations and secondary contacts in the dominant temperate deciduous tree genus *Fagus*": SupplR&D

### Supplementary Results & Discussion

This supplementary file contains detailed results, including further in-depth analyses, as well as a discussion of all aspects of nuclear and plastome differentiation underlying the assumptions, derivations and conclusions presented in the main text.

***Used terminology***—The following label conventions are used: in contrast to the common usage in molecular-phylogenetic literature, we distinguish (cf. Felsenstein 2004, chapter 10) between inferences – reconstructed **clades**, i.e. subtrees in inferred and rooted phylogenetic trees – and their evolutionary interpretation: monophyly; the assumption that *all* members of an identified group share an *exclusive to them* common ancestor (Hennig 1950, 1965, 1982; Platnick 1982). At and above the species level and in the face of potential primary (followed by incomplete lineage sorting) and secondary gene flow, *inclusive* (monophyly in a strict sense) and *exclusive* (paraphyly) common origins may be difficult or impossible to discern. Reasons include signal issues, non-dichotomous speciation processes, and explicit ancestor-descendant relationships, which cannot be modelled via dichotomous trees. In such situations, a clade observed in an inferred tree is neither a *sufficient* nor a *necessary* criterion for monophyly. For instance, all trees inferred from complete chloroplast genome (plastome) data show unambiguous support for splitting the oaks (genus *Quercus*) into two clades, resolved as sister lineage(s) (e.g. Zhou et al. 2022) to New World (2 monotypic genera) and Old World castaneoids (3 genera), respectively, or grade(s) embedding them (Yang et al. 2021). Nuclear data, on the other hand, unambiguously supports each modern-day Fagaceae genus, including *Quercus*, as genetically high-coherent clade (intra-clade diversification << inter-clade divergence; genus roots with unambiguous support; cf. Hipp et al. 2020). Following the common assumption that inferred clades are *sufficient* criteria for monophyly (Farris 1983), oaks are at the same time a polyphyletic, comprising two mono- or paraphyletic subgroups, and monophyletic genus.

Hence, we here apply the nomenclature proposed by Ashlock (1971): '**monophyly**' is used in the original, pre-Hennigian sense (Haeckel 1866) to generally denote a putative common origin, and '**holophyly**' for an assumed inclusive common origin (monophyly fide Hennig non Platnick). Only the combination of high genetic coherence (low intra-group differentiation, high inter-group divergence triggered by evolved, group-specific sequence patterns) resulting in (near-)unambiguous branch support for a clade is considered a *sufficient* criterion for holophyly. Evolutionary **paraphyly** is a natural consequence of gradual and 'budding' speciation: a progenitor species, originally holophyletic, is partly replaced by one to several descendant species or a part of a species gets isolated accumulating unique genetic and morphological traits. Co-existing ancestors and descendants are per definition paraphyletic (ancestors) and holophyletic (each descendant). In addition, the evolution of beech involved secondary contacts and gene exchange (see also Mallet 2008). Hybridisation, i.e. the crossing of lineages directly leads to (primary) paraphyly, as the hybrid lineage shares an ancestor with each of the two parental or donor (in case of allopolyploidisation) species or species lineages. Finally, intra-species homogenisation may lead to secondary holophyly, all (surviving) populations of a species can be considered sharing an exclusive, but polymorphic ancestor.

To accommodate such particular situations, Wheeler (2014) proposed additional types of phyly (‘**epiphyly**’, ‘**periphyly**’) as intermediate concepts between paraphyly and holophyly in a classic sense. Being the product of (highly) reticulate speciation processes (cf. **Fig. SR1-1)**, most modern-day beech species as defined by Denk et al. 2024 are either epiphyletic or paraphyletic.

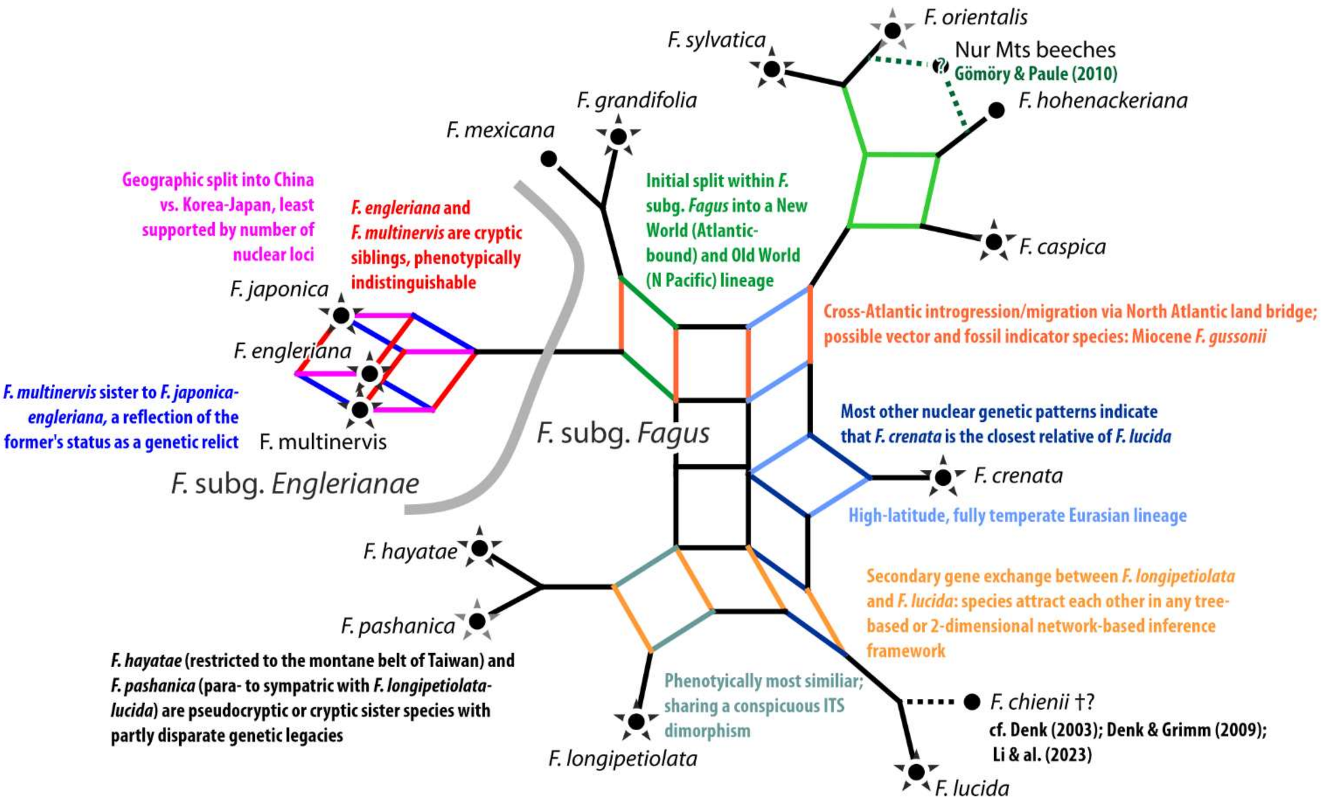


**Figure SR1-1 | Consensus network summarising all known inter-species relationships in beeches** based on various nuclear and morphological studies (Denk 2003, Denk et al. 2005, Grimm et al. 2007, Denk & Grimm 2009, Renner et al. 2016, Cardoni et al. 2022, Jiang et al. 2022, Denk et al. 2024). *Fagus chienii* Cheng is probably extinct: a recent recollection in the original area (Li et al. 2023) only retrieved individuals morphologically and genetically indistinguishable from *F. pashanica.* Star highlight species covered in this study including newly generated data (black stars) and data harvested from gene banks (grey stars).

### 1 A nuclear-based quasi species tree of *Fagus*

Accumulated nuclear data has demonstrated that the evolution of *Fagus* cannot be modelled via a species tree (coalescent) but represents a highly reticulate species network, a 'species coral' (Cardoni et al. 2022; and literature cited therein). We do not have the necessary resources to dig deep into the nuclear genomes of the individuals for which we have obtained complete plastome sequences, to further refine our knowledge about this species network (as shown in **Fig. SR1-1**). The purpose of sampling nuclear data in this study was mainly to confirm the nuclear identity of the main plastome lineages and rule out recent reticulation. Our main objective is to establish a timed divergence framework of the species/individuals covering the main lineages in our plastome sample to assess whether plastid divergences predated or not the species divergences. For this, a nuclear data set is obligatory, because fossil-taxa can only be objectively assigned to nuclear (signatures correlated with phenotypes) but not plastid clades (decoupled from traceable speciation processes). Based on the data available at the start of the project (→ **SupplM&M.pdf**, *section 2*), two nuclear gene regions, a part of the *Crabs Claw* gene (*CRC*) and the 2$^{nd}$ intron of the *Leafy* gene (LFYi2) were targeted that have been successfully amplified in beeches (*CRC:* Oh & Manos 2008; LFYi2: Renner et al. 2016) and showed promise to test the nuclear identity of our samples and main plastid lineages.

#### *1.1 General sequence features*

Matrix dimensions and basic run statistics of the maximum likelihood (ML) tree inference and non-parametric bootstrapping (BS) analysis (cf. **SupplM&M.pdf**, *section 4*) of the full data are summarised in **Table SR1-1**. Both the *CRC* and LFYi2 data show a relatively high diversity (> 300 distinct alignment patterns for a total sequence length of > 1000 bps) and can be generally well aligned leading to a low matrix gappiness. Identical sequence variants are rare and typically restricted to the same individual (cloned data) or same species (cloned and gene bank data). The notable exception is one of our sample of *F. grandifolia* from Quebec, Canada, showing the same *CRC* sequence as the sample of *F. mexicana* included in Oh & Manos (2008; accession no. EU189865). One LFYi2 variant is shared by each one of our *F. engleriana* Seemen ex Diels (accession #34, Anhui, China) and *F. japonica* Maxim. samples (Tohoku prov., Japan) and an individual (*F. japonica,* Oh 82818 TUT) included in Oh et al. (2016; accessions KU565443/KU565444).

**Table SR1-1 | Matrix dimensions and basic maximum likelihood/bootstrapping (BS) statistics of the total *CRC* and LFYi2 data** (new cloned data + harvested gene bank accessions). Full details are provided in RunStats.xlsx included in the Online Data Archive. Abbrev.: LPR = length-polymorphic region; Ts = transition; Tv = transversion.

| **Parameter** | ***CRC*** | | **LFYi2** | |
|---|---|---|---|---|
| | **LPRs incl.** | **LPRs excl.** | **LPRs incl.** | **LPRs excl.** |
| Matrix dimensions — OTUs | 103 | | 238 | |
| Matrix dimensions — Characters | 1702 | 1527 | 1297 | 1146 |
| Distinct alignment patterns | 318 | 224 | 465 | 370 |
| Proportion of undetermined characters[a] | 3.2% | 0.1% | 4.1% | 1.0% |
| Number of necessary BS pseudoreplicates | 700 | 800 | 500 | 550 |
| Approximate optimised substitution model | TVM—abcdbe | TVM—ab**a**cbd | HKY—abaaba | |
| Relative rates for transversions and transitions | $Tv_{CG}<Tv_{AT}\lesssim Tv_{AC}\lesssim Tv_{GT}<<Ts$ | | $1\lesssim Tv<<Ts$ | $Tv_{AC/AT}\lesssim Tv_{GT}\lesssim Tv_{CG}<<Ts$ |
| Number of identical accessions | 6; 4 pairs, 2 triples | | 16; 10 pairs, 3 triples, 4 quartets; 1 quintet | |
| Species sharing identical accessions | *F. grandifolia–F. mexicana* | | *F. engleriana–F. japonica* | |
| Species where different individuals share identical accessions | *F. multinervis, F. japonica* | | *F. crenata, F. longipetiolata, F. sylvatica* All spp. of *Fagus* subg. *Englerianae* | |

[a] Mostly alignment gaps.

##### 1.1.1 CRC

The *CRC* data show a relatively high inter-species diversity but, like earlier studied ITS data, involve intra-individual polymorphism. A substantial part of the intra- and inter-individual variation, most of which is phylogenetically informative[1], is expressed in the form of generally length-polymorphic regions (LPRs). Thus, we have relied exclusively on cloned sequence data able to capture and reflect inter- and intra-individual variation. In total, the sequenced *CRC* gene includes ten portions showing length-polymorphism (**Table SR1-2**; for sequential details see **SupplGenetics_ncDNA.xlsx**, sheet *CRC LP-patterns*): three (simple) indels (2 deletions, 1 duplication); four regions with length-varying mononucleotide repeats (2 multi-A, 2 multi-T); and four with dinucleotide repeats including secondary SNPs (2 $[TA]_x$, 1 upstream-$[TA]_x$-downstream-$[GA]_x$, 1 $[CA]_x$). In contrast to simple indels (insertions, duplications, deletions), the other LPRs cannot be unambiguously aligned across the entire genus but rather represent ± complex but phylogenetically sorted sequence re-organisations/modifications (example provided in **Fig. SR1-2**). By comparison across the genus and on the background of earlier nuclear-phylogenetic analyses (summarized in **Fig. SR1-1**), putatively ancestral and probably derived sequence variants can be identified. None of the LPRs contrasts with the

[1] With “phylogenetically informative” (not to be confused with “parsimony informative”), we address sequential patterns that can inform phylogenetic relationships, either by triggering phylogenetic splits – taxon bipartitions, represented by edges/branches in inferred phylogenetic graphs – or by providing direct evidence for holophyly or monophyly in general in the context of implicit ancestor-descendant relationships: complex, sequential and species-sorted and -conserved mutation patterns such as genetic synapomorphies.

overall tree topology (→ **Section 1.2**). In addition, the differentiation patterns establish ancestral-descendant relationships indicating that *(i) F. multinervis* Nakai appears to be the least-evolved and most-isolated species within *Fagus* subgenus *Englerianae* Denk & G.W.Grimm, and *F. engleriana* the most evolved species sharing a potentially exclusive last common ancestor (LCA) with *F. japonica*; (*ii*) The North American species of *F.* subgenus *Fagus* show evidence for (pseudo)cryptic speciation and are generally still closer to the common ancestor of *F.* subgenus *Englerianae* than any other modern-day member of their subgeneric lineage; (*iii*) within the Eurasian clade of *Fagus* subgenus *Fagus*, *F. hayatae* Palib., *F longipetiolata* Seemen and the Western Eurasian species accumulated more lineage- and/or species-unique mutations than *F. crenata* Blume; (*iv*) the Western Eurasian species are better sorted: both *F. sylvatica* L. and *F. caspica* Denk & G.W.Grimm have each one unique, conserved, derived LPR motive, two are lineage-specific (i.e. shared by both), compared to none for any of their East Asian siblings (**Table SR1-2**; **SupplGenetics_ncDNA.xlsx**, sheet *CRC LP-patterns*).

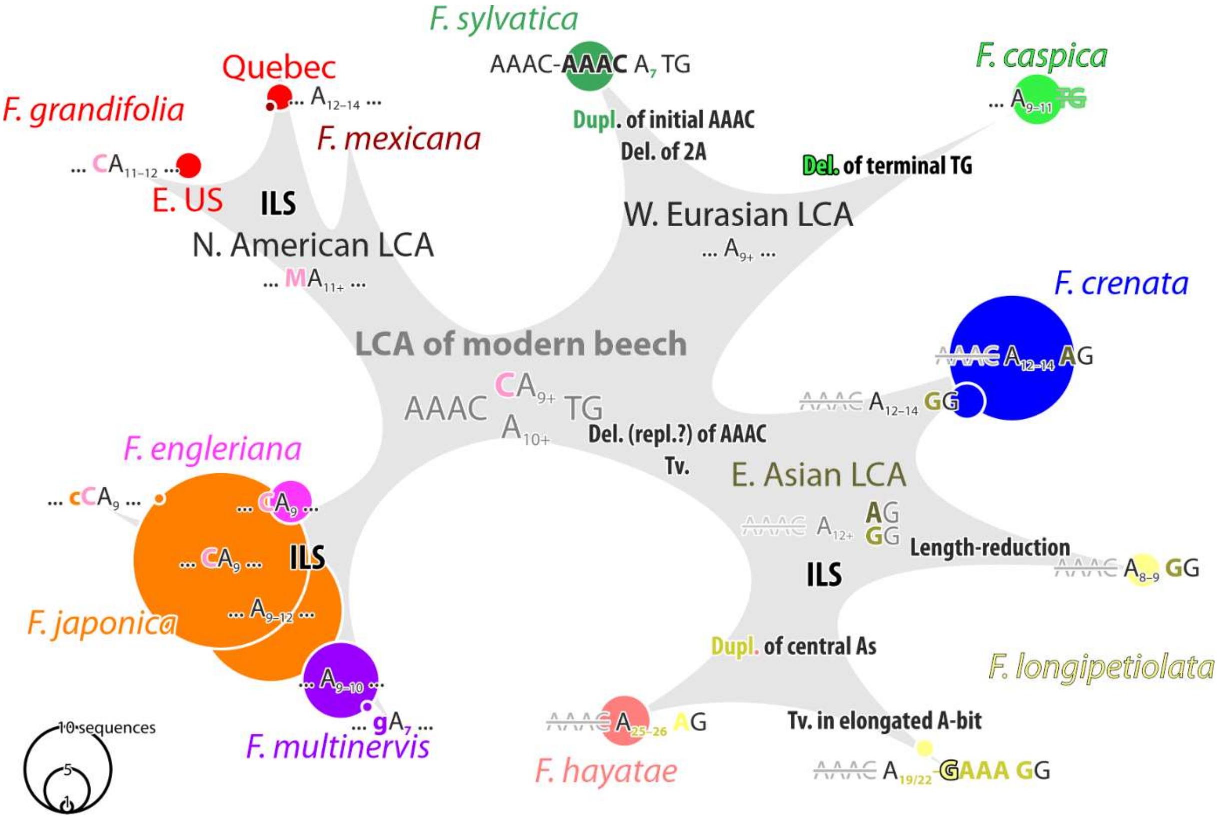


**Figure SR1-2 | Hypothetical evolution of the length-polymorphic oligonucleotide *CRC* motif #5, an A-dominated sequence motif.** The main split is a semi-sorted phylogenetic-geographical A↔T transversion, differing between Eurasian members of *Fagus* subg. *Fagus* and their American sibling, the latter showing the same type as less-evolved *CRC* variants found across all species of *F.* subg. *Englerianae*. Mutational events: Del. = deletion; Dupl. = duplication; Repl. = replacement; Tv. = transversion. Other abbrev.: ILS = incomplete lineage sorting; LCA = last common ancestor.

**Table SR1-2 | Sequential composition of length-polymorphic regions (LPRs) in *CRC* gene sequences.** Currently there is no *CRC* data for four species: *F. hohenackeriana, F. orientalis*, *F. lucida,* and *F. pashanica*.

| Nr[a] | Type | Main split | Shared, ancestral type (Anc.) | Unique, derived type(s) | *Mu* | *Ja* | *En* | *Gr* 1[b] | *Gr* 2[b] | *Cr* | *Ha* | *Lo* | *Ca* | *Sy* |
|---|---|---|---|---|---|---|---|---|---|---|---|---|---|---|
| 1 | [KA]$_x$ | WEA \| rest | [TA]$_{6–9}$-TACA-[GA]$_3$ | ***Me*: [TA]$_2$GA**-[TA]$_8$-**[CA]$_2$**-[GA]$_3$<br>**WEA:** [TA]$_3$-**GT**-[TA]$_4$-**[CA]$_2$**-[GA]$_3$ | Anc. | Anc. | Anc. | Anc. | ***Me*** | Anc. | Anc. | Anc. | **WEA** | **WEA** |
| 2 | [TA]$_x$ | *Gr* \| rest | TA-T-TA | **Tv.**: TA-T-T**T**<br>***Gr*: [TA]$_{2–3}$** | Anc. | Anc.<br>**Tv.** | Anc. | **[TA]$_2$** | **[TA]$_3$** | Anc. | Anc. | Anc. | Anc. | Anc. |
| 3 | T-dom. | None | T$_{10–31}$-CT-GTTT | **Del.**: T$_{21–22}$-GTTT<br>**Red.**: T$_{15–18}$-C<br>**EAS**: T$_6$-**G**-T$_8$-CT-GTTT<br>**ENG**: T$_8$-**G**-CT-GTTT | Anc. | Anc.<br>**ENG** | **ENG** | Anc. | Anc. | Anc.<br>**Del.** | **EAS** | **Red.**<br>**EAS** | Anc. | Anc. |
| 4 | Indel | None | No modification | **Del.**: 28 nt-long deletion<br>**Tv**: A→**T** transversion | Anc. | Anc. | Anc. | Anc. | **Del.** | Anc. | Anc. | Anc. | Anc. | **Tv.** |
| 5 | A-dom. | *Gr*+ENG \|<br>EuAs | AAAC-A$_{10}$-TG | **PAC:** AAAC-**C**A$_{11}$-TG<br>**EAS:** A$_{10–18}$-**R**G<br>***Ha***: A$_{25/26}$-**GA**AG<br>***Lo***: A$_{19/22}$-**GAAA**G**G**<br>***Ca***: AAAC-A$_{9–11}$<br>***Sy***: **[**AAA**C]$_2$**-A$_7$-TG | Anc. | Anc.<br>**PAC** | **PAC** | **PAC** | Anc. | **EAS** | | **EAS**<br>***Lo*** | ***Ca*** | ***Sy*** |
| 6 | Indel | None | CTG | **Del.:** 3 nt-long **deletion** | Anc. | Anc. | Anc. | **Del.** | Anc. | Anc. | Anc. | Anc. | Anc. | Anc. |
| 8 | Indel | WEA \| rest | No modification | **Du.:** 7 nt-long **duplication** | Anc. | Anc. | Anc. | Anc. | Anc. | Anc. | Anc. | Anc. | **Du.** | **Du.** |
| 9 | [TA]$_x$ | *Gr*+ENG \|<br>EuAs[c] | [AT]$_2$ | **Du.:** [AT]$_\mathbf{4}$<br>***Lo***: [AT]$_3$**-AC** | Anc. | Anc. | Anc. | Anc. | Anc.<br>**Du.**[c] | **Du.** | **Du.** | **Du.**<br>***Lo*** | **Du.** | **Du.** |
| 10 | SNLP | None | T$_4$ … T$_3$ | **Tv.**: T**G**TT … T$_3$<br>**Red.: TTT** … **TT** | Anc. | Anc.<br>**Red.** | **Red.** | Anc. | Anc. | Anc. | Anc. | Anc. | Anc.<br>**Tv.** | Anc. |
| 11 | [CA]$_x$ | ENG \| FAG | – | ENG**: [AC]$_\mathbf{4}$**<br>FAG**: [AC]$_\mathbf{3}$** | ENG | ENG | ENG | FAG | FAG | FAG | FAG | FAG | FAG | FAG |

[a] Pattern number; → **SupplGenetics_ncDNA.xlsx**, sheet *CRC LP-patterns*
[b] There are two basic *CRC* variants in North American beeches: *Gr* 1 is private to *F. grandifolia*; *Gr* 2 shared by our Canadian sample and the sample of *F. mexicana* generated by Oh & Manos (2008). The extent of (pseudo)cryptic speciation in North American beeches has so far not been studied (see also Denk et al. 2024).
[c] The [AT]$_4$ pattern is only found in the directly sequenced *F. mexicana* by Oh & Manos (2008). It may be a sequencing/editing artefact.

##### 1.1.2 LFYi2

In general, LFYi2 data is more homogenous than *CRC* data and includes only four major length-polymorphic patterns, two of which are shared exclusively by all accessions of *Fagus* subgenus *Englerianae*: a duplication at pos. 199–205 and a 45 nt-long deletion at pos. 573ff. In the central part of the intron, an A-dominated generally length-polymorphic region (pos. 671ff) is followed downstream (36 bp) by a T-dominated counterpart. In addition, newly studied *F. crenata* samples may show an 8-nt long duplication at pos. 889ff. Both the A- and T-dominated LPR show lineage-conserved (specific) sequence patterns as well as limited intraspecific (old + new data)/-individual (new cloned data) variation (**Table SR1-3**). The intra-individual variation may have caused amplification/sequencing problems encountered in our earlier study (Renner et al. 2016).

At pos. 1130ff, a lineage-specific 6-nt long inversion coupled with a 2-nt deletion discriminates the Western Eurasian species (*F. caspica*, *F. orientalis* Lipsky*, F. sylvatica*) from all other taxa. Based on its sequences, the putative *F.* (×) *moesiaca* specimen (arboretum accession number 617-92, from a cultivated plant of non-georeferenced wild origin) is a *F. sylvatica*[2]: both the upstream A-dominated as well as the downstream T-dominated LPR are identical to *F. sylvatica* samples from France, Spain, and NE. Italy, in striking contrast to *F. orientalis* (the supposed other donor) and *F. caspica* samples (*grandifolia*-similar A-dom. LPR; unique T-dom. LPRs: elongated in the *F. orientalis* sample and much reduced in *F. caspica*, showing a pattern also found in two of Oh et al.'s [2016] *F. crenata* accessions; → **SupplGenetics_ncDNA.xlsx**, sheet *LFY LP-patterns*).

In addition to diagnostic motives in the central LPRs, a number of species-conserved point mutations can be observed, including point mutations consistently separating *F. orientalis* and *F. caspica* from *F. sylvatica* (old & new data) but not *F. pashanica* C.C.Yang (Hubei, P.R.C., old data) from *F. hayatae* (Taiwan, new data).[3]

[2] Up to now there is no genetic evidence for a presumed hybrid origin of *F.* (×) *moesiaca,* a putative hybrid between *F. sylvatica* and *F. orientalis*, reported from the south-eastern Balkans; see e.g. Gömöry & Paule (2010), discussion in Denk et al. (2024).

[3] The *F. pashanica* sequence shows however several polymorphic calls including at pos. 619f, a polymorphism encompassing the difference between *F. hayatae* and *F. longepetiolata*; and a few but singleton point mutations. Newer, cloned data would be needed to verify these patterns and assess whether *F. pashanica* originated from continental *F. hayatae* populations that have been introgressed by or mixed with *F. longipetiolata.*

**Table SR1-3 | Summary of oligonucleotide motif differentiation patterns in *CRC* and LFYi2** (mostly involving inter- and intra-individual length-polymorphism). Numbers refer to **SupplGenetics_ncDNA.xlsx**, sheets *CRC LP-patterns*, *LFY LP-patterns.* Numbers in brackets indicate less-frequent variants forming part of observed intra-individual/-specific dimorphism, typically evolved from the dominant subgeneric or genus consensus (putative primitive, ancestral variant). Italic numbers: co-dominant patterns. Bold-underlined numbers: specific, lineage-conserved patterns (potential genetic synapomorphies).

| **Species** | **Provenance** | ***CRC*** | | | | **LFYi2** | | |
|---|---|---|---|---|---|---|---|---|
| | | Species-unique | Exclusive to 'subg. Fagus' | Characteristic for 'subg. *Englerianae*' | Primitive | Species-unique | Exclusive to either subgeneric lineage | Primitive |
| *F. grandifolia* | Quebec | 1,**2** | 4,**11** | 9 | 3,5–8,10 | **2**,**4** | 6,7 | 1,3,5,8–10 |
| | Pennsylvania[a] | **2**,6 | **11** | 5,9 | 1,3,5,7,8,10 | **2**,**4** | 6,7 | 1,3,5,8–10 |
| | Florida[b] | N/A | N/A | N/A | N/A | **2**,**4** | 7 | 1,3,5,6,8–10 |
| *F. mexicana* | Mexico | **2** | 4,9,**11** | — | 1,3,5–8,10 | N/A | N/A | N/A |
| *F. crenata* | Japan | (3) | 5,9,**11** | — | 1–4,6–8,10 | **2**,**5**,6,(7) | (4),6,(7),10 | 1,3,4,7–9,(10) |
| *F. lucida* | C. China | N/A | N/A | N/A | N/A | | 5,6,10 | 1–4,7–9 |
| *F. longipetiolata* | C. China | 3,(5),9 | (3)[c],5,(7)[c],(9),**11** | — | 1,2,4,6–8,10 | (4),5 | 5,6,10 | 1–4,(5)[f],7–9 |
| *F. pashanica* | C. China | N/A | N/A | N/A | N/A | | 5,6,10 | 1–4,5,7–9 |
| *F. hayatae* | Taiwan | **5** | 3[c],7[c],9,**11** | — | 1,2,4,6,8 | | 5,6,10 | 1–4,5,7–9 |
| *F. sylvatica* | Europe | **4**,**5** | **1**[d],**8**[d],9,**11** | — | 2,3,6,7,10 | **5**,**9** | **2**[e],(4) | 1,3,4,7,8,10 |
| *F. orientalis* | NW. Turkey | N/A | N/A | N/A | N/A | | **2**[e],4,10 | 1,3,5,7–9 |
| *F. caspica* | N. Iran | **5**,10 | **1**[d],**8**[d],9,**11** | — | 2–4,6,7,(10) | **2**,(4) | 4,6 | 1,3,(4),5,7–10 |
| *F. multinervis* | Ulleung Is. | — | — | 9,**11** | 1–8,10 | | **1**,2,**3**,**5**,**6**,10 | 4,7–9 |
| *F. japonica* | Japan | *2* | — | *3*,*5*,9,*10*,**11** | 1,2,*3*,4,*5*,6–8,*10* | (4) | **1**,2,**3**,**5**,**6**,(8),10 | (2),4,7–10 |
| *F. engleriana* | China | — | — | 3,5,9,10,**11** | 1,2,4,6–8 | | **1**,2,**3**,**5**,**6**,10 | 4,7–9 |

[a] Including *CRC* gene bank accession from New York (*F. grandifolia*)

[b] Including LFYi2 gene bank accession by Oh et al. (2016), identical in all oligonucleotide motives.

[c] Same derived motif in *F. longipetiolata* and *F. hayatae.*

[d] Shared by sibling species *F. sylvatica* and *F. caspica* (potential synapomorphy of the holophyletic Western Eurasian lineage)

[e] Shared by sister species *F. sylvatica* and *F. orientalis*

[f] Primitive type only detected in a single directly sequenced individual (Denk 2006010 M; sequenced by Renner et al. 2016), for which no complete intron was obtained due to overlap issues with 5' and 3' reads (possibly triggered by intra-genomic length-polymorphism).

#### 1.2 All-accession ML trees

Each on its own, the two nuclear markers allow focussing on different aspects of the putative species network. In general, *CRC* shows a higher diagnostic capacity at and below the species level, while LFYi2 strongly supports species and species groups at the cost of ambiguity within according subtrees.

To account for potential alignment bias, we inferred trees including and excluding length-polymorphic regions (LPRs). In *Fagus* subgenus *Englerianae* the cloned *CRC* variants form three sublineages, two of which (**E1**, **E2**) represent derived genotypes in *F. japonica* individuals, one (**E2**) shared with the *F. engleriana* individual from Anhui Province. The third, least-derived genotype (**E0**) is private to *F. multinervis* (**Fig. SR1-3**). The LFYi2 variants can be classified analogously but cannot be assigned to distinct clades within the according subtree (**Fig. SR1-4**) being generally more similar to each other and less sorted. The potentially most ancestral variant (**E0**) with respect to the sister subgenus is found in the *F. japonica* Chugoku individual #31 as intra-individual variation (4 out of 6 clones) and two gene bank individuals Oh 5451 TUT (intra-individual variation; 2 accessions) and Oh 5448 TUT (all three accessions).

Clones obtained from our *F. longipetiolata* individual (#36) reflect a consistent *CRC* dimorphism (cf. **Table SR1-2**) including *F. crenata* (genotypes **Cr0** + **Lo1**) vs. *F. hayatae*-related *CRC* variants (**Lo2**) (cf. Grimm, Denk et al. 2007), and three basic LFYi2 variants (**CN1–CN3**) distinguished in the most diverse A-dominate LPR, one of which (**CN1**) is shared with the other three Chinese-Taiwanese species: *F. hayatae, F. lucida* Rehder & E.H.Wilson and *F. pashanica.*

The unambiguously supported and prominent North American-Western Eurasian clade (‘Euamerican clade’), and its sister, the ‘East Asian-subgenus-*Fagus*’ clade, in the clone-based *CRC* tree are supported by a conspicuously low number of conserved SNPs at #328 (A↔C), #1228 (A↔G; East Asia | North America + Western Eurasia) and #1069 (A↔G; East Asian-subg.-*Fagus* | rest). Considering the overall sequence structure – slightly more evolved *F. crenata CRC* genotype **Cr1** not recognised as clade – these SNPs may have been convergently evolved or represent deep divergences that were incompletely sorted. The reason why these splits persist during tree inference and bootstrapping is because incompatible alignment patterns do not consistently support an alternative (e.g. a Eurasian *F.* subg. *Fagus* clade; → **Section 1.4**). The LFYi2 data and tree sets them apart. Clear evidence for introgression of North American gene variants into the Western Eurasian gene pool (as seen in some of Jiang et al.’s [2022] nuclear genes, see supplements to Cardoni et al. 2022 and Denk et al. 2024) are not found in the *CRC* (or LFYi2) data. However, if ancient (ancestral) *CRC* gene variants with yet undiagnostic LPRs (cf. **Fig. SR1-2**) introgressed from a North American (cross-Atlantic) parent into the Miocene gene pool of the Western Eurasian species (→ **Section 3.3.3**), they would trigger reciprocal clades as seen in the *CRC* tree as well (note the placement of *F. caspica* clones in the LFYi2 tree; see also **Section 1.3**).

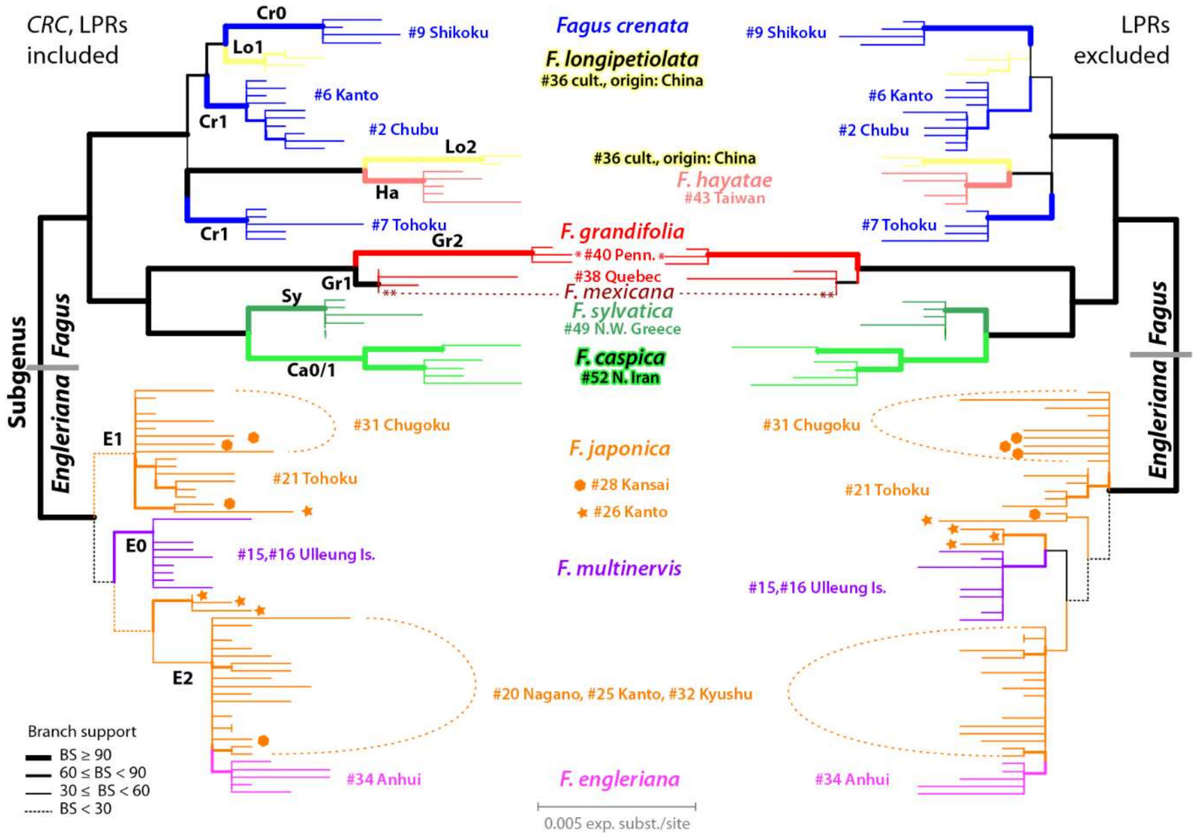


**Figure SR1-3 | Maximum likelihood (ML) trees based on all available *CRC* data**; mostly new cloned data + two directly sequenced accessions by Oh & Manos (2008): *, *F. grandifolia* from New York State, **, representing *F. mexicana*. Left: Length-polymorphic regions (LPRs; cf. **Table SR1-2**) included; right: LPRs excluded. The trees are rooted under the assumption of reciprocal holophyly of the two subgenera (Denk al. 2005, Renner et al. 2016, Jiang et al. 2021). "**Cr0**" etc. refer to main genotypes that can be characterised based on oligonucleotide motives including length-polymorphism (**Table SR1-3**; see also **SupplGenetics_ncDNA.xlsx**, sheet *CRC LP-patterns*)

Even when LPRs are excluded (**Table SR1-2**), the Canadian individual of *F. grandifolia* groups with the *F. mexicana* accession of Oh & Manos (2008; more primitive *CRC* genotype **Gr1**; note the generally shorter root-tip distances compared to *CRC* genotype **Gr2**). The LFYi2 genotypes of the *F. grandifolia* individuals are nonetheless near-identical (compare branch-lengths in **Figs SR1-3** and **SR1-4**; no data on *F. mexicana*): the *CRC* splitting likely represent incomplete lineage sorting of an ancestral *CRC* variant (*CRC* genotype **Gr1,** detected in *F. mexicana* and the Canadian individual) and its potentially *F. grandifolia*-unique derivative (*CRC* genotype **Gr2**).

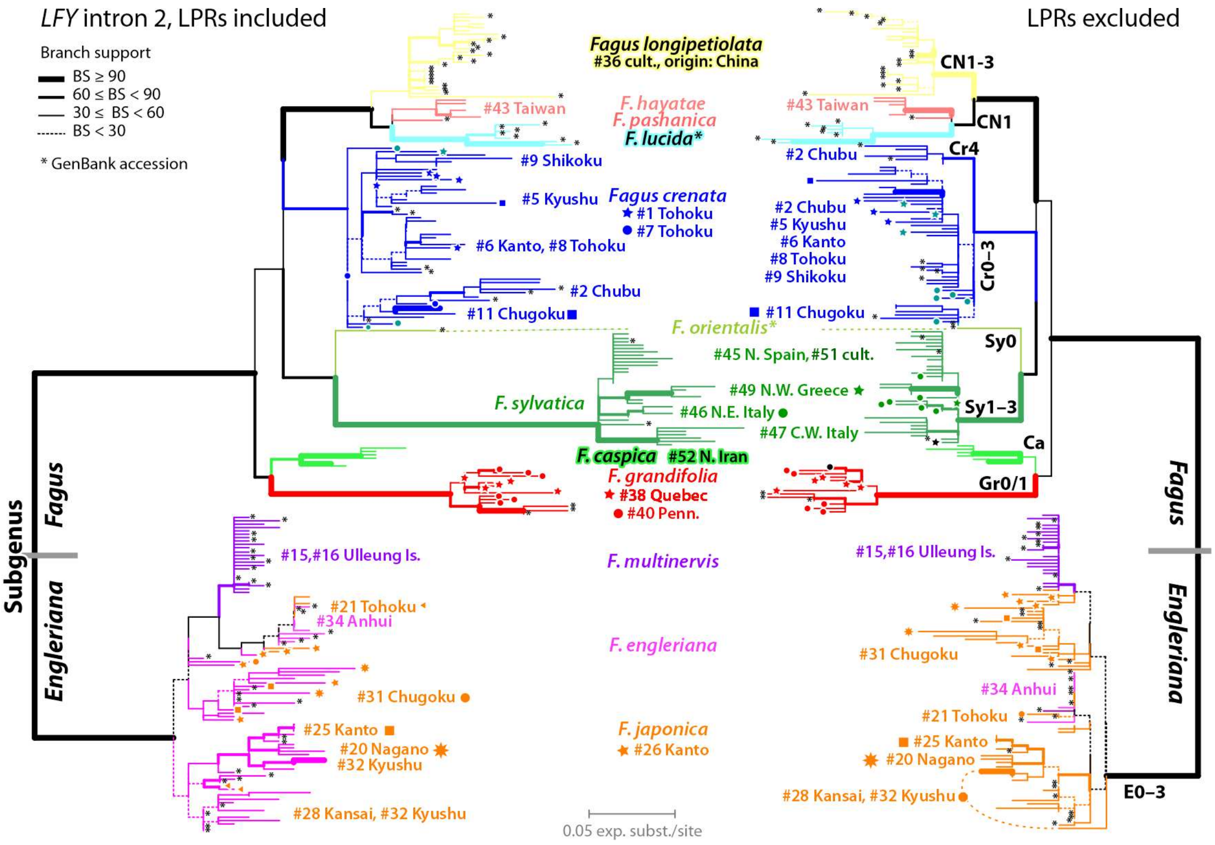


**Figure SR1-4 | Maximum likelihood (ML) trees based on all available *LEAFY* intron 2 data.** "**Cr0**" etc. refer to main genotypes that can be characterised based on oligonucleotide motives including length-polymorphism (**Table SR1-3**; see also **SupplGenetics_ncDNA.xlsx**, sheet *CRC LP-patterns*). Asterisks indicate directly sequenced accessions obtained from gene banks (Renner et al. 2016; Oh et al. 2016)

#### *1.3 Individual-based analyses*

##### 1.3.1 Individual-based phylograms

Individual-level 'ML-A' (cf. Potts et al. 2014) trees are shown in **Figure SR1-5**. The overall topology is equivalent to the cloned-data based trees, i.e. most phylogenetic information in the primary data is captured by using individual consensus sequences. The *CRC*-favoured East Asian-subgenus-*Fagus* clade collapses further; the relationships between the sampled individuals (four *F. crenata*, one each of *F. longipetiolata* and *F. hayatae*) remain ambiguous.

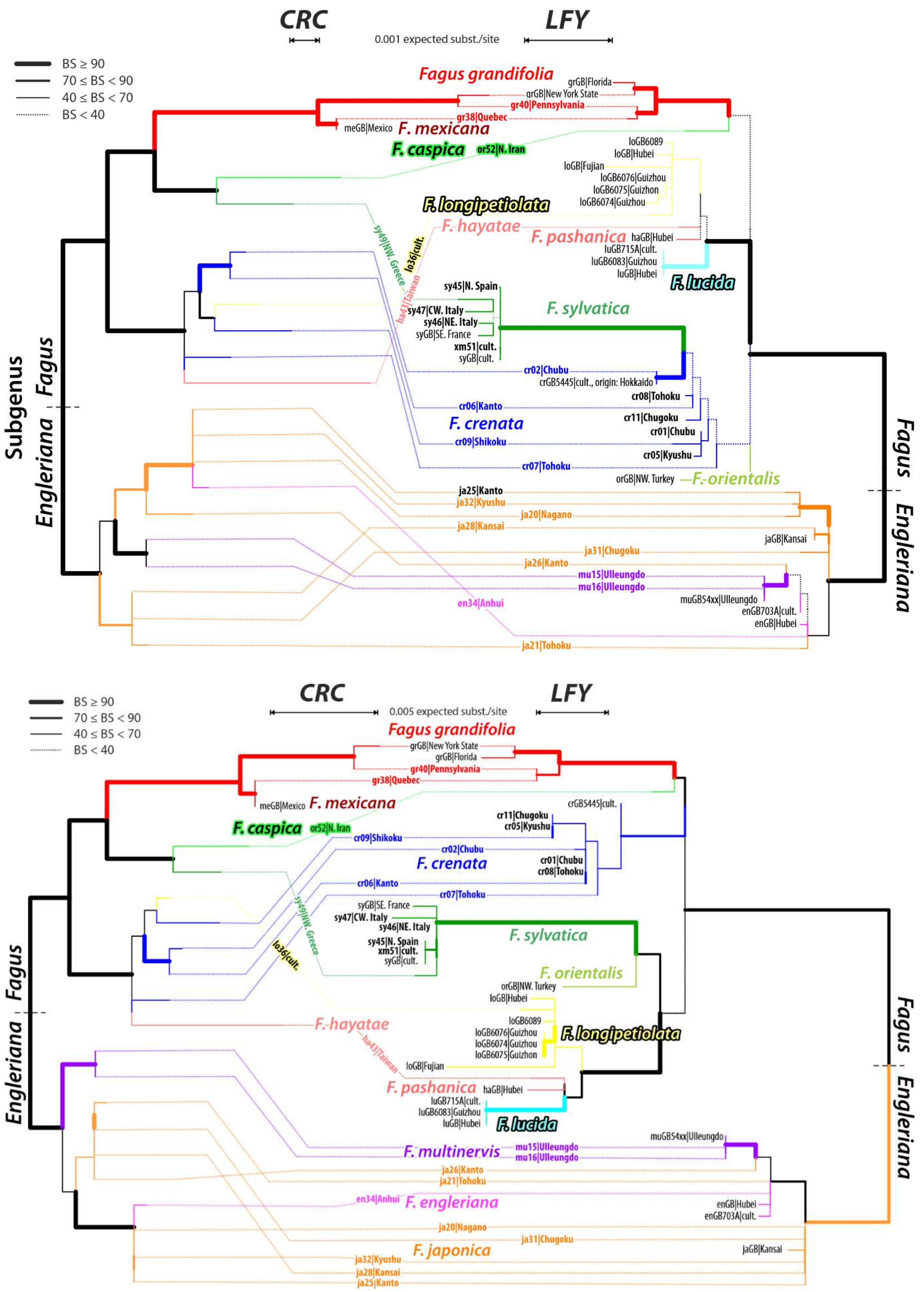


**Figure SR1-5 | *CRC* vs LFYi2 tanglegrams based on individual-consensus sequences.** Top, modal consensi; bottom, strict consensi. Gaps treated as missing data (cf. Göker & Grimm 2008).

The characterisation of *F. crenata* as a species that has retained more of the ancestral polymorphism and the deep divergence within the Eurasian species gradient becomes more apparent: using modal LFYi2 consensi, *F. sylvatica* is nested in a poorly supported *F. crenata* grade, while using strict consensi, it remains as sister to *F. orientalis.* This shift reflects the lack of species-wide conserved and private SNPs in *F. crenata*; species-unique 2ISPs (intra-individual site polymorphism, cf. Potts et al. 2014) are more common but limited to a minority of the *F. crenata* clones, hence, not covered in the modal consensi. It also indicates that the most-distinct LFYi2 variants in the Western Eurasian species can be traced back to a *F. crenata*-like ancestor. As in the case of the clone-based inferences, several ambiguously supported clades indicate unlikely sister relationships (*F. caspica* + *F. grandifolia*), triggered by data-induced (ancestral sequence similarity) branching-artefacts (i.e. paraphyletic clades, for the theoretical background see Grimm 2017).

##### 1.3.2 Individual-based planar (meta-)phylogenetic networks

The PBC-transformed[4] neighbour-nets (**Fig. SR1-6**) recover all ± unambiguous splits in the ML trees (based on primary, untransformed and individual-consensi data) and capture their resolution issues: (*i*) the subgeneric split (*CRC* and LFYi2) with a high-coherent *Fagus* subgenus *Englerianae* and diverse *F.* subgenus *Fagus*; (*ii*) the *CRC*-only split between East Asian and non-East Asian species of *F. s*ubgenus *Fagus*, reduced to *F. hayatae-pashanica, F. longipetiolata* and *F. lucida* when using LFYi2 data; (*iii*) the distinctness of *F. sylvatica* (and *F. orientalis*) and miscellaneous (non-tree like) signal from *F. caspica*; (*iv*) the miscellaneous signal from *F. crenata*, placed relatively close to the centre of the graph as reflection of its generally least evolved genotypes (as outlined above, **Section 1.1**).

Not obvious from the phylogenetic trees is the notable difference between the *F. orientalis* (singleton, directly sequenced) and *F. caspica* (cloned data), which are placed on opposite sides of the LFYi2 graph: the Iranian individual is genetically closer to *F. crenata* and the potential root of *Fagus* subgenus *Fagus* – indicated by the position of the North American *F. grandifolia* in the graph – than its equally low-evolved western sibling, which shows a clear attraction to the most-evolved (within *F.* subg. *Fagus*) set of tips: individuals representing *F. sylvatica.*

[4] Distances between individual clone samples transformed in distances between individuals using the 'phylogenetic Bray-Curtis' distance proposed by Göker & Grimm (2008) for datasets with high proportion of 2ISPs (see also **SupplM&M.pdf**, section 4.2)

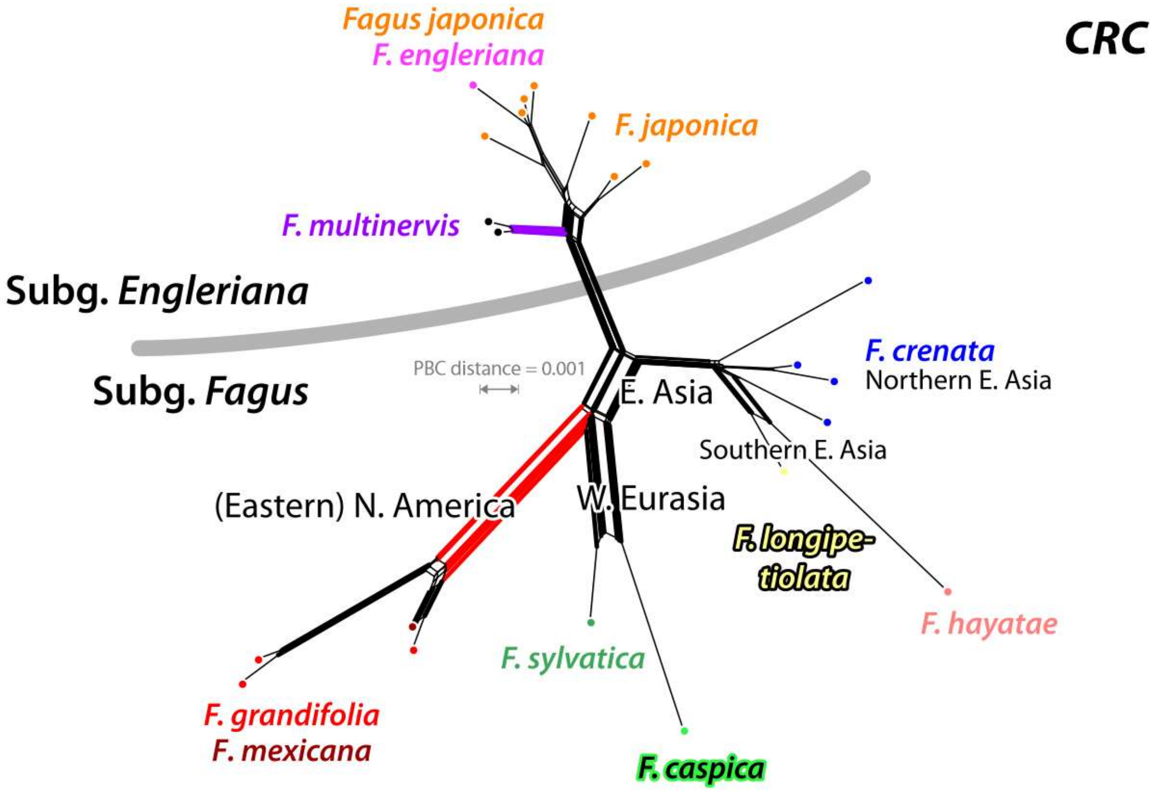


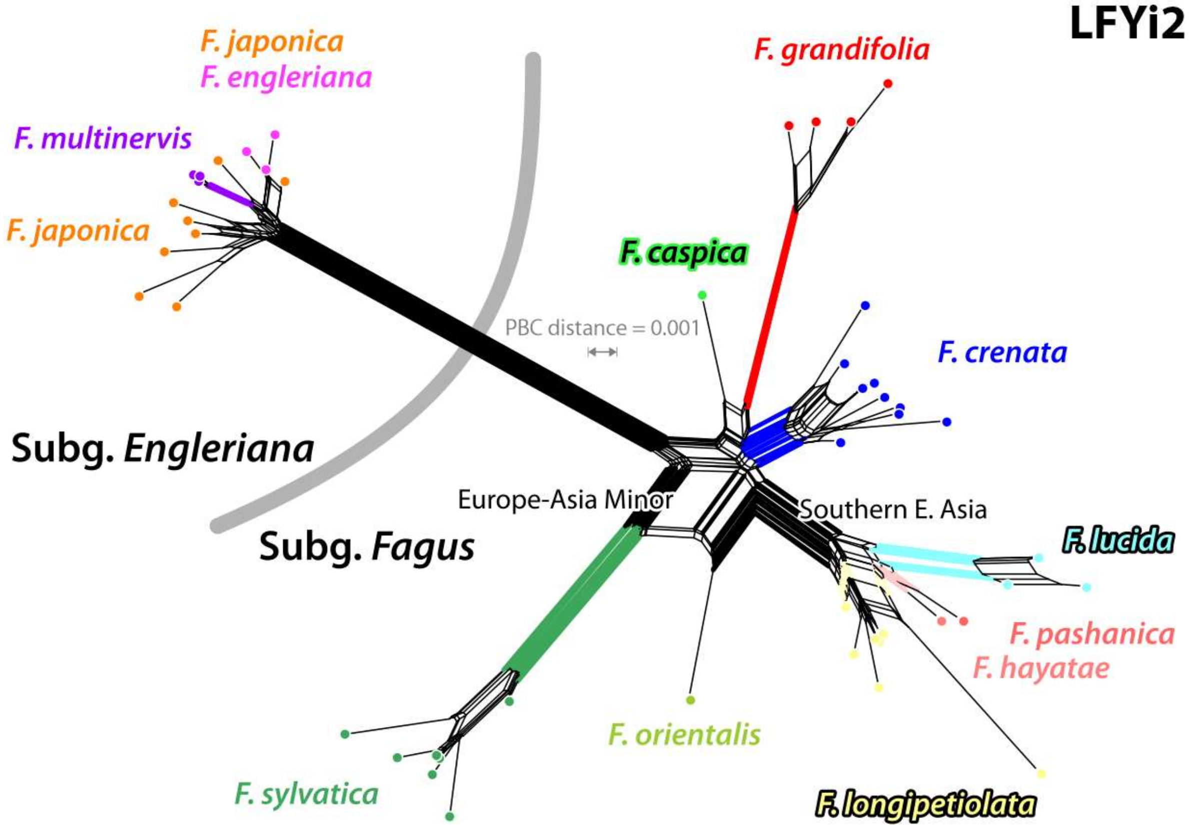


**Figure SR1-6 | Neighbour-nets based on inter-individual 'phylogenetic Bray-Curtis' distances.** Top, *CRC*; bottom, LFYi2. * Individuals ('hosts') represented by a single sequence (as 'associate').

Diversification in the holophyletic Western Eurasian lineage (→ **Section 3.3.3**) must have started right after or even before its isolation from its East Asian sisters (modern-day *F. crenata*).

Within the (continental) East Asian species cluster, *F. lucida* is most distinct, and the only species characterised by a well-developed 'trunk'. The cryptic sister pair *F. hayatae-pashanica* (cf. Denk et al. 2024) are closer to *F. lucida* than their probable sister species *F. longipetiolata* (cf. **Fig. SR1-1**). Based on the observed overall genetic diversity, the divergence in the Western Eurasian species (*F. caspica, F. orientalis* and *F. sylvatica*) is higher than in the East Asian species of *Fagus* subgenus *Fagus* (*F. crenata, F. hayatae–pashanica, F. longipetiolata, F. lucida*), and differ by their genetic affinity with respect to the North American *F. grandifolia.* Applying the empirical species concept (Mallet 1995, 2001), both *F. orientalis* as well as *F. caspica* warrant recognition as species (Denk et al. 2024).

#### 1.4 Discussion of branching/split patterns

**Reciprocal holophyly of subgeneric lineages**—The longest (or second-longest) internal branch (internode) or edge bundles in all phylogenetic graphs, unambiguously supported irrespective of the data used, refers to the split between Shen's (1992) informal subgenera, recently formalised (Denk et al. 2024): *Fagus* subgenus *Englerianae* and *F.* subgenus *Fagus*. This data-wise trivial split is supported by highly conserved sequence patterns in both *CRC* and LFYi2 (**Fig. SR1-4**). While there are no close-enough outgroups for *Fagus* – beeches diverged at least 80 Ma from the remainder of the contemporary Fagaceae (Grímsson et al. 2016) – the consistency and prominence of the subgeneric split in any nuclear data set, low- or high-divergent, ensures that any outgroup (e.g. Jiang et al. 2021; including genes not resolving the subgeneric split, cf. Cardoni et al. 2022, data S5; molecular diagnoses in Denk et al. 2024) or clock-based rooting (Renner et al. 2016) places the genus' root between the two subgenera. Hence, we rooted all trees under the assumption of reciprocal holophyly (Denk et al. 2005, Renner et al. 2016, Jiang et al. 2021), and use this evolutionary scenario as starting point for the discussion of the plastid divergence patterns.

***Fagus* subgenus *Englerianae* species aggregate**—While individual species of *Fagus* subgenus *Fagus* show a generally high coherence and sequential identity, accessions of *F.* subgenus *Englerianae* are mixed. *Fagus japonica* variants typically encompass the genotypic range of *F. engleriana* (our individual and gene bank accessions); the *F. engleriana* individuals fall within a *F. japonica* 'trunk' (*CRC*) or 'fan' (LFYi2) in the individual-based neighbour-nets. Conversely, *F. multinervis* is genetically distinct from the various genotypes observed in *F. japonica* and shows a high coherence in both *CRC* and LFYi2 data. Despite this high coherence, tree inferences fail to unambiguously resolve putative sister relationships (but see Jiang et al. 2021, and the re-analysis of their data by Cardoni et al. 2022) because (*i*) the *F. multinervis* genotype is still relatively close to the common ancestor(s) of the subgenus, (*ii*) the intermediate nature of *F. japonica* accessions bridging between and embracing more ancestral (*F. multinervis*-similar) and most-evolved *F. engleriana* types; and (*iii*) notable intra-specific variation in the *F. japonica* across the Japanese archipelago. Nuclear-genetically, *F. multinervis* seems to represent an early isolate (ancient relict), while *F. japonica* is an originally paraphyletic species from which *F. engleriana* evolved (by 'budding speciation'), currently undergoing secondary homogenisation within the Japanese archipelago.

**Monophyly but not holophyly of East Asian species of *Fagus* subgenus *Fagus*—**Within *Fagus* subgenus *Fagus*, *F. hayatae* (insular, endemic to Taiwan) and *F. pashanica* (mainland, C. China) are resolved as close relatives of *F. longipetiolata* and *F. lucida*, which matches ITS differentiation patterns (Grimm et al. 2007, Göker & Grimm 2008) but also recently compiled single-/low-copy nuclear gene data (Jiang et al. 2021). The latter provides (direct) evidence for secondary gene flow between sympatric *F. longipetiolata* and *F. lucida*, triggering their placement as sister clades in simple combined trees, while a combination of unique and primitive gene variants pull the cryptic sister species *F. hayatae* and *F. pashanica* away from them (discussed and visualised in Cardoni et al. 2022). These four species form a (today) quasi-holophyletic group characterised by a polytomous sibling relationship. The three species lineages differ in their evolutionary history and the extent of (secondary) contact with each other and *F. crenata* and its precursors.

The Japanese *F. crenata* is their closest living relative. *CRC*-wise it forms part of the East Asian-subgenus-*Fagus* clade/neighbourhood. Its *CRC* and LFYi2 genotypes furthermore show a variation that would place the species at or close to the basis of a possible holophyletic East Asian lineage (as found by Jiang et al. 2021; supported by a total of three compatible SNPs and near-unambiguously by a single of the 28 genes, 2 genes producing BS = 50–60 vs. 14 rejecting an East Asian-subgenus-*Fagus* clade with BS = 54–99.7; see also **SupplDating.xlsx**, sheet *pre-FBD topology test*). In addition to the patterns seen in the LPRs, the shifting placement of *F. crenata* in the various nuclear trees could indicate that *F. crenata* represents the (then paraphyletic) source species from which *F. longipetiolata, F. hayatae–pashanica* and *F. lucida* evolved (cf. Cardoni et al. 2022); only secondarily homogenised to some degree being today restricted to Japan and disjunct from its siblings. However, despite being a factor >10 more divergent than any of Jiang et al.'s (2022) 28-genes, neither *CRC* nor LFYi2 show any conserved, uniquely derived LPR feature exclusively shared between *F. crenata* genotypes and any of the East Asian species of *Fagus* subgenus *Fagus* that could pinpoint exclusive sister or ancestor-descendant relationships (**Table SR1-3**). Instead, the species-diagnostic sequence patterns seem to be derived directly from the (Eurasian) *F.* subgenus *Fagus*-ancestor. Earlier nuclear, morphological and historical (fossil) data (Grimm et al. 2007, Denk & Grimm 2009, Renner et al. 2016) identified *F. crenata* as the closest living relative of the Western Eurasian species, which represent an originally holophyletic lineage going back to the Oligocene †*F. castaneifolia* Unger as the first common ancestor (West-Eurasian-FCA; see also Denk 2004). This ultimately Western Eurasian lineage originated in north-eastern Asia close to the precursor(s) of *F. crenata,* migrating west north of the Paratethys (Denk & Grimm 2009, Cardoni et al. 2022, this study); a scenario that is consistent with differentiation patterns seen in the more specific LFYi2 LPRs (motives #2, #4, and #6) but also certain SNPs. In sum, the *CRC* and LFYi2 differentiation patterns would agree with the hypothesis that *F. crenata* shares an exclusive common origin with the Western Eurasian lineage, but, in contrast to them, (repeatedly?) exchanged genetic material with other East Asian species of *F.* subgenus *Fagus* or their precursors. Hence, the failure of the LFYi2 tree to resolve the position of *F. crenata,* namely the Western Eurasian (siblings) and the southern East Asian species (interbreeding cousins) in contrast to (narrower sampled, potentially more biased) *CRC* data (**Fig. SR1-6**).

As in the case of the early assembled and deeply sampled ITS data and Jiang et al.'s (2022; cf. Cardoni et al. 2022) gene sample, the distribution and differentiation of *CRC* and LFYi2 genotypes in the contemporary remaining East Asian species is the product of incomplete lineage sorting and probably primary (early) and secondary (not recent) lineage mixing between the precursors of the modern-day (continental) East Asian species and eastern populations of the *F. crenata*–West-Eurasian species lineage. The notable intra-individual/-specific variation could be partly triggered by speciation processes predating the formation of the modern-day species. Persistent, frequent contact and episodes of backcrossing between (precursors of) *F. crenata* and possibly more ancient continental East Asian species lineages can explain the overall nuclear-genetic similarity of the East Asian *Fagus* subgenus *Fagus* species, which will be reflected by a high-supported clade with relatively short root branch in nuclear data sets showing little phylogenetic structuring in the East Asian species complex such as *CRC* (cf. **Fig. SR1-6**).

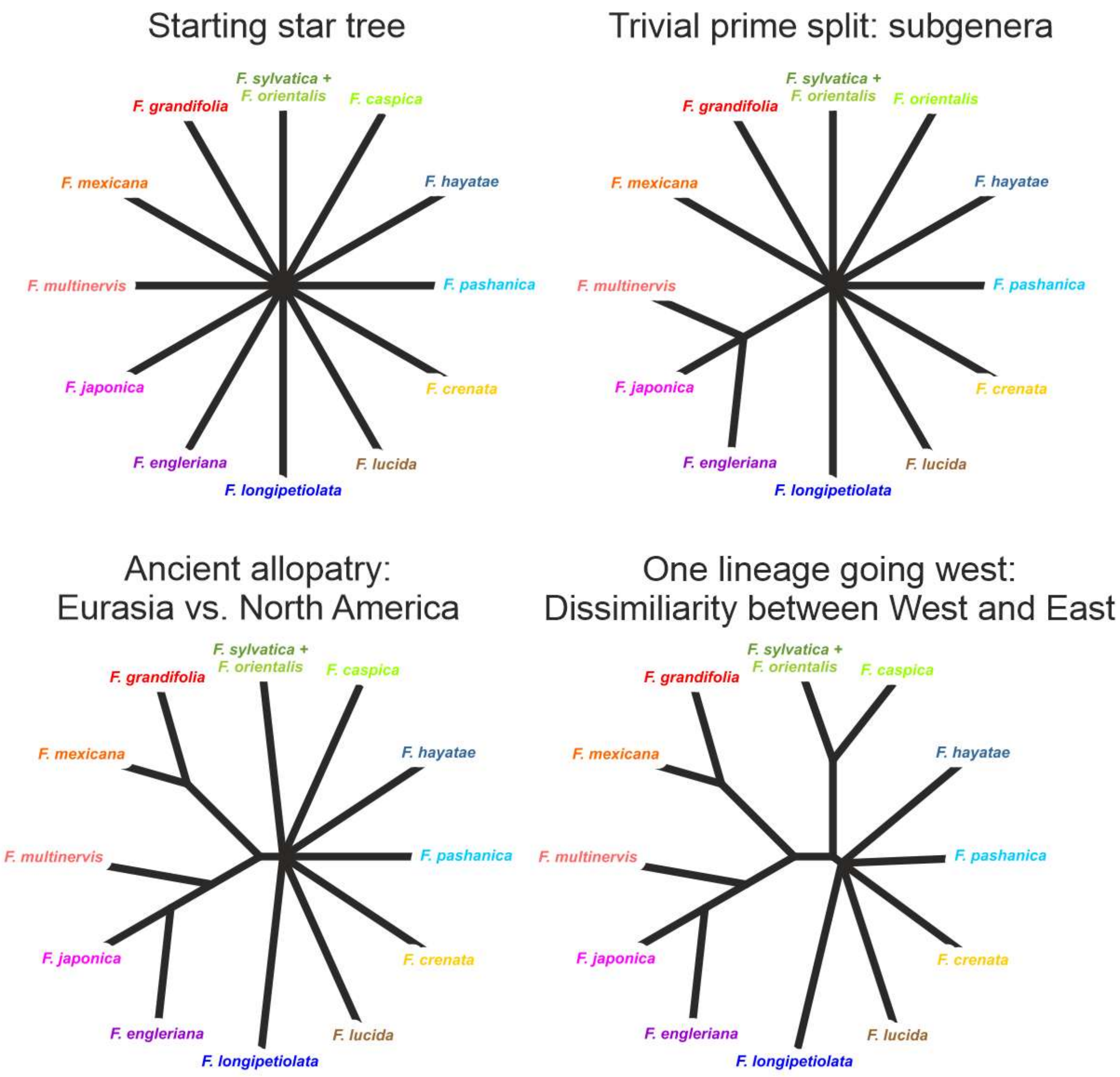


**Figure SR1-8 | Species tree reconstruction in the case of genus *Fagus*.** All species are ± coherent using nuclear-molecular data sets but part of their intra-individual/-specific variation may show disparate affinities. Resulting overall dissimilarity and similarity patterns favour a given sequence of splits pre-defining the topology of any inferred combined-data tree: both (likely) paraphyletic (East Asian spp.) and (putatively) holophyletic groups (W. Eurasian spp.; N. American spp.; subgenera) can form clades.

In contrast, the longer and more persistent geographic isolation of the North American and Western Eurasian lineages facilitated the accumulation and fixation of specific and/or lineage-unique sequence characteristics in past (precursors) and modern-day species (genetic drift). Accordingly, the *CRC* clone-based tree (**Fig. SR1-3**) and the LFYi2 tree (**Fig. SR1-4**) resolve/place the probably paraphyletic East Asian species in a clade, while placing their North American and (part of their) Western Eurasian siblings, being significantly more distinct, as sister clades (inevitable ingroup-outgroup long-branch attraction; 'short-branch culling';

**Fig. SR1-8**). The five extant East Asian species constitute a monophyletic but probably not a holophyletic group.

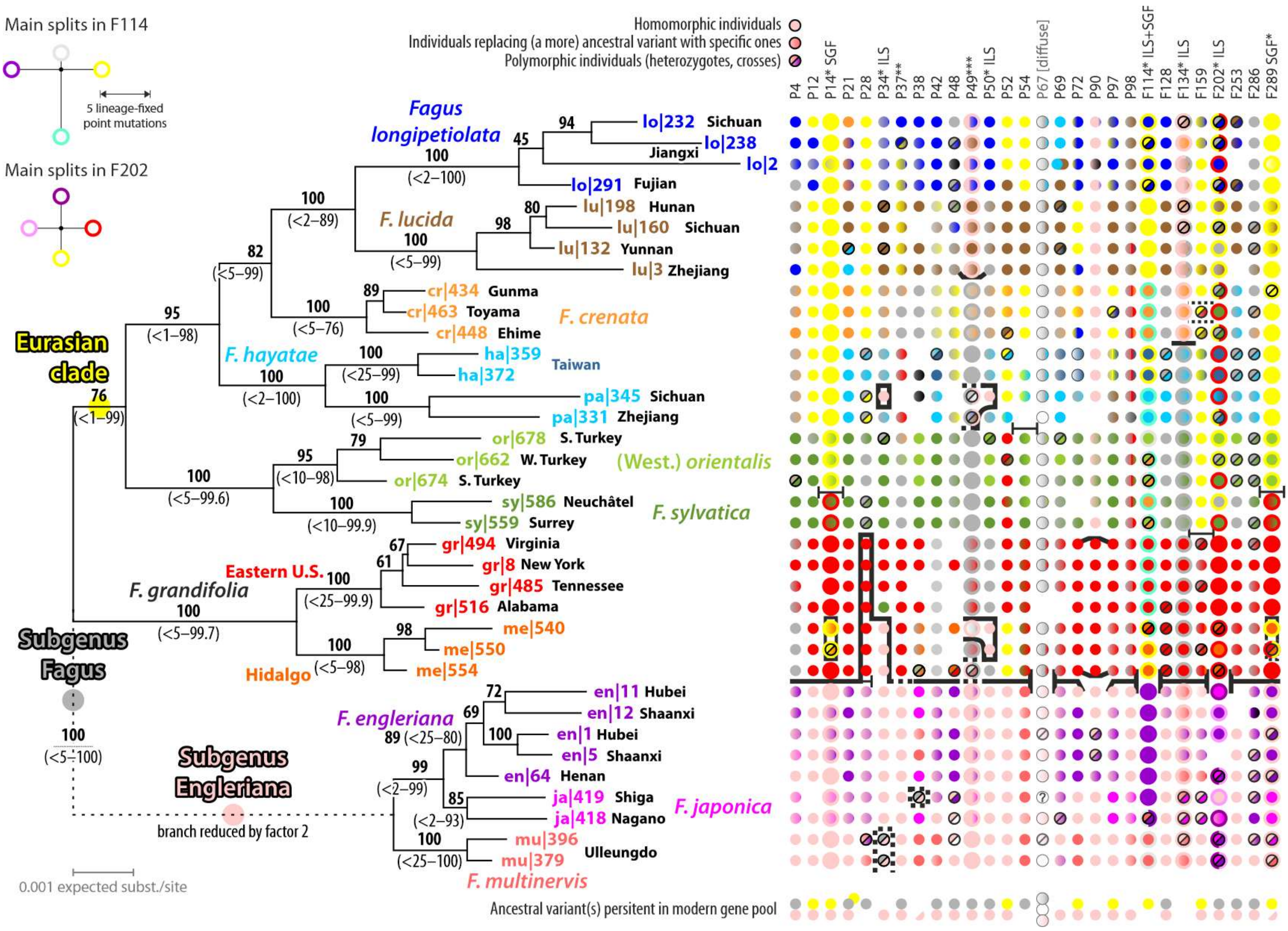


**Figure SR1-9 | Visualisation of per-gene support for branches recovered in an individual consensus sequence based tree** using the (artificial) phased data of Jiang et al. (2022), coloured dots refer to sequence variants (putative alleles) shared between or unique to species, colour gradients reflect shifts from ancestral variants to (specific) evolved variants encoded as 2ISPs (intra-individual site polymorphism; Potts et al. 2014). From Cardoni et al. (2022), data S5.

Having retained most of the ancestral variation and sequence characteristics (see **Fig. SR1-9** for Jiang et al.'s 2022 nuclear gene sample) *F. crenata* acts as a 'rogue taxon' collapsing deep branches and decreasing branch support within the *Fagus* subgenus *Fagus*-clade and its Eurasian subclade (as depicted in **Fig. SR1-9**). It collectively attracts less evolved, more primitive variants of others species, while more evolved, derived variants are attracted by the other members of the subgenus, hence, resolved as sister lineages. The same phenomenon can be observed when combining the 28-gene data of Jiang et al. (2022, fig. 1): *F. hayatae–pashanica* – see also **Figure SR1-10** for first 5S-IGS data, 5S-IGS main types shared between

*F. crenata* and *F. hayatae* or *F. pashanica* but not between *F. hayatae* and *F. pashanica* – are placed as sister clade to *F. crenata* and *F. longipetiolata–lucida* because they are, when summing up across all 28 genes, least similar to *F. crenata* (but see the dated cloudogram in Jiang et al., 2022, showing a faint attraction between *F. crenata* and *F. hayatae-pashanica*[5]).

**Pseudo-cryptic and cryptic speciation in Western Eurasian beeches and *Fagus* subgenus *Englerianae*—**Both *CRC* and LFYi2 clone-based inferences recognise the deep split between *F. caspica, F. orientalis* (only LFYi2 data available) and *F. sylvatica* confirming the under-taxonomisation of the Western Eurasian species, often treated as a single species with two arbitrarily defined subspecies (following Greuter and Burdet 1981).[6] Within *Fagus* subgenus *Englerianae*, *F. multinervis,* endemic to the volcanic Korean island of Ulleungdo and morphologically indistinguishable (Denk 2003) from Chinese *F. engleriana* (i.e. an example for cryptic speciation in beeches), shows the highest level of genetic coherence at the species level within its subgeneric lineage. The *F. multinervis* populations appear to be more isolated from their disjunct sibling species *F. engleriana* (~1300 km min. air-distance; China, widespread) and *F. japonica* (~300 km; W. and S. Japan, till C. Honshu) than the latter two from each other (~1000 km min. air-distance). In contrast to the narrow-endemic Taiwanese *F. hayatae,* genetic drift because of small effective population size cannot explain the genetic distinctiveness of *F. multinervis.* As in the case of *F. hayatae-pashanica*[7]*,* Jiang et al.'s (2022) data and tree resolved *F. multinervis* as sister to *F. engleriana* + *F. japonica* but with the shortest subgenus-*Englerianae*-MRCA – tip distances: in contrast to *F. hayatae*, *F. multinervis* gene variants lack evolved sequence patterns which are more common in *F. engleriana-japonica* (**Fig. SR1-9**). The same can be observed in the more than 10-times more divergent *CRC* and LFYi2 data used here.

In contrast, the Western Eurasian species-level taxa are clearly genetically distinct in both *CRC* and LFYi2, with specific variants being ± evolved from a common ancestor shared with *F. crenata* but not any of the other Asian species and still sequentially close to the consensus

[5] The authors did not discern between the two species despite their notable genetic divergence, possibly for political reasons (One-China-Policy; see Grimm 2025). Accordingly, *F. hayatae-pashanica* is represented by a single tip in their cloudogram. The selected age priors were ≥ 28 Ma (*Fagus* stem age), ≥ 17 Ma (*Fagus* crown age), and ≥ 14 Ma (outgroup crown age) too young (cf. Grímsson et al. 2016, Renner et al. 2016, Hipp et al. 2020).
[6] A paper collecting genetic and morphological evidence for a four-species concept for Western Eurasian beeches has recently been publlished (Denk et al. 2024).
[7] Placed as sister clade to the other three East Asian spp. of 'Subgenus Fagus' with unambiguous support. The continental cryptic sibling *F. pashanica* showing generally lower root-tip distances than the insular *F. hayatae.*

of all Eurasian *Fagus* subgenus *Fagus* species, which is expressed in the long terminal tip branches in the cloned-data trees (**Figs SR1-3, SR1-4**).

**Limitations, sample bias, and blank spots**—Based on the *CRC* sequence patterns and the single-/low-copy nuclear gene data of Jiang et al. (2022) and with respect to earlier (cloned) ITS data (Denk et al. 2005), it can be expected that the currently available (gene bank and new) nuclear data only superficially captures the actual gene pool diversity in the North American beeches, particularly with respect to the isolated Mexican populations (pseudo-cryptic species *F. mexicana*; cf. Denk et al. 2024). It possibly underestimates the diversity in Chinese species, especially in *F. engleriana* but also the equally scattered Chinese species of *Fagus* subgenus *Fagus* (**main-text fig. 1**; see also maps in Worth et al. 2021). Further cloned *CRC* and LFYi2 data but also ITS and 5S-IGS data (**Fig. SR1-10**) would be instrumental to evaluate the permeability of species boundaries in the Chinese species, identify and discern reticulation due to primary and secondary contact and (re-)evaluate the species status of distinct (nuclear) genotypes. While producing (near-)unambiguously resolved (species) trees (**Fig. SR1-9**; Jiang et al. 2022, fig. 1) concatenated phylogenomic data from singleton or a few placeholders (one per acknowledged species) will always be incomprehensive: to infer the likely complex and reticulate inter-species relationships in East Asian beeches, as many as possible individuals from different provenances need to be sampled and a proper assessment of intra-genomic variation (as done by Cardoni et al. 2022 for the 28-gene data generated by Jiang et al. 2022) will be critical.

For the Western Eurasian beeches, the current data set does not include any or little genetic information on three critical provenances for understanding the evolution and speciation of *Fagus* in Western Eurasia: the Pontian (Caucasian) Oriental beech populations of *F. hohen-ackeriana* Palib. (Denk et al. 2024, included in **Fig. SR1-10**), *F.* (×) *taurica* Popl. from the Crimean Peninsula, and the isolated populations in the Nur Mountains (Hatay province, S. E. Turkey; sample included in **Fig. SR1-10**; cf. Gömöry & Paule 2010; Kurz et al. 2023).

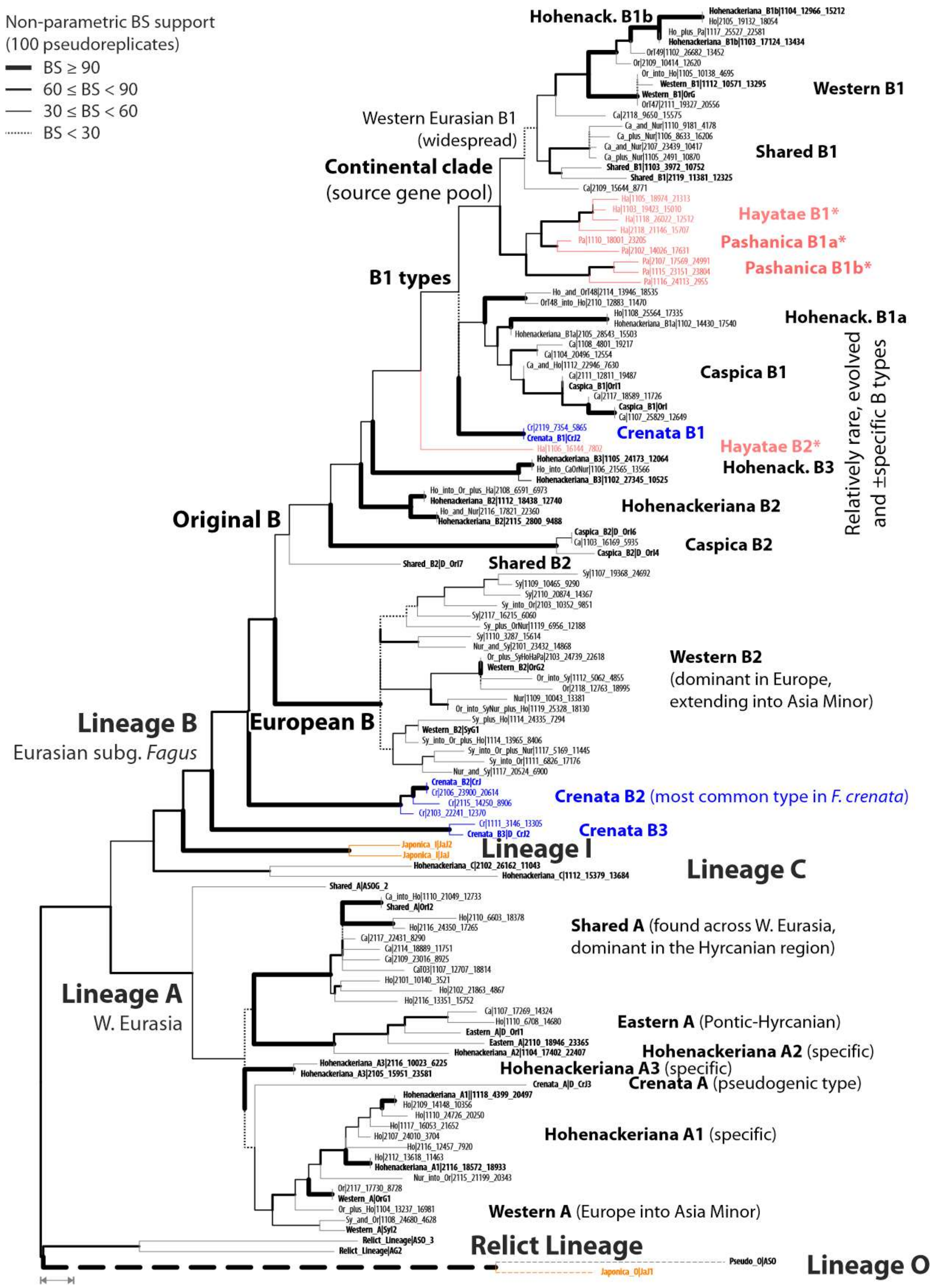


**Figure SR1-10 | Maximum likelihood tree including most-frequent 5S-IGS variants from 19 samples covering the range extremes of *Fagus* across Western Eurasia as well as two samples of the cryptic sister species *F. hayatae* and *F. pashanica*.** While most of their variants form part of the same subtree within the core clade of Lineage B (cf. Cardoni et al. 2022), all these variants are exclusive to either *F. hayatae* or *F. pashanica*. With respect to these new data, Cardoni et al.'s Lineage A represents an evolutionary source near-exclusive to the Western Eurasian species. Taxonomic reference tips for main diagnostic types (Denk et al., submitted) in bold font. Source: S. Cardoni, M. C. Simeone, G. W. Grimm (work in progress).

### 2 A comprehensive chloroplast genome tree

Our complete chloroplast genome (plastome) data includes 82 accessions, 49 newly generated for this study. *Fagus* plastomes have the classic quadripartite structure of angiosperm plastomes consisting of a large single copy (LSC) 88,826 bp in length, a small single copy (SSC) unit of 19,150 bp, and the two inverted repeat regions (IR) of 25,932 bp and encode for 129 genes (**Fig. SR2-1**).

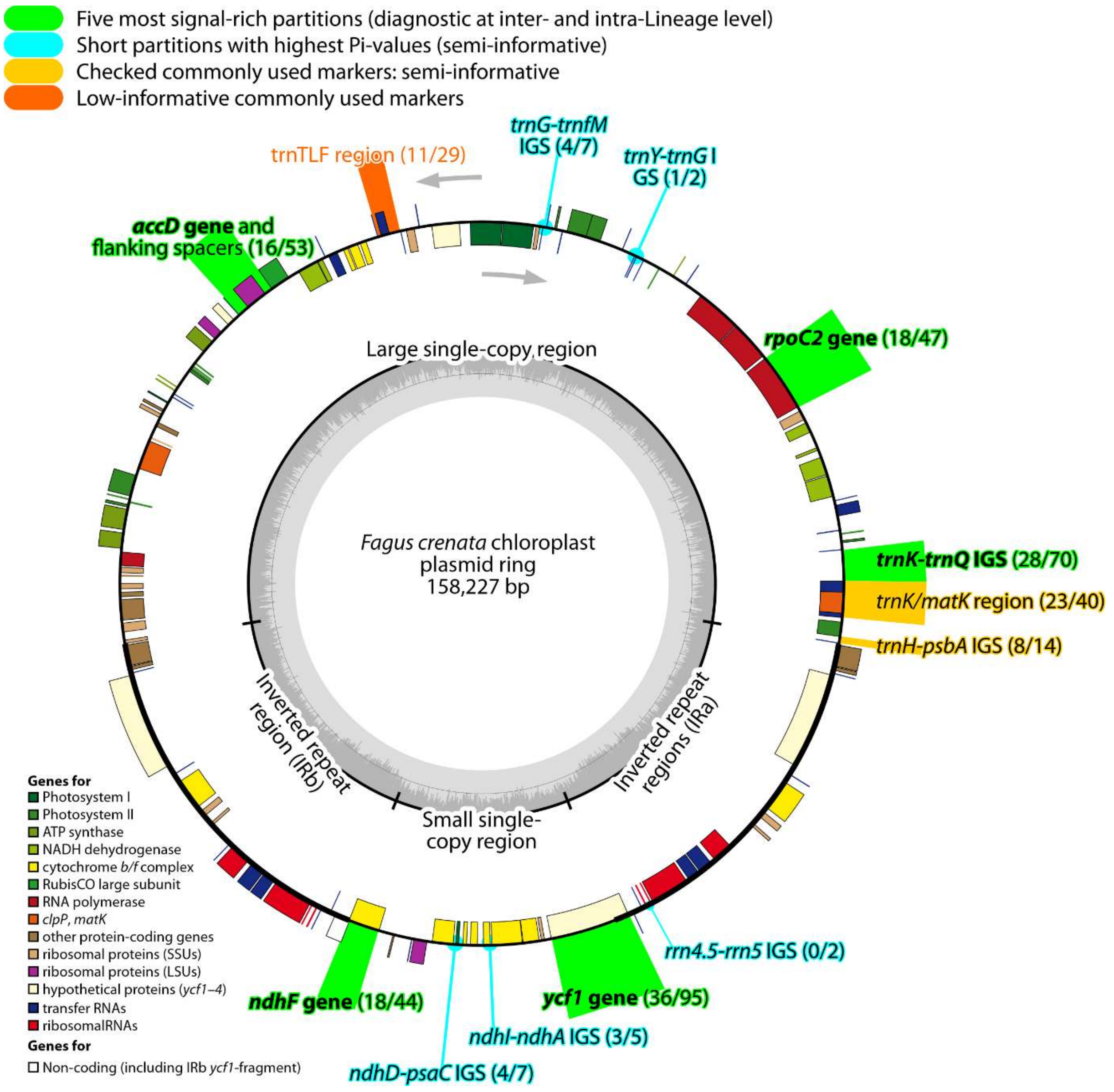


**Figure SR2-1 | The chloroplast genome of beech** with most signal-rich regions highlighted (modified after Worth, Liu et al. 2019). Fully annotated mutation tables for all marked regions are included in the supplement to Denk et al. (2024).

The total alignment of 82 whole chloroplast genomes was 159,840 bp in length and 113,902 bp when excluding the second IR region. There were a total of 1642 polymorphic sites of which 407 were singletons and 1235 were parsimony informative. The overall nucleotide diversity (*Pi*) was 0.00174 and the average number of nucleotide differences (*k*) was 227.9. Insertion/deletion events numbered 664 in total including SSR repeats with indel diversity per site, *Pi*(i), being 0.00093. Sliding-window analysis (**ODA**, file *SlidingWindowResult.xlsx*) and per-partition divergence statistics (**SupplGenetics_cpDNA.xlsx**, sheet *cp gene stats*) revealed that most variation is ± equally scattered across the entire plasmid ring, with only isolated and small diversity hotspots. The five most signal-rich partitions included four long protein-coding genes and only one, but very long intergenic spacer (IGS): (*i*) *rpoC2* gene, (*ii*) *trnK-trnQ* IGS, (*iii*) *ycf1* gene, (*iv*) *ndhF* gene, and (*v*) *accD* gene (and flanking IGS regions). Traditionally used plastid 'barcode' regions (such as *trnH-psbA* IGS, *trnK/matK* and trnTLF regions[8]) have very little discriminative signal (for a tabulation of mutations in these plastid gene regions see **ODA**, file *00_bits.xlsx*).

**Figure SR2-2** shows the result of the ML/BS analysis (GTR+Γ, with 6 partitions defined by function) coloured for major plastid lineages and species. The preliminary results of Worth et al. (2021) are confirmed by the enlarged data set: the plastomes are not sorted by subgenera or species in the East Asian taxa, with both Japanese species showing by far the highest plastome diversity. In contrast, the plastomes of the widespread and accordingly represented North American and European species, *F. grandifolia* and *F. sylvatica,* are near-identical (≤ 23 SNPs per whole plastome) despite ≤ 1980 km air-distance between our samples and ~ 2600 km max. air-distance between *F. sylvatica* samples generated and used by Ulaszewski et al. (2021; cf. **SupplGenetics_cpDNA.xlsx**, sheet *PlstmDissim*). As in other Fagaceae and northern hemispheric extra-tropical tree genera (e.g. *Acer*, Grimm 2022a,b and literature cited therein; *Quercus*, Yan et al. 2019, Li et al. 2025; see Yang et al. 2021 and Zhou et al. 2022 for Fagaceae and Fagales in general), the beech plastomes show a strong geographic sorting.

[8] Comprising the *trnL* gene and flanking intergenic spacers: *trnT-trnL* IGS, *trnL-trnF* IGS.

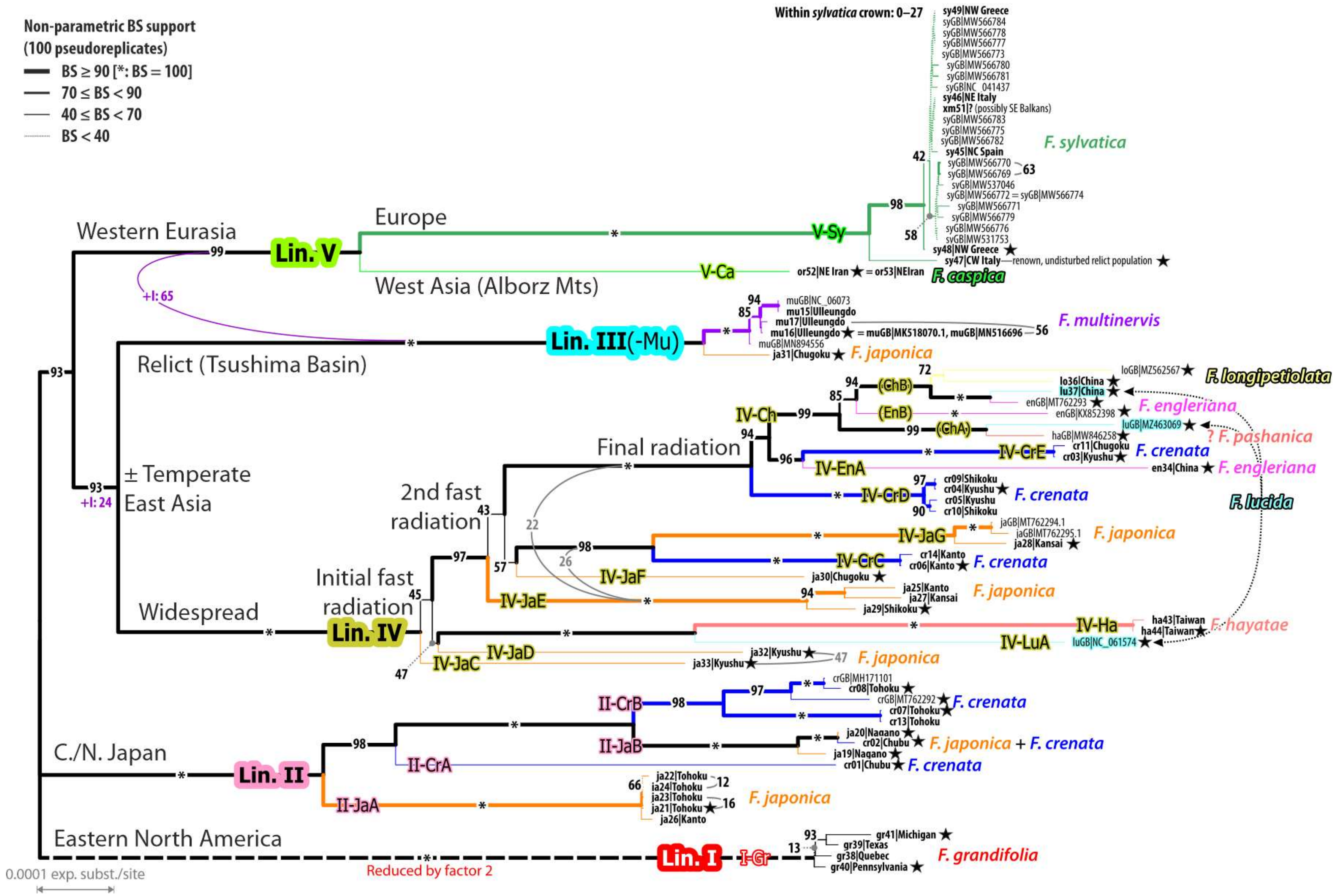


**Figure SR2-2 | Maximum Likelihood tree inferred from the full tip-set including data from 77 complete plastomes**; new accessions in bold, label including provenance. BS support annotated, including topological alternatives in grey. Purple "+I" tangles: alternative topology preferred by ML /BS analyses including a parameter modelling the proportion of invariant sites placing Lineage III as sister to Lineage V. II-JaA etc. denote here defined 'species-level plastid types' (cf. **Section 2.2**).

The five main plastome lineages (denoted by Roman numerals) reflect trans- and intra-continental disjunctions of various scales: (*i*) New World, Lineage I (eastern N. America) vs. Old World (Eurasia), Lineages II–V; (*ii*) East Asia, Lineages III+IV vs. Western Eurasia, Lineage V; (*iii*) adjacent to Tsushima Basin, Sea of Japan, Lineage III vs. continental and oceanic (north-)eastern Asia, Lineage IV; (*iv*) subtropical to temperate (south-)western Japan, Lineage IV vs. (cold-)temperate central and northern Honshu + Hokkaido, Lineage II; (*v*) China + southern Japan (Kyushu and Shikoku islands) vs. Taiwan + Japan, intra-Lineage IV diversification; (*vi*) Alborz Mountains and adjacent montane regions facing the Caspian Sea (V-Ca) vs. Europe (V-Sy). The latter, notably pronounced split, is the only major plastid divergence that can be linked to a speciation event expressed in modern-day beech species (*F. caspica* ↔ *F. sylvatica*[-*orientalis*]; **Fig. SR2-2**; see also Cardoni et al. 2022).

#### 2.1 Partition effect

Overall, beech plastomes are low-divergent (note the scale in **Fig. SR2-2; → SupplGenetics_cpDNA.xlsx**, sheets *PlstmDissim* and *cp gene stats*), hence, saturation issues or strong alignment bias are irrelevant for the inference. Nonetheless, to assess possible under- and over-parameterisation effects, we performed:

1. an unpartitioned ML analysis, same model optimised for the complete data;
2. a fully partitioned ML analysis, defining each functionally and product-wise different plastome segment as individual partitions, i.e. three partitions for each protein-coding gene ($1^{st}$, $2^{nd}$, and $3^{rd}$ codon positions of each gene's exon), and one partition each for each coding (exon) and non-coding (intron) tRNA part, each rRNA gene (*rrn4.5, rrn16, rrn5, rrn23*), and each intergenic spacer (total of 389 partitions);
3. a 'logical' partitioned analysis using six partitions, each grouping base pairs by function and primary genetic constraints: (*i*) $1^{st}$ + $2^{nd}$ codon position, defining amino acid sequence of proteins; (*ii*) $3^{rd}$ codon position, susceptible to synonymous mutations (and back mutations); (*iii*) structurally strongly constrained transfer RNA gene (tRNAs) and (*iv*) ribosomal RNA genes (rRNAs); (*v*) noncoding, usually relatively conserved (in temperate trees) introns of protein-coding genes and tRNAs; (*vi*) the neutrally evolving junk DNA interspersing the genes, the intergenic spacers; (*vii*) the infA pseudogene.

The number of distinct alignment patterns (DAP) is generally low in case of the fully partitioned analysis and for the tRNA, rRNA and infA partitions in case of the logical-partitioned analyses, leading to unreasonably high α-values for the Gamma distribution. When adding the proportion of invariant sites as an additional parameter to the fully partitioned analysis, RAxML failed to infer meaningful branch lengths (**Fig. SR2-3**). Aside from that, there were only two notable differences between the partitioning schemes and GTR+Γ vs. GTR+Γ+I inferences: (*i*) the placement of *F. multinervis* plastomes (Lineage III); and (*ii*) the sequence within the initial and secondary fast (ancient) radiations in Lineage IV.

When the proportion of invariant sites is included as a parameter or in case of the fully partitioned analyses, the East Asian relict Lineage III is placed as sister to the Western Eurasian Lineage V (↔ **Fig. SR2-2**). The competing and well supported East Asian clade (**Fig. SR2-2**) comprising Lineage III and IV (unpartitioned and logical-partitioned GTR+Γ: BS > 90) is, however, not rejected in these analyses but represents the 2nd best alternative captured in the BS pseudo-replicate sample (**Table SR2-1**). Such signal ambiguity can be related to implicit quasi-ancestor-descendant relationships, with Lineage III representing the ± primitive (underi-ved, non-evolved) plastomes and Lineages IV and V its stronger evolved sibling lineages that underwent generally higher genetic drift (see discussion and interpretation in **Section 3.3**).

**Table SR2-1 | BS support for competing topological alternatives by model and partitioning scheme.** Full list is included in **SupplGenetics_cpDNA.xlsx**, sheet *cp splits*. Abbrev.: NW = New World (eastern N. America), OW = Old World (Eurasia); JaC, etc. refer to 'species-level plastid types' (SLPTs; → **Section 2.2.1**) annotated in **Fig. SR2-2**.

| Split, clade | Logical | | Unpartitioned | | Fully partitioned | |
|---|---|---|---|---|---|---|
| | GTR+Γ | …+Γ+I | GTR+Γ | …+Γ+I | GTR+Γ | …+Γ+I |
| **Initial divergences** | | | | | | |
| NW (Lin. I) \| OW | 100 | 100 | 100 | 100 | 100 | 100 |
| Lin. I/II \| III–V | 93 | 80 | 68 | 56 | 99 | 80 |
| Lin. III+IV | 93 | 24 | <13 | <13 | 99 | 22 |
| Lin. III+V | <7 | 65 | 87 | 34 | <1 | 74 |
| Lin. IV+V | <7 | <11 | <13 | 53 | <1 | <4 |
| **First divergence in Lin. IV** | | | | | | |
| IV-JaC \| rem. IV | 45 | <20 | 24 | <20 | 40 | <13 |
| IV-JaC/-JaD \| rem. IV | 37 | 52 | 64 | 33 | 46 | 79 |
| IV-JaD/-LuA/-Ha \| rem. IV | 47 | <20 | <20 | <20 | 42 | <13 |
| IV-LuA/Ha \| rem. IV | <20 | 42 | <20 | 36 | <20 | <20 |

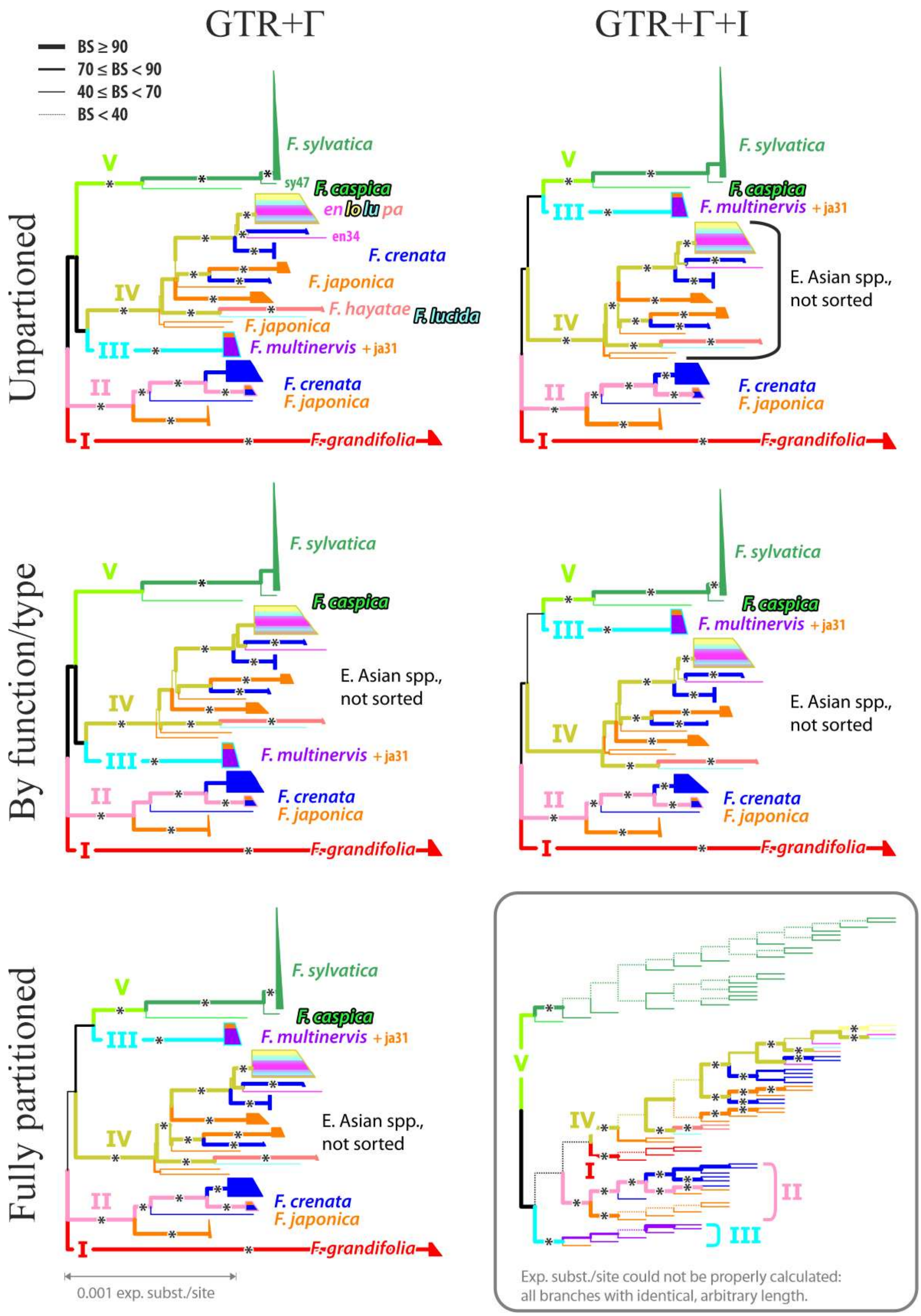


**Figure SR2-3 | ML phylograms using different partitioning schemes and models.** Left: GTR+Γ, right: GTR+Γ+I; top-row: using a single partition for the entire data, middle: using six genetic-biologically informed partitions, bottom: over-parametrising, fully partitioned analysis with 389 partitions, most of which have <10 distinct alignment patterns.

The reason for the ambiguous relationships linked to the primary radiations within Lineage IV is a general lack of discriminate (note the short backbone branch lengths in **Figs SR2-2, SR2-3**) and sorted differentiation signal. The BS pseudoreplicate samples always show the same competing alternatives but with varying frequency (**Table SR2-1**). Such a signal and inference situation is typical for fast (and ancient) radiations: the breaking up of a widespread species (or species aggregate) with near-simultaneous isolation and incomplete sorting of plastid pools along a geographic gradient. In the case of Lineage IV, three of these fast radiation events left their imprint on the plastomes found in modern-day East Asian species, mainly in the Japanese archipelago:

1. The initial radiation involves genetically unique Japanese relict populations of *F. japonica* (species-level plastid types, SLPTs, 'IV-JaC' and 'IV-JaD'; **Fig. SR2-2**), the plastome lineage leading to and conserved in the modern-day *F. hayatae* ('IV-Ha'), an insular isolate, and the main lineage of (originally continental?) Lineage IV plastomes (China-Japan). A newly reported plastome and unique SPLT of *F. lucida* ('IV-LuA') is the product of this radiation as well. Given that the plastid HTs isolated during this radiation cover individuals from both subgenera and today allopatric species (*F. japonica, F. lucida, F. hayatae*), this radiation must have been followed by later hybridisation/introgression events (so-called "chloroplast capture" in phylogenetic literature).
2. The subsequent 2nd fast ancient radiation is only reflected in plastome variation within the Japanese archipelago. Also in this case, hybridisation or introgression must have played a role: the various *F. crenata* and/or *F. japonica* haplotype lineages, representing four SLPTs ('IV-JaE', '-JaF', '-JaG', and '-CrC') do not form a well-supported clade but a grade. It seems that at least *F. japonica* introgressed into earlier diverged (and otherwise extinct) species (lineages) that were isolating during this radiation. Since three of the four SPLTs are exclusive to *F. japonica*, *F. crenata* is the more likely (final) introgressor with SLPT IV-CrC representing an (± ancient) *F. japonica* plastome, i.e. a secondary pick-up, or a primary pick-up from the (extinct) sister species of the (extinct) species from which *F. japonica* got its IV-JaG plastomes.
3. The final radiation led to the formation of distinct plastomes (up to six SLPTs, possibly more[9]) within the remaining East Asian species of *Fagus* subgenus *Fagus* (*F. crenata* 'IV-CrD' ↔ *F. lucida* and *F. pashanica* 'IV-ChA' ↔ *F. longipetiolata* 'IV-ChB'). Since both *F. engleriana* specimens carry divergent plastomes formed during this final phase

[9] The plastid diversity within the clade labelled IV-Ch in **Fig. SR2-2** is higher than the diversity observed in all other SPLTs.

(‘IV-EnA’ and ‘IV-EnB’), *F. engleriana* (or its precursors) must have introgressed at a large scale into populations of *F.* subgenus *Fagus* capturing their plastomes, or have formed as an inter-subgeneric hybrid between a continental lineage of *F.* subgenus *Englerianae* (paternal donor) and a precursor/late sibling species of the contemporary East Asian species of *F.* subgenus *Fagus* as maternal donor(s). In either case, *F. engleriana,* likely represents the most recently speciated or established lineage of *F.* subgenus *Englerianae* that has migrated into an area inhabited by various members of *F.* subgenus *Fagus*.

There are two other main deliverables from the assessment of the partitioning effect regarding the evolution of the beech chloroplast genome pool.

First, the differentiation signal supporting the five main lineages (Lineages I–V) as well as the putative species-level plastid types (SLPTs, cf. **Section 2.2**) is trivial: irrespective of the used partitioning scheme or whether a proportion of invariant sites was modelled or not, the according branches received unambiguous BS support (BS = 100). Hence, the lineages and SLPTs can be viewed as the primary biological units when interpreting and studying plastid differentiation on the background of beech evolution.

Second, the differentiation pattern and divergence sequence within Lineage II are signal-wise equally trivial (all intra-lineage, super-SLPT splits with unambiguous BS support); Lineage II reflects a well-structured differentiation process: a sequence of geographic isolation and homogenisation, such as a step-wise allopatric speciation in course of geographic expansion that left its imprint in the modern-day species of the Japanese archipelago (→ **Section 2.2**). In contrast, the hierarchically analogous relationships resolved within Lineage IV can be model-dependent, the signal is diffuse regarding many internal aspects, as expected for fast (ancient) radiations. The exception are several ± terminal inter-SLPT relationships in Lineage IV: (i) a southern plastid lineage including the *F. hayatae* plastomes and the current reference plastome for *F. lucida* (gene bank acc. no. NC_061574); (ii) a taxonomically and geographically disjunct ‘Lineage IV core clade’ comprising individuals from East Asian species of *Fagus* subgenus *Fagus* (Japanese *F. crenata* + Chinese *F. longipetiolata, F. lucida* and *F. pashanica*) and all three sequenced *F. engleriana* plastomes (Chinese *F.* subgenus *Englerianae*) plastomes; which (iii) includes a purely Chinese clade collecting individuals of all Chinese species (including one of the three *F. engleriana* plastomes currently available and the other *F. lucida* plastome, MZ463069).

#### *2.2 Characterisation of main plastid lineages*

##### 2.2.1 Defining 'species-level plastid types' (SLPT)

The massive decoupling between nuclear and plastid phylogenies in wind-pollinated tree genera like beech cannot be explained merely by incomplete lineage sorting (ILS) of ± polymorphic, widespread LCAs (last common ancestors), occasional take-over of one species by a (invading) sibling species ("chloroplast capture" via asymmetric introgression) or genetic stasis (much-decreased genetic drift). While not being sorted by modern-day species and species groups, the divergence observed between and within the here defined plastid lineages is too consistent across the beech species tree to be the product of random-stochastic processes. Cases like *F. sylvatica* vs. *F. caspica* but also the unique *F. hayatae* plastomes or the signal-wise trivial differentiation within Lineage II (see below) suggest a causal correlation between geographic isolation, i.e. allopatric speciation, and plastome divergence patterns as, e.g., reflected in the total number of differing SNPs (**Fig. SR2-4**). Recent introgression and hybridisation can be ruled out. Despite not being sorted at the subgenus or species level, sym- or parapatric species such as *F. japonica* and *F. crenata* do *not* generally share (highly similar) haplotypes. Plastomes of the para-to sympatric Chinese species (*F. engleriana* of *F.* subg. *Englerianae; F. longipetiolata, F. lucida* and *F. pashanica* of *F.* subg. *Fagus*) belong to the same major lineage (Lineage IV) and can be relatively similar (Lineage IV core clade, SLPTs ChA, ChB, EnA and EnB; **Figs SR2-2, SR2-4**), but there is so far no evidence of shared identical or near-identical plastid haplotypes. Thus, it is a valid hypothesis that mid- and deep divergences observed in the plastome trees represent past (allopatric) speciation events overprinted by later radiations and lineage mixing that have led to the establishment of the modern-day species; all of which have had a hybrid/mixed origin.

Today unsorted plastomes may have been specific to geographically restricted species, at a certain point in time. Species were then consumed by or evolved into the modern-day species by fusing with congeners migrating into their area. Asymmetric backcrossing with one of the parents subsequently homogenised the nucleome but differential maternal legacies were passed on via the plastomes.

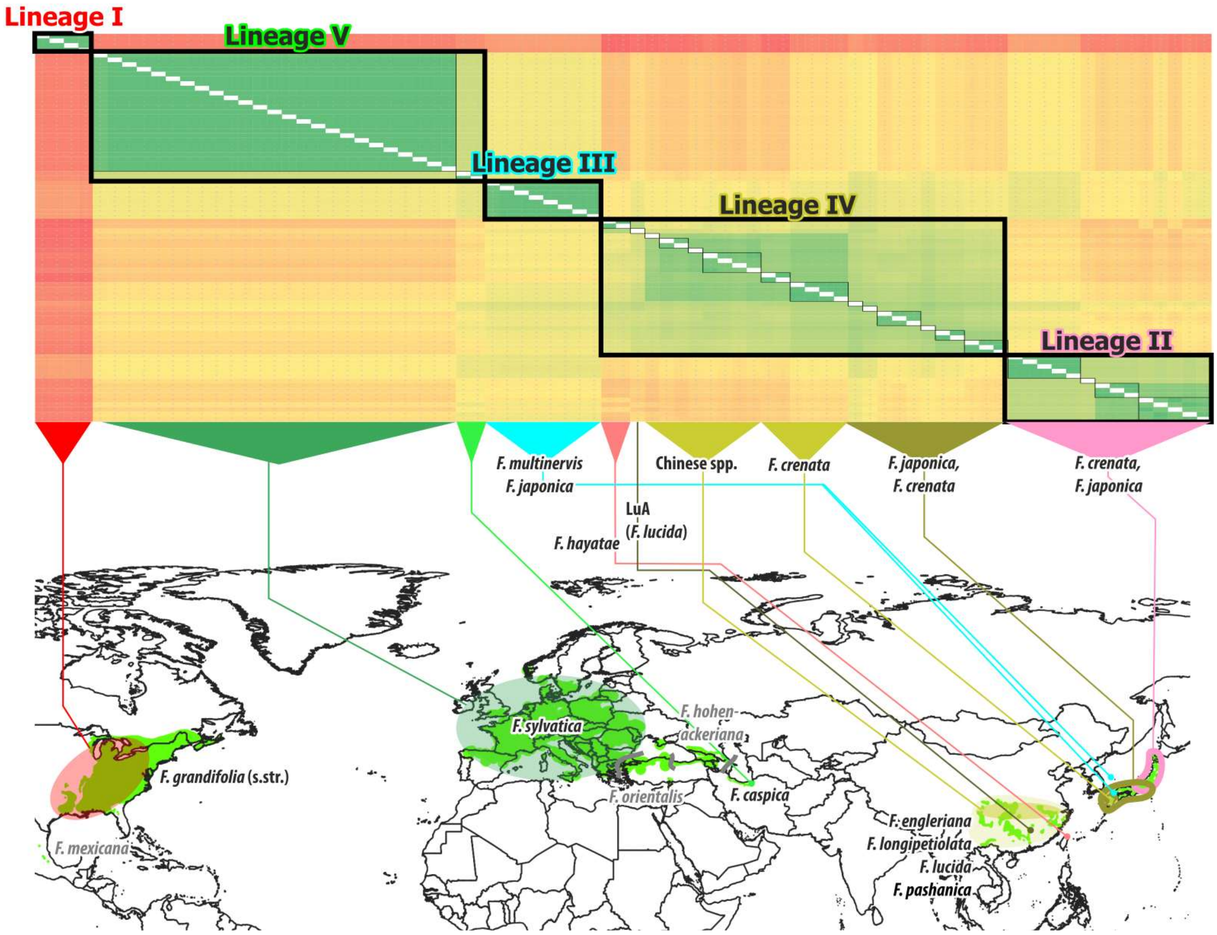


**Figure SR2-4 | Plastome-wide SNP dissimilarity matrix (heat-map) put in a geographic context.** Colour gradient in the heat map (top) ranges from ≤ 4 SNPs difference (dark green) to red >400 SNPs difference (dark red). Thick black boxes: main plastome lineages, thin black boxes: 'species-level plastid types' (see text). No complete plastome data are available for *F. orientalis*, the Caucasian *F. hohenackeriana* and *F. mexicana.* The exact provenance of most Chinese plastomes ('Chinese spp.' group, part of the Lin. IV core clade) is unknown/not reported in the original papers (cf. **Suppl_Genetics.xlsx**, sheet *New and used plastomes*).

On the background of reticulate evolution, the branching patterns in the plastid trees identify 'last common mothers' (LCMs): ancestral species or near-species groups of mother populations that acted as maternal donor(s) for the modern-day taxa and their geographic subgroups and/or precursors. Based on the number of differing SNPs and the phylogenetic framework, the identification of phylogenetic SLPTs – 'species-level plastid types' – within each main lineage is straightforward: unambiguously supported as clades (cf. **Fig. SR2-2**), genetically coherent and distinct (**Fig. SR2-4**) terminal subtrees corresponding to neighbourhoods defined by a prominent edge-bundle ('trunk'; **Fig. SR2-5**), the combination of which is a *sufficient* criterion (in a logical sense) for holophyly. Thus defined SLPTs denote a group of (highly) similar plastid haplotypes within an evolutionary lineage defined by mutually exclusive LCMs. The sequence dissimilarity threshold for recognising a single haplotype or group of similar haplotypes as SLPT can be derived from the recorded plastome differentiation in *F. sylvatica*, which is the only widespread, well-sampled and plastome-wise coherent and sorted species of beech (**Fig. SR2-4**; **SupplGenetics_cpDNA.xlsx**, sheet *PlstmDissim*), that is, representing a species, where the LCM equals the modern species' LCA.[10]

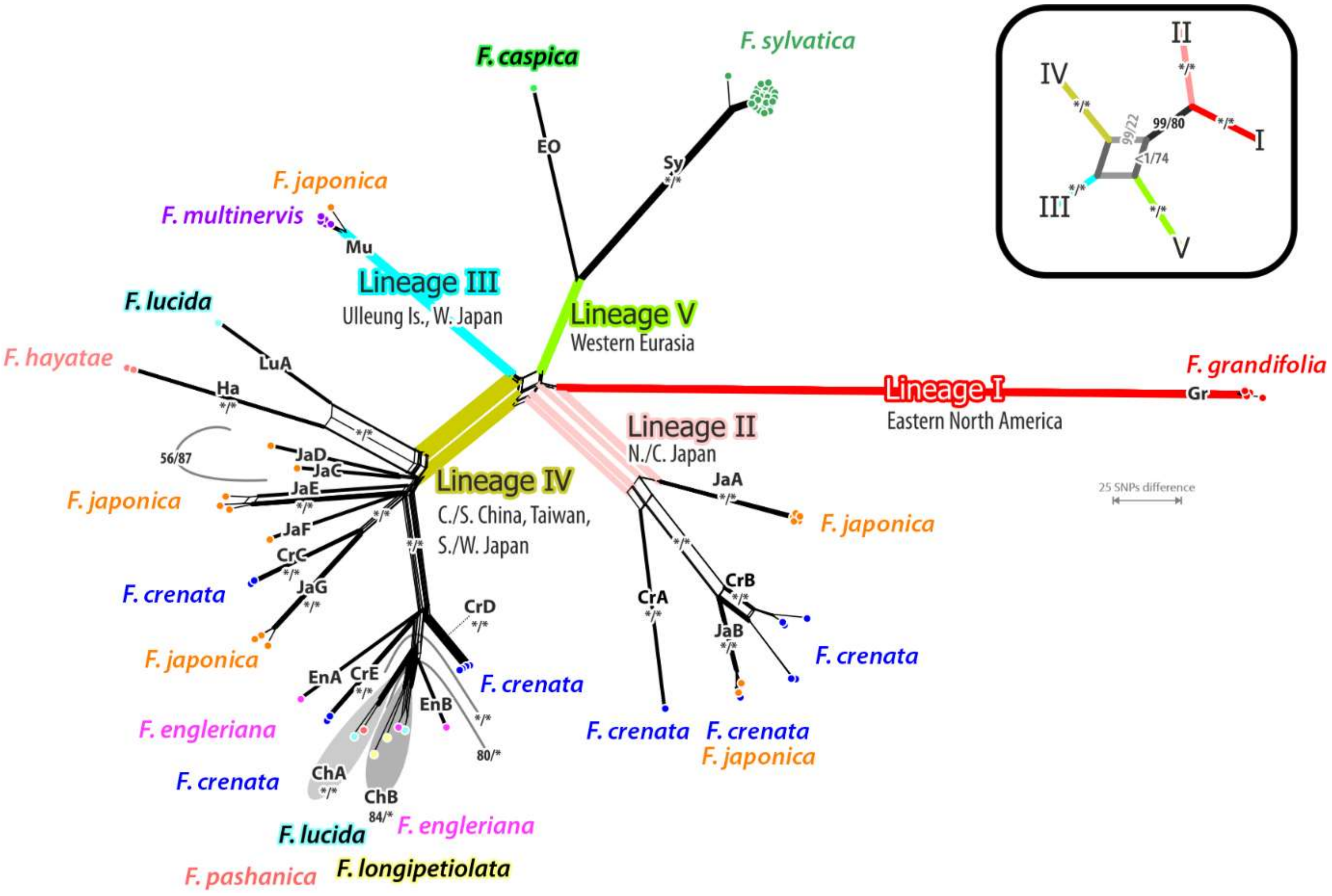


**Figure SR2-5 | Neighbour-Net based on absolute sequence dissimilarity,** expressed by the number of differing SNPs. Numbers at edge bundles and brackets give non-parametric bootstrap support of corresponding branches under maximum likelihood (unpartitioned analyses, asterisks = unambiguous support; see also **Table SR2-1**). * Individuals selected for nuclear sequence analyses of two low-copy genes (**Section 1**).

##### 2.2.2 Composition and geographic patterns of the five main plastid lineages

Of the five main plastid lineages (denoted by Roman numerals), three are sorted by species—Lineage I: *F. grandifolia*, Lineage III (p.p.): *F. multinervis*; and Lineage V: *F. caspica* + *F. sylvatica* (**Fig.SR2-5**). Two, Lineage I and III, comprise (highly) similar haplotypes forming a single SLPT each (I-Gr, III-Mu), and the third, Lineage V, two very distinct SLPTs (V-Ca,

[10] I.e. holophyletic in a strict sense. The complete sorting of nuclear and plastid differentiation signals implies that all modern-day *F. sylvatica* go back to the same parental population/precursor; they share an exclusive to them direct (last) common ancestor. Based on currently available genetic data, the only other strictly holophyletic modern species is (*i*) the N. American *F. grandifolia* (equally widespread, so far only four plastomes available and very limited plastid sequence data; no data on *F. mexicana*), and the highly endemic insular relicts: (*ii*) the Taiwanese *F. hayatae* (in contrast to its continental sister species *F. pashanica*) and (*iii*) the Korean *F. multinervis*.

V-Sy). Of the East Asian beech species, only the Taiwanese *F. hayatae* has a distinctly unique and specific plastome (IV-Ha). The Japanese species *F. crenata* and *F. japonica* comprise six and eight SLPTs from two (Lineage II and IV: CrA–CrF) and three of the main lineages (Lineages II–IV: JaA–JaG, III-Mu). The Chinese species are plastome-wise polymorphic with (at least) three sibling SLPTs in *F. engleriana* (IV-EnA, -EnB, -ChB)*,* one (so far) SLPT in *F. longipetiolata* (IV-ChB) and three in *F. lucida* (IV-LuA, -ChA, -ChB). The plastid composition of *F. pashanica,* the continental counterpart and cryptic sister species of *F. hayatae*, is so far understudied; accession MW846258 (labelled as *F. hayatae*; with a IV-ChA) most probably represents a *F. pashanica* plastome but the provenance of the sequenced tree would need to be verified. It can be expected that more broadly sampled plastome data would reveal more diversity in the Chinese species than currently captured by the available complete plastome data, in analogy what has most recently been found for *Quercus* subsection *Campylolepides*, comprising the East Asian species of *Q.* section *Cerris* (Li et al. 2025).

**Lineage I** is characteristic for the (eastern) North American species *F. grandifolia* of *Fagus* subgenus *Fagus* and represents the most-distinct of all modern-day beech plastomes (**Fig. SR2-4**; **SupplGenetics_cpDNA.xlsx**, sheet *PlstmDissim*) and the initial split within the plastid gene pool of modern beeches (genus *Fagus*; see also according subtrees in Yang et al 2021; Zhou et al 2022). The here sequenced four individuals cover the entire range (up to ~2420 km air-distance) of *F. grandifolia* but show very little intra-lineage divergence (≤ 13 SNP differences per complete plastome) and can be included in a single SPLT: **I-Gr**. There is little plastid data on the disjunct southern-most North American populations, *F. mexicana*, which, so far, show no difference and might represent the same SPLT or a closely related sister SPLT but more comprehensive data is needed to assess their position within Lineage I (**SupplGenetics_cpDNA.xlsx**, sheet *AddHTs_tKmK_ LinI*). Nucleome-wise, *F. mexicana* differs from its (pseudo-cryptic) sister species *F. grandifolia* by retention of ancestral gene variants (**Fig. SR1-9**; Cardoni et al. 2022, data S5). Applying the principle of consilience, the uniqueness of Lineage I plastomes reflects a (northern) North Atlantic-LCM of eastern North American beeches in contrast to the northern North Pacific-LCM shared by current-day Eurasian beeches and a divergence within genus *Fagus* that must have preceded the establishment of the modern-day subgeneric lineages. This hypothesis is in perfect fit with the fossil record (→ **Section 4**).

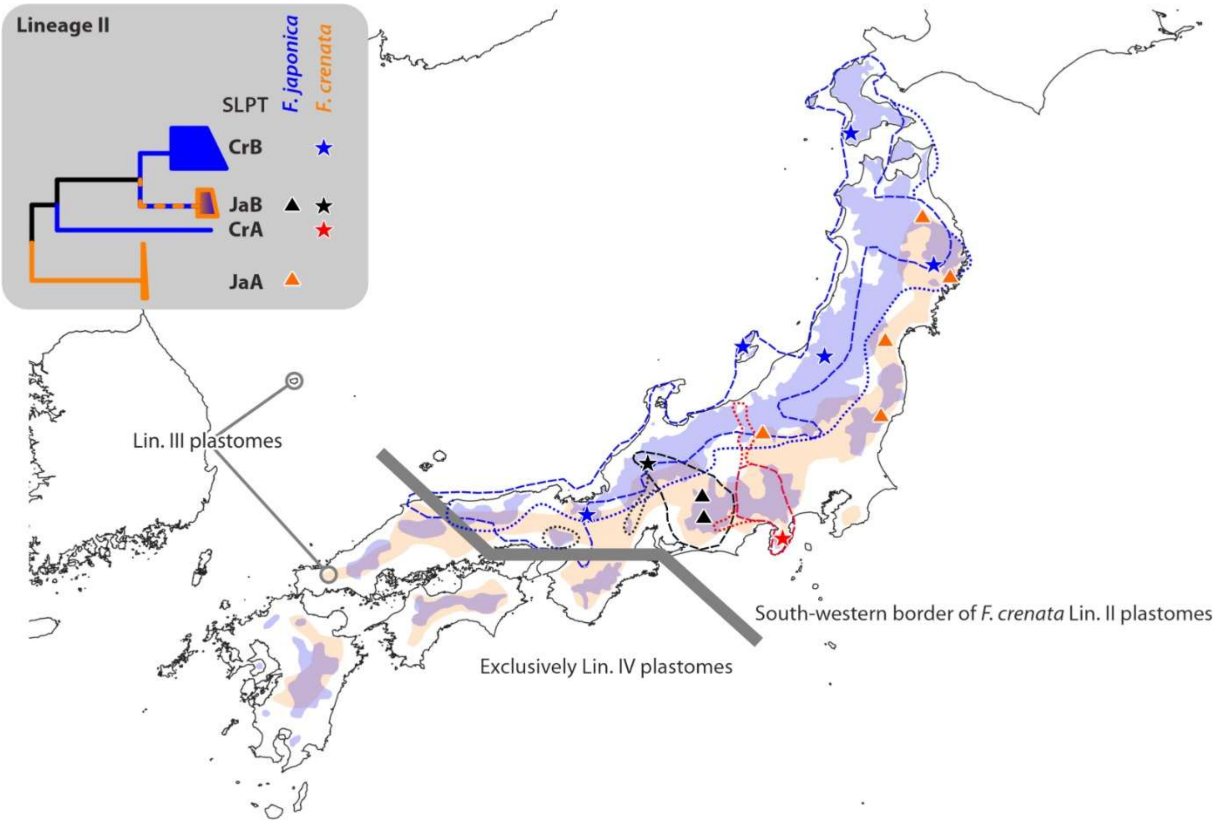


**Figure SR2-6 | Modern-day geographic distribution of Lineage II plastomes and their species-level plastid types (SPLTs).** Provenances of completely sequenced plastomes indicated by stars (*F. crenata* individuals) and triangles (*F. japonica*). Stippled lines give the distribution of according *F. crenata* haplotypes based on Fuji et al. (2002), dotted lines their emendation by Tsumura & Suyama (2015), who covered the entire range of *F. crenata* in Japan. Only point data available so far for *F. japonica*.

**Lineage II** is exclusively shared by *F. crenata* (*F.* subg. *Fagus*) and *F. japonica* (*F.* subg. *Englerianae*) individuals from central and north-eastern Honshu and Hokkaido (**Figs SR2-4, SR2-6**) and represents the 2nd diverging plastid lineage. Lineage II includes four SPLTs (≤ 161 SNP differences per complete plastome), one exclusive to *F. japonica,* **II-JaA** in north-eastern Honshu (para- to sympatric with *F. crenata* carrying Lin. IV plastomes), and two allopatric SLPT exclusive to *F. crenata*: **II-CrA**, geographically restricted (= HT E), and **II-CrB** (HTs A–C of Fujii et al. 2002), the most widespread of all *F. crenata* SLPTs ranging from central Honshu to southwestern Hokkaido along the mountain chains and slopes facing the Sea of Japan. The 2nd *F. japonica* SPLT, **II-JaB**, is shared by sympatric *F. crenata* in central Honshu (**Fig. SR2-6**; HT D of Fujii et al. 2002), and can only represent a relatively recent (see also **Section 3.3**) introgression of *F. crenata* (precursors) into the *F. japonica* species lineage. Based on its current-day distribution (**Fig. SR2-6**), we interpret Lineage II as the plastome originally associated with a group of high-latitude, potentially cross-Beringian beeches and the, or one of the, original plastome(s) of *Fagus* subgenus *Englerianae*. In contrast to Lineage IV, the plastid differentiation in Lineage II reflects a clearly structured sequence of divergences, subsequent ± ancient allopatric speciation events, resulting in a clear phylogenetic sorting of mutational patterns and accordingly high support of all branches in the subtree (BS ≥ 98 in all analyses): (*i*) west-east vicariance into a Pacific-bound species (II-JaA) and a species focussed on the Sea of Japan side, followed by (*ii*) migration and isolation of the southernmost populations (II-CrA) and a (*iii*) final speciation event between a lineage today distributed in central Honshu (II-JaB) and a northbound, ± montane species (II-CrB), whose descendants form the northern Japanese populations of modern-day *F. crenata* thriving on the side of the Sea of Japan in the mountains of central and north(west)ern Honshu into Hokkaido.

With respect to the altitudinal, ecological and morphological differences between modern-day *F. crenata* and *F. japonica* (Hara 2010), the SLPT II-CrB is the most likely candidate for the plastome of the last common mother of modern-day *F. crenata* populations, the *Crenata*-LCM the primordial *F. crenata* 'Eve(s)'; a plastome the *F. crenata* 'Eve' picked up when it migrated into the original area of the early *Fagus* subgenus *Englerianae* (N.E. Siberia, Kamchatka, northern Japan; see main-text figs 9 and 10 and **Section 4**).

**Lineage III** comprises one SLPT, **III-Mu** (≤ 20 SNPs), and is exclusively shared by *F. multinervis* from Ulleungdo, a relatively young volcanic island in the Sea of Japan ~130 km away from the South Korean mainland at the northern margin of the Tsushima Basin, and a single individual of *F. japonica* (ja31) from a population in western Honshu at the southern side of the Tsushima Basin (**Fig. SR2-6**). It clearly represents a relatively early divergence event and probably a 'genetic relict' predating differentiation within the more diverse lineages (Lineages II and V; → **Sections 2.3.1** and **3.3**; see also main-text fig. 9). The haplotypes from Ulleungdo stored in gene banks are near-identical to sequences generated here (0–4 SNP difference between all *F. multinervis* samples), while the Japanese haplotype on the opposite side of the Tsushima Basin (~350 km air-distance to Ulleungdo, no further contemporary stepping stones for plant and seed dispersal) differs by an additional 16 SNPs. Lineage III is phylogenetically intermediate between East Asian Lineage IV and the Western Eurasian Lineage V (**Figs SR2-2, SR2-3, SR2-5**), a reflection of the position of Ulleungdo within the North-East Asia region (→ **Section 4**): the island lies on the cross-roads between the earliest (Oligocene) members of the Western beech lineage within *Fagus* subgenus *Fagus,* the West-Eurasian clade (Lin. V plastomes), *F.* subgenus *Englerianae* and the earliest East Asian members of *F.* subgenus *Fagus* (**SupplFossilTable.xlsx**, sheet *main list*).

Despite being geographically highly restricted and their accordingly small active population size compared to their Japanese sisters and cousins, Lineage III plastomes are the least evolved of all contemporary beech plastomes: they are overall the least dissimilar to any other lineage (**Table SR2-2**) and the mutational patterns in signal-rich regions are typically closest to the genus' or Eurasian plastome (modal) consensus (→ **Section 2.3**). Notably, the area today forming the Tsushima Basin and the rest of the Sea of Japan has not only been at the cross-roads but also the centre of the long-lasting biodiversity hot spot of beeches in North-East Asia, thus, a likely refuge for the survival of a genetic relict. The lack of genetic drift observed in Lineage III plastomes can be attributed to the lack of bottlenecks and geographic stasis, and a much larger active population size until or into the Pleistocene, especially during global low sea level situations: However, despite being exclusive to species of *Fagus* subgenus *Englerianae*, the phylogenetic position of Lineage III plastomes (part of the same lineage as Lin. IV and Lin. V plastomes) make it an unlikely candidate for the original plastome of *F.* subgenus *Englerianae*.

**Table SR2-2 | Coherence and differentiation of major plastome lineage in beech.** Above diagonal, minimum number of differing SNPs between plastomes of either lineage; below diagonal, maximum number; diagonal, dissimilarity range within each lineage preceded by the number of included complete chloroplast genomes. Genus-wide inter-lineage minima highlighted by grey backgrounds.

| | Lineage I | Lineage II | Lineage III | Lineage IV | Lineage V |
|---|---|---|---|---|---|
| **Lineage I** | 4: 5–13 | 357 | 360 | 369 | 384 |
| **Lineage II** | 413 | 14: 0–161 | 215 | 201 | 222 |
| **Lineage III** | 372 | 264 | 8: 0–20 | 193 | 210 |
| **Lineage IV** | 440 | 315 | 262 | 28: 0–231[a] | 170 |
| **Lineage V** | 409 | 290 | 237 | 316 | 27: 0–173[b] |

[a] Maximum intra-lineage dissimilarity found between SLPT IV-EnA (Lin. IV core clade, 100–118 differing SNPs to other Lin. IV core clade plastomes) and IV-Ha (generally most-distinct and genetically drifted Lin. IV plastomes: [135–]180–231 differing SNPs to other Lin. IV plastomes).
[b] Maximum intra-lineage dissimilarity only between plastomes of SLPT V-Sy (data on 25 individuals) vs. V-Ca (171–173 differing SNPs, 2 individuals with identical plastomes).

**Lineage IV** is the most diverse plastid lineage (14 SLPTs differing by ≤ 231 SNPs), and can be found in all East Asian species of *Fagus* subgenus *Fagus* and is shared with the Chinese *F. engleriana* (late diverging HTs, cf. **Fig. SR2-2**; **IV-EnA**, **IV-EnB**) as well as the Japanese *F. japonica* (early diverging HTs; **IV-JaC–G**) of *F.* subgenus *Englerianae*. Lineage IV is the only lineage showing a combination of early isolation of several subtypes, a fast ancient radiation (IV-JaC, -JaD, -JaE; **IV-Ha + IV-LuA; IV-CrC** + IV-JaG), and a late radiation event (Lin. IV core clade: **IV-CrD/CrE**, **IV-ChA/ChB**, IV-EnB; **Fig. SR2-7**).

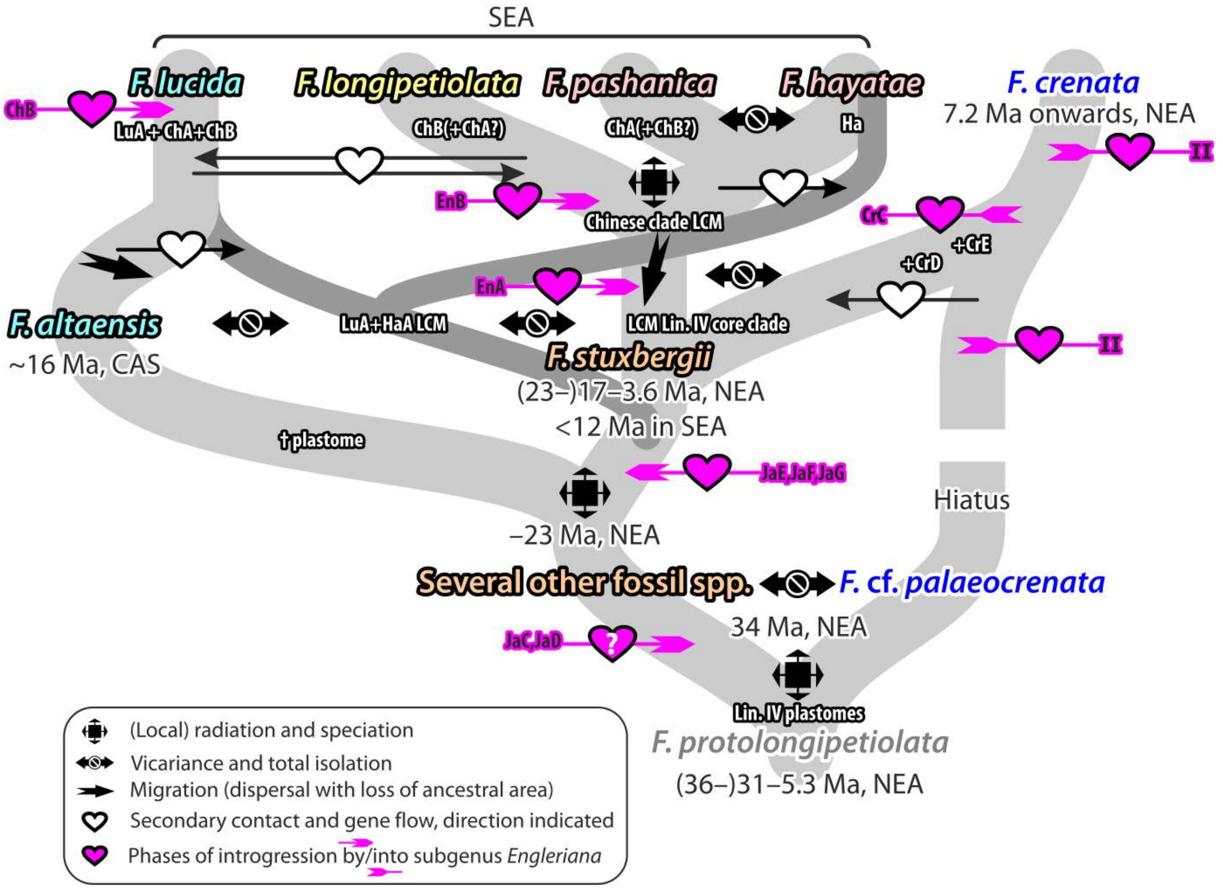


**Figure SR2-7 | Possible spatio-temporal framework for Lineage IV plastomes.** Fossil species with affinity to modern-day East Asian species embedded in a simplified species network of East Asian species of *Fagus* subgenus *Fagus*; same colouring of names reflect phenotypic affinities. Abbrev.: LCM = last common mother; NEA = North-East Asia; SEA = southern East Asia. "LuA" etc. refer to SLPT (species-level plastid types) within Lineage IV. For an explicit dating of plastid and nuclear divergences see main-text, **SupplDating.xlsx**, sheet *final dating* and **Section 3.3**.

Notably, all individuals of Chinese species studied so far (*F. engleriana, F. longipetiolata, F. lucida, F. pashanica*) carry Lineage IV plastomes, either shared between species (IV-ChA, IV-ChB) or of distinct evolutionary sources (IV-LuA, IV-EnA, IV-EnB). Thus, the working hypothesis for any plastid-phylogenetic or population-level analysis in China is that *none* of these species can be traced back to an exclusive last common mother; notably, species that are morphologically distinct even in sympatric stands, show a high genetic coherence in their nucleomes and no sign of ongoing hybridisation (Shen 1992, Denk 2003, Jiang et al. 2022, Cardoni et al. 2022; this study). As in the case of its cousin, the oaks (*Quercus*; all plastid Sanger-sequence data generated so far, see e.g. Simeone et al. 2016, 2018, Vitelli et al. 2017; see Li et al. 2025 for broad-sampled complete plastomes), plastid data of Chinese beeches, whether based on individual gene markers or complete chloroplast genome sequences, are useless to infer intra-generic and inter-species relationships for the Chinese species. Likewise, any broadly sampled population-scale study of a single species using plastid haplotypes must include at least several sympatric and parapatric individuals of the other three species to be able to interpret the found haplotype differentiation patterns. Only the non-Chinese (insular) species, *F. hayatae* from northern Taiwan (IV-Ha) and the Japanese *F. crenata* (IV-CrC–CrE), can be characterised by distinct, exclusive SLPTs forming high-supported clades in phylogenetic trees and according neighbourhoods in distance-based networks (**Figs SR2-2, SR2-5**).

There is currently no sufficient, georeferenced chloroplast haplotyping data on the Chinese species to assess patterns of geographic sorting within their Lineage IV plastomes. For the Japanese archipelago, comprehensive plastid haplotype data are available (Fujii et al. 2002, Tsumura & Suyama 2015), and can be put in the context of the here defined SLPTs and their phylogenetic context (**Fig. SR2-8**).

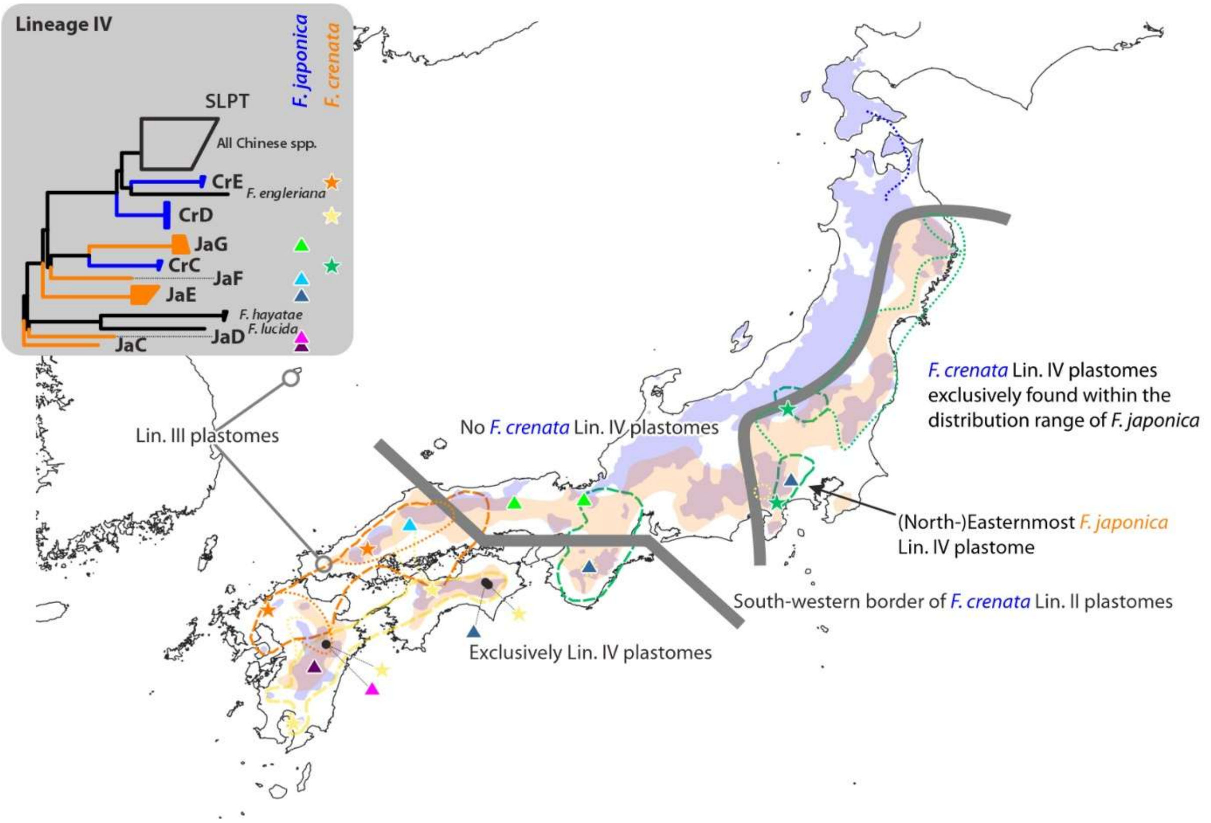


**Figure SR2-8 | Modern-day geographic distribution of Lineage IV plastomes in Japan.** *Fagus crenata* with late-diverging Lin. IV plastomes (Lin. IV core clade comprising all continental East Asian plastomes) are limited to westernmost Honshu and the southern islands of the Japanese archipelago. Early diverging Lin. IV plastome of both *F. crenata* and *F. japonica* can be sym- or parapatric with Lin. II plastomes.

In contrast to Lineage II-carrying *F. crenata* (see above, **Fig. SR2-6**), *F. crenata* with Lineage IV plastomes are chiefly sym- to parapatric with *F. japonica*. In eastern and north-eastern Honshu, the earliest diverging *F. crenata* Lineage IV SLPT, IV-CrC, covers the entire range of *F. japonica* in this area, none of which showed its sister SLPT, IV-JaG. In the southern-most

*F. japonica* individual sequenced from this area, the IV-CrC area overlaps with IV-JaE, an isolated, early diverged SPLT within Lineage IV, and in the remainder of eastern Honshu, it appears to be sympatric with II-JaA, the earliest diverging SLPT within Lineage II. Sympatric sister SPLTs IV-CrC and -JaG can be found in central Honshu. The south-western border of *F. crenata* Lineage II plastomes coincides with the transition zone between *F. crenata* SLPT IV-CrC and SLPTs that form part of the Lineage IV core clade (IV-CrD and -CrE), a clade that also includes all completely sequenced plastomes of the continental, Chinese species *F. engleriana, F. longipetiolata, F. lucida* and *F. pashanica* with the exception of the gene bank reference plastome for *F. lucida* (IV-LuA), originating from south-central China (Hunan). IV-CrD and -CrE are allopatric with the exception of central Kyushu, where they marginally overlap. The range of the northern Ryushu and western-most Honshu IV-CrD encompasses or is close to the localities of plastid-wise ± distantly related *F. japonica*, the population carrying Lineage III plastomes closely similar to those of *F. multinervis* (III-Mu) and the individual with the strongly isolated SLPT IV-JaF, a plastid type originating in the second radiation wave within Lineage IV. Its southern counterpart, IV-CrE from the mountains of Kyushu and Shikoku, is para- to sympatric with early diverging *F. japonica* Lineage IV SLPTs: IV-JaC and -JaD in Kyushu and IV-JaE in Shikoku. The latter, IV-JaE appears to be stretching along *F. japonica* populations along the entire southern coast of Honshu (**Fig. SR2-8**). Putting together, the strong geographic sorting observed in Japan within the chloroplast genomes of *F. crenata* and *F. japonica* can only be the legacy of allopatric speciation processes, most of which predate the manifestation of the modern-day species, and later (re-)migration of distantly related mother populations (extinct species).

With respect to the divergence observed between sister SLPTs in *F. crenata* and *F. japonica* (II-CrB + II-JaB, the latter being the only SLPT shared by both species, vs. IV-CrC + IV-JaG) and the allopatry of ± close vs. sym-/parapatry of ± distant SLPTs, large-scale (sub-)recent introgression can be ruled out as explanation for the complex geographic structuring of the plastid gene pool across the Japanese archipelago. A recent massive and repeated introgression or large-scale hybridisations should also have left some detectable imprint in the nucleomes of both species but any diverse-enough nuclear gene region sequenced so far reflects the primary dichotomy into the two subgeneric lineages.

The high plastid divergence observed in Japanese Lineage IV-carrying beeches coincides with a high phenotypic diversity that persisted into the Pliocene of the Japanese archipelago, covering both ancestral, or taxonomically ambiguous, as well as modern-day phenotypes; fossil species that comprise probably the maternal donors of the various Lineage IV SLPTs. The Lineage IV plastomes of *F. japonica* and *F. crenata* appear to be the legacy of extinct Japanese species and beech lineages of North-East Asia, some of which (IV-CrC and IV-CrD) were sister species (the eastern, oceanic counterparts) of the precursors or were the precursors of the modern-day continental Chinese species of *Fagus* subgenus *Fagus*. The relative genetic coherence of the mostly Chinese Lineage IV core clade in conjunction with the mutational patterns observed in selected plastid gene regions (see following section) would fit to the hypothesis that a single lineage of Lineage IV carrying beeches migrated from Japan (North-East Asia) into (central) China (southern East Asia; → **Section 4**; main-text figs 8, 10), where it diversified further and disintegrated into locally restricted species or clusters of mother trees. Some of these first modern-day Chinese beeches were apparently introgressed by the modern members of *Fagus* subgenus *Englerianae* (SLPTs IV-EnA and -EnB), while others (SLPTs IV-ChA and -ChB) were the precursor or maternal donors of the modern-day Chinese species of *F.* subgenus *Fagus*: *F. longipetiolata, F. lucida* (p.p.) and *F. pashanica.*

The Western Eurasian **Lineage V** is the only geographically *and* phylogenetically/taxonomically perfectly sorted plastid lineage. Its two strongly divergent SLPT (**V-Ca** and **V-Sy**; varying between 170–173 SNP differences) reflect both the geographic distance between the sampled individuals as well as their taxonomy and (nuclear-)phylogenetic distance (*F. sylvatica* V-Sy vs. *F. caspica* V-Ca) and the evolutionary history of Western Eurasian beeches (as discussed in Cardoni et al. 2022, Schulze & Grimm 2022, Denk et al. 2024).

Based on the available data (gene banks, Ulaszewski et al. 2021, and new data), *F. sylvatica* plastomes[11] are near invariable across the entire sampled range (north-central Spain to north-western Greece and eastern Bulgaria) with the exception of our central Italian individual sy47, and, to a lesser degree, an individual from north-western Greece, sy48. The uniqueness of sy47 can be explained by its provenance: it comes from a recognised old relict forest in the central Apennines. Northern Greece has also been suggested to represent a Pleistocene relict area and refuge for beech trees. Work in progress hints to an increased plastid divergence in this region (Hatziskakis et al. 2009; Papageorgiou et al. 2014; Tsiripidis et al. 2024). The high similarity of *F. sylvatica* plastomes outside these two refugial areas fits to the hypothesis that most modern-day beeches of Europe stem from a single population that survived the Last Glacial Maximum (LGM) and re-colonised Europe within the last 7000 years (Magri et al. 2006, 2008). Their similarity also implies that *F. sylvatica* is one of the few strictly holophyletic species

[11] And including specimen representing the alleged hybrid taxon *F.* × *moesiaca*.

within the genus, i.e. all modern-day beeches of Europe go back to a shared and exclusive to them common ancestor.[12] The genetic homogeneity of the European *F. sylvatica* appears to have only one living analogue: *F. grandifolia* in eastern North America that covers essentially the same climatic envelope and an accordingly large area (Maycock 1994; Grimm & Denk 2012). If the lack of plastid and nuclear intra-species diversity in *F. sylvatica* is a product of Pleistocene bottleneck situations, the same would apply to *F. grandifolia*.

The plastomes of *F. caspica* (new species; formalised in Denk et al. 2024) are here represented by two individuals collected in Gorgan County, north-eastern Iran (Golestan province). Earlier morphological and genetic results (Denk 1999, Gömöry & Paule 2010) indicate that these populations represent a species on their own right, which has been recently confirmed by high-resolution nuclear genotyping (Kurz et al. 2023 and Denk et al., 2024). The Iranian beeches are the most distant relatives of *F. orientalis* and its sister species, *F. sylvatica,* within the Western Eurasian beech lineage (Cardoni et al. 2022, Denk et al. 2024). Unfortunately, the currently available plastid data on *F. orientalis* and its eastern counterpart *F. hohenackeriana* (mostly *trnL* + *trnL-trnF* IGS; Paffetti et al. 2007) does not provide enough resolution for resolving the two SLPT within Lineage V (**SupplGenetics_cpDNA.xlsx**, sheet *GB-HTs_trnTLF*) but accessions obtained from *F. orientalis* (**Table SR2-3**; A. Papageorgiu and co-workers, unpublished data) show haplotypes not covered in the complete plastomes so far. Complete plastomes from these areas may reveal the presence of further Lineage V SLPTs within and diagnostic for the remaining Western Eurasian beech species (*F. orientalis*, *F. hohenackeriana*, and, potentially, the beeches of the isolated Nur Mts in S. E. Turkey).

**Table SR2-3 | Potential additional SLPT – species-level plastid types – within Lineage V.** Mutational patterns diagnostic for Lineage V plastomes and their two here identified SLPTs, *F. caspica* vs. *F. sylvatica* plastomes, highlighted by green shading.

| | | | *trnH-psbA* IGS | | | | | | *matK* gene | | | | | downstream *trnK* intron | | |
|---|---|---|---|---|---|---|---|---|---|---|---|---|---|---|---|---|
| **Species** | | **Region** | **60** | 128–139 | 169–185 | 240–251 | **362** | **376** | **377** | **540** | **560** | 977 | **1248** | **2095–2108** | *2188–2198* | **2369** |
| Genus consensus/ancestral sequence | | | C | $A_9$ | $T_4$-TA-$T_8$ | $A_9$-TG | T | T | T | T | T | T | G | $A_9$-CC | **$A_{10}$**-T | A |
| *F. caspica* | New data only | N. Iran | C | $A_{10}$ | $T_4$-TA-$T_8$ | $A_7$-**GA**-TG | T | **C** | T | **C** | **C** | T | **A** | $A_9$-CC | $A_9$-T | A |
| Indet. (*F. orientalis* s.l.) | AY042396 | ? | N/D | N/D | N/D | N/D | N/D | N/D | N/D | N/D | N/D | N/D | N/D | $A_{10}$-CC | **$A_{10}$**-T | |
| *F. orientalis* (s.str.) | MT136566, MT114335 | Romania | N/D | $A_9$ | $T_4$-TA-$T_8$ | $A_7$-**GA**-TG | T | T | N/D | N/D | N/D | g | **A** | N/D | N/D | N/D |
| *F. sylvatica* (s.str.) | Complete plastomes | Europe | **T** | $A_9$ | $T_4$-T**C**-$T_8$ | $A_7$-**GA**-TG | **G** | T | **C** | **C** | **C** | T | **A** | $A_{10}$-CC | **$A_{10}$**-T | **C** |
| | Gene bank accessions | Europe | **T** | $A_9$ | $T_4$-T**C**-$T_8$ | $A_7$-**GA**-TG | **G** | T | **C** | **C** | **C** | T | **A** | $A_{10}$-CC | **$A_{10}$**-T | **C** |

Diagnostic for Lineage V plastomes
… for *F. caspica* plastomes
… for *F. sylvatica* plastomes
Possible sequencing/editing artefact

[12] The genetic mosaic found in the nuclear-encoded rDNA arrays is, hence, inherited from a polymorphic ancestor.

#### 2.3 Evolution of selected plastome regions

Multigene and phylogenomic analysis typically produce high branch support and may propagate and enforce first-level error (Delsuc et al. 2005). This is aggravated when using concatenated data matrices to infer trees for plant groups that are affected by deep and flat reticulate evolution such as beech. This was also the case for the nuclear 28-gene data of Jiang et al. (2022): the authors inferred a (near-)fully resolved, perfectly sorted but aspect-wise incorrect species tree (Jiang et al. 2022, fig. 1) by concatenating nuclear genes with strongly incongruent histories and differential variation. Accordingly, all branches in their multi-species coalescent (Jiang et al. 2022, fig. 3) were poorly supported, albeit showing the same inter-species relationships within genus *Fagus*.[13] Given the overall low-diversity of beech plastomes, we hence explored the mutation patterns in selected regions (listed according to their position in the plasmid):

- three commonly used plastid 'barcodes': the *trnH-psbA* intergenic spacer (IGS), the *trnK* intron (the comprised *matK* gene is near-invariable), and the *trnL* intron + *trnL-trnF* IGS;
- the five partitions with the highest number of segregating sites (SNPs; **SupplGenetics_ cpDNA.xlsx**, sheet *cp gene stats*): *trnK-trnQ* IGS, *rpoC2* gene, the *accD* gene and flanking intergenic spacers (*rbcL-accD*; *accD-psbI*),[14] *ndhF* gene, and the *ycf1* gene (the latter is the longest, and mutation-richest, chloroplast gene region and partition);
- in addition, the five partitions with the highest *Pi*-values identified by sliding-window analysis (**ODA**, file *SlidingWindowResults.xlsx*), very short intergenic spacers with a single or few high-divergent mutation patterns: *trnY-trnE* IGS, *trnG-trnfM* IGS, *rrn4.5-rrn5* IGS, *ndhD-psaC* IGS, and *ndhI-ndhA* IGS.

The set of partitions cover all three main regions of the plasmid ring, the large single-copy unit (LSC), the short single-copy unit (SSC) and the (identical) inverted repeat region(s) (IRa and IRb). The mutation patterns of the selected plastid regions are tabulated in **00_Bits.xlsx**, included in the ODA.

##### 2.3.1 General fit of individual mutations with five main lineages

Most mutation patterns in the selected plastid regions are congruent with the splitting of the plastomes into five main lineages (**Table SR2-4**): the great majority of observed character splits

[13] Both trees differ, however, in the topology of their outgroup subtrees.
[14] Due to a now corrected annotation error, we initially overlooked the *accD* gene and treated it as part of an artificial *rbcL-psbI* IGS.

are fully compatible with the inferred complete plastome tree. Non-lineage-sorted patterns notably include all observed inversions in the selected partition sample or are linked to mononucleotide repeat length-polymorphism and probable backmutations (predominantly C↔T transitions in distantly related lineages). While not sorted across the entire tree or within the main lineages, these patterns can be haplotype-conserved, thus, would have a certain diagnostic value or could be informative for establishing phylogeny-flat biogeographic patterns (**Fig. SR2-9**).

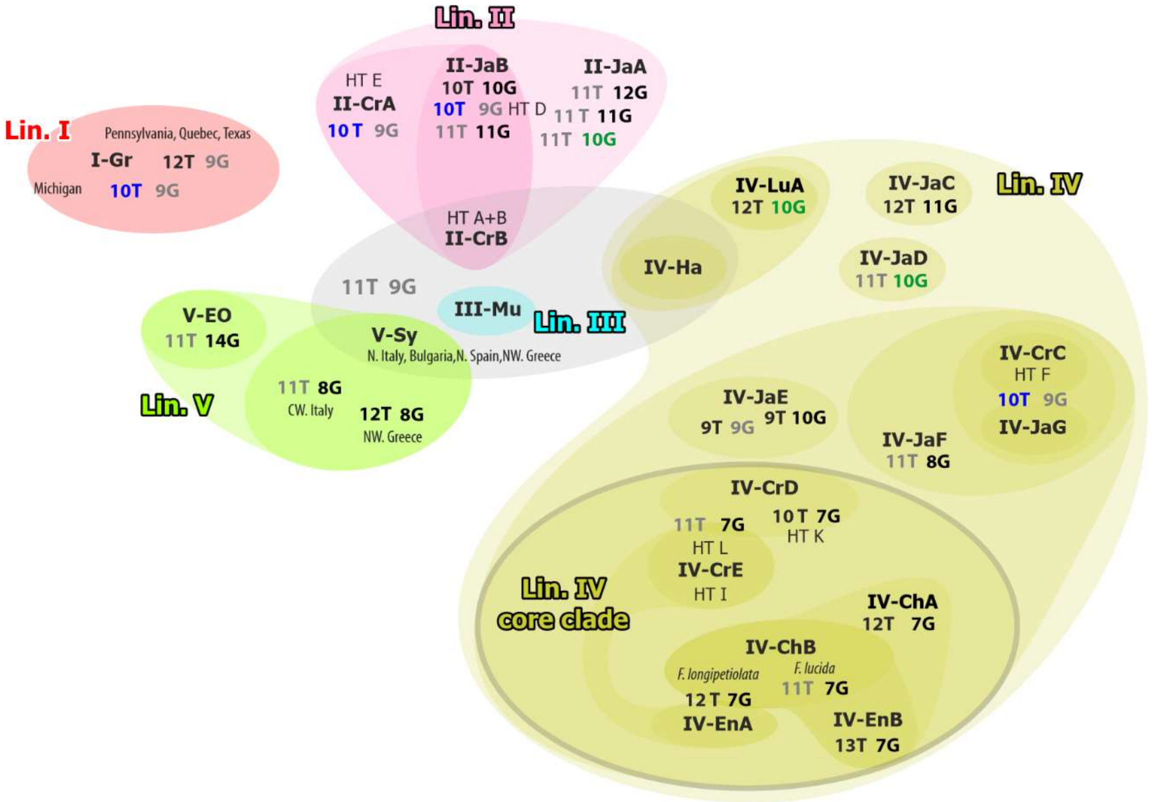


**Figure SR2-9 | Example for signal-depth in plastid microsatellite markers ccmp4 and ccmp7** (cf. **ODA**, file *00_bits.xls*) **traditionally used in species-level haplotype studies**, located in the *atpF* intron and *atpB-rbcL* IGS of beech plastomes. The combination of these two markers may be informative to map species-level plastid types (SLPTs) within Lineage IV and, possibly, intra-SLPT patterns in *F. sylvatica* (SLPT V-Sy) and *F. grandifolia* (I-Gr) but would fail at the (quasi-)species-level. Note the wide sharing of the putative ancestral combination (11T + 9G, in grey) and its first modifications (→10T, blue+grey; →10G, green font).

The mutation-rich partitions (shaded grey in **Table SR2-4**) include up to 17 lineage-conserved mutation patterns (mostly SNPs), the majority of which discriminate between the North American Lineage I (*F. grandifolia*) and the Eurasian samples.

**Table SR2-4 | General statistics regarding the number and diagnostic stringency of mutation patterns in 13 selected plastid regions**. #DMP – Number of distinct mutation patterns (SNPs, mononucleotide repeats, indels, other length polymorphic patterns); #ST – thereof singletons (uninformative); #LCP – thereof lineage-characteristic: uniquely derived and conserved within a main lineage/species; #NLS – not lineage-sorted: convergent, stochastically distributed patterns, incompatible with main plastid lineages. For more statistics see **Pi values by gene etc. .xlsx**

| Partition, gene | Type | Region | Sites | #DMP | #ST | #LCP | | | | | | | | | #NLS | | | Remark |
|---|---|---|---|---|---|---|---|---|---|---|---|---|---|---|---|---|---|---|
| | | | | | | | I | II | III | IV | | V | | | All | Mult.X | SNP | |
| | | | | | | | | | | | Ha | | Sy | Ca | | | | |
| *trnH-psbA* | IGS | LSC | 422 | 14 | 2 | 8 | 3 | 1 | 1 | 0 | 0 | 1 | 2 | 1 | 0 | | | |
| *trnK*i | Intron | LSC | 1,023 | 17 | 3 | 8 | 3 | (G) | 0 | 0 | 3 | 0 | 1 | 0 | 1 | 1 | 0 | (G) – Some with genus consensus |
| *trnL*i | Intron | LSC | 540 | 8 | 2 | 4 | 0 | 1 | 0 | 0 | 1 | 1 | 0 | 0 | 1 | 1 | 0 | |
| *trnL-trnF* | IGS | LSC | 194 | 7 | 1 | 0 | 0 | 0 | 0 | 0 | 0 | 0 | 0 | 0 | 0 | | | |
| *trnK-trnQ* | IGS | LSC | 2,199 | 70 | 15 | 28 | 13 | 3 | 4 | 2 | 3 | 0 | 1 | 3 | 3 | 3 | 0 | |
| *rpoC2* | Coding | LSC | 4,131 | 47 | 7 | 18 | 8 | 3 | 0 | 0 | 1 | 0 | 2 | 3 | 1 | 0 | 1 | |
| *rbcL-psbI* | Mixed* | LSC | 2,877 | 47 | 11 | 16 | 6 | 1 | 2 | 0 | 1 | 0 | 3 | 2 | 1* | 0 | 0 | * Inversion within pseudo-stem |
| *ndhF* | Coding | SSC | 2,266* | 44 | 11 | 18 | 5 | 3 | 1 | 2 | 6 | 1 | 0 | 2 | 1 | 0 | 1 | * Excl. overlap with *ycf1* |
| *ycf1* | Coding | SSC | 5,700 | 117 | 26 | 44 | 17 | 2 | 3 | 4 | 5 | 4 | 7 | 1 | 5* | 0 | 3 | * Incl. only inversion |
| *trnY-trnE* | IGS | LSC | 59 | 2 | 0 | 1 | 0 | 0 | 0 | 1 | 0 | 0 | 0 | 0 | 0 | | | |
| *trnG-trnfM* | IGS | LSC | 185 | 7 | 1 | 4 | 2 | 0 | 0 | 3 | 0 | 0 | 0 | 0 | 1 | 0 | 1 | |
| *rrn4.5–rrn5* | IGS | IR | 257 | 2 | 1 | 0 | | | | | | | | | 1* | 0 | 0 | * Inv. or insert within pseudo-stem |
| *ndhD-psaC* | IGS | SSC | 136 | 7 | 0 | 4 | 0 | 1 | 0 | 0 | 1 | 0 | 2 | 0 | 1* | 0 | 0 | * Inv. within pseudo-stem |
| *ndhI-ndhA* | IGS | SSC | 82 | 5 | 0 | 3 | 0 | 0 | 0 | 1 | 0 | 0 | 1 | 0 | 0 | | | |

* Including the *accD* gene and flanking intergenic spacers; the *accD* gene was overlooked in the original gene annotation.

On the background of the phylogenetic tree, 'genetic synapomorphies' (in a Hennigian sense) could thus be identified: uniquely derived sequence patterns identifying a plastome as member of one of the five main lineages, unambiguously supported as clades/taxon bipartitions in all phylogenetic tree inferences. In addition, SLPTs within each lineage could be diagnosed by autapomorphic SNPs and other mutational patterns. 'Genetic synapomorphies' and lineage- or SLPT-sorted suites of mutations are of high-diagnostic value for densely sampled population-scale studies as they allow to identify the plastome lineage, or SLPT, at hand of single gene regions/ relatively short DNA sequences ("fragments"). Observed haplotype diversity within a species can then be put in a more general evolutionary-phylogenetic context (see example of *F. crenata* haplotypes; **Figs SR2-6, SR2-8**; Li et al. 2025 for *Quercus* subsection *Campylolepides*). Furthermore, by exploring the (putative) mutation history of accordingly informative gene regions using population genetic approaches, one can get further insights in the evolutionary quality of the observed plastome divergences.

##### 2.3.2 Uniqueness of *F. hayatae* plastomes: long isolation and increased genetic drift

In contrast to the other East Asian species forming part of Lineage IV, the plastomes of *F. hayatae* show a consistent pattern of evolved sequence patterns. This genetic uniqueness will trigger a placement of *F. hayatae* as sister to all other or most members of Lineage IV during phylogenetic tree inference due to inevitable 'inclade-outclade' (long-)branch attraction. From the observed, tabulated mutation patterns, one can deduce that the placement as sister to the rest of Lineage IV is likely to be a tree-inference artefact. We didn't observe a single mutational pattern supporting an all-but-*hayatae* subtree, i.e. would support reciprocal holophyly. Instead, subtypes within Lineage IV (**Fig. SR2-10**) are more or less derived and can be easily traced to a shared ancestral sequence; which, in most plastid gene regions, is still found in accessions of *F. crenata* and *F. japonica*. The haplotypes of *F. hayatae* in all most signal-rich plastid regions can be derived from these ± ancestral haplotypes (**Table SR2-4**). Rather than representing a genuine sister lineage to all other Lineage IV plastomes, the plastomes of *F. hayatae* are strongly evolved from the common ancestor, the precursor plastome of the Lineage IV-LCM (last common mother of all beeches carrying Lin. IV plastomes).

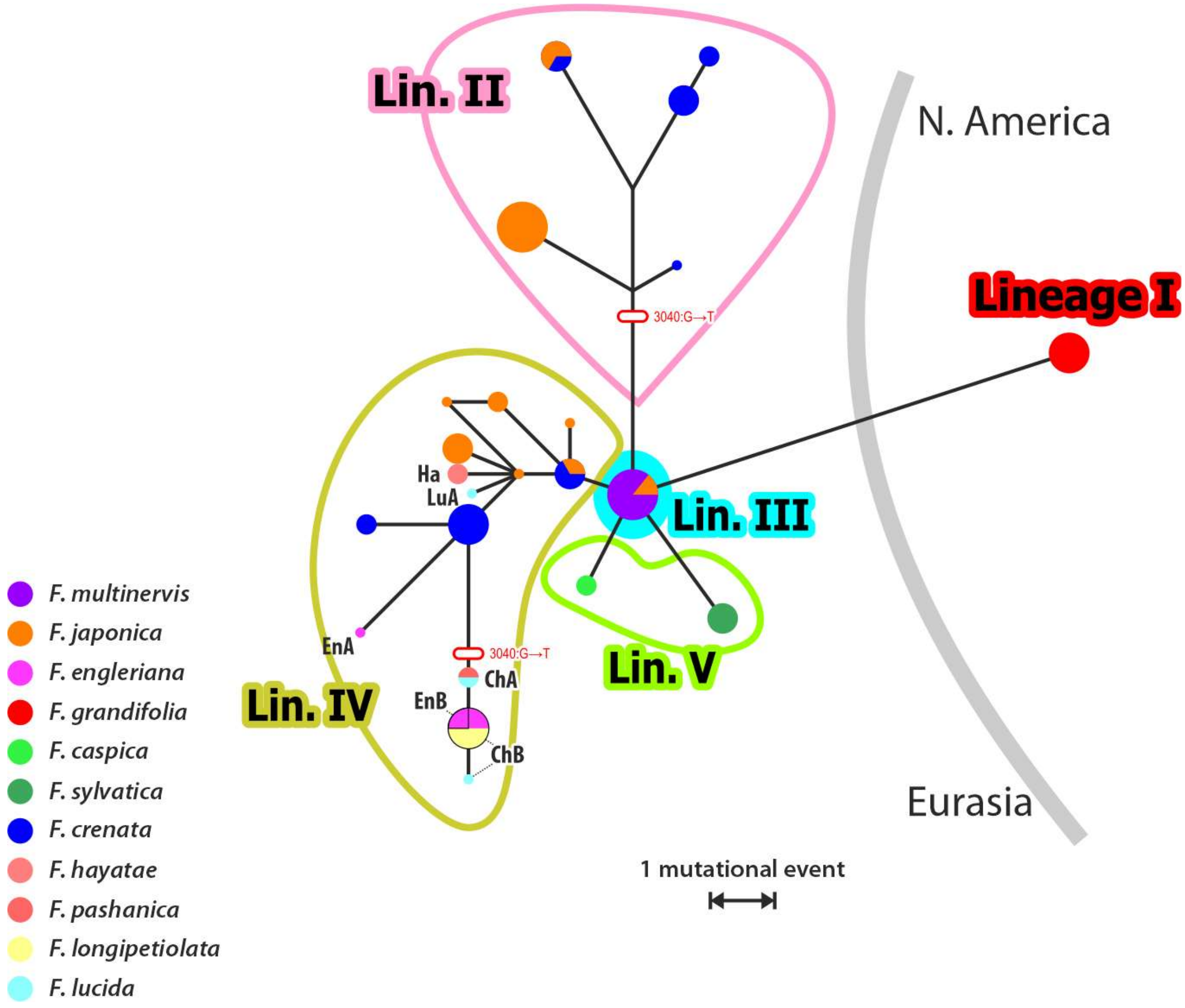


**Figure SR2-10 | (Quasi-)Full median network (cf. Bandelt et al. 2000) for *rpoC2.*** The single convergent mutation is indicated. Note the satellite haplotype position of the *F. hayatae* (IV-Ha)*, F. engleriana* (IV-EnA, -EnB, -ChB)*, F. longipetiolata, F. lucida* and *F. pashanica* (IV-ChA, -ChB) genomes within the Lineage IV and the placement of the Lineage III(-Mu) plastomes as the central median, i.e. showing a *rpoC2* sequence identical to that of the reconstructed ancestor of all Eurasian beech plastomes, the median connecting the other four main lineages.

Given that they probably always have had a small population size combined with a strong geographic isolation – *F. hayatae* only occurs in the north and central Taiwanese mountains, at altitudes between 1300–2000 m a.s.l. – high genetic drift can be assumed. The closest suitable habitats would be the mountain tops in south-eastern China (~400–500 km air-distance; 430 km air-distance to the putative south-easternmost populations of its continental cryptic sister species *F. pashanica*[15]).

[15] *Fagus hayatae* may be restricted to Taiwan or not. Focussing on *F. chenii,* the best-sampled study on Chinese *F. pashanica* (not recognised as distinct species and included in *F. hayatae*; Grimm 2025), Li et al. (2023), did not include material from the south-easternmost populations and relied on a subset of their earlier 28 nuclear gene set (Jiang et al. 2021) targeting only a single gene region (F128) that is species-diagnostic within East Asian species of *F.* subgenus *Fagus* (cf. Cardoni et al. 2022, data S5; **Fig. SR1-9**).

**Table SR2-4 | Mutations exclusive to *F. hayatae* plastomes (IV-Ha) and/or their sister plastome IV-LuA within Lineage IV in most signal-rich plastid gene regions.** SLPT = species-level plastid type; rLaD = *rbcL-accD* IGS; trnKQ = *trnK-trnQ* IGS.

| | **trnKQ** | | | | | ***rpoC2*** | **rLaD** | ***accD*** | ***ndhF*** | | | ***ycf1*** | | | | | | |
|---|---|---|---|---|---|---|---|---|---|---|---|---|---|---|---|---|---|---|
| **Alignment pos.** | **260ff** | **1132** | **1360…67** | **1649** | **2059ff** | **1917** | **177ff** | **695** | **145** | **244…76…82f** | **886** | **167** | **304** | **317** | **430** | **2472** | **3489** | **4831ff**[c] |
| Lin. I | $A_{14}$ | A | A…C | C | TT$A_{10}$CA | T | $A_9$… | G | T | G…A…GA | T | T | C | T | T | T | C | $[XY]_2$ |
| Lin. II, III, V | **$A_x$**GAAA[b] | A | A…C | C | TT$A_{8–9}$CA | T | $A_{9–\mathbf{10}}$… | G | T | G…A…GA | T | T | C | T | T | T | C | II/V:$[XY]_2$<br>III:$[XY]_{\mathbf{(3)5}}$ |
| Lin. IV[a] | $A_{9–11}$GAAA | A | A…C | C | TT$A_{(8)9–10(11)}$CA | T | $A_{7/9}$… | G | T | G…A…GA | T | T | C | T | T | T | C | $[XY]_{\mathbf{3}}$ |
| IV-LuA | **$A_{12}$**GAAA | A | A…**G** | **T** | TT$A_9$CA | T | **$A_{10}$**… | G | **C** | G…A…GA | T | **C** | C | **G** | T | T | **T** | XYX**X**XY |
| IV-Ha | **$A_{12}$**GAAA | **G** | **G**…**G** | C | **TT$A_{13}$** | **G** | **$A_{10}$**… | **A** | T | **T**…**T**…**TC** | **G** | **C** | **T** | T | **C** | **C** | **T** | XYX**X**XY |

[a] Semi-strict consensus, excluding singleton mutations and mutations restricted to a single SLPT other than IV-LuA and IV-Ha.
[b] $A_x$ = (10)12–13 A in Lin. II, 10 A in Lin. III and V-Sy plastomes; 9 A in V-Ca plastomes.
[c] X = AGA codon; Y = AGG codon

An equally isolated but less drifted sister plastome (SLPT IV-LuA) has recently be recorded from this area in an individual addressed as *F. lucida* (Yang et al. 2021). While being generally closer to the consensus haplotype of Lineage IV (**Table SR2-5**), this plastome shares a notable amount of exclusive mutations with the *F. hayatae* plastomes in all checked gene regions, separating these two SLPTs as an isolated sublineage from the remainder of the Lineage IV plastomes (**Tables SR2-4**, **SR2-5**; **ODA**, file *00_Bits.xlsx*).

**Table SR2-5 | Mutations exclusive to *F. hayatae* plastomes (IV-Ha) and/or their sister plastome IV-LuA within Lineage IV in plastid gene regions commonly used as 'barcodes' and species-level haplotype studies.** SLPT = species-level plastid type; tHpA = *trnH-psbA* IGS; trnKi = *trnK* (3') intron; trnLi = *trnL*(XXX) intron; trnTL = *trnL-trnF* IGS.

| | **tHpA** | | ***matK*** | | **trnKi** | | | **trnTL** | **trnLi** |
|---|---|---|---|---|---|---|---|---|---|
| | 75ff | 161ff | 1217…1265 | 1700 | 2006 | 2085ff | 2375 | 20 | 1409 |
| Lin. I | $A_8$ | $A_7$ | A…C | C | T | $A_{10}$TC | A | C | T |
| Lin. II, III, and V | $A_9$/**$A_{12}$**/$A_{9–10}$ | $A_8$ | A…C | C | T | $A_{9–11}$CC | A | C | T |
| Lin. IV consensus[a] | $A_{9–10}$ | $A_8$ | A…C | C | T | $A_{9–11}$CC | A | C | T |
| IV-LuA | **$A_{12}$** | **$A_{11}$** | A…C | **T** | **C** | **$A_7$** | **C** | C | T |
| IV-Ha | $A_9$ | **$A_{11}$** | **C…A** | **T** | **C** | $A_{10}$CC | **C** | **T** | **C** |

[a] Semi-strict consensus, excluding singleton mutations and mutations restricted to a single SLPT other than IV-LuA and IV-Ha.

In some gene regions, e.g. the *ndhF* and *matK* genes, the IV-LuA plastome shows a sequence representing the modal consensus of the Lineage IV, the putative ancestral haplotype; haplotypes highly similar to (near-)identical to early diverging and moderately evolved *F. japonica* (and *F. crenata*) Lineage IV plastomes (IV-JaC, -JaD, -JaG, -CrC); much in contrast to other Lineage IV plastomes from Chinese species including the other two sequences from *F. lucida* individuals (all part of the sequentially strongly evolved Lin. IV core clade) or its sister SLPT IV-Ha (*F. hayatae*).

Irrespective of their exact position within Lineage IV and in relation to the early diverging Lineage IV plastomes found in western-most and southern Japanese islands (IV-JaC and -JaD), it is clear that these two sister SPLTs (IV-LuA + IV-Ha) were isolated from the plastome pool carried by the rest of the Chinese beeches (Lin. IV core clade) before the latter evolved and radiated. One branch of this lineage seems to have survived in south-central China, where it was consumed by the modern species of *Fagus* subgenus *Fagus* when they migrated in that region. Its most remote populations were trapped and subsequently isolated in Taiwan, where they provided the maternal ancestor of the extant *F. hayatae*, the *Hayatae*-LCM.

##### 2.3.3 Further general trends within Lineage IV

As exemplarily seen in **Figure SR2-10**, the ancestral ('median' in the context of median networks; Bandelt et al. 1995) haplotypes are concentrated in *F. crenata–japonica* exclusive sublineages of Lineage IV (**Table SR2-4**). This further supports the hypothesis (→ **Section 2.2.2**) that the Japanese archipelago has conserved a lot of the original variation of Lineage IV plastomes and must be closer to the area of origin of the Lineage IV 'Eve', while the plastomes of the Chinese species, irrespective of their subgenus, represent a single sublineage that evolved ("budded") from this North-East Asian stock and migrated to lower latitudes in East Asia.

The continental Asian haplotypes (*F. engleriana, F. longipetiolata, F. lucida, F. pashanica*) show consistently evolved satellite types. Together with certain *F. crenata* plastomes, they form the crown-group of Lineage IV ('Lin. IV core clade'). The lack of branch and character support in this part of the tree can be attributed to a relatively recent fast radiation event, in which several sublineages established simultaneously and accumulated few lineage-segregating mutations. While relatively few, often a single or few SNPs or other mutations per gene region (coding or non-coding), these lineage-segregating mutations are highly conserved across the entire plastome and convergences or parallelisms are rare. This is illustrated in **Figure SR2-11**, which shows a map of bipartition-segregating mutations in the high-conserved *trnL* intron + *trnL-trnF* IGS (traditionally used as a "barcode" for plants at various hierarchical levels) and the signal-rich *ycf1* gene on the Lineage IV subtree. As a general rule, convergent mutational patterns are often associated with long terminal branches (long isolation time, increased genetic drift due to small active population sizes), while being extremely rare on internal branches. Exceptions are, as detailed above, inversions and mononucleotide repeat related length-polymorphism.

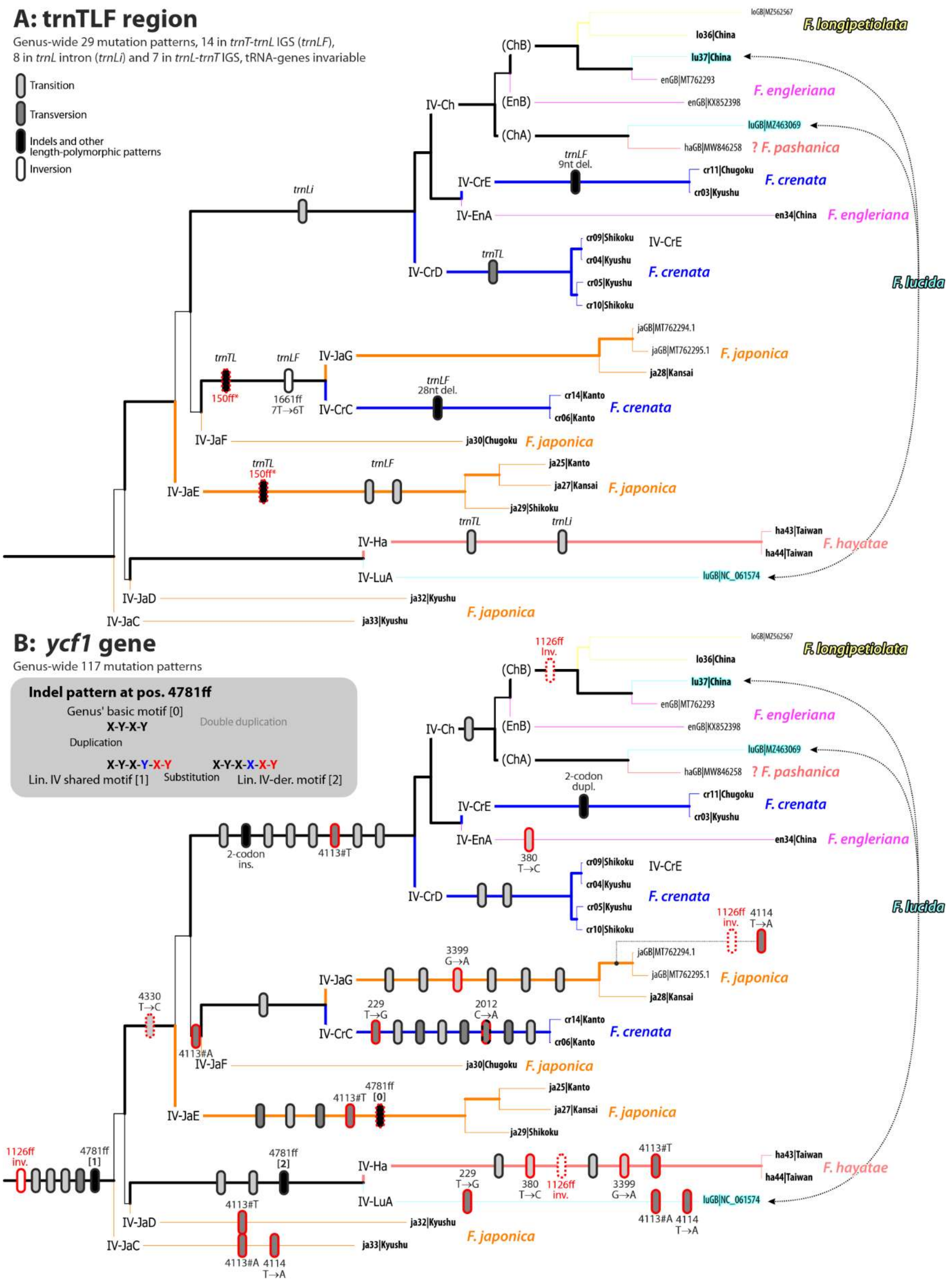


**Figure SR2-11 | Mutations mapped on the Lineage IV subtree in Fig. SR2-2. A.** The high-conserved *trnL* gene and flanking spacers, a commonly used plastid "barcode". **B.** Most signal-rich partition, *ycf1* gene. Signatures Light grey , transitions, medium grey, transversion, black, other mutational patterns. Black outlines, clade-specific mutations; red outlines, mutations accumulated in parallel/convergently (number gives position in *ycf1* gene); stippled outlines, detected backmutations (reversals). * Lin. III and Lin. V plastomes share with most of the Lin. IV plastomes a 43 nt-long duplication in at the 5' end of the *trnL-trnF* IGS, which is missing in JaE and CrC+JaG plastomes.

##### 2.3.4 Deep east-west division and white spots in the West-Eurasian Lineage V

The plastid differentiation structure of Western Eurasian beeches, traditionally perceived as one or two species ('European beech' *F. sylvatica* s.str., 'Oriental beech' *F. orientalis* s.l. ranging from SE. Europe to N. Iran) has been intensively studied using cpDNA microsatellite data (Demesure et al. 1996; Magri et al. 2006; Hatziskakis et al. 2009; Papageorgiou et al. 2014; Tsiripidis et al. 2024) with a particular focus on identifying Pleistocene refugia for *F. sylvatica*. The now available complete plastome data (Ulaszewski et al. 2021; this study) renders some of their conclusions obsolete or even problematic: the complete plastomes of Spanish, French, trans- and cis-Alpine, Carpathian and north-western Greek *F. sylvatica* obtained by three independent working groups are near-identical. An exception is the plastome of our Italian relict individual from the central Apennines. The lack of a clear geographic or phylogenetic sorting of mutation patterns (see also **SupplGenetics_cpDNA.xlsx**, sheet *LinV HTs*) makes microsatellite differentiation patterns – "allelic" variation restricted mostly to singletons, cf. column "Marker ratio" in Ulaszewski et al. 2021, table 4 – difficult to interpret above the level of the here defined SLPTs, and results in the accordingly poorly resolved V-Sy subtrees (**Fig. SR2-2**; Ulaszewski et al. 2021, fig. 2, showing only two branches with BS support > 24).[16]

**Figure SR2-12** gives the geographic context of a haplotype network inferred from two common plant "barcodes" (*trnK/matK* region and trnTLF region) of the individuals for which complete plastomes have been generated. The data shows no geographic structure within *F. sylvatica* and their SLPT V-Sy. Moreover, mutations occurring in some of the *F. sylvatica* plastomes are randomly distributed: for instance, a 4 nt-long deletion in the *ndhF* gene, a mutation exclusive to most accessions generated by Ulaszewski et al. (2021), can be found from the Cantabrian Mountains via Central Europe to the Bulgarian Carpathians and include an individual from the Italian Alps but is missing in all our plastomes and in four plastomes sampled close-by. A possible split into two V-Sy *ndhF* haplotypes (deleted and undeleted) is, however, correlated with a 4 nt-long duplication in the *ycf1* gene (**SupplGenetics_cpDNA.xlsx**, sheet *LinV HTs*; **Fig. SR2-12**), which may be indicative for a general plastid dimorphism of *F. sylvatica* populations outside its most southern areas but would need to be verified by target sequencing of the according fragments using a much larger sample of populations and individuals. In

[16] The lack of tangible results may be the reason, why the study of Ulaszewki et al. (2021), despite its impressing sampling and data set, was published in a MDPI journal, *Genes*. MDPI is a massive-dump, profit-optimised open-access publisher with a dubious reputation (e.g. *Science Insider* 2018; Web of Science delisting 50 journals; English Wikipedia: Evaluation and Controversies), employing publication strategies that can be described as predatory (see e.g. Oviedo-Garcia 2021, *Res. Eval.* **30**: 405–419a). Concerns about their publishing process has led to black-listing of some of their journals in e.g. Norway and South Africa by the local science councils.

conclusion, while there is a profound and consistent (**Fig. SR2-12**) divergence between the western- (*F. sylvatica*) and easternmost species (*F. caspica*) of the West-Eurasian beech lineage, neither one shows any further substructure, despite their much different extant ranges; and the deep split is not apparent from traditionally applied plastid "barcodes" or microsatellite markers.

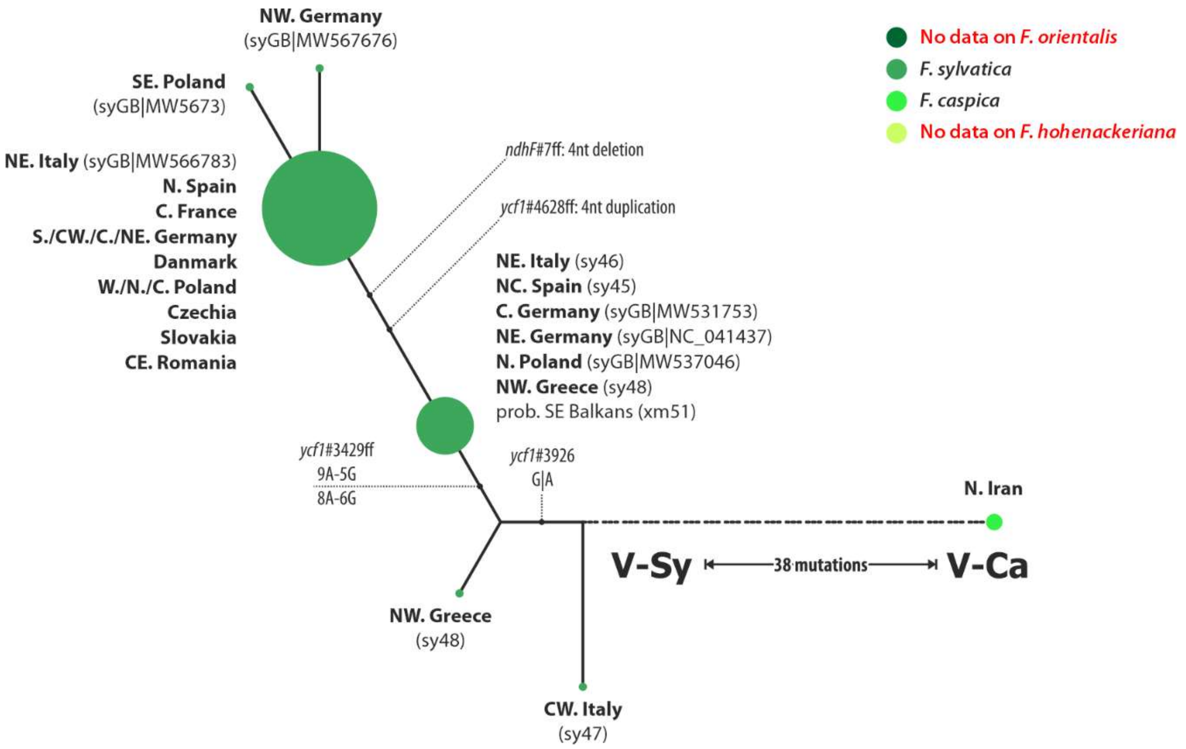


**Figure SR2-12 | Full Median network based on mutational patterns in the five most signal-rich plastid regions plus two classic "barcodes" (*trnK/matK* cistron; trnTLF region).** The barcode regions show no variation within *F. sylvatica* plastomes (in contrast to the sequences reported by Paffetti et al. 2007, cf. **Fig. SR2-13** below).

Thus, the plastomes carried by the species bridging between *F. sylvatica* and *F. caspica*, the western Euxinian *F. orientalis* and the Pontic-Caucasian *F. hohenackeriana*, remain a wildcard. They are only partly covered by existing chloroplast based studies (i.e. Demesure et al. 1996) and the problematic Sanger sequence data compiled by Paffetti et al. (2007; **Fig. SR2-13; Table SR2-6**).

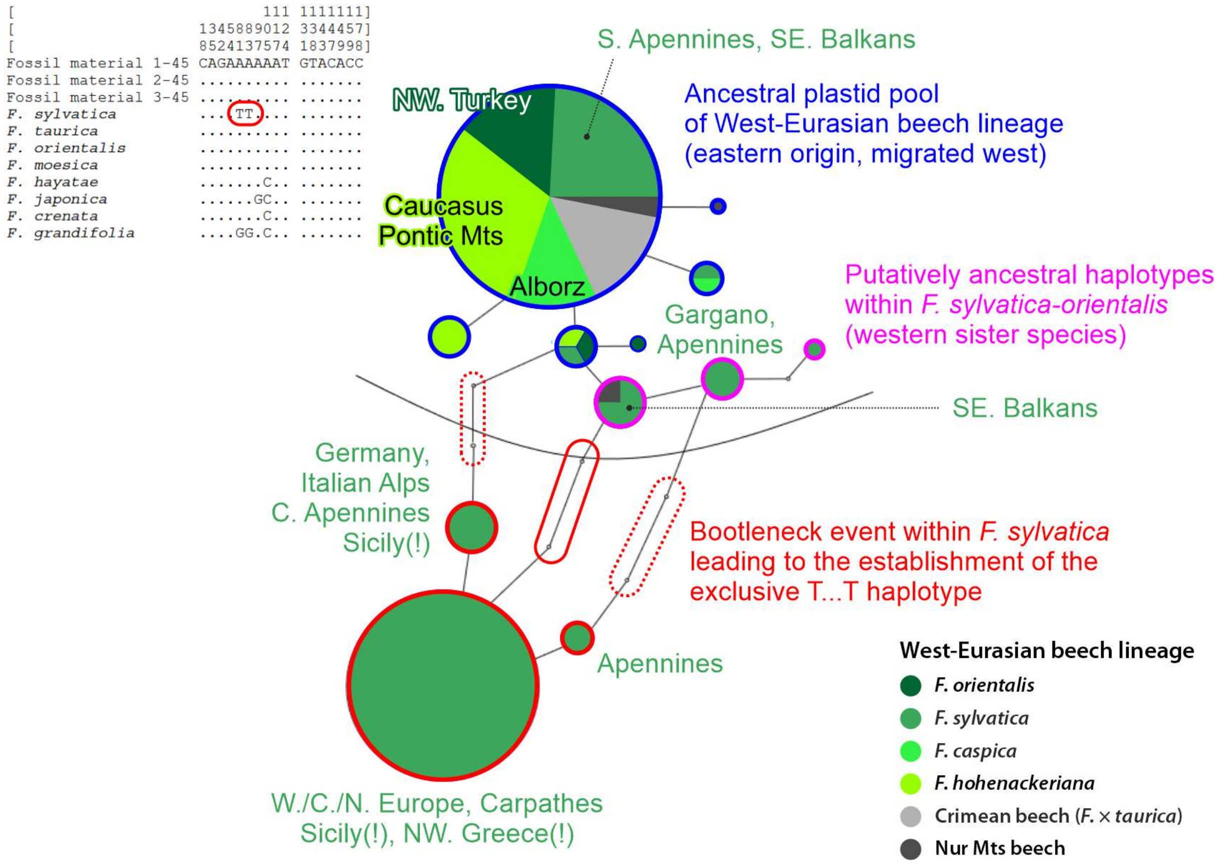


**Figure SR2-13 | The haplotype network shown in Paffetti et al. (2007), with the provenance of individuals annotated and coloured according to the new species concept for Western Eurasian beeches.** Species concept after Denk et al. (2024). Stippled outline, unlikely mutational pathways; if a genuine pattern, the T…T mutation would have been established a single time, in a single mother population of *F. sylvatica* and was propagated across the entire range of the species, while *F. sylvatica* populations in putative Pleistocene relict areas and close to the area of their sister species *F. orientalis* (C./S. Italy, SE. Balkans) (partly) conserved the ancestral haplotypes. Paffetti et al. (2007) erroneously concluded that all *F. sylvatica* retaining the ancestral West-Eurasian haplotype are *F. orientalis* with a *sylvatica*-phenotype. However, there are no naturally occurring *F. orientalis* present in any of these areas and all nuclear data from these provenances confirm their phenotypic assignment as pure *F. sylvatica* populations.

**Pitfalls and prospects of broadly sampled cpDNA "barcodes"**—Paffetti et al. (2007) argued that it is possible to distinguish between *F. sylvatica*, the European beech, and *F. orientalis* s.l., the Oriental beeches, based on two point mutations in the *trnL* intron (reproduced in **Fig. SR2-13**), and, using this argument, inferred the occurrence of late Pleistocene *F. orientalis* in the Venetian laguna area based on degraded sequences they attributed to subrecent material. Their data basis included *F. sylvatica* individuals from across the entire range of the species and a comprehensive sample covering the major provenances of the Oriental beeches, with individuals that represent probably all three species forming this complex (according Denk et al. 2024): *F. orientalis* (individuals from NW. Turkey), *F. hohenackeriana* (individuals from Georgia and Armenia) and *F. caspica* (individuals from N. Iran, covering populations not far from the populations we sampled for V-Ca plastomes). The main conclusion of Paffetti et al. (2006) can already be rejected by their own analysis as the supposedly *orientalis*-specific haplotypes are shared by half of their Italian and Greek *F. sylvatica* individuals (**Fig. SR2-13**; see also the haplotype map in Magri et al. 2006, fig. 5), which they explain by their ancientness and/or disparate pheno- and genotypes: the *F. sylvatica* in "cluster A" are only phenotypically *F. sylvatica,* but genetically *F. orientalis*, a hypothesis that can be rejected based on nuclear and morphological data (Denk et al. 2024 and literature cited therein). A very simple, alternative explanation for the structure of the haplotype network would have been that the "*F. orientalis*-complex"[17] A…A haplotype ("cluster A") represented a genetic symplesiomorphy of West-Eurasian beeches, non-evolved haplotypes that persisted in the Oriental beeches as well as in the relict areas of *F. sylvatica* (cf. Magri et al. 2006, Magri 2008; note the microsatellite diversity in the regions with A…A haplotypes in *F. sylvatica*), while the T…T haplotype ("cluster B") would be a recent mutation that started to manifest itself in *F. sylvatica* (equivalent to the red haplotype 1 in Magri et al. 2006, fig. 5). Generally, the *F. sylvatica* populations are more homogenous and less drifted from their eastern relatives in their putative south-eastern glacial refugia than elsewhere.

However, *any* conclusion based on only these data would have been problematic. While few point mutations may be highly informative, they need to be put in the context of the complete plastome phylogenies to discern between high- (sorted along the complete plastome tree) and low-informative (convergences, backmutations) SNPs (cf. **Figs SR2-6, SR2-8, SR2-12**) as well as potentially ancestral vs. evolved mutation patterns. A comparison with *trnL* intron (and *trnT-trnK* and *trnL-trnF* IGS) sequences from other beech species (complete plastomes and classic Sanger sequences) revealed that the allegedly *F. sylvatica-orientalis* s.l. segregating SNPs are not present in data generated by any other study so far (**SupplGenetics_cpDNA.xlsx**, sheet *GB-HTs_trnTLF*). If they exist at all in the *trnL* intron, they would just represent satellite haplotypes of the widespread haplotype(s) (cf. **Fig. SR2-12**) of *F. sylvatica* V-Sy plastomes. Which, in the *trnL* intron and *trnL-trnF* IGS, are difficult to distinguish from most other Eurasian plastomes, specifically Lineage III and least-drifted Lineage IV plastomes (**Table SR2-6**).

[17] Paffetti et al.'s "*F. orientalis* complex" includes all Orient beeches as well as potential hybrid taxa with *F. sylvatica*: the S.E. European *F.* (×) *moesiaca* and the Crimean *F.* (×) *taurica.* For *F.* × *moesiaca* all assembled molecular data supports its inclusion as a low-land ecotype in *F. sylvatica*. The status of *F. taurica* as a potential hybrid between *F. sylvatica* and *F. orientalis* or *F. hohenackeriana* remains uncertain (Denk et al. 2024).

**Table SR2-6 | Mutation patterns in the *trnL* intron based our complete plastome data set and data harvested from gene banks.** Mutations restricted to a plastid lineage or West Eurasian species in bold, the haplotype/species-defining transitions (Pafetti et al. 2007, fig. 2) are underlined; mutations reported by Pafetti et al. that could not be confirmed by other studies in red colour.[18] Singleton mutations and likely artefacts not listed; full list provided in **SupplGenetics_cpDNA.xlsx**, sheet *GB-HTs_trnTLF*.

| Species, lineage | Mutational pattern | | | | | | | | |
|---|---|---|---|---|---|---|---|---|---|
| *F. grandifolia*/Lineage I plastomes | G… | $A_{11/13}$-GGATA[a] | A | A | C | C | T | A | A/**G** |
| Japanese spp. w/Lin. II plastomes | G… | $A_{9/13}$-GGATA | **G** | A | C | C | T | A | A |
| *F. multinervis*/Lin. III plastomes | G… | **$A_8$**-GGATA | A | A | C | C | T | A | A |
| East Asian spp. w/Lin. IV plastomes | G… | $A_{9–13}$-GGATA | A | A/**G** | C | C/**T** | T/**C** | A | A |
| V-Sy plastomes | G… | $A_{13–14}$-GGATA | A | A | **A** | C | T | A | A |
| *F. sylvatica* Italy, SE. Europe | G… | $A_{10–15}$-GGATA<br>$A_{11}$**G**-GGATA<br>$A_{12}$**G**-GG**T**T**T** | A | A | **A** | C | T | A/**C** | A |
| Other *F. sylvatica* | G… | $A_{11–13}$**G**-GG**T**T**T** | A | A | **A** | C | T | A | A |
| *F. orientalis* | G… | $A_{10–11}$-GGATA<br>$A_{11}$**G**-GGATA | A | A | **A** | C | T | A | A |
| *F. hohenackeriana* | G…/**C**… | $A_{10}$-GGATA | A | A | **A** | C | T | A | A |
| *F. caspica* | G… | $A_{9–10}$-GGATA | A | A | **A** | C | T | A/**C** | A |
| Nur Mts beech | G… | $A_{11}$**G**-GGATA | A | A | **A** | C | T | A | A |
| V-Ca plastomes | G… | $A_{10}$-GGATA | A | A | **A** | C | T | A | A |

[a] According to the data uploaded by Pafetti et al. (2007) *F. grandifolia* is characterised by a distinct G-enriched motif: $A_{12}$**G**-GG**G**T**G**, but all data from other studies and our complete chloroplast genomes failed to retrieve this pattern. Unless confirmed by new data, it has to be treated as a sequencing/editing artefact.

Most importantly for West-Eurasian beeches, the set-up used by Paffetti et al. (2007) cannot discriminate between the most-divergent *F. sylvatica* V-Sy plastomes and *F. caspica* V-Ca plastomes. Thus, in stark contrast to the data assembled for *F. crenata* (Fujii et al. 2002, Tsumura & Suyama 2015; **Section 2.2.2**), we cannot assign *ad hoc* the found *trnL* intron/ *trnL-trnF* IGS haplotypes to the here defined SLPTs, and potential additional SLPTs or geographically restricted haplotypes cannot be detected using these data.[19] Only by using a framework based on the complete trnTLF region sequences extracted from the complete plastomes, the Sanger sequence data could be put into some taxonomic context **(Fig. SR2-14)**,

[18] These mutations are missing from data sets generated by complete chloroplast genome assembling as well as older and younger Sanger sequences, and including material from similar provenances. There are several possible explanations for such study-restricted mutations: *(i)* Paffetti et al. (2007) like most authors of the time may have used a non-proofreading polymerase, which can account for artificial mutations (in course of PCR) restricted to a single or few accessions, but not a major pattern such as the T…T and G…G haplotypes. (*ii*) their sequence reads may have been of mediocre quality, and they misedited the raw chromatograms (the authors didn't report which sequencer was used); sequences differing by the number of nucleotides in mononucleotide repeats or showing towards one nucleotide (in this case G) are a common bias of the time. (*iii*) they manipulated (part of) the data in order to enforce some originally random segregating pattern.

[19] The sequences obtained from the subrecent material by Paffetti et al. (2007) are equally uninformative; however, they show an increased amount of pseudogenic mutations as expected for degraded DNA. Pafetti et al. assigned them to their HT 6, which is exclusive to *F. sylvatica*.

making use of the circumstance that the few SNPs in this region are generally sorted by phylogenetic lineages, while the mononucleotide repeat patterns are consistent only at the level of SLPTs (cf. **Fig. SR2-11**). The A…A pattern relied on by Paffetti et al. (2007) has no discriminative power being shared across the entire genus' plastomes, the derived T…T pattern likely represents a data artefact. All West-Eurasian species share the same or near-identical trnTLF haplotypes that only differ by a single but highly conserved point mutations from the other two main continental Eurasian plastid lineages (Lin. III and IV) located in the centre of the *trnL* intron. The diagnostic SNP to distinguish the two divergent SLPT within the West-Eurasian Lineage V, V-Ca and V-Sy, is located in a mononucleotide repeat in the *trnT*-*trnL* IGS, for which only limited Sanger data are available.

The ancestral trnTLF haplotype shared across West-Eurasian species and a derived *F. sylvatica*-only trnTLF haplotype appear to differ solely by the number of As in the mononucleotide repeat in the 5' part of the *trnL* intron, whether this difference correlates with the V-Ca and V-Sy SLPTs cannot be judged at this point. Also in this case, data from two length-polymorphic A-repeats in the *trnT*-*trnL* IGS may be crucial to identify haplotypes within the SLPT V-Sy (or additional SLPTs between V-Sy and V-Ca). The sequences obtained from the subrecent material represent degraded versions of a putatively *F. sylvatica*-restricted haplotype. Moreover, it is clear that the entire trnTLF region only accumulated very few lineage-consistent mutational patterns across the continental Eurasian clade that comprises Lineage III, IV and V plastomes: any broadly sampled haplotype study using this regions can only be interpreted on the background of the complete plastome phylogeny (**Fig. SR2-14**).

Regarding inter-species diversification in the West-Eurasian lineage as well as the potential intra-species biogeographic patterns encoded in their plastomes, the necessary next step is hence to sample complete plastomes from across the range of the other two species of West-Eurasian beeches: *F. orientalis* (westernmost populations in SE. Bulgaria and NE. Greece; easternmost populations in N. Turkey) and *F. hohenackeriana* (N. to NE. Turkey, Georgia, Armenia, southern Russia) and the western-most populations of the newly recognised species, *F. caspica*: individuals originating from the Talysh region in south-eastern Azerbaijan. Such a reference data set could be complemented by individuals from the isolated populations on the Crimean Peninsula (the putative hybrid *F.* × *taurica*) and the Nur Mountains in Hatay province, southernmost Turkey (uncertain status; cf. Denk et al. 2024). This will allow to identify the best-possible sequence markers and most-informative mutations for broadly sampled population-level studies as it is already the case for the Japanese species (**Figs SR2-6** and **SR2-8** above).

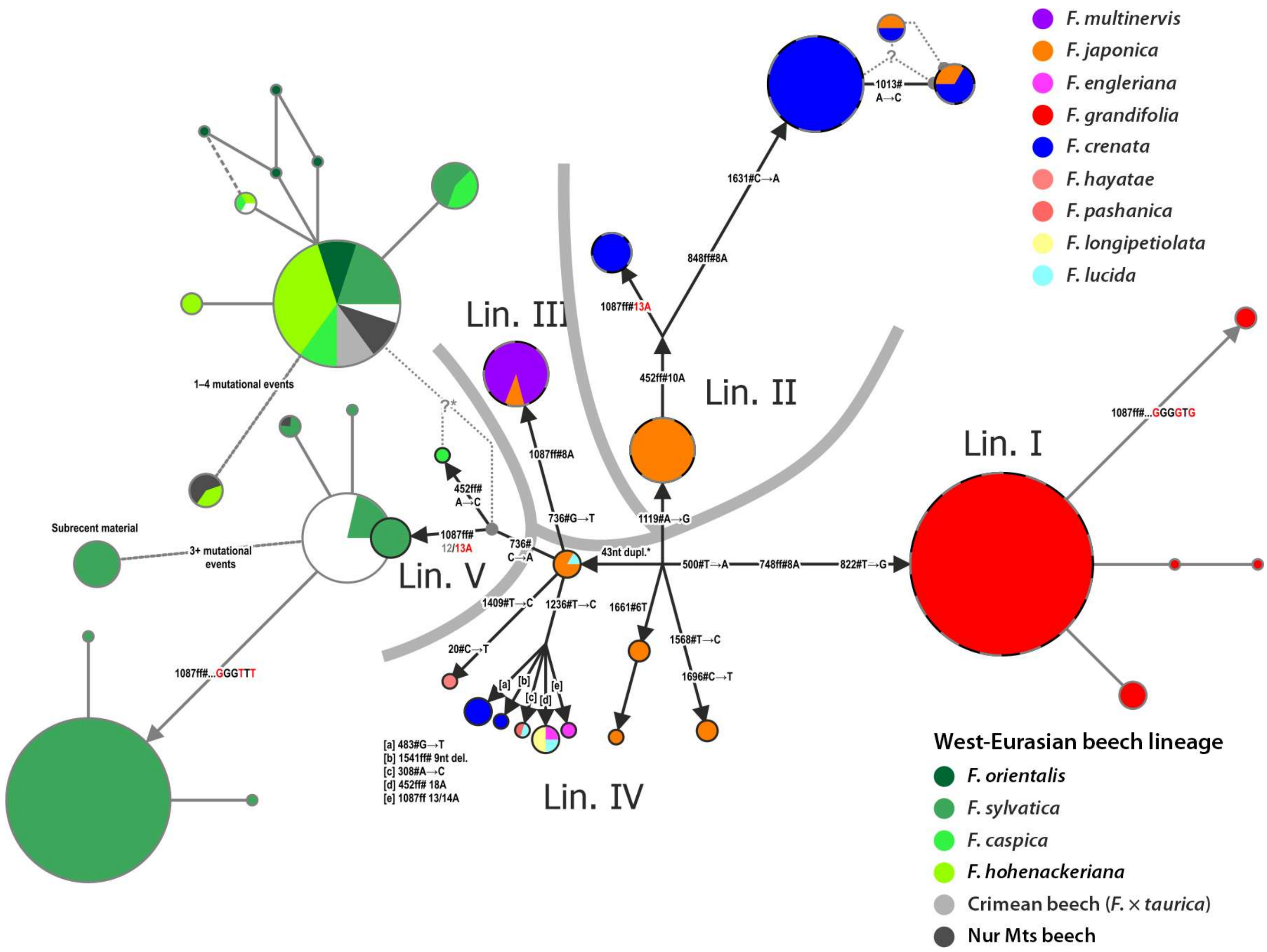


**Figure SR2-14 | Reduced Median network for the entire trnKLF region using data from complete plastomes and Sanger sequences.** The core graph represents a quasi-full median network inferred from the complete trnTLF region (→ **ODA**, file *00_Bits.xlsx*, sheet *trnTLF*), which includes three plastid regions traditionally advocated as angiosperm "barcodes": the *trnT-trnL* IGS, the *trnL* intron and the *trnL-trnF* IGS. Black outlines, sequences extracted from complete plastomes; grey outlines, Sanger sequences covering (part of) the trnTLF region (Lineage IV Sanger sequences not included for simplicity).

Equally important, complete and geo-referenced plastomes are crucial to select the micro-satellite markers with the capacity to distinguish between the species-level plastid types and their main haplotypes (**Fig. SR2-9**). While providing more differential signal, the microsatellite differentiation patterns will inevitably include a higher level of stochasticity, within and outside the West-Eurasian Lineage V. Only by mapping the used microsatellite markers on the complete plastome phylogenies including all main types, it is possible to assess the information content provided by their differentiation signal for a certain taxonomic and biogeographic level. To put the microsatellite differentiation patterns found in *F. sylvatica* in an explicit evolutionary and Pleistocene refugia context, one needs to add comparative data across the entire range of the Oriental beeches, most importantly, the populations that form *F. orientalis*, the sister species of *F. sylvatica.* The critical question in such a context is whether relict haplotypes are restricted to *F. sylvatica* and not shared with its sister species or even their eastern relatives.

### 3 Molecular dating

#### 3.1 Initial node dating of plastid vs nuclear data from the same individuals

As baseline for the molecular dating experiments and results, we conducted prior-poor dating using our new data. The tip set only included individuals for which we generated both the complete chloroplast genomes (plastomes) and nuclear data (LFYi2 + *CRC*) allowing for a 1:1 comparison of the divergence signal in the two principal data sets. Even when using the same root age prior, i.e. under the assumption that both the nucleome and plastome of modern-day individuals and species started to diverge at the same, the temporal decoupling of speciation events (in addition to their evolutionary decoupling) is evident (**Fig. SR3-1**).

Crown-group radiation – divergence and radiation of modern-day species manifesting as dist-inct nuclear genotypes – happened after the splitting of the plastome pool into main SLPTs (species-level plastid types; **Section 2.2**). For instance, a late Oligocene stem age and mid-Miocene crown age of *F. crenata* (see also Renner et al. 2016), four individuals from across the Japanese archipelago, stands against a late Eocene divergence of the plastomes (Lin. II vs Lin. IV; two SLPTs each) carried by these four individuals. The same holds for the putative sister species *F. hayatae* and *F. longipetiolata* (one individual each, two most distinct Lin. IV SLPTs) and divergences within *Fagus* subgenus *Englerianae* (Lin. II, III, and IV plastomes, various SLPTs ↔ recently radiated and incompletely sorted modern-day species). Of the main plastid lineages, only Lineage V, the plastome of the West-Eurasian beech lineage, showed a good correlation, with a slightly older stem (split from sister lineages) and crown age (divergence between the genomes of an Italian *F. sylvatica* and Iranian *F. caspica* individual reflecting the primary split within this lineage; Cardoni et al. 2022, Denk et al. 2024) in the nuclear than the plastid chronogram. Because we used much more realistic root age constraints (see Grímsson et al. 2016, Renner et al. 2016; **SupplM&M.pdf**, *section 5.1*) already the estimates of this simple dating experiment do not contrast the fossil record and are in general (much) older than

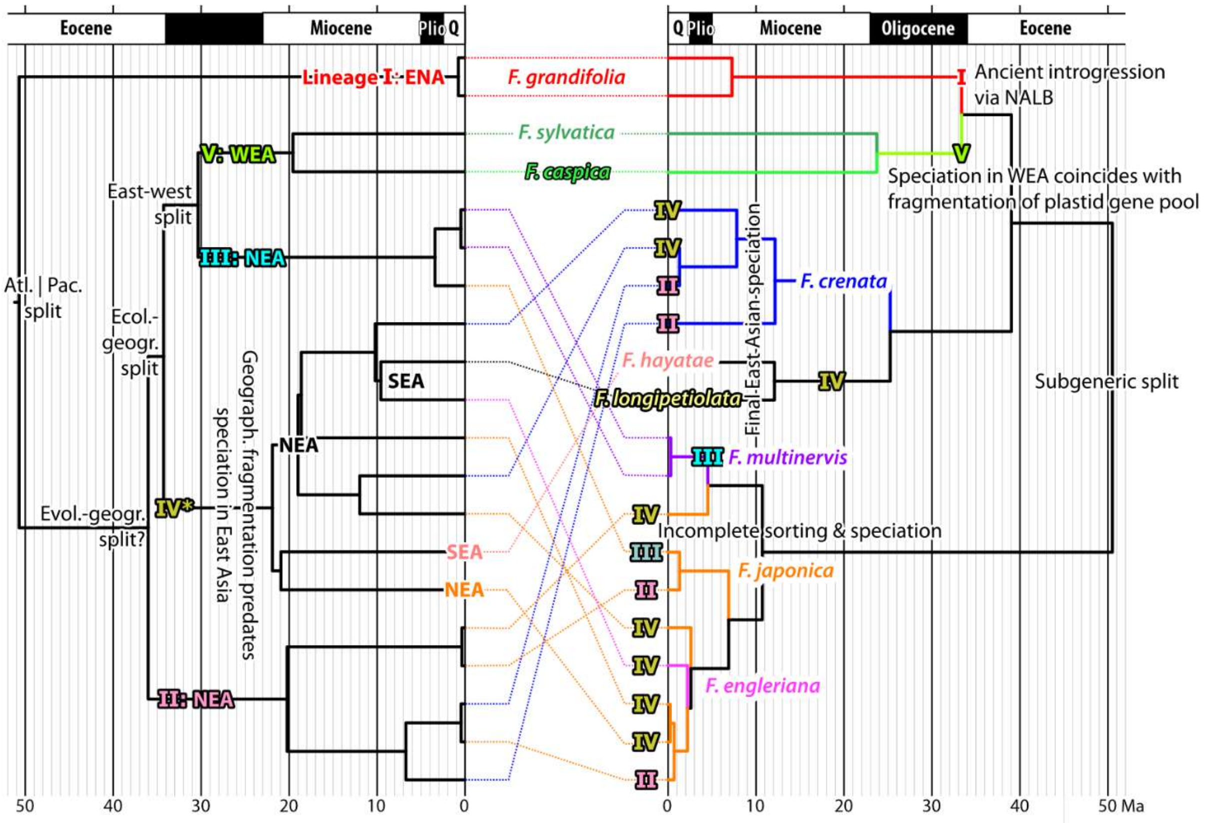


**Figure SR3-1 | Tangled chronograms using the same root age constraint** of 52±2 Ma for comparison with earlier dating studies (*F. langevinii* representing starting crown-group radiation in modern beeches). Plastid crown-group radiation, reflecting potential past speciation processes, generally preceding formation of modern-day species lineages. Primary divergences incongruent but ± coeval. Roman numerals: main plastome lineages. Biogeographic regions (spatiotemporal units; cf. **SupplM&M.pdf**, *section 5*) abbreviated by three letters: ENA = eastern N. America; NEA = northern E. Asia; SEA = southern E. Asia; WEA = Western Eurasia. Other abbrev.: Atl.|Pac. = split between northern N. Atlantic (NALB = North Atlantic land brigde) and northern N. Pacific populations (cross-Beringia; → **Section 4**); the latter further radiating in(to) NEA and ultimately colonising WEA and SEA.

those reported by Jiang et al. (2022, fig. 2), who, on the basis of outdated secondary literature used in other dating papers, assumed a >20 Ma too young age stem and crown age of genus *Fagus* and the crown age of their outgroup clade comprising several core Fagaceae.

Based on the fossil record, the palaeogeographic history (summarised in **Section 4**; **Suppl-FossilTable.xlsx**, sheet *main list*; data set emended from Denk & Grimm 2009) and modern-day differentiation patterns (→ **Sections 1, 2**), the assumption that plastomes and nucleomes of modern-day species started to diverge at the same time is likely wrong (note that Fagales studies using dated trees were all inferred from combined but incongruent nuclear and plastid data set; Sauquet et al. 2012, Xing et al. 2014, Xiang et al. 2014, Larson-Johnson 2016[20]). The modern-

[20] While all these papers have principal issues regarding their approach and conclusions, Larson-Johnson (2016) sticks out by being fundamentally flawed in *any* respect, be it the used data, methodology, results or interpretation.

day species can be grouped into two mutually holophyletic subgenera characterised by non-overlapping suites of morphological and genetic (sequence-wise) traits (Shen 1992, Denk & Grimm 2009; Denk et al. 2024). However, these subgeneric differences only manifest in fossil phenotypes at the Eocene-Oligocene boundary, at c. 34 Ma (**SupplFossilTable.xlsx**), i.e. nearly 30 myrs after genus *Fagus* reached a cross-Arctic distribution (→ **Section 4**, see also main-text fig. 8). It is thus conceivable that the primary split in the plastomes between (eastern) North American and Eurasian beeches established when *Fagus* started to radiate in the late Paleocene (→ **Section 4: Fig. SR4-2**), while the differentiation and primary sorting of nucleomes reflecting the subgeneric split coincided with the phenotypical manifestation of the modern subgenera at or shortly before the Eocene-Oligocene boundary (→ **Section 4: Fig. SR4-5**; see also **Section 3.3**).

Under this assumption, the Eurasian plastomes started to radiate ~15 myrs before the establishment of the modern subgenera (**Fig. SR3-2**). When the subgenera manifested at the Eocene-Oligocene boundary, all main five plastome lineages were already present or about to diverge (last: relict Lin. III from W. Eurasian Lin. V). The Lineage I plastome was picked up by the ancestors of *F. grandifolia* after *Fagus* subgenus *Fagus* radiated during the Oligocene. The earliest diverging Lineage II is a probable candidate for the plastome of the first common ancestor (FCA) of *F.* subgenus *Englerianae*. The alternative would be the Lineage III plastome, which would require, however, that both the Eurasian branch of *F.* subgenus *Fagus* as well as early members of *F.* subgenus *Englerianae* picked up plastid signatures from stem group beeches and extinct lineages (Lin. II, Lin. IV, Lin. V plastomes) during their initial radiations.

The divergences within the Eurasian plastomes and subsequent radiations of Lineages II and IV seem to reflect mid-Oligocene to Early Miocene allopatric speciation events predating the formation of the modern-day East Asian species lineages by at least ~5 myrs. Irrespective of which plastome lineage(s) is (are) the original plastomes of the first (FCAs) or last common ancestors (LCAs) of either subgenus, secondary contact involving "chloroplast capture" appeared to have been common in the Miocene or post-Miocene of East Asia (**Fig. SR3-2**): (*i*) the unique Lineage IV plastome of *F. hayatae* (IV-Ha) is substantially older than the species as a genetically isolated entity; (*ii*) the modern-day species of *Fagus* subgenus *Englerianae* only diverged in the Late Miocene to Pliocene but their plastomes had been by the Early Miocene at last; and (*iii*) *F. crenata* carries plastomes that diverged from the Oligocene till sub-recent times (IV-JaB plastome in *F. crenata* of C. Honshu; cf. **Fig. SR2-6**).

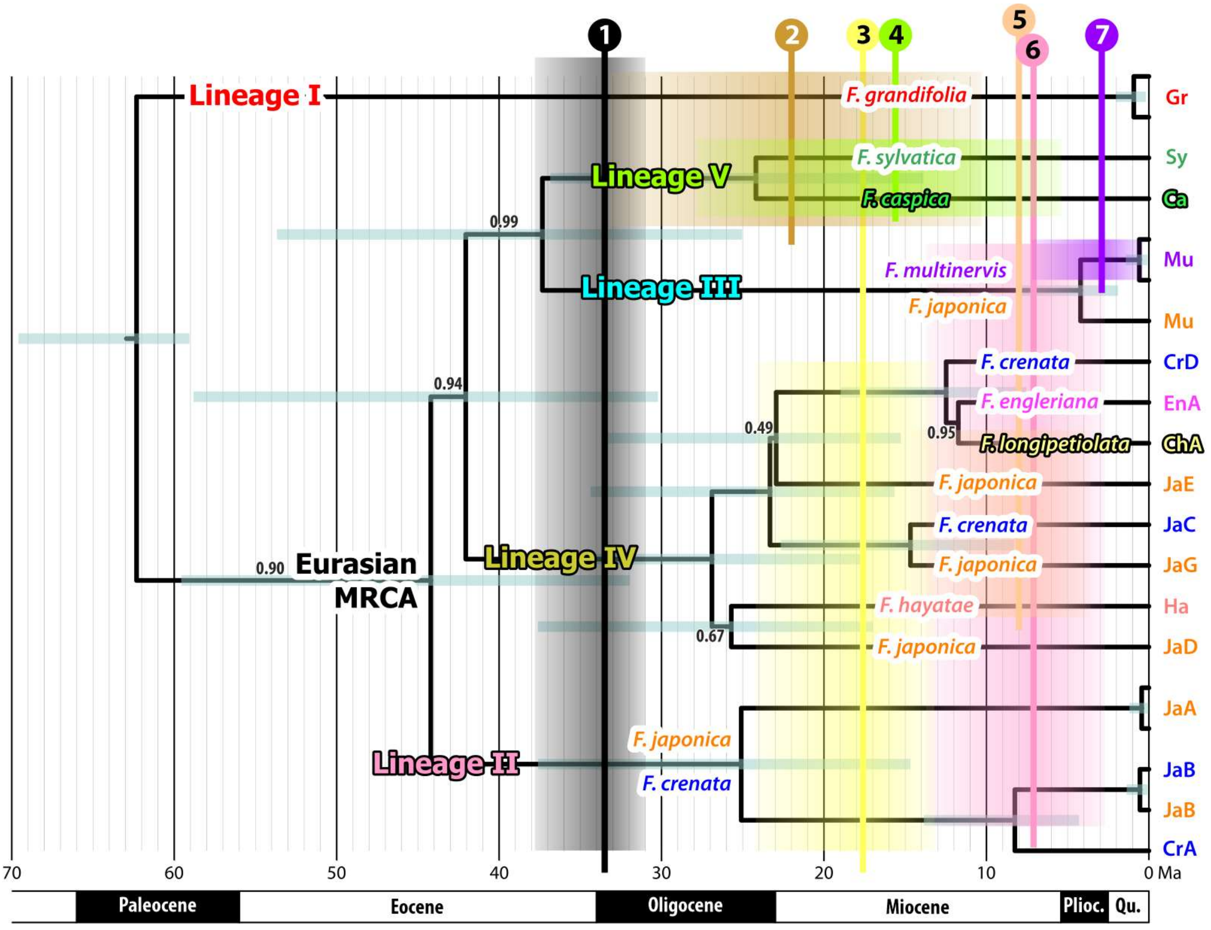


**Figure SR3-2 | Overlaid chronograms using different root-age constraints for plastid and nuclear divergences**: initial plastid divergence correlated to trans-continental, high-latitude Paleocene radiation of modern beech (N.E. Siberia to W. Greenland; cf. **Section 4**); subgeneric-nuclear split (❶) matched to fossil record: near-coeval appearance of fossils at the Eocene-Oligocene boundary that can be attributed to either *Fagus* subgenus *Fagus* or *F.* subgenus *Englerianae*. Roman numerals: main plastome lineages; biogeographic regions abbreviated as in **Fig. SR3-2.** Plastid chronogram shown in full, nuclear chronogram reduced as vertical fields: ❶ subgeneric split; ❷ last contact between North American and Western Eurasian lineages of *F.* subgenus *Fagus*; ❸ first recorded divergence within the East Asian-subgenus-*Fagus* clade; ❹ radiation starts within Western Eurasian lineage (here: stem age of *F. crenata*); ❺ split between *F. hayatae* and *F. longipetiolata*; ❻ *F.* subgenus *Englerianae* crown age; ❼ isolation of *F. multinervis.*

The notable exception from the general pattern of temporal and phylogenetic decoupling of plastomes and species lineages is the West-Eurasian lineage where the 95% HPD intervals still overlap to a large degree and the medians indicate a mid-Miocene MRCA of *F. sylvatica* and *F. caspica* and late Oligocene divergence between their V-Ca and V-Sy Lineage V plastomes. Albeit being a crude dating experiment, this result is in full agreement with the palaeogeographic and -taxonomic history of the Western Eurasian beeches (summarised in **Fig. SR4-3** in **Section 4**) as well as the modern-day situation (four species characterised by unique genetic features as well as genetic clines; Cardoni et al. 2022, Denk et al. 2024).

#### 3.2 Association of fossils with plastid lineages

##### 3.2.1 Theoretical-conceptual conundrum

Plastome differentiation in beech and other Fagaceae/Fagales requires geographic disjunction, which facilitates speciation. Heterogeneity observed in the plastomes of the same (empirical) species, hence, can have two evolutionary causes: *(i)* limitation of seed dispersal leading to isolation in course of vicariances and culminating in allopatric speciation or *(ii)* evolutionary

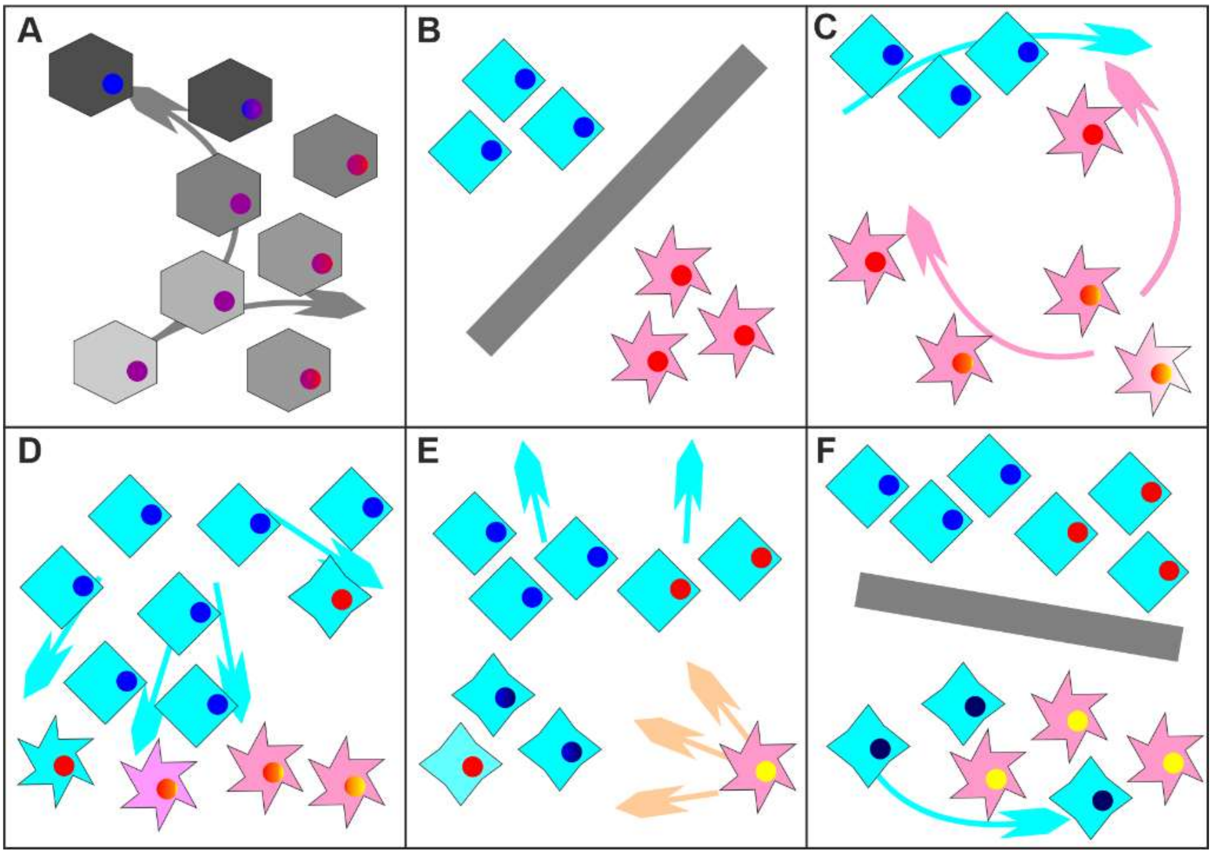

**Figure SR3-3 | "Chloroplast capture" via asymmetric introgression.** A species' populations (signature form representing populations' phenotypes, colour their nucleome genotype) expand into a large area, leading to a genetic gradient in their plastomes (internal red to blue dots replacing the ancestral purple; **A**) culminating in allopatric speciation: a northwestern species (diamonds, blue plastomes) and southeast-bound species (stars, red plastomes; **B**). **C.** External factors (e.g. global warming) allow the stars to expand their distribution area while weaken the contact between their populations leading to an increased intra-species drift reflected firstly in their plastomes inherited from only their mother trees (propagated via short-dispersed by sedentary animals seeds in case of beech). **D.** A change (e.g. global cooling) favours the expansion of the diamonds, which start to introgress into surviving star populations leading to a complete take-over (enforced by backcrossing of introgressed star-offspring with their pure-diamond parents) of the most remote ones; and retreat of the remainder of the star-species, which undergoes increased genetic drift (yellow plastome). **E.** Reset of the climate forces the majority of the diamond populations to retreat to higher latitudes. Intra-species homogenisation leads to the complete loss of any sign of the early introgressed northern stars' parentage in the east, except that they carry and have propagated the eastern, "captured", red plastomes. At low latitudes a similar plastid dimorphism establishes but only populations of introgressed stars survive. Because of their relative rarity and lower active population size, their red plastomes get eventually lost while their blue plastomes have drifted (dark blue circles), when the final isolation and allopatric speciation within the diamonds is terminated (**F**); the descendants of the southern introgressed stars, phenotypically, nuclear-genetically largely and plastid-wise unambiguous members of the diamonds may co-exist with their genetically now most distinct red star cousins.

reticulation, the crossing of lineages via introgression and hybridisation: hybrid speciation ("chloroplast capture" in phylogenetic literature).

In beech, hybrid speciation can be expected to happen in course of any large-scale dispersal event that leads to secondary contacts of beech species (**Figs SR3-3**, **SR3-4**). Phylogenetically, the two processes may be or may not be distinguished by mapping plastid types on nuclear phylograms: vicariance or isolation of a remote mother population will always lead to sister plastid haplotypes; evolutionary reticulation may involve sisters (characterised by sister haplotypes) but also more distant cousins. Moreover, extinctions leading to the loss of a maternal lineage and corresponding plastid haplotypes, may obscure both events (**Fig. SR3-5**).

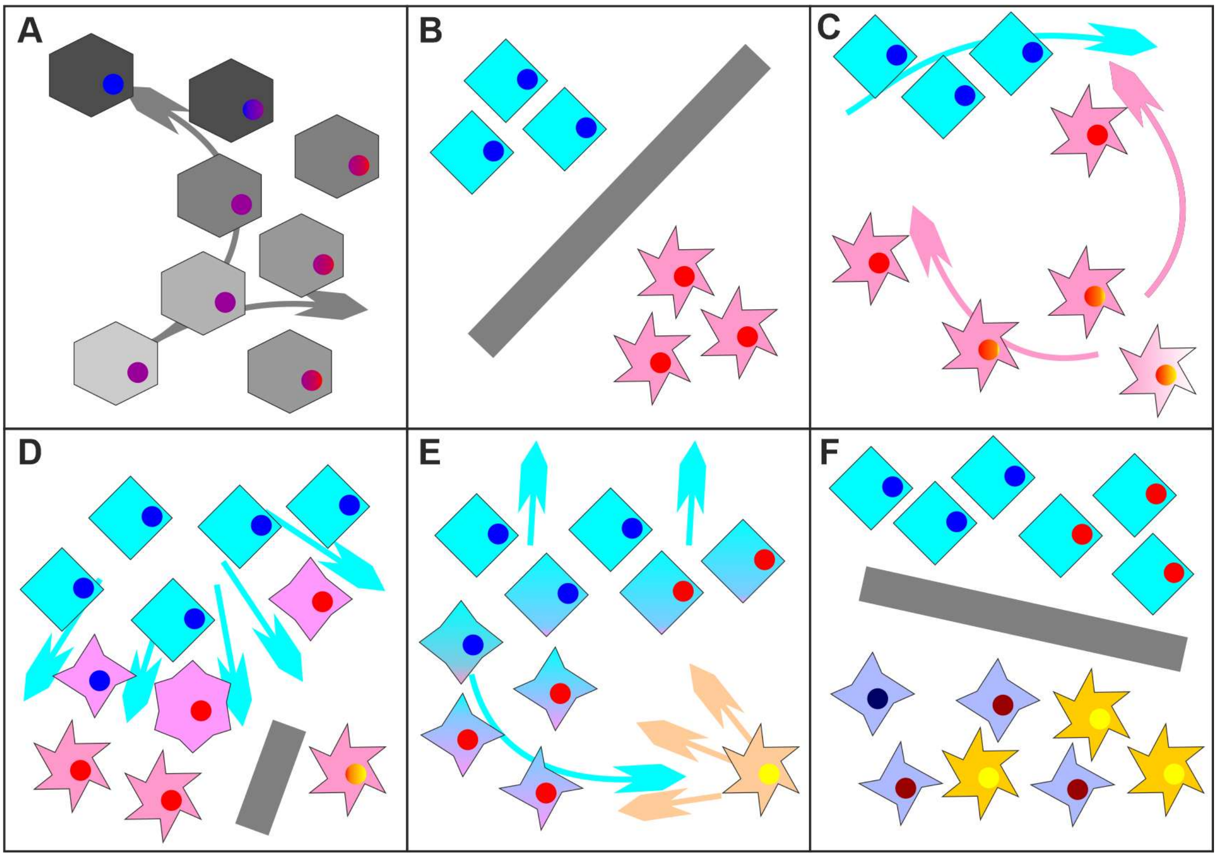


**Figure SR3-4 | "Chloroplast capture" via hybridisation.** Same set-up as before for the asymmetric introgression scenario (**A–C**), only that the two species, diamonds and stars, form ± stable hybrids, a hybridisation zone/swarm, when coming into contact (purple populations with intermediate outlines, **D**). Within the hybrid zone, individuals and later populations can include any combination of diamond and star phenotypic and genotypic traits (**D**). The boundary between the hybrid species (4-tip stars) and their sym- then parapatric parent (the diamonds) will remain permeable for some time, allowing original star-genes to introgress into the diamond core population (colour gradients in **E**). The final isolation leads to a similar pattern of phenotypic, nucleome and plastid diversity than for the asymmetric introgression scenario (**Fig. SR3-3**), only that now the hybridogenous species (4-tipped stars) have retained a plastid polymorphism stemming from introgressed (pure-)diamond populations and are characterised by drifted diamond-origin and star-origin plastomes (dark blue and dark red circles).

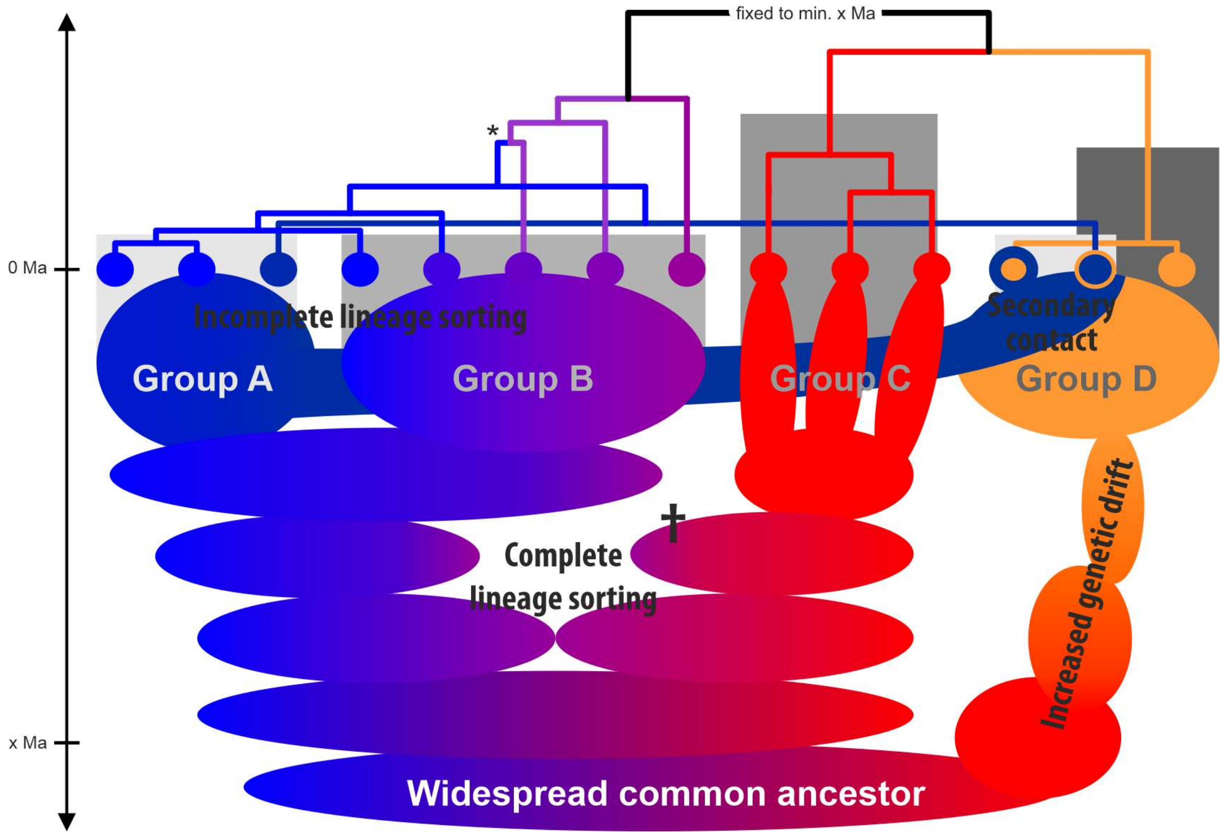


**Figure SR3-5 | Schematics of evolutionary events leading to the gain or loss of plastid haplotype lineages within a species lineage (taxonomic decoupling).** The resultant plastid chronogram at the top can reflect the fragmentation sequence of the plastome pools but not the speciation events or the interspecies phylogenetic relationships: the plastome tree is decoupled from the species tree. The grey fields represent the according (quasi-)holophyletic species groups. The situation in Group D would be addressed as "chloroplast capture" but simply represents the hybrid origin of some of its members: they share exclusive common ancestors with the remainder of Group A (periphyletic according Wheeler 2014) *and* (epiphyletic) Group D. Secondary contacts are often asymmetric, hence, introgressed members of Group D may or may not have intermediate/mixed phenotypic and nuclear-genotypic traits. They may also pass just as members of Group A. A future bottleneck may even eliminate the original Group D plastomes, thus, removing any evidence of the orange lineage except for surviving plastomes in members of Group A.
* Defined by two independently drifted plastid blue sublineages, each of which is closer to the main plastid type than they are to each other, this branch will typically receive ambiguous support while being elongated.

##### 3.2.2 Palaeobiogeographical framework

In plant and animal phylogenetics, well-dated trees can only rely on a fossil record that can be instrumentalised as age priors: fossils that can be assigned to clades in the molecular trees and can inform minimum ages for node dating (ND) or a lineage's age distribution for the "fossilized-birth death" (FBD) dating. In most optimal cases (only possible for trait-rich animal groups), individual fossils can be used as extinct tips that are placed within the molecular tree using a morphological partition for extant and extinct tips (total-evidence dating). When the plastid genealogy is largely decoupled from the nuclear genealogy, the species coalescent (in the case of beech rather a network than a tree), and the phenotype of nuclear-coherent groups and species, the association of fossils in an FBD dating framework using plastid data require a prior (palaeo-)biogeographical hypothesis ('TaxSets' listed in **SupplFossilTable.xlsx**, sheet *cpFBD*) as outlined below (see also main text and **Sections 1, 2**).

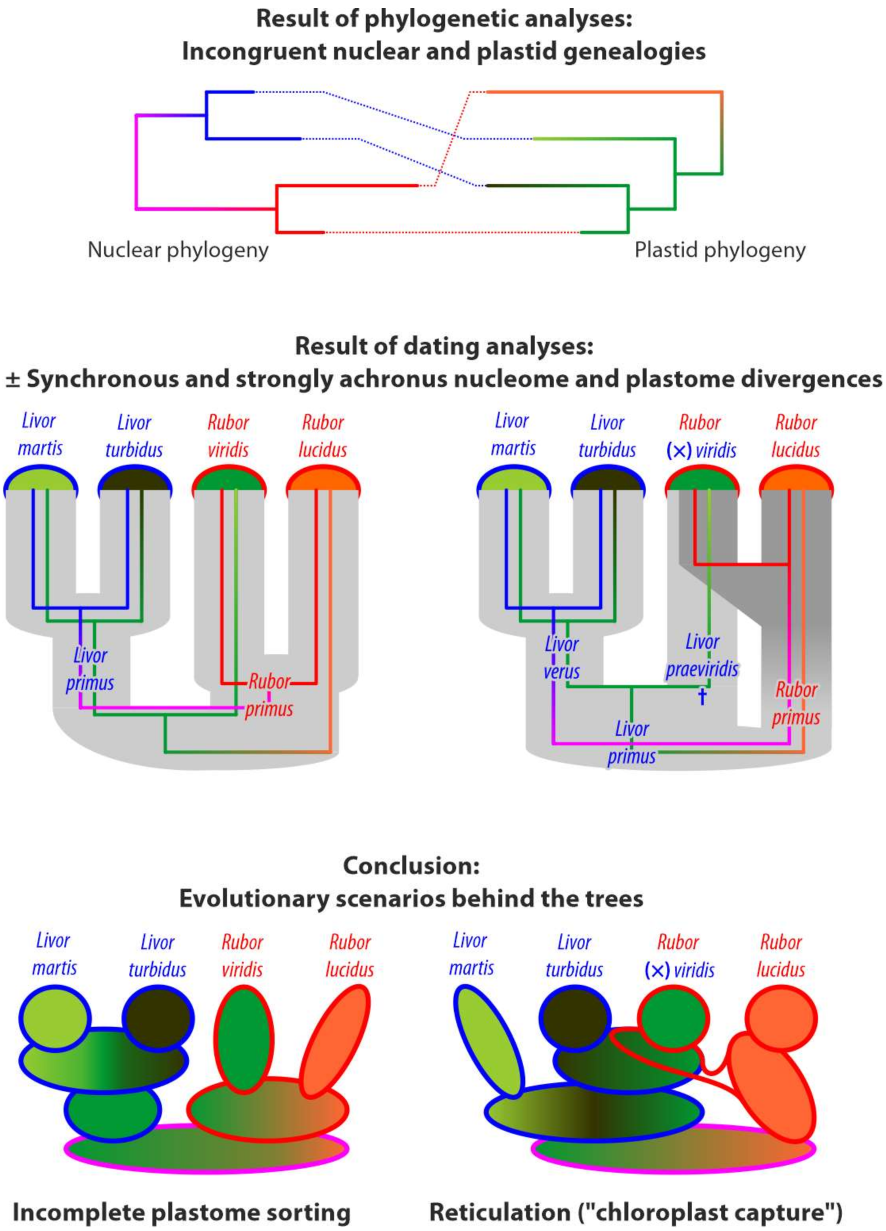


**Figure SR3-6 | How dating can change the interpretation of nuclear-plastid topological incongruences.**
**A.** A tanglegram illustrating the incongruence between nuclear and plastid data-based genealogies. The two species of the blue clade representing the taxon *Livor* (a hypothetical genus) have green lineage plastomes, while the two species of its red sister taxon (genus *Rubor*) either have a strongly distinct orange or a (nested, short-branching) green plastome.
**B, C.** Two possible outcomes (chronograms, dated trees) of explicit dating analyses. The plastid and nuclear divergence may be resolved as ± synchronous (**B**), i.e. relating to the same evolutionary episodes (taxonomic and geographic radiations) or (strongly) achronous, i.e. temporally decoupled (**C**) and involving past speciation events within one lineage (†*Livor praeviridis* | *L. verus*) and reticulation because of secondary contact of sister/cousin lineages (*Rubor viridis* originating from massive introgression of the *R. primus* lineage into *L. praeviridis* lineage).
**D, E.** Based on the dating results and adding further context (palaeogeographic and -ecological framework, exploratory data analysis of nuclear divergence patterns) one may be able to preclude certain evolutionary scenarios. The synchronicity observed in B fits to an incomplete lineage scenario, with the modern-day species representing old (in *Rubor*) and young speciations (*Livor*), and having evolving from either a geographically restricted and homogenous or widespread and plastome-diverse progenitor species (**D**). The much younger divergence between the *Rubor* species compared to that of the plastomes they carry can only be explained by secondary contact and lineage crossing. *Rubor viridis*, originating from *Livor* mother trees introgressed by members of the *Rubor primus* lineages have conserved the only molecular evidence for another and ancient speciation event within the *Livor* lineage (**C**,**E**).

**TaxSets 'American_lineage' including modern-day Lineage I vs. 'Eurasian_lineage'**— Modern-day beeches show five main plastid lineages (Lineages I–V; **Section 2**); the most distinct (Lineage I) is restricted to the eastern North American species. Primary and deep New World | Old World plastome splits can be observed across the entire Fagaceae family (Zhou et al. 2022) and the order Fagales (**Fig. SR3-7**).

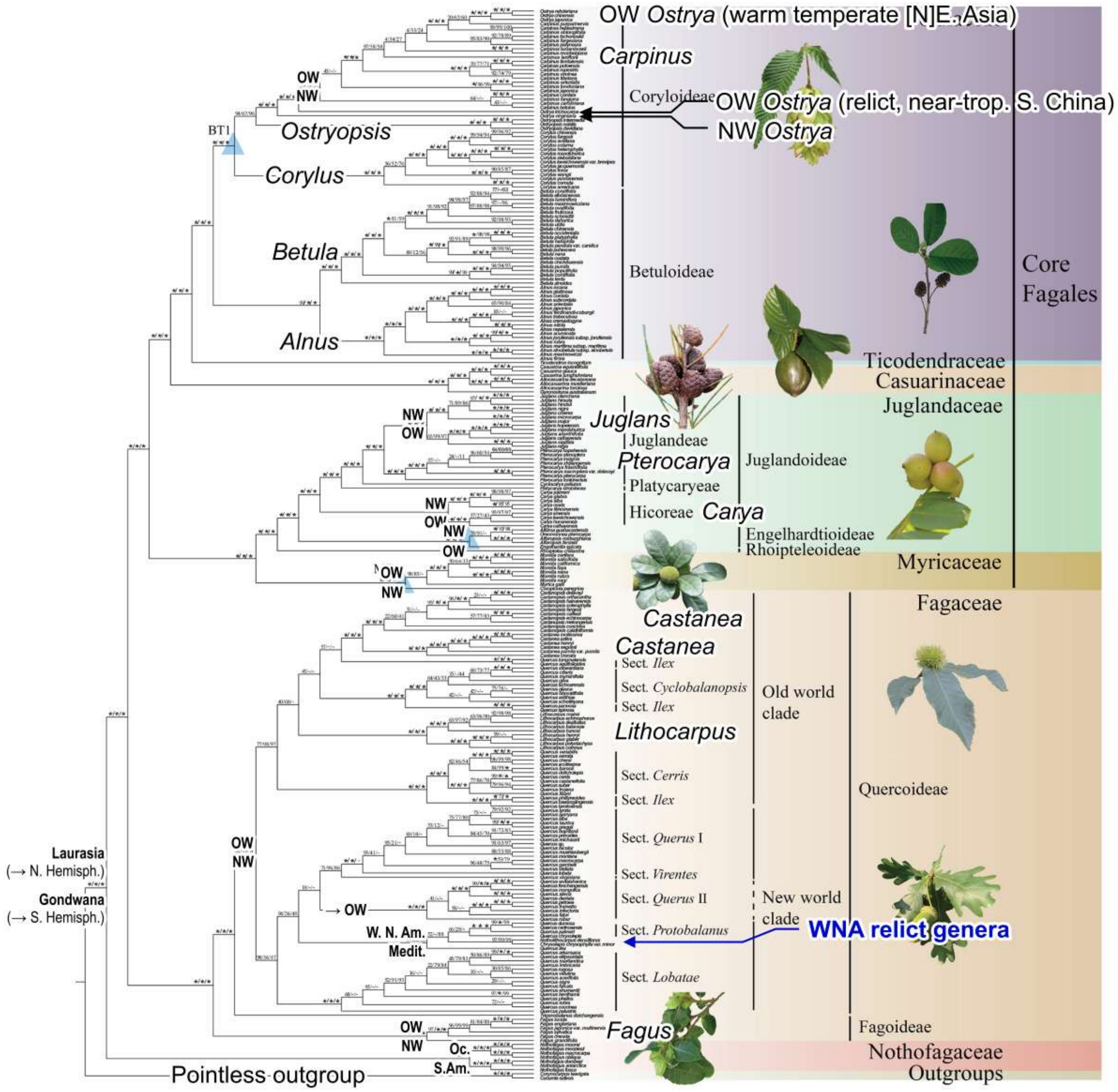


**Figure SR3-7 | Yang et al.'s (2021), fig. 1, with crucial missing information added.** Main geographic splits imprinted in Fagales plastomes in congruence with nuclear phylogenies: (*i*) Gondwana (Nothofagaceae) | Laurasia (all other Fagales), (*ii*) Oceania (Oc.) | South America (Andes and west of them; S. Am.) within Nothofagaceae, Late Cretaceous; (*iii*) within Myricaceae; and (*iv*) Juglandoideae genera. Geography-correlated plastome splits in (stark) conflict with systematic relationships and the nuclear genealogy: Old World (Eurasia; OW) | New World (Americas; NW) splits within (*v*) *Fagus,* (*vi*) core Fagaceae (mislabelled "Quercoideae"), (*vii*) Engelhardioideae (misspelled "Engelhardtioideae"), (*viii*) *Ostrya* and (*ix*) Mediterranean (Medit.) oaks | western N. American (W. N. Am.) Castanoideae.

Thus, we treated any fossil from eastern North America (ENA) as carrying Lineage I beech plastomes, as well as those of the northern North Atlantic (NAT, extinct region; see also **Section 4: Fig. SR4-2**). Their Old World counterparts were associated to the sister clade of Lineage I, the clade comprising all Lineage II to V plastomes and Eurasian-Beringian fossils. Fossils from western North America (WNA), representing an extinct lineage of New World beeches with ± affinity to the modern-day ENA species and co-eval fossils from Beringia (BER) and North-East Asia (NEA), were not assigned to any of the plastid clades and TaxSets. They may have carried plastomes representing a sister lineage of ENA Lineage I or more closely related to the Eurasian lineages (Lin. II–V, following section).

**TaxSet 'Oceanic_lineage'/Lineage II plastomes**—During most of the Eocene and Oligocene, the Beringian Land Bridge connected western North America with North-East Asia; extinct, similar morphotypes can be found along the entire northern Pacific coastline and its hinterland (see also Denk & Grimm 2009). It is hence likely that Lineage II represents the original plastome of coastal northern Pacific beech populations and was carried or picked up by the northeastern Japanese beeches when they migrated into their modern-day area. As there are no survivors of stem beeches on either side of the northern North Pacific, we cannot rule out that the earliest beeches of this region and adjacent western North America (WNA, e.g. the Eocene †*F. langevinii* Manchester & R.M.Dillhoff) carried a different, now extinct lineage of beech plastome. Accordingly, we associated only those northern North Pacific fossils (Beringia, BER) with Lineage II that represent or show affinity to the modern-day subgenera.

**TaxSet 'Continental_lineage'/Lineage III–V plastomes**—For East Asia, two scenarios are conceivable: (*i*) Lineage III is the relict of mid- to high-latitude, ± continental beech populations (northern Asia, NAS, Central Asia, CAS; cf. **Section 4**), separated by mountain chains and early intra-continental pre-steppe regions from the North-East Asia (NEA) biodiversity hotspot, home of Lineage IV; (*ii*) Lineage III is the genuine haplotype of NEA, and Lineage IV represents populations from outside NEA and later (re-)migrated into the region. Based on the phylogenetic position of Lineage IV, possibly related to Lineage III but distant from Lineage V to the west, and with respect to the observation that the most-distinct haplotype within Lineage IV is from the Taiwanese island isolate *F. hayatae*, southern East Asia (SEA) is more plausible as source area for Lineage IV plastomes than CAS.

At 23–21 Ma, crown-group radiation must have set in, ± simultaneously in lineages II and IV, either triggered by multiple speciation events, isolation of geographically relatively close populations (genetic drift via bottlenecks), or massive range extension (genetic drift via geographic distance). Morphologies evolved at that time were passed on into modern-day continental Eurasian species (CAS †*F. altaensis* Kornilova & Rajushkina → SEA *F. lucida*; NEA †*F. protolongipetiolata* [A] Huzioka → SEA *F. longipetiolata*) but apparently not their plastid signatures. The latter, plastid signatures from ancient allopatric speciation events, only survived in the precursor(s) of *F. japonica* (NEA), which conserved this ancient diversity until today. The last East Asian range expansion appears to be directly linked to the latest radiation in *Fagus* subgenus *Fagus*, the establishment and isolation of the modern-day East Asian morphotypes and species, *F. crenata, F. longipetiolata,* and *F. lucida*.

To minimise bias, we hence did not distinguish between the two East Asian Lineages III and IV, and assigned all Eurasian fossils with affinity to either subgenus (WEA + CAS + NEA + SEA) to the Lineage III–V clade (TaxSet 'Continental lineage') and only discerned two sub-clades within it:

**TaxSet 'Southern_lineage'/IV-Ha and IV-LuA Lineage IV plastomes**—The Lineage IV plastomes of *F. hayatae* and their sister plastome reported for a south-central *F. lucida* individual are notably distinct from those of the other East Asian species, reflecting an ancient divergence (cf. **Section 2.2**). Our working hypothesis is that these plastomes are the remainder of the original low- to mid-latitude beeches of mountainous East Asia. Our fossil dataset includes three macrofossil species from the Late Miocene to Pliocene of southern East Asia (SEA), two of which are (very) similar to the modern-day *F. longipetiolata* and *F. hayatae-pashanica.* Given their place and time (Jiangxi, Guizhou provinces; Taiwan) they most probably carried the same plastid lineage as the modern-day *F. hayatae*.

**TaxSet 'WestEurasian_lineage'/Lineage V plastomes**—Being the only aspect of the beech species network showing high congruence across nuclear, plastid and phenotypic diversity patterns, it is trivial to assume that all fossils of the Western Eurasian lineage (†*F. castaneifolia* in CAS and †*F. castaneifolia,* †*F. haidingeri* Kováts and †*F. gussonii* Massalongo*,* Western Eurasia, WEA) can be linked to the Lineage V plastome clade.

#### *3.3 A dated species network of beech*

##### 3.3.1 General limitations

As expected, the FBD- and node-dated (ND) estimates differ strongly from each other for the most part, with the meta-calibrated FBD estimates being double as high as the conservatively calibrated ND estimates for most internal nodes in case of the nuclear dated trees

(**Table SR3-1**). Keeping in mind that ND estimates at best represent minimum divergence ages, the ND estimates make, overall, more sense than the FBD estimates based on what we already know about beech (fossil record, evolution of Earth's surface and climate during the Late Cretaceous and Cainozoic).

**Table SR3-1 | 'Fossilized-Birth-Death' (FBD) and node dating (ND) estimates for each optimised topology.** For the full list see **SupplDating.xlsx**, sheet *final dating*. ∆ = Difference in medians, rounded to full myrs.

| Node | Topology | FBD | ND | ∆ |
|---|---|---|---|---|
| MRCA of all extant plastomes | Plastid | 66.0 (73–64) | Set to ≥ 63 Ma | -3 |
| Eurasian MRCA | Plastid | 43.6 (49–40) | Set to ≥ 36 Ma | -7 |
| MRCA of Lin. III–V | Plastid | 34.5 (35–34) | 31.4 (33–31) | -3 |
| MRCA of Lin. IV | Plastid | 19.0 (20–19) | 22.3 (27–18) | +3 |
| MRCA of all extant spp. | Atl \| Pac | 60.8 (66–56) | Set to ≥ 34 Ma | -26 |
| | High \| Low | 67.7 (77–63) | Set to ≥ 34 Ma | -33 |
| | NW \| OW | 73.0 (89–63) | Set to ≥ 34 Ma | -37 |
| MRCA of *F.* subg. *Englerianae* | Atl \| Pac | 5.7 (9–3) | 5.8 (11–3) | ±0 |
| | High \| Low | 18.8 (15–14) | 9.7 (17–5) | -9 |
| | NW \| OW | 6.7 (10–5) | 5.5 (11–2) | -2 |
| MRCA of *F.* subg. *Fagus* | Atl \| Pac | 51.8 (59–46) | 29.4 (35–22) | -22 |
| | High \| Low | 51.5 (59–46) | 31.0 (36–26) | -21 |
| | NW \| OW | 59.1 (73–48) | 36.5 (59–30) | -24 |
| MRCA of W. Eurasian spp. | Atl \| Pac | 24.8 (33–18) | 14.3 (22–6) | -10 |
| | High \| Low | 33.1 (38–37) | 17.3 (26–11) | -16 |
| | NW \| OW | 30.5 (43–20) | 24.8 (33–13) | -6 |
| MRCA of E. Asian spp. | Atl \| Pac | 39.8 (44–36) | 18.8 (22–16) | -21 |
| | NW \| OW | 41.1 (46–37) | 27.2 (32–24) | -14 |

Our FBD estimates, as well as those by Renner et al. (2016), are probably overestimating major speciation events, the crown ages of both subgenera and Old World vs. New World lineages and the stem ages of (some) modern-day species lineages, for two reasons. First, by using the entirety of the fossil record, our lineages are oversampled with respect to the FBD model: in the FBD model, each fossil is regarded to reflect a speciation event leading to an additional tip, which is not necessarily the case for beech. To comprehensively compile the fossil record of beech, we ensured that differently time-and-place-stamped fossils of the same fossil species have been included. As we have no possibility to judge whether the same fossil species represents a single (empirical) species in space and time and not a complex of ± closely related species (ancestors and their descendants and/or sister species), we refrained from reducing the fossil sample to one fossil per fossil species for the FBD dating.

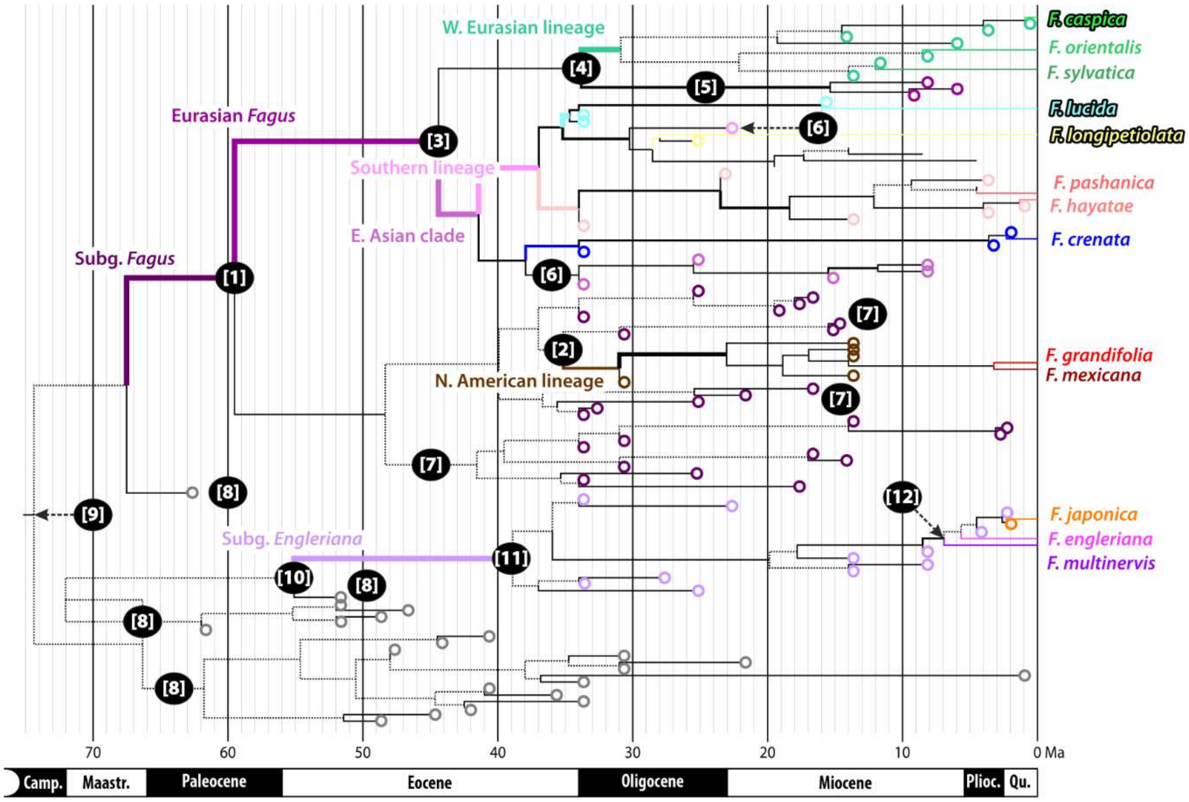


**Figure SR3-8 | Example for modelled (artificial) extinct sister lineages in the FBD chronograms**. Shown is a full maximum clade credibility tree, with fossils semi-arbitrarily, i.e. within the constraints of the topological priors, added as additional tips by the FBD algorithm. The crown age of *Fagus* subgenus *Fagus,* equalling the MRCA of its N. American and Eurasian subclade*,* is placed in the Paleocene **[1]**, predating the first unambiguous fossils of both modern-day subgenera (coloured circles) and the modelled stem age of the N. American subclade of *F.* subgenus *Fagus* **[2]** by ~25 myrs. The extant MRCA-based stem age of the W. Eurasian lineage **[3]** is placed ~10 myrs before the modelled stem age **[4]** involving a modelled extinct sister clade **[5]** collecting young fossils (†*F. gussonii* fossils from W. Eurasia, < 10 Ma old) assigned to the Eurasian *Fagus* clade but not any of its subclades. These fossils are modelled as a sister clade to the W. Eurasian lineage rather than the East Asian-subgenus-*Fagus* clade (EASF) because they, and the fossils assigned to the W. Eurasian clade, are generally younger than fossils assigned to the EASF clade **[6]** or its modern-day species lineages. All fossils assigned to *F.* subgenus *Fagus* but not any of its sublineages are modelled as extinct sister lineages of the N. American lineage **[7]**, which, in addition to the *(i)* Eocene-Oligocene age of fossils assigned to Eurasian species lineages and *(ii)* the genetic dissimilarity between N. American and Eurasian members of *F.* subgenus *Fagus* and older fossils not assigned to either subgenus **[8]**, accounts for the probably too old crown age of *F.* subgenus *Fagus* **[1]**. The latter also trigger a Campanian root age **[9]**. Being not part of either subgeneric lineage but numerous and covering a large time span, the FBD algorithm models them as speciation events linked to extinct beech lineages, placing them in between both subgenera **[8]**. Accordingly, the modelled stem **[10]** and crown ages **[11]** of *F.* subgenus *Englerianae* are again much younger than the tree's root age equalling the MRCA of all modern-day species and both subgenera, i.e. the MRCA-based stem age of both subgenera. The temporal distribution of fossils of *F.* subgenus *Englerianae* furthermore contrasts with the genetic similarity of its modern-day species expressed in a very young MRCA-based crown age **[12]**.

The modern-day genetic mosaic implies that a lot of speciation processes in beeches are cryptic or pseudo-cryptic and strongly influenced by phenotypically untraceable secondary contacts. On the other hand, we have today widespread species like *F. sylvatica* and *F. grandifolia* showing very little intra-species differentiation; they are morphologically and genetically homogenous. Identical or highly similar fossils from the same time period but different provenances may hence belong to the same species (FBD models too many speciation events and will be overestimating) or different species (FBD estimates will be more to the point). Nonetheless, oversampling within the conceptual framework for FBD dating expresses itself in the resulting chronograms; all of which include modelled extinct sister lineages to accommodate the high number of fossils per time period (example shown in **Fig. SR3-8**).

Second, and more importantly, phenotypic affinities likely have different systematic quality, the further one goes back in time. For instance, fossils highly similar to identical to modern-day *F. crenata* from the Messinian to Pleistocene of Japan most probably come from plants that were direct precursors of the modern-day populations of *F. crenata* and, thus, inform the age distribution of the *F. crenata* lineage in an FBD framework. However, first fossils with unambiguous affinity to *F. crenata* have been recorded from the early Oligocene Kraskino palaeoflora from the Russian Far East (described as †*F.* 'cf. *palaeocrenata*' by Pavlyutkin et al. 2014, cf. **SupplFossilTable.xlsx**, sheet *main list*), where they co-occur with various other modern phenotypes. In a consistently applied FBD framework, where fossils are assigned to clades according to their morphological affinities, these fossils extend the age distribution of the *F. crenata*-lineage beyond the Oligocene-Eocene boundary, irrespective of the topology that was dated (**Fig. SR3-8**; **Table SR3-2**).

**Table SR3-2 | Stem and (modelled) crown age of the *F. crenata* lineage, inferred using FBD dating.** Listed are median divergence ages, brackets give the 95% highest probability density interval (rounded to 1 myrs).

| Topology | Sister lineage | Stem age Modelled | MRCA-based | Crown age Modelled |
|---|---|---|---|---|
| Atlantic \| Pacific | *F. longipetiolata* | 34.6 (36–34) Ma | Same | 34 Ma[a] |
| High- \| Low-latitude | W. Eurasian lin. | 35.1 (38–34) Ma | Same | 34 Ma[a] |
| New World \| Old World | Southern lin.[b] | 37.6 (42–34) Ma | 41.1 (46–37) Ma | 34 Ma[a] |

[a] Directly informed by †*F.* cf. *palaeocrenata*: in every sampled Bayesian-inferred chronogram, the fossil marks the onset of speciation within the *F. crenata* lineage (hence the 95% HPD interval has a width of zero).
[b] The 'southern lineage' comprises all other extant E. Asian species of *F.* subgenus *Fagus* and a number of fossils with affinities to them including †*F. galbanifolia.*

Since unambiguous *crenata*-like fossils have not been reported from the Oligocene and Early to Middle Miocene, *F.* cf. *palaeocrenata* may as well *(i)* represent a temporal parallelism, i.e. a cousin lineage thriving in a similar niche than modern-day *F. crenata*, or *(ii)* an ancestral (stem) lineage of *F. crenata* (and other East Asian species), which's phenotypic legacy has been conserved in *F. crenata*. Which is the case, is impossible to ascertain. Since modern-day beech species are genetic mosaics, they carry signals of a highly complex and dynamic reticulate evolution (hence, the use of three topologies for the nuclear dating). Even if extant species do not hybridize anymore, (some of) their precursors have done so in the past, and repeatedly. Some gene regions may reflect different speciation event as others, and some may have been homogenised by more recent introgression and backcrossing. The last common mothers (plastid genealogy, unknown part of the nuclear genealogies) and fathers (nuclear genealogies), which we model during dating may be of hybrid origins themselves. Past hybrids, especially ancient ones showing less evolved modern traits in general, would not be recognizable and discernible in the fossil record as such. Lastly, as a tree genus excelling in its climax niche, beeches are phenotypically restricted in their traits (Shen 1992, Denk 2003); parallelisms are likely but, in addition, hybrids and introgressed populations, will often be cryptic to some degree. We may have cases where a distant cousin lineage is phenotypically more similar to a modern-day taxon than its precursors as well as cases where the phenotypic similarity reflects a topological aspect of the species coalescent network only captured by a (small) portion of the available genetic data.

Rather than representing an evolutionary dead-end and temporal convergence (or parallelism), †*F.* cf. *palaeocrenata* could be an exclusive ancestor of *F. crenata,* which's genetic legacy has been largely lost or overprinted in the modern-day species' nucleome due to later speciation events (involving reticulation with other modern species lineages) but which passed on unique, beneficial phenotypic characteristics. Characteristics that allowed beeches of the *F. crenata* lineage to thrive in a niche not occupied or available to their siblings, i.e. reflecting 'hybrid vigour' (McKown & Guy 2018) of a or the ultimate *F. crenata* precursor. That some of their ancestors were not shared with other extant species of *Fagus* subgenus *Fagus* is evident by the plastomes carried by modern-day *F. crenata* (**Section 2.2.2**; **Figs SR2-6**, **SR2-8**). Also in such a case, assigning †*F.* cf. *palaeocrenata* to the *crenata*-lineage would overestimate the stem age of this species lineage, the latter being defined by the divergence between *F. crenata* and its extant sister lineage(s) (the West-Eurasian spp. and the southern East Asian spp. of *F.* subg. *Fagus*). Nonetheless, it could inform an ancient split between *F. crenata*-specific plastome and the plastid lineage characteristic for the last common ancestor of the remaining East Asian or Eurasian species of *F.* subgenus *Fagus.* Also, some nuclear genes support a 'Southern lineage', a common ancestry of all other East Asian species but *F. crenata*; a signal that may relate to the same speciation/radiation event. †*Fagus* cf. *palaeocrenata* may be part of a North-East Asian species (lineage) within or outside *F.* subgenus *Fagus* that later fused with an East Asian member of the modern-day *F.* subgenus *Fagus* forming the or a precursor of the modern-day *F. crenata* (**Fig. SR3-9**).

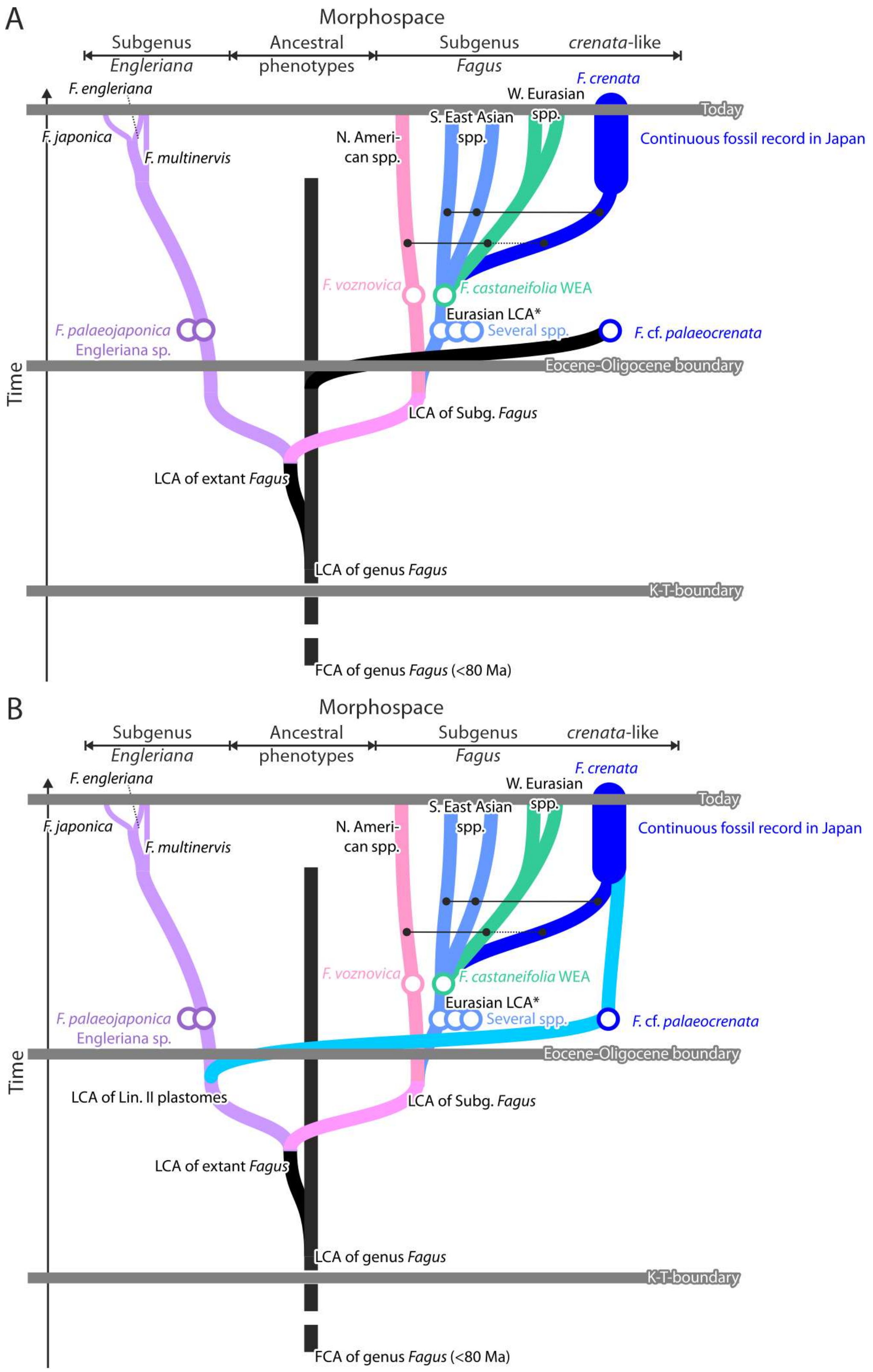
A
Morphospace
Subgenus Engleriana
Ancestral phenotypes
Subgenus Fagus
crenata-like
F. engleriana
F. crenata
W. Eurasian spp.
S. East Asian spp.
Today
F. japonica
F. multinervis
N. American spp.
Continuous fossil record in Japan
F. voznovica
F. castaneifolia WEA
Eurasian LCA*
Several spp.
F. palaeojaponica
Engleriana sp.
F. cf. palaeocrenata
Time
Eocene-Oligocene boundary
LCA of Subg. Fagus
LCA of extant Fagus
LCA of genus Fagus
K-T-boundary
FCA of genus Fagus (<80 Ma)
B
Morphospace
Subgenus Engleriana
Ancestral phenotypes
Subgenus Fagus
crenata-like
F. engleriana
F. crenata
W. Eurasian spp.
S. East Asian spp.
Today
F. japonica
F. multinervis
N. American spp.
Continuous fossil record in Japan
F. voznovica
F. castaneifolia WEA
Eurasian LCA*
Several spp.
F. palaeojaponica
Engleriana sp.
F. cf. palaeocrenata
Time
Eocene-Oligocene boundary
LCA of Lin. II plastomes
LCA of Subg. Fagus
LCA of extant Fagus
LCA of genus Fagus
K-T-boundary
FCA of genus Fagus (<80 Ma)

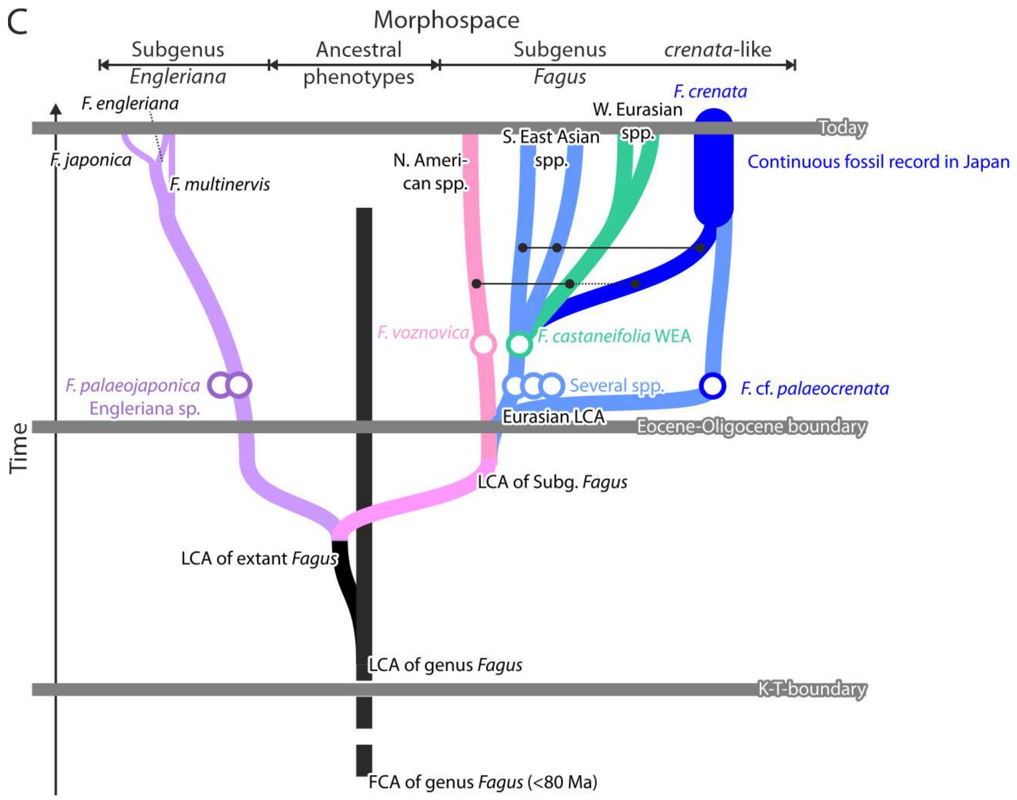


**Figure SR3-9 | Scheme illustrating possible placements of †*F.* cf. *palaeocrenata* in relation to modern-day *F. crenata* and its use in FBD or ND approaches. A** (preceding page)**.** The *crenata*-like Kraskino fossils represent a temporal convergence: an early and extinct beech lineage occupying a similar niche than modern-day *F. crenata* that had evolved the same leaf morphology*.* Both ND, fossil informing minimum stem age of the *crenata*-lineage, and FBD estimates using these fossils would be erroneous by default and overestimating. **B, C**. The fossils represent one of the evolutionary sources of modern-day *F. crenata* from a lineage outside or inside of *Fagus* subgenus *Fagus.* In this case, the fossil could, using ND, inform a minimum age for the e.g. the plastome Lineage II crown age (**B**, preceding page) or the crown age of the Eurasian subclade of *F.* subgenus *Fagus* (C); overestimation using FBD could be avoided by re-assigning the fossil to the according larger clade (cf. Table **SR3-3**).

Assigning †*F.* cf. *palaeocrenata* to the *F. crenata* species lineage may be as wrong as not assigning it, always pending the differentiation signal in the used molecular data matrix.

A possibility to remedy this conceptual problem would be an iterative FBD approach, in which putatively too old fossils such as †*F.* cf. *palaeocrenata* from the Kraskino palaeoflora would be subsequently removed from a lineage (with respect to topological aspects of the date trees) and assigned to the next larger clade. The result of an according experiment is shown in **Table SR3-3**. The new estimates would then be checked again, and further fossils moved. However, such an iterative reinterpretation of fossils would add a strong implicit bias to the chronogram with each iteration, already meta-calibrated due to the high number of fossils used to inform the age distribution. Eventually, this would render any explicit dating approach pointless beyond what one would directly deduce by interpreting the fossil record in a systematic-phylogenetic context. Thus, we decided to report the likely overestimating but standard-obtained FBD estimates assigning each fossil strictly by its morphological affinities and their alternatives, obtained by removing the fossils from the Kraskino palaeoflora from the species-level TaxSets.

Traditional node dating has been used as a comparative corrective. ND divergence age estimates rely more strongly on implicit assumptions invoked by the selection of particular fossils as age priors for nodes in the input topologies (as detailed in **SupplM&M.pdf**, section 5.2.1). They can, per se, only inform minimum estimates for divergence ages. In this, they perfectly complement the putatively overestimating FBD-dated divergences.

**Table SR3-3 | Effect on the FBD estimates when treating modern-type Kraskino beech fossils as a temporal convergence.** D = Difference in medians, rounded to full myrs. Blue background: expected younger estimates, ≥ 5 myrs, orange: older estimates obtained by treating the Kraskino fossils as product of ancient phenotypic parallelism.

| Node | Topology | Standard approach[a] | First iteration[b] | Δ |
|---|---|---|---|---|
| E. Asian clade crown age (EAS crown) | Atl \| Pac | 39.8 (44–36) | 37.6 (43–34) | -2 |
| | NW \| OW | 41.1 (46–37) | 36.5 (40–34) | -5 |
| MRCA of *F. crenata-longipetiolata* | Atl \| Pac | 34.6 (36–34) | 26.4 (35–28) | -8 |
| MRCA of *F. longipetiolata-lucida* | High \| Low | 35.9 (40–34) | 35.5 (39–34) | ±0 |
| | NW \| OW | 35.0 (37–34) | 29.6 (34–26) | -5 |
| Modelled stem age of *F. crenata* lineage | Atl \| Pac | = *cr-lo* MRCA | 6.7 (11–5) | -28 |
| | High \| Low | 35.1 (38–34) | 34.0 (34–34) | -1 |
| | NW \| OW | 37.6 (42–34) | 36.5 (40–34) | -1 |
| Modelled stem age of *F. longipetiolata* lin. | Atl \| Pac | 28.9 (33–26) | = *cr-lo* MRCA | -2 |
| | High \| Low | 28.6 (35–25) | 34.0 (34–34) | +5 |
| | NW \| OW | 28.0 (33-26) | 29.6 (34–25) | +1 |
| Modelled stem age of *F. hay.-pash.* lin. | Atl \| Pac | = EAS crown | 28.0 (34–26) | -12 |
| | NW \| OW | 36.7 (40–34) | 24.8 (27–24) | -12 |
| Modelled stem age of *F. lucida* | Atl \| Pac | 37.1 (40–35) | 30.9 (35–28) | -6 |
| | High \| Low | = *lo-lu* MRCA | 34.0 (34–34) | -2 |
| | NW \| OW | = *lo-lu* MRCA | 21.7 (25–20) | -13 |

[a] Modern-type fossils of the Kraskino palaeoflora treated as members of modern species lineages: †*F.* cf. *palaeo-crenata* → *F. crenata* lineage; †*F. koraica* → *F. hayatae-pashanica* lineage; †*F.* cf. *evenensis,* †*F. olejnikovii* → *F. lucida* lineage; †*F. protolongipetiolata* A → East Asian-subgenus-*Fagus* clade.
[b] Kraskino fossils treated as members of *F.* subg. *Fagus.*

Aware of the method-inherent and fossil-triggered uncertainties, we do refrain from naïve reliance on our dating results and reporting impossible precision as commonly seen in molecular dating papers, especially in the case of Fagales (e.g. Sauquet et al. 2012, Xiang et al. 2014, Xing et al. 2014, Larson-Johnson 2016). Instead, we discuss and interpret our FBD-inferred maximum and ND-inferred minimum estimates on the background of the basic molecular differentiation patterns in the used data (**Sections 1.1, 2.2, 2.3**; for nuclear oligogene data see Cardoni et al. 2022, data S5), the palaeogeographic history of beech (time-spans summarised in **Fig. SR3-10**), and the tectonic-climatic history of the Northern Hemisphere (→ **Section 4**).

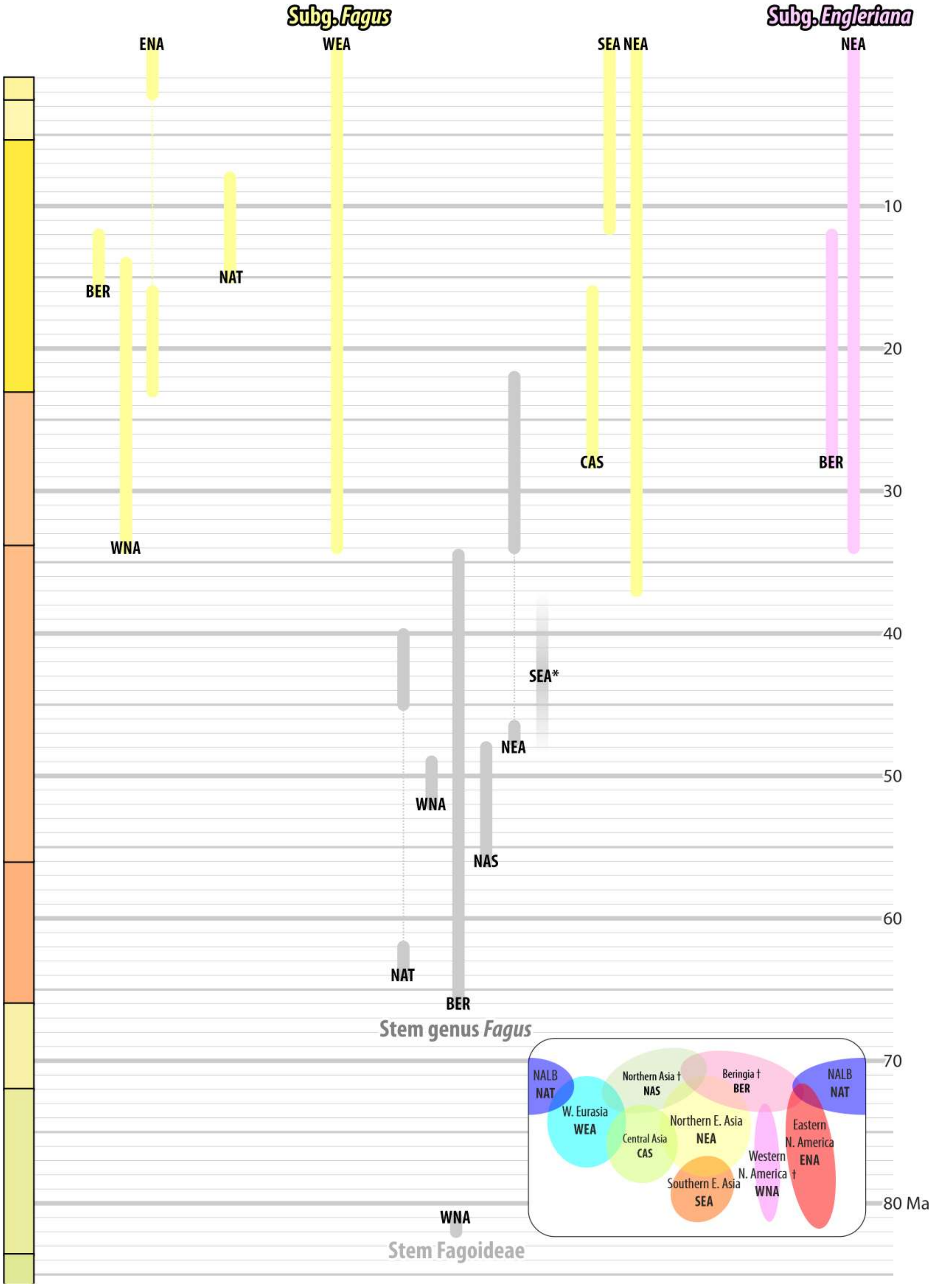


**Figure SR3-10 | *Fagus* over time and space, a summary of the fossil record of beech.** Shown are the stratigraphic ranges of main lineages within *Fagus* per biogeographic region as defined in **Section 4** (abbreviated by three letters, see inlet). * Space- and time-wise isolated pollen from the Eocene of Hainan Is. Additional abbrev.: NALB—North Atlantic land bridge.

##### 3.3.2 Hybrid origin and long isolation of eastern North American beeches

The maternal lineages that gave rise to the modern-day *F. grandifolia* and *F. mexicana* started to diverge in the Paleocene, when the high-latitude eastern (northern North Atlantic, NAT) populations were geographically isolated from their western (northern North Pacific, Beringian, BER) counterparts (**Fig. SR3-11**). This split represents a divergence that preceded the formation of the two modern subgenera in the (late?) Eocene. Based on all available data, the divergence can only have happened in the northern North Pacific region and North America at (very) high latitudes. The Lineage I plastomes, the 'New World' chloroplast genomes, are the legacy of the eastern, northern North Atlantic-Artic branch of the first beeches, beeches thriving since the late Paleocene in subtropical to temperate high-latitude North America (Canadian Arctic + western Greenland).

Phenotypically, the two subgenera are conspicuously linked to the Eocene-Oligocene boundary (Denk & Grimm 2009; **SupplFossilTable.xlsx**). While it can be expected that the subgeneric divergence is (slightly) older than its phenotypic manifestation in the fossil record, it is not as old as the split between Old World and New World plastomes (see also **Section 3.1** for a fossil-record unbiased estimate). As a nucleome-wise and morphologically unambiguous member of the modern *Fagus* subgenus *Fagus,* the first common ancestors (FCA) of *F. grandifolia-mexicana* must have migrated into eastern North America (ENA) *after* the subgeneric divergence, where they then introgressed more ancient (stem) beeches with Lineage I plastomes. A continuous high-latitude beech distribution connecting eastern and western North America (WNA) via Beringia with North-East Asia (NEA) could have maintained trans-continental gene flow between the (high-eastern) North American and Pacific members of the then newly formed *F.* subgenus *Fagus* shaping the distinct *F.* subgenus *Fagus* nucleome signature of the eastern introgrades. *Grandifolia*-like (ENA-type) fossils occur at high latitudes on both sides of the Beringian land bridge at that time part of the "Pacific lineage" of Denk & Grimm (2009) and extend to montane regions in North-East Asia and western North America. On the other hand, the geographic distance coupled with a probably large effective population size of introgressed local mother trees and few invasive *F.* subgenus *Fagus* mother trees ensured that only the 'New World' plastome survived in the *F. grandifolia-mexicana* lineage, while their original (Lineage II?) *F.* subgenus *Fagus* plastomes got lost. If any *F.* subgenus *Fagus* plastome signatures persisted (e.g. in the WNA beeches that can be associated with the *F. grandifolia-mexicana* lineage and *F.* subg. *Fagus*), they got lost during the Miocene and Pleistocene bottlenecks and extinctions.

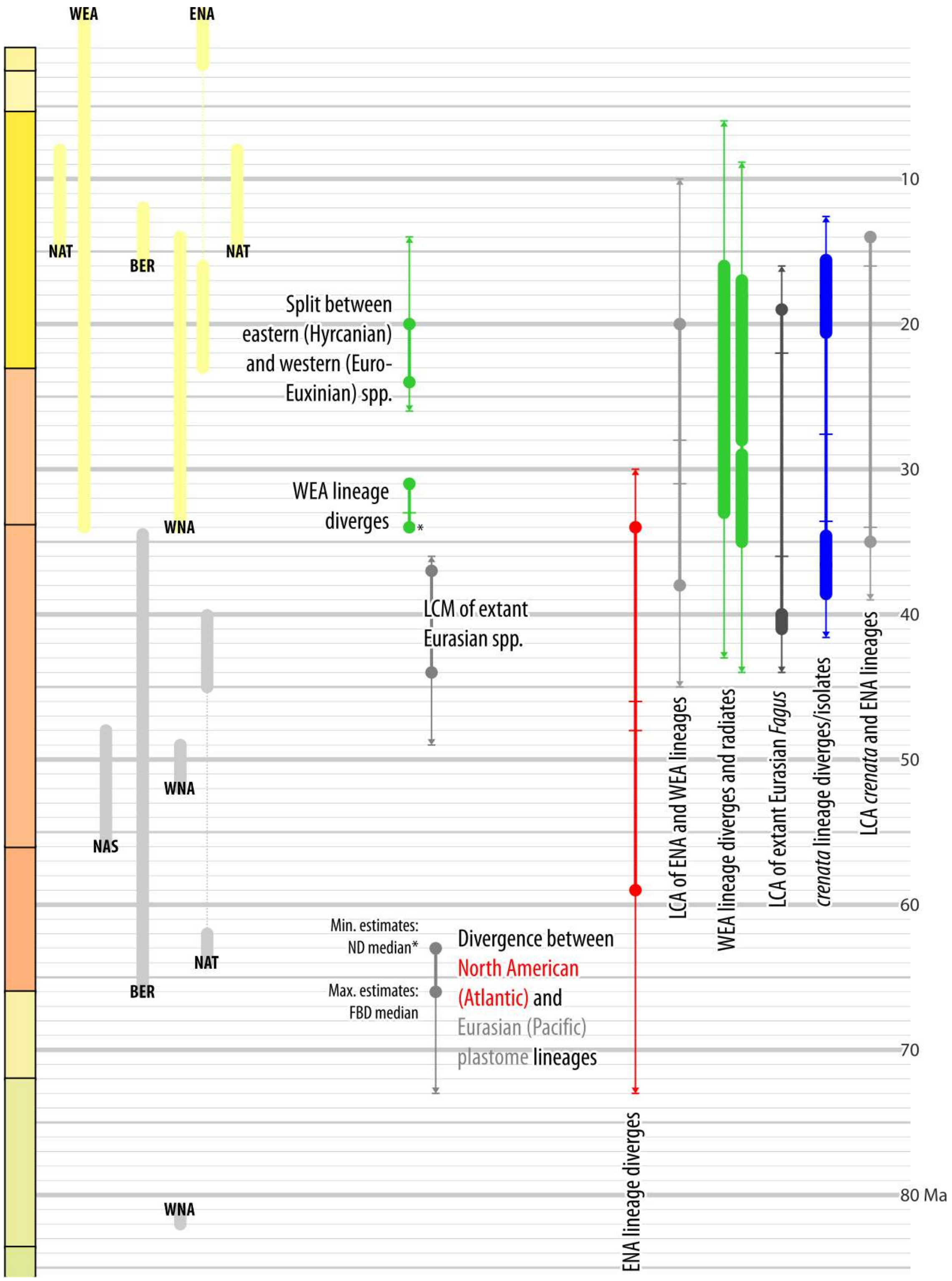


**Figure SR3-11 | Timing of transcontinental disjunctions and gene flow across the North Atlantic land bridge.** Graphed are minimum (using node dating, ND) and maximum estimates (using fossilized birth-death dating, FBD) for selected divergences and nodes, dots and bars give median estimates, whiskers the minimum/-a and maximum/-a of the 95% HPD intervals (in case of nuclear data, from up to three topological scenarios). A summary of the relevant fossil record per biogeographic unit is shown on the left (cf. **Fig. SR3-10; Section 4**). Further abbr.: LCA, LCM = last common ancestor or last common mother, here equalling the most-recent common ancestor of a clade in the chronogram(s) that may include fossil species assigned to the according lineage (see e.g. **Fig. SR3-8**).

The last contact between the eastern North American and West-Eurasian (WEA) beeches or their eastern high-latitude precursors (Atlantic | Pacific topology; see **SupplM&M**, *section 5.2*) – reflected by a sister relationship supported by a number of nuclear loci (CRC, this study; several of Jiang et al.'s 2022 loci, cf. Cardoni et al. 2022) – can be placed in the Oligocene (**Fig. SR3-11**), a time of global cooling. The High-Arctic beech populations that thrived during the Eocene across northern North America and the adjacent land bridges to Eurasia (Beringian land bridge and NALB) got stuck in dead ends (W. North America) or retreated and survived at lower latitudes (Eurasia, E. North America), thus severing any transcontinental gene flow. Nevertheless, possible gene flow across the northern North Atlantic may have continued into the Miocene. A late Early Miocene beech species in Iceland with clear links to North American high latitude beech populations, †*F. friedrichii* Grímsson & Denk was replaced by the younger †*F. gussonii* during the Late Miocene (Grímsson and Denk 2005, Denk et al. 2011, 2013). The latter had a Middle to Late Miocene distribution along the shores of the Mediterranean Sea and the Atlantic including Turkey, Greece, Italy, northern Spain and Iceland (Güner et al. 2017) and may have been a possible vector for gene exchange between North American and Western Eurasian beech populations.

As consequence of the long history of beech in North America, their plastomes as well as some nuclear most-variable gene regions such as the non-coding but transcribed ITS1 and ITS2 are strongly distinct from those of their Eurasian sisters (Denk et al. 2002, 2005), and gene regions supporting an 'Euamerican' clade (Atlantic | Pacific topology) are characterized by relatively distinct sister genotypes (or alleles, see e.g. **Figs SR1-2** and **SR1-3**).

##### 3.3.3 Holophyletic origin and reticulate fate of West-Eurasian beeches

The Western Eurasian beech lineage is the only lineage within genus *Fagus* that has congruent plastid and nuclear differentiation patterns, notably including the estimates from dated trees (see green signatures in **Fig. SR3-11**). Thus, we can assume that, initially, the Western Eurasian beeches had been holophyletic, there was an exclusive West-Eurasian-LCA (last common ancestor) shared by the precursor(s) of all extant Western Eurasian beeches. Based on the fossil record and both nuclear- and plastid-based dating estimates, the West-Eurasian-LCA, represented by the older records of the fossil-species †*F. castaneifolia,* had already been geographically and genetically isolated from its East Asian sisters since the Oligocene. Crown-group radiation within the West-Eurasian lineage must have started simultaneously or shortly after this isolation (**Fig. SR3-11**) when †*F. castaneifolia* expanded westwards north of the Paratethys into the upbuilding Pontic and Hyrcanian mountain chains and those of the Euro-

Mediterranean region; a legacy of these early speciation processes are the markedly drifted plastomes of *F. sylvatica* vs. *F. caspica*. High-resolution genetic analyses indicate that speciation in the Western Eurasian lineage always was more reticulate than dichotomous but generally followed a western-eastern gradient (Cardoni, et al. 2022, Kurz et al. 2023, Denk et al. 2024). The genetic mosaic of modern-day West-Eurasian beeches, combining considerable intra- and interspecific variation while going back to a single (or two) evolutionary sources, is best explained by a reticulate-dynamic speciation model, in which populations get isolated for a certain time (starting allopatric speciation) but never to a degree that they would lose genetic compatibility (allowing for secondary, sympatric hybridisation and introgression) as it is the case of modern-day *F. crenata* (*F.* subg. *Fagus*) and *F. japonica* (*F.* subg. *Englerianae*; Okaura & Harada 2002, Suzuki et al. 2024).

After reaching Europe and colonisation of the proto-Mediterranean region, the then still young West-Eurasian lineage apparently came into contact with a second evolutionary source (see above), the eastern populations of a once trans-Atlantic (or trans-polar?) beech lineage and extinct siblings of the (eastern) North American beeches (nuclear genes supporting the Atlantic | Pacific topology). The age of an Euamerican MRCA (Atlantic | Pacific topology) is estimated to be ~15 myrs younger than the stem age of the North American lineage and ± coeval with the stem age of the Western Eurasian lineage in the New World | Old World topology supported by the majority of nuclear gene regions (**Fig. SR3-11**). The morphologically ambiguous fossil-species †*F. gussonii* found in the Miocene of the proto-Mediterranean region and the Late Miocene of Iceland is a fitting candidate for a late representative of this extinct lineage and a secondary contact. The fact that we do not find plastomes with North American affinity in the beeches of Europe and the Euxinic-Hyrcanian region could be explained by unilateral introgression (pollen-mediated) from the Atlantic lineage (†*F. gussonii*, carrying Lin. I-like plastomes) into the West-Eurasian lineage (coeval fossil-species †*F. haidingeri*, carrying Lin. V plastomes). However, †*F. haidingeri* is a polymorphic species that includes but may not be restricted to the direct precursors of the modern-day species in Europe, Asia Minor (*F. sylvatica-orientalis*) and the Pontic-Hyrcanian region (*F. caspica, F. hohenackeriana*). It includes phenotypes gradually shifting from †*F. castaneifolia* morphotypes into early modern species-shared and late species-restricted morphotypes (see Denk 2004). Part of this morphological heterogeneity could have been due to Atlantic × West-Eurasian hybrids, or introgressed Atlantic-lineage beeches, still carrying 'New World' plastomes, coexisting with the eastern immigrants during the Miocene into the Pliocene. An alternative explanation is hence that the species and populations with North American/Atlantic (Lin. I) plastomes, introgressed descendants of the cross-Atlantic beech lineage, simply lacked the necessary number of mother trees to pass their genetic legacy through the Plio-Pleistocene bottlenecks, being outmatched by the large numbers of *F. sylvatica* mothers carrying the Old World Lineage V(-Sy) plastomes. Recently accumulated data on West-Eurasian beeches has shown that the European *F. sylvatica* is genetically the most homogenous of all West-Eurasian species and only recently diverged from its eastern sister(s) (Gömöry & Paule 2010, Gömöry et al. 2018, Cardoni et al. 2022). All modern-day *F. sylvatica* appear to go back to a small number of mother populations, if not a single one or two (see also **Figs SR2-2, SR2-12**).

An analogy may be found in the oaks; the today exclusively North American red oaks (*Quercus* sect. *Lobata*) were common elements of the pre-LGM (Last Glacial Maximum) European deciduous mixed temperate forests but did not survive the LGM and did not leave any genetic legacy in European oaks. In contrast, the 'Euro-Med' plastid lineage of the westernmost populations of the cross-Mediterranean *Quercus ilex* could be of North American origin, being not part of the Old World core Fagaceae plastome clade (Simeone et al. 2016; placement in Yang et al.'s 2021 cladogram, **Fig. SR3-7**).

##### 3.3.4 East Asian crossroads and melting pot

**An abominable beech mystery**—The Kraskino palaeoflora (dated to 34 Ma, Eocene-Oligocene boundary) of the Russian Far East harbours not only the earliest unambiguous fossil species that can be assigned to either *Fagus* subgenus *Fagus* or *F.* subgenus *Englerianae.* The two principal phenotypes of East Asian species of *F.* subgenus *Fagus* are present as well: fossils with striking resemblance to modern-day *F. crenata* of Japan or *F. longipetiolata* of central China. This sudden peak of distinctly modern and co-occurring phenotypes at the Eocene-Oligocene boundary can only be the product of earlier speciation processes and a first sorting of species lineages. The first common ancestors of both subgenera must have diverged earlier. The plastome Lineage II found in north-bound Japanese beech populations of *F. crenata* (*F.* subg. *Fagus*) and *F. japonica* (*F.* subg. *Englerianae*) is notably different from the Lineage IV and Lineage V plastomes of the continental Eurasian species and southern or western-most populations of *F. crenata* and *F. japonica*. The FBD-estimated divergence age, treating Lineage II plastomes as a legacy of a high-latitude beech lineage represented by fossils ranging from 36–16 Ma, is ~44 Ma, i.e. younger than the FBD-estimated subgeneric split but older than the phenotypical manifestation of the modern subgenera (used as age prior for ND of the Eurasian plastid MRCA). The oldest pollen records in this area go back to the early Paleocene (66–62 Ma), the oldest fossil assigned to represent Lineage II is †*F. napanensis* Fotjanova from the latest Eocene (Priabonian) of Kamchatka, a fossil that may represent a very early, phenotypically still ± primitive member *Fagus* subgenus *Englerianae* (not assigned to either subgenus in the nuclear dating experiments). Further evidence for a deep subgeneric split comes from the ITS region, with all *F.* subgenus *Englerianae* species having sequentially much different ITS variants in contrast to the Eurasian species of *F.* subgenus *Fagus* (Grimm et al. 2007). Likewise, the CRC and LFYi2 sequences of both subgenera a notably distinct (**Section 1**; see also Renner et al. 2016), and in any of the low-divergent nuclear marker compiled by Jiang et al. (2022) with sufficient diversity, the subgenera are clearly sorted, the number of SNPs and mutations patterns being up to ~5-times higher than the maximum observed within *F.* subgenus *Fagus* (between N. American and E. Asian spp.) It is hence probable that the split between highest-latitude Eurasian Lineage II and lower latitude Lineage III–V plastomes coincided with the subsequent geographic and genetic isolation of the FCAs of the modern-day subgenera, and that the Lineage II plastomes were carried by the high-latitude Subgenus-*Englerianae*-LCA, while Lineage IV plastomes come from a lower latitude East Asian-subgenus-*Fagus*-LCA. The alternative is that a polyploidisation event rendered the early members of both subgeneric lineages breeding-wise incompatible. Both the ITS regions (Denk et al. 2005) as well as the dominant 5S-IGS arrays (Cardoni et al. 2022; see also **Fig. SR1-10**) are much more divergent in *F.* subgenus *Englerianae* than in species of *F.* subgenus *Fagus*, which could reflect an early auto- or allopolyploidisation or ancient hybridisation via secondary contact: the *S*ubgenus-*Englerianae*-LCA may have been plastid-wise heteromorphic from its start.[21]

Either way, the fact that the plastomes are not sorted in East Asian beeches along the boundaries of the subgenera can only be explained by a hybrid origin (reticulate evolution): the early members of *F.* subgenus *Englerianae* or both subgenera must have come into contact and introgressed (extinct) lineages of beeches that already had started to diversify in the northwestern Pacific region. Hybridisation and widespread introgression between the two modern subgeneric lineages can be largely ruled out because (*i*) their phenotypes remain stable and distinct from the Eocene-Oligocene boundary onwards despite being found close-by or even in the same fossil lagerstätte (cf. **SupplFossilTable.xlsx**, sheet *main list*) and (*ii*) the lack of introgressed or exchanged, sequentially evolved nuclear alleles and variants of most-divergent multi-copy nuclear gene region such as the ITS region (Denk et al. 2005) and the

[21] In other sampled, low- or single-copy nuclear markers (*CRC*, LFYi2, markers studied by Jiang et al. 2022) such a signal could have been largely lost and overprinted by a recent radiation of the Modern-Beech-LCA.

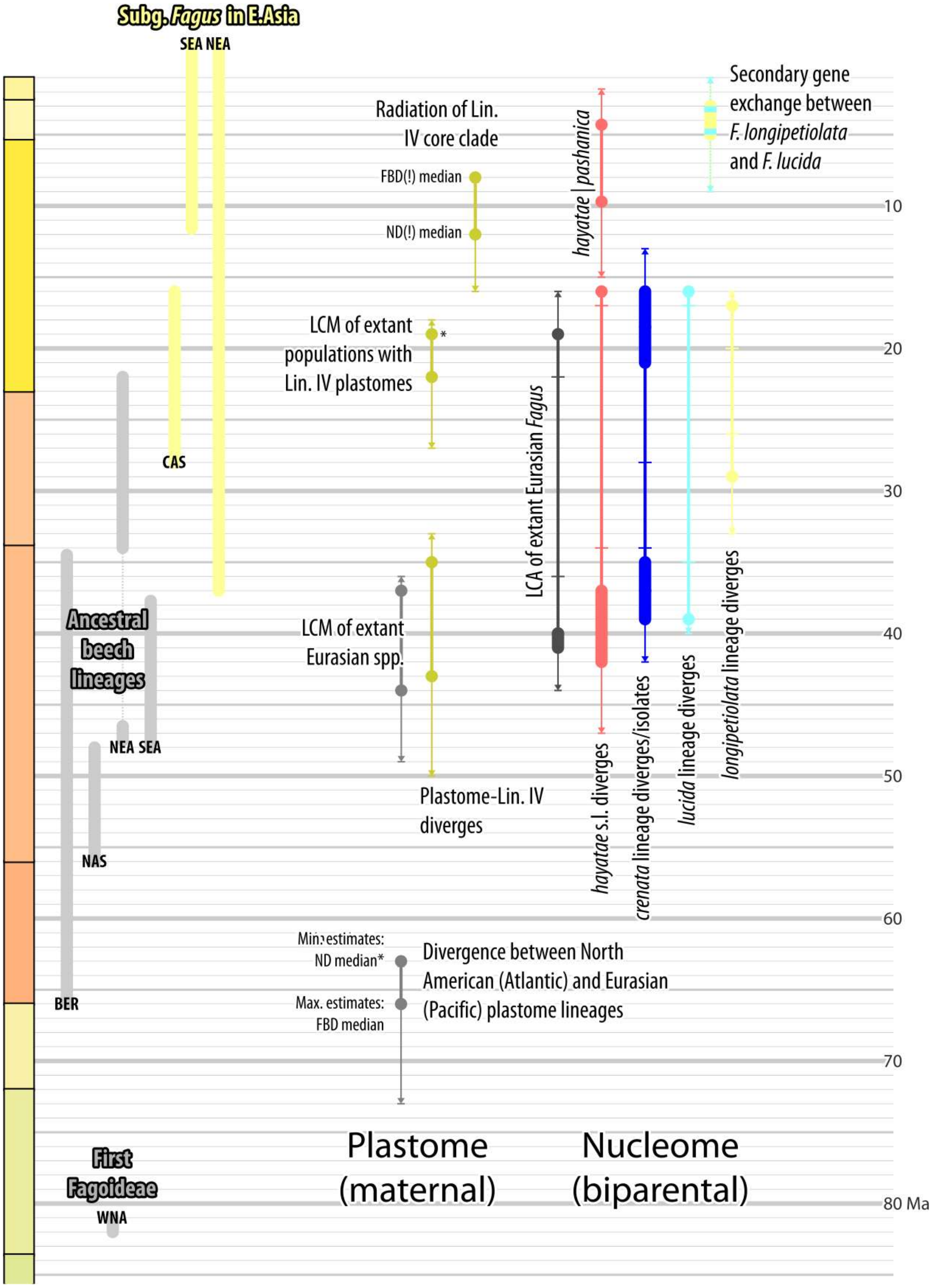


**Figure SR3-12 | Timing of radiations within the East Asian members of *Fagus* subgenus *Fagus*.** Graphed are minimum (using node dating, ND) and maximum estimates (using fossilized birth-death dating, FBD) for selected divergences and nodes, dots and bars give median estimates, whiskers the minimum/-a and maximum/-a of the 95% HPD intervals (in case of nuclear data, from up to three topological scenarios). A summary of the relevant fossil record per biogeographic unit is shown on the left (cf. **Fig. SR3-10; Section 4**).

intergenic spacers of the 5S rDNA arrays (Cardoni et al. 2022). Even when growing in mixed stands, there is no evidence for genetic exchange between species of either subgenus (Okaura & Harada 2002, Suzuki et al. 2024). While sym- and parapatric *F. crenata* and *F. japonica* share sister plastomes within a lineage or the same SLPT (species-level plastid type; **Section 2.2**), they always differ consistently by at least a few SNPs and length-polymorphic patterns.

The primary and secondary radiation in Lineage IV plastomes (**Fig. SR3-12**) may be linked to early diversification of beeches predating the subgeneric split or primordial members of *Fagus* subgenus *Fagus*, which were then introgressed by early members of *F.* subgenus *Englerianae*, the latter providing the parental lineage(s) of the modern-day *F. japonica*. The notably young age for the MRCA of the modern-day species of *F.* subgenus *Englerianae* and the geographic differentiation in their plastomes can be explained by a species bootleneck and re-radiation of the Modern-Beech-LCA, the last common ancestor of all extant beeches. Early members of *F.* subgenus *Englerianae* carrying different plastomes went extinct by being introgressed by (a) (Late) Miocene species, the precursor(s) of modern-day *F. engleriana, F. japonica* and *F. multinervis.*

The latest trans-subgeneric introgressive episode, which our estimates place in the latest Miocene, lead to the formation of *F. engleriana*. In contrast to its north-eastern siblings, *F. engleriana* only carries plastomes of the Lineage IV core clade (SLPTs IV-EnA, -EnB and -ChB), a plastid lineage shared by all continental southern East Asian species. At this time, all extant species lineages of *Fagus* subgenus *Fagus* had been diverged and isolated from each other, even when we consider only the youngest estimates among our dating experiments. As in the case of *F. japonica*, there is no evidence for relatively recent or ongoing gene flow between *F. engleriana* and sympatric species of *F.* subgenus *Fagus*; thus, all Lineage IV core clade SLPTs must have come from extinct sisters or cousins of the modern-day Chinese species.

The mixing with different evolutionary sources (Lineage II-donor[s]; Lineage IV-donor[s]) linked with hybrid vigour (McKown & Guy 2018) would serve as an explanation for the co-occurrence and subsequent rapid spread of *crenata*- and *longipetiolata*-like leaves shortly after the two modern subgeneric lineages formed and manifested themselves. Our dating of the nuclear and plastid genealogies place the onset and primary radiations of *Fagus* subgenus *Fagus* in East Asia in the Oligocene (oldest FBD estimates late Eocene, < 42 Ma), with the final divergences, the formation of the modern-day species lineages, as late as the mid-Miocene (youngest ND estimates >15 Ma; **Fig. SR3-12**). While decoupled, and likely reflecting at least partly different speciation events, the primary and following main radiations seem to have happened within the same time frame in both the nucleomes and plastomes, and can be largely linked to the Oligocene cooling (**Table SR3-4; Fig. SR3-12**). Last contacts and final speciation events can be as late as the latest Miocene (> 5 Ma). The decoupling within the East Asian species of *Fagus* subgenus *Fagus* oscillates between two extremes: on one side, the Japanese *F. crenata* carrying both Lineage II and mid- to late radiating Lineage IV plastomes, evolving in parallel; on the other the Chinese-Taiwanese species, carrying an early diverged Lineage IV plastome in their south-easternmost, oceanic populations (Taiwan, south-eastern China?) and most-recently diverged Lineage IV core clade plastomes in their continental range. The Lineage IV core clade SLPTs are sister types of SLPTs found in the only Chinese species of *F.* subgenus *Englerianae*, *F. engleriana*, and south-westernmost *F. crenata.*

**Table SR3-4 | Median FBD (Kraskino fossils treated as crown members, maxima) and ND estimates (minima, rounded to full myrs) for major divergences within East Asian beeches of *Fagus* subgenus *Fagus*.** MRCA = most-recent common ancestor, i.e. the head node linked the subtree including all tips of the respective species; YSA = youngest shared ancestor, the youngest node connecting representatives of the respective species in the tree, i.e. the last inferred contact between members of two or more species.

| Node | Plastid tree | Nuclear trees | | |
|---|---|---|---|---|
| | | Atl \| Pac[a] | High \| Low[b] | NW \| OW[c] |
| MRCA of extant E. Asian spp. | 66 /63 | 40 /19 | 52 /31 | 41 /27 |
| YSA of extant E. Asian spp.[d] | 18 /22 | = MRCA | = MRCA | = MRCA |
| *F. crenata-F. longipetiolata* YSA/MRCA[e] | 8 /11 | 35 /19 | 52 /31 | 41 /27 |
| MRCA of SEA spp.[f] | 18 /22 | 40 /19 | 42 /23 | 37 /24 |
| *F. longipetiolata-F. lucida* YSA/MRCA[e] | 5 /6 | 37 /17 | 36 /5 | 35 /9 |
| MRCA of *F. hayatae-pashanica* | 18 /22 | 10 /5 | 30 /16 | 12 /23 |

[a] In the Atlantic | Pacific topology, the *F. hayatae-pashanica* lineage is sister to *F. lucida* + *F. crenata-longi-petiolata*.
[b] In the High | Low topology, the *F. crenata* lineage is part of the 'high-latitude clade' (together with the W. Eurasian and N. American spp.), while the other E. Asian species form a 'low-latitude'/'Southern clade'.
[c] In the New World | Old World topology, the *F. crenata* lineage represents the first diverging species lineage within an E. Asian clade, followed by *F. hayatae, F. pashanica* (forming a clade, FBD-chronogram, or grade, ND-chronogram), and *F. longipetiolata* + *F. lucida.*
[d] In case of the plastomes, close to the MRCA of Lin. IV plastomes: MRCA of SLPTs IV-Ha + IV-LuA and Lin. IV core clade.
[e] In the nuclear trees, the YSA of two species equals their MRCA, as each species is represented by a single tip.
[f] Species lineages today restricted to southern E. Asia: *F. hayatae-pashanica, F. longipetiolata, F. lucida,* all of which have fossil records in N. E. Asia and, in case of the *F. lucida* lineage, the Miocene of Central Asia (†*F. altaensis*)

**The many mothers of *F. crenata***—The many and distinct plastomes carried by modern-day *F. crenata* can only come from beeches that were at some point introgressed and taken over by the direct ancestor(s) of *F. crenata*. Both the pre-Oligocene *Crenata*-FCA and Late Miocene or younger *Crenata*-LCA, the first *and* last common ancestors of today's *F. crenata* (**Fig. SR3-12, Tables SR3-4, SR3-5**), may have been holophyletic but the modern-day populations do not share the same primordial mothers. Since there is so far no evidence for post-divergence gene flow between both subgeneric lineages and *F. crenata* and *F. japonica* do not interbreed even in close contact (Okaura & Harada 2002, Suzuki et al. 2024; see also **Section 2.2.2**), it is unlikely that *F. crenata* picked up any plastome directly from sym- to parapatric *F. japonica*: shared plastome lineages and SLPTs (II-JaB) can only represent ancient introgression (shortly after the subgenera evolved and came into contact) or, more probably all data considered, parallel but temporally different introgression of the same extinct beech lineages; primordial beeches that were genetically compatible with both the subgenus-*Fagus*-FCA and the subgenus-*Englerianae*-FCA and/or the precursors of modern-day *F. crenata* and *F. japonica* being from the same lineage than one of their parental genetic donors.

Morphologically and genetically (all nuclear markers considered), *F. crenata* is the closest modern relative of the West-Eurasian species (Grimm et al. 2007, Renner et al. 2016, data produced by Jiang et al. 2022; see also main-text fig. 3). *Fagus crenata* is also the only species in East Asia that covers the entire bioclimatic niche of the West-Eurasian species (May-cock 1994, Peters 1997, Grimm & Denk 2012). Using the High- | Low-latitude topology, we estimated that gene flow between the precursors of *F. crenata* (and the New World beeches) and their western relatives broke down latest by the Burdigalian (Early Miocene, ~17 Ma). In contrast, none of the many SLPTs detected in *F. crenata* represents a close or closest relative of the West-Eurasian Lineage V plastomes. Lineage II and one Lineage IV SLPTs are sister types of sym- to parapatric *F. japonica*; and the remaining Lineage IV SLPTs sister to or part of the Lineage IV core clade, which characterise all continental East Asian beeches. The last shared mother of (modern-day) *F. crenata* and the West-Eurasian beeches, the High-latitude-clade-LCM, is at least 8 myrs older than the putative High-Latitude-LCA (late Eocene, median 44 Ma vs. 38 Ma using FBD; earliest Oligocene, median 35 Ma vs. 17 Ma using ND; **Fig. SR3-12, Table SR3-5**). With respect to the phylogeographic history of West-Eurasian beeches, shared genetic characteristics could have been maintained longer via intermediate Central and North Asian beech populations. A ± continuous distribution area and stable large effective population sizes would have slowed down genetic drift between the western Paratethyan and eastern Pacific members of a cross-Eurasian (cool-)temperate beech lineage, which would explain *(i)* why some nuclear gene regions, such as the ITS regions of the nuclear-encoded 35S rDNA cistrons, are still very similar in Western Eurasian beeches and their East Asian sisters and (*ii*) the limited genetic drift observed in far the most nuclear gene regions of *F. crenata*. On this background, both the FBD- and ND-estimates from the High- | Low-latitude topology make sense.

*Fagus crenata* is also much more similar to the other East Asian species of *Fagus* subgenus *Fagus* than are the West-Eurasian species. Nonetheless, the other two nuclear topologies, both with an East Asian-subgenus-*Fagus* clade lead to similar divergence estimates when we focus on the (MRCA-based) stem age of the *F. crenata* lineage (**Table SR3-2**; see also **Fig. SR3-12**). As a trend, the stem age estimates (maximum FBD, minimum ND: **Fig. SR3-12**) for the *F. crenata* lineage fall in the same time frame as the primary (fast) radiation within Lineage IV plastomes; a radiation which does not involve *F. crenata* SLPTs at all. Rather than being a sister species of any of the Chinese-Taiwanese species of *F.* subgenus *Fagus*, as implied by the certain nuclear trees and the East Asian-subgenus-*Fagus* clade, *F. crenata* may be a distant (high-latitude, cool-temperate) cousin that only secondarily came into contact with and introgressed their actual Japanese sisters/oceanic counterparts. Their genetic signatures imprint the signal for the East Asian-subgenus-*Fagus* clade favoured by some of the nuclear markers of Jiang et al. (2022), supported by relatively few conserved mutations, and rejected by others (cf. **SupplGenetics_ncDNA.xlsx**, sheet *SummaryJiangEtAl*, see also Cardoni et al. 2022, S5).

Notably, even the youngest FBD-estimates for the MRCA of *F. crenata* and the rest of the East Asian species of *Fagus* obtained with the Atlantic | Pacific topology (treating the modern-type Kraskino fossils as members of *Fagus* subgenus *Fagus* but not as members of the modern-day species lineages) are older than the age of the MRCA of Lineage IV plastomes which are carried by all other East Asian species of *F.* subgenus *Fagus* (26 vs. 19 Ma). The youngest divergence between *F. crenata* plastomes and sister plastomes carried by *F. japonica* or other East Asian species may have been as late as the mid- to Late Miocene (~ 10 Ma), i.e. reflect (allopatric) speciation (or intra-species geographic sorting) ≥ 15 myrs after the FCA of *F. crenata* diverged from the remainder of *F.* subgenus *Fagus*. The precursor(s) of *F. crenata* must have introgressed several local beech species when they reached Japan and established their modern-day distribution area. Based on the current data, it is impossible to explicitly date the takeover but it may be very recent. When not including †*F.* cf. *palaeocrenata* from the Kraskino palaeoflora in the *F. crenata* lineage, we obtained a Late Miocene *modelled* stem age of 7 (11–5) Ma for the *F. crenata* lineage within the East Asian-subgenus-*Fagus* clade of the Atlantic | Pacific topology (9 Ma in the New World | Old World topology), which reflects the fossil hiatus of *crenata*-like leaf fossils in- and outside Japan.[22] Probably not coincidental, Late

[22] The difference between modelled and MRCA-based stem ages is due to shadow lineages modelled in course of the FBD optimisation to accommodate the surplus of fossils assigned to (the next) larger clade(s) and subsequent birth-death events. Since no morphological partition was used, these shadow lineages are pure model-constructs and accordingly modelled divergence ages must be viewed as such. The young *modelled* stem age of the *F. crenata* lineage relates to the age distribution of fossils assigned to this lineage, which will be embedded in a larger clade

Miocene Japanese palaeofloras also include the last *F.* subgenus *Fagus* leaf morphologies that lack a clear affinity to any modern-day species lineages (†*F. protolongipetiolata* A, ranging from Eocene-Oligocene boundary till the Late Miocene of the Sea of Japan and Taiwan) or show a high similarity to *F. hayatae-pashanica* (†*F. florinii* Huzioka & Takahashi*,* †*F. stux-bergii* [A] (Nathorst) Tanai; the latter found until the Zanclean [Pliocene] in Japan). Finally, all *F. crenata* Lineage IV SLPTs were more recently diverged (FBD ≥ 10 myrs, ND ≥ 8 myrs) than the primary radiation reflected in Lineage IV SLPTs found in *F. japonica*; putting the introgressive event in the mid-Miocene or later. This also means that the (putatively holophyletic) *Crenata*-LCA is much younger than the MRCA-based stem age estimated for the *F. crenata* species lineage but in line with the young modelled stem ages, when treating the Kraskino fossils not as part of the modern species lineages (**Tables SR3-2**, **SR3-3**). The modern-day *F. crenata* appears to be only the Plio-Pleistocene reincarnation of an old lineage of (cool-)temperate Eurasian-Beringian beech species that originally thrived at the high latitudes.

A ± simultaneous or concerted invasive and introgressive phase represents the simplest possible explanation not only for the complex plastid diversity seen in *F. crenata* but also the partly contradicting aspects of its nuclear differentiation patterns. Even though the sample was limited, Cardoni et al. (2022) detected three main types of 5S-IGS variants, one being part of lineage shared with the West Eurasian species (Crenata B1, part of the 'Original B' lineage) but the dominant ones (Crenata B2 and Crenata B3) being exclusive to *F. crenata* and clearly distinct from the (co-)dominant West-Eurasian types. Meanwhile, first data has been obtained from the Taiwanese *F. hayatae* and its cryptic Chinese sister species *F. pashanica.* Also in this case, the main 5S-IGS types appear to be specific (we found no exclusively shared 5S-IGS variants between our two samples, one for *F. hayatae,* one for *F. pashanica*); three of the four new *F. hayatae* and *F. pashanica* types are part of the 'Original B' lineage (**Fig. SR1-10**) and the forth is more similar to it than the Crenata B2 and B3 types, which could indicate that they either come from a sister lineage of the modern-day *Fagus* subgenus *Fagus* or from a genetically much drifted or early diverged species lineage within *F.* subgenus *Fagus.* We furthermore found a CRC polymorphism in *F. crenata* and dimorphism in *F. longipetiolata* (**Fig. SR1-3**), a link that is corroborated by some of the nuclear markers sequenced by Jiang et al. (2022) and expressed in the Atlantic | Pacific topology (**SupplM&M**, fig. SM5-3): such a

comprising fossils assigned to the East Asian clade of subgenus *Fagus*, most of which are older than the *F. crenata* lineage-fossils. The MRCA of the East Asian subgenus *Fagus* clade, defines the MRCA-based stem age of the *F. crenata* lineage.

pattern may be the result of secondary gene exchange between their lineages or mixing of shared parental lineages. The observation that *F. crenata* constitutes a possible sister species not only of one but several other modern-day species (**Fig. SR1-1)** could be, in analogy to its plastid diversity, a legacy from the various, not directly related, native Japanese species that were simultaneously introgressed by the precursor of modern-day *F. crenata*. The existence of Japanese sister species (or regional subspecies) of the modern-day Chinese and Taiwense species and their precursors would follow a common phytogeographic pattern observed in many subtropical to temperate tree genera stretching from Central China and Taiwan to Japan (Latham & Ricklefs 1993).

Provided better sampled phylogenomic data and data from population-level studies covering all *crenata*-SLPTs, it may become possible to discern the probable number and even the phylogenetic affinities of the many mothers of *F. crenata.* Just based on the plastomes and our dating experiments, at least five or six different species (lineages) where introgressed by the precursor(s) of *F. crenata* (**Table SR3-5**) including the plastid sister clade of the clade comprising plastomes of all continental 'southern' East Asian (SEA) species (*F. longipetiolata, F. lucida*, *F. pashanica*).

**Table SR3-5 | Species-level plastid types (SLPT) of *F. crenata* and their phylogenetic affinity.** Stem and crown ages give median FBD- and ND-estimates for respective MRCAs. NA = not applicable (single tip).

| SLPT | Affinity | Stem age | Crown age | Evolutionary hypothesis |
|---|---|---|---|---|
| II-CrA | 2nd diverging branch | 22 /18 | NA | Ancient high-latitude (Beringian) lineage |
| II-CrB[b] | Final radiation | 10 /8 | 5 /8 | Ancient high-latitude (Beringian) lineage |
| II-JaB[b] | Recent introgression | 10 /8 | 1 /1 | Ancient high-latitude (Beringian) lineage |
| IV-CrC+JaF | Early *F.* subg. *Fagus* | 15 /18 | 10 /11 | Native subgenus *Fagus* species |
| IV-CrD | Lin. IV core clade | 9 /12 | NA | Miocene radiation within subgenus *Fagus* |
| IV-CrE+EnA | Sister of SEA spp. | 8 /11 | 6 /9 | Extinct ± subtropical species[c] |

[a] The SLPTs II-CrB and -JaB could represent as well a geographic differentiation within a single, once widespread and dominant species (cf. **Fig. SR2-6**).
[b] IV-CrE plastomes are exclusively found in the mountains of Shikoku and Kyushu Islands (among warmest stands of *F. crenata* across Japan); in central China, *F. engleriana* can occur from 200–300 m a.s.l. (hot subtropical *Cfa* climate) to mid-altitudes, i.e. below and as part of the beech forest belt (cf. supplement to Grimm & Denk 2012).

**Episodic migration into lower latitudes**—Based on today's distribution and the molecular differentiation pattern as well as the fossil record (**SupplFossilTable.xlsx, → Section 4**; see also Grímsson et al. 2016), it is reasonable to assume that beech originated and radiated at (very) high latitudes during the global greenhouse climate of the Paleocene-Eocene (Zachos et al. 2001; Scotese et al. 2021) and then migrated to lower latitudes in course of the late Eocene-Oligocene cooling. The beeches that produced the middle Eocene (47–37 Ma) Hainan pollen (~19° N palaeolatitude) left no traceable imprint in the plastome of their modern-day relatives.

The first tangible legacy of a southward expansion of modern-day beeches are the Lineage IV plastomes carried by *F. hayatae*, endemic to the northern mountains of Taiwan (SLPT IV-Ha) and a putatively south-eastern Chinese individual of *F. lucida* (SLPT IV-LuA). A plastid divergence placed in the Early Miocene by both FBD- and ND-estimates and in line with a similar ND-estimate for the crown age of an East Asian-subgenus-*Fagus* clade. At the time a suitable mountainous area stretched between 25–29° N along the Pacific coastline and hinterland (Scotese 2014b). The divergence of this southern East Asian plastid lineage is ≥ 8 myrs younger than FBD-estimates for the stem age of the *F. hayatae-pashanica* lineage (**Table SR3-6**), a modern species lineage which may include as oldest representatives a fossil from the Kraskino flora (†*F. koraica* Huzioka; triggering Eocene stem ages for this and other E. Asian species lineages; see above: **Section 3.3.1**, **Table SR3-3**) as well as †*F. florinii* from the Oligocene-Miocene boundary of Jilin (N.E. China; **SupplFossilTable.xlsx**). A Late Miocene fossil from Taiwan (†*F. protolongipetiolata* A) lacks, however, a distinct *hayatae-pashanica* morphology. In contrast, the only *F. pashanica* plastome sequenced so far (one of two plastomes of SLPT IV-ChA; the other reported from another *F. lucida* individual), is part of the Lineage IV core clade.

The Lineage IV core clade diverged and radiated > 10 myrs later (in or by the Late Miocene), placing the origin of the SLPT IV-ChA and the only reported *F. pashanica* plastome in about the same time frame as the inferred MRCA of *F. hayatae* and *F. pashanica* (minimum 5 Ma, maximum 10 Ma; **Table SR3-6**). At the time the island populations (*F. hayatae*) got completely isolated from the continental populations (*F. pashanica*), their maternal lineages had been long diverged and sorted. Two migratory scenarios can explain our results. Either the *Hayatae-Pashanica*-LCA, the last common ancestor of *F. hayatae* and *F. pashanica*, migrated from its originally coastal distribution area (IV-Ha plastomes; fossils with *hayatae-pashanica* phenotypes are known from Japan from the Middle Miocene till the Pleistocene, and the Pliocene of Jiangxi province, China) into the mountains of central China, where it introgressed other beech species of *Fagus* subgenus *Fagus* (picking up the IV-ChA SLPT; forming *F. pashanica*). Or vice versa: > 10 myrs after beeches established in Taiwan and adjacent areas, the *Hayatae-Pashanica*-LCA migrated south-east into this area, where it introgressed the IV-Ha plastome carrying local species to form the modern-day *F. hayatae*. Current available data on nuclear differentiation point to a relatively early migration and isolation of the Taiwanese beeches from their continental Chinese counterparts (precursors of *F. longipetiolata* and *F. lucida*). *Fagus hayatae* distinguishes itself by uniquely evolved alleles (details provided in Cardoni et al. 2022, data S5) and two, drifted from each other, 5S-IGS main types while *F. pashanica* shows a higher proportion of shared alleles, ancestral within Eurasian species of *F.* subgenus *Fagus,* and two similar, less drifted 5S-IGS main types (**Figs SR1-9, SR1-10**). Accordingly, the age estimation for the MRCA of *F. hayatae-pashanica* differs strongly between topologies and according gene samples (**Table SR3-6**). *Fagus pashanica* could be the product of (Miocene) mixing and secondary and/or longer persisting gene flow between more continental-montane and oceanic low(er) latitude beech lineages of *F.* subgenus *Fagus*. Further support for such a 'unequal sisters' hypothesis, early/strongly isolated *F. hayatae* vs. once-promiscuous *F. pashanica*, comes from a recent study by Li et al (2023). Despite relying on suboptimal nuclear markers, their data and comprehensive sampling indicates a generally low genetic coherence of *F. pashanica* across its contemporary range while confirming its genetic isolation from *F. hayatae*. Recent mixing and ongoing gene flow can be so far ruled out, since *F. longipetiolata, F. lucida* and *F. pashanica* occasionally form mixed stands, but no intermediate phenotypes or conspicuously mixed genotypes have been reported so far.

**Table SR3-6 | Median FBD (as maxima) and ND estimates (as minima, in Ma) for the stem and crown age of *F. hayatae* (*ha*)-*pashanica* (*pa*) lineage and associated species-level plastid types** (*F. hayatae* SLPT IV-Ha, *F. pashanica* shared SLPT IV-ChA). Number in brackets give estimates treating the modern-type Kraskino fossils not as part of the respective species lineage but as member of the entire *Fagus* subgenus *Fagus* clade (†*F. koraica,* in case of the *F. hayatae-pashanica* lineage). Red numbers: MRCAs (reconstructed most-recent common ancestors) do not represent exclusive LCAs (last common ancestors).

| **Node/aspect** | **Topology** | | | |
|---|---|---|---|---|
| | Atl \| Pac | High \| Low | NW \| OW | Plastid |
| MRCA of E. Asian subg. *Fagus* spp. | 40 (38) /19 | 52 (53) /23 | 41 (37) /21 | 44 /≥ 36[a] |
| Stem age of *ha-pa* lineage | 40 (38) /16 | 42 (–) /–[b] | 37 (25) /– | – /– |
| Modelled crown age of *ha-pa* lineage[c] | 36 (24) /– | 37 (–) /– | 34 (25) /– | – /– |
| MRCA of *ha-pa* | 10 (9) /5 | 29 (45) /15 | 12 (11) /13 | 18 /22 |

[a] Constrained by age prior
[b] If no age estimate are given, the according node/aspect does not apply for the respective dated tree.
[c] Modelled age refers to the foot node of the root of the *F. hayatae-F. pashanica* clade including fossils as defined in the according TaxSet in the xml-file (see **SupplGenetics.xlsx**, sheet *ncFBD*).

So far, the plastomes of three individuals of *F. lucida* have been sequenced and revealed one divergent and two similar Lineage IV SLPTs: IV-LuA, sister plastome of *F. hayatae* plastomes vs. IV-ChA, shared with (some?) *F. pashanica,* and IV-ChB, shared with *F. longipetiolata* and an individual of *F. engleriana* (of *F.* subg. *Fagus*). Thus, dating-wise and plastome-wise *F. lucida* is analogous to *F. hayatae-pashanica*. With respect to the geographic history of the *F. lucida* lineage, characterised by a unique leaf phenotype, it may well be that the precursors of *F. lucida* introgressed the ancestors of *F. hayatae-pashanica,* when they migrated into their current distributional area. Their south-easternmost populations introgressed the continental counterpart of what would become *F. hayatae*,[23] and the remainder early or primordial *F. pashanica* carrying a Lineage IV core clade plastome coming from a different mother population (as discussed above). The *F. lucida* species lineage can be traced via †*F. altaensis* from the Middle Miocene of central Asian Altai Mountains to the Kraskino palaeoflora from the Eocene-Oligocene boundary of the Russian Far East (†*F.* 'cf. *evenensis*', †*F. olejnikovii* Pavlyutkin), but has not been found in the Japanese archipelago in contrast to *longipetiolata* and *hayatae-pashanica* phenotypes (**SupplFossilTable.xlsx**). It may represent an early continental lineage within *Fagus* subgenus *Fagus* that originally thrived outside the range and niche of the precursors and extinct sisters of *F. longipetiolata* or *F. hayatae-pashanica,* the putative donors of Lineage IV plastomes in *F. crenata* and *F. japonica*. If such a continental lineage carried accordingly drifted plastomes, which should be closer to Lineage III and V plastomes than Lineage IV plastomes, they have been lost when the precursor of the modern-day *F. lucida* migrated into their current range in Central China, where they have become sym- or parapatric to *F. longipetiolata* and *F. pashanica*. Extensive mixing with (an)other, genetically already isolated, *F.* subgenus *Fagus* lineage(s) reflects its migratory history: Jiang et al. (2022) encountered increased levels of heterozygosity linked to (strongly) evolved but divergent alleles shared between *F. lucida* and *F. longipetiolata* in addition to alleles and nuclear gene regions rejecting any sister relationship between the two species. The modern-day *F. lucida* is clearly of hybrid origin and underwent at least two major phases of reticulate evolution, with some of its evolutionary roots well going back to the time of the first radiations of *F.* subgenus *Fagus* close to the Eocene-Oligocene boundary (~34 Ma) as well as inter-species gene flow in course of secondary contact as late as the latest Miocene to Pliocene (~ 5 Ma; **Table SR3-7**). It can be expected that more data on the species, a better sampling across its entire range, will reveal more aspects of its complex history and also the timing of the final migration into its modern area.

As in the case of *F. hayatae-pashanica* and *F. lucida* lineage, also the *F. longipetiolata* lineage originated in North-East Asia during the Late Eocene-Oligocene transition and underwent possibly several phases of reticulation. One parental lineage of the modern-day *F. longipetiolata* had been diverged latest by the Early Miocene (≥ 17 Ma; **Table SR3-7**; **Fig. SR3-12**). The genetic legacy of this divergence are its highly specific and evolved nuclear alleles (**Fig. SR1-9**; **OSA**, sheet *00_bits.xlsx*; see also Cardoni et al. 2022, data S5) as well as the ITS dimorphism

[23] The current plastid data on *F. pashanica* is too limited to judge the possibility that IV-LuA SLPT is in fact a SLPT shared by south-eastern *F. lucida* and

shared with *F. pashanica* (Denk et al. 2005; Grimm et al. 2007), and, possibly, also the *CRC* dimorphism that links *F. longipetiolata* to *F. crenata* (this study).

**Table SR3-7 | Median FBD and ND estimates for the stem and crown age of *F. longipetiolata* (*lo*) and *F. lucida* (*lu*), and the plastid Lineage IV core clade.** Red numbers: MRCAs (reconstructed most-recent common ancestors) do not represent exclusive LCAs (last common ancestors).

| **Node/aspect** | **Topology** Atl \| Med | High \| Low | NW \| OW | Plastid |
|---|---|---|---|---|
| MRCA of *lo* + *lu* | 37 (31) /17 | 40 (40) /3 | 35 (30) /5 | 18 /22[a] |
| Stem age of *lo* lineage | 29 (26) /17 | = MRCA | 28 (30) /5 | <5 /6 |
| Stem age of *lu* lineage | 37 (21) /16 | = MRCA | = MRCA | – /–[b] |
| Lin. IV core clade stem age | – /– | – /– | – /– | 15 /18 |
| Lin. IV core clade crown age[c] | – /– | – /– | – /– | 9 /12 |
| MRCA of IV-ChA + IV-ChB[d] | – /– | – /– | – /– | 6 /8 |
| MRCA of IV-ChB | – /– | – /– | – /– | 5 /7 |

[a] MRCA including SLPT IV-LuA; topology-wise identical to the plastid MRCA of *F. hayatae-pashanica* (**Table SR3-6**)
[b] If no age estimate are given, the according node/aspect does not apply for the respective dated tree.
[c] as defined by the MRCA of the comprising SLPTs
[d] SLPT IV-ChA is shared by *F. lucida* and *F. pashanica*; SLPTs IV-ChB can be found in individuals of *F. longipetiolata, F. lucida* and *F. engleriana* of *F.* subg. *Engeleriana*.

In contrast to *F. lucida* and the cryptic sister species *F. hayatae* and *F. pashanica,* the so far still limited plastid data on *F. longipetiolata* shows a relatively high coherence (complete plastomes representing a single SLPT, IV-ChB, the two *F. longipetiolata* plastomes placed as sister tips; cf. **Fig. SR2-2**). The last common mothers (LCM) of *F. longipetiolata* and the SLPT IV-ChB are estimated to be of latest Miocene age (**Table SR3-7**). The *Longipetiolata*-LCM may have been exclusive to the precursor of the extant species, coming from the same lineage that also acted as a maternal donor of some *F. lucida* and *F. engleriana*; a lineage formed during the latest radiations within the Lineage IV core clade. This maternal donor lineage, with an estimated Early Miocene stem and Middle Miocene crown age (**Table SR3-7**) must have had a cross-North-East Asian distribution as its constituent SLPTs include the SLPTs IV-CrD (first diverging branch) and IV-CrE (sister type) of southern and central Japanese *F. crenata*. The palaeo-air-distance to the region of IV-Ha and IV-LuA plastomes was > 1100 km, and > 3000 km over land when avoiding the then hot-subtropical, near-tropical, beech-hostile lowlands (Scotese 2014b, Scotese et al. 2021). All available nuclear genetic data on *F. longipetiolata* indicates that while its ancestors underwent reticulate evolution, the modern-day species is likely holophyletic: the *Longipetiolata*-LCM was also the last common father of the extant populations, i.e. represents the *Longipetiolata*-LCA. As in the case of *F. crenata*, the *Longipetiolata*-LCA marks the endpoint of a North-East Asia lineage. Among the *F. longipet-*

*iolata*-like fossils of and prior to the Late Miocene of North-East Asia are the ancestors of the extant species; this lineage as well migrated into southern East Asia during the Miocene. During the migration process, it came into secondary contact with and was introgressed by the ancestors of *F. lucida* (shared nuclear alleles + SLPT IV-ChB), and eventually evolved into the *Longipetiolata*-LCA.

##### 3.3.5 The legacy of lost lineages and radiations

The Lineage IV plastomes carried by *F. japonica* and *F. engleriana*, the most recently evolved sister species pair within *Fagus* subgenus *Englerianae* (**Table SR3-9**), differ fundamentally in their genetic distinctness (**Fig. SR2-5**). They represent different radiations within East Asian beeches predating *F. engleriana* | *F. japonica* speciation event. In case of the *F. japonica* Lineage IV plastomes (SLPTs IV-JaC–G; **Fig. SR2-2**), the total data (molecular differentiation patterns and fossil record, **SupplFossilTable.xlsx** and **Section 4**) and resulting age estimates easily fit within an ancient introgression of a lost lineage scenario: a group of ancestral beeches that radiated and speciated across (northern) East Asia since the late Eocene and whose plastomes had started to drift apart, which were then introgressed at a large scale by early members of *F.* subgenus *Englerianae* when migrating south from their high-latitude cradles. The plastid heterogeneity was passed on to later species of *F.* subgenus *Englerianae* until the final radiation and speciation that formed the modern-day *F. japonica.* Two basic processes could have stabilised the coexistence of various maternal lineages at a small regional scale: (*i*) lack of severe extinction events, such as the availability of multiple refugia until and during the Pleistocene climate fluctuation in the Japanese archipelago (Tsukada, 1982; Kimura *et al.*, 2014) and (*ii*) subsequent speciation events within *F.* subgenus *Englerianae* leading to a temporary sorting of SLPTs, followed by intra-subgeneric introgression, hybridisation and homogenisation of the nucleome. The latter is evidenced by the high diversity and polymorphism of the non-coding nuclear-ribosomal rDNA spacers (Grimm et al. 2007) and poor sorting of allelic nuclear variation in the extant species of *F.* subgenus *Englerianae* (data of Jiang et al. 2022; see Cardoni et al. 2022, data S5, **SupplGenetics_ncDNA.xlsx**, sheet *JiangEtAl*; for a detailed list of mutational patterns see **OSA**, file *00_bits.xlsx*). Some remnants of this ancestral beech lineage and early radiation may have survived in the Japanese archipelago until the latest Miocene or Pliocene as species, being finally introgressed by *F. crenata* (SLPT IV-CrC, sister type of IV-JaG).

The case of its sister, the continental, southern East Asian *F. engleriana,* is more puzzling. Here, we find plastomes that are much more similar to those of sym- to parapatric species of *Fagus* subgenus *Fagus* thriving in the same larger area, close to each other, and occasionally forming mixed stands. They are all part of the Lineage IV core clade, reflecting the last major radiation event within this lineage, a radiation event that did not involve any of the maternal donors of its sister species *F. japonica.* However, each of the three completely sequenced plastomes of *F. engleriana* has different phylogenetic affinities within the Lineage IV core clade.

- SLPT IV-EnA is a sister type of the most recently diverged Lineage IV plastome of the Japanese *F. crenata*. It could come from a North-East Asian species of *F.* subgenus *Fagus* that diverged after the large-scale introgression by the ancestors of *F. japonica* (no Lineage IV core clade plastomes in *F. japonica*); carried southward when the precursors of *F. engleriana* migrated into their current distribution area.
- SLPT IV-EnB is an isolated plastome type within an exclusively Chinese (southern East Asia) *F.* subgenus *Fagus* clade. Its source can only be a species of *F.* subgenus *Fagus* that was already in place, when the precursors of *F. engleriana* moved in.
- SLPT IV-ChB comprises a *F. engleriana* plastome highly similar to a plastome obtained from a *F. lucida* individual. It either represents an introgressed precursor of *F. lucida* or coevally introgressed (by the precursors of *F. engleriana* and *F. lucida*) southern(?) East Asian member of *F.* subgenus *Fagus.*

On the background of dated phylogenies and with respect to the palaeogeography/-topography and paleoclimate framework (Scotese 2014b; Scotese et al. 2014, 2021), the simplest explanation would be that *F. engleriana* introgressed the other Chinese species at a large scale, thus, picking up their plastid signatures. However, there is no evidence for ongoing or (sub-)recent uni- or bidirectional gene flow between the two subgenera. If *F. engleriana* would have picked up these SLPTs from the extant species of *Fagus* subgenus *Fagus*, there should be some evidence for introgressed alleles and ITS variants of the other subgenus in the nucleomes of *F. engleriana* and/or the (today) Chinese species of *F.* subgenus *Fagus* and at least a few reports of phenotypically ambiguous individuals in accordingly mixed stands.

An alternative scenario is that (the precursor of *F. multinervis*-)*F. japonica,* when extending its range on the continent, introgressed some of the local, now extinct beech species; the same did the precursors of *F. longipetiolata* and *F. lucida.* Being only genetically compatible with their introgressors, the nucleomes of the introgressed native populations would have been quickly homogenised by their respective paternal lineage; leaving only their plastomes evidencing their shared ancestry (cf. **Fig. SR3-3**). The higher fitness of the introgressors and, in particular, the introgrades regarding a changing environment would have led to the extinction and total replacement of the original native species, the non-introgressed populations. Latest with the Pleistocene fluctuation, the source (North-East Asian) populations of the introgressors, and subsequently, their plastome lineages, would have died out, while the introgressed southern populations (in modern-day central and southern China) would have much less affected by any Ice Age triggered bottlenecks. Thus, one cannot find any Lineage II or III(-like) plastomes, or plastomes closer to these lineages and Lineage V, in contemporary *F. engleriana, F. longipetiolata* and *F. lucida* as one would expect under a northern introgressor scenario. Nonetheless, a more in-depth exploration of the nucleome differentiation of the Chinese species across their entire range and focussing on potential relict stands (such as Sichuan, Long Xi Shan, and Shennongjia Forest District) may reveal alleles and gene regions reflecting this lost southern, lower-latitude lineage; nuclear genes that can be correlated with the plastid Lineage IV core clade.

For the simultaenous-introgression-and-replacement hypothesis, we can indirectly time-stamp the introgressive event. "Chloroplast capture" requires the migration of one or several species into the area of another lineage that carries already a distinct, diverged plastome (cf. **Figs SR3-3, SR3-4**). When "captured" by different introgressors, the plastomes will independently drift further apart from each other. Plastomes of the same introgressor may diverge further, too, but will remain part of the same clade and species-exclusive (form sister/sibling haplotypes, as in the case of the two *F. longipetiolata* plastomes). The introgressive event must have happened after the latest divergence observed in the "captured" plastome lineage (in this case the Lineage IV core clade) between introgressors from different evolutionary lineages or species of the same lineage. For the Japanese archipelago, the introgressive event eliminating Lineage IV core clade as a distinct biological entity can be placed in the Late Miocene (≤ 10 Ma; divergence between SLPTs IV-CrC and -JaG) while that of their continental, southern East Asian counterparts can be linked to (early) Pleistocene climate and range fluctuations (≤ 1 Ma, between *F. engleriana* and *F. lucida* individuals carrying SLPT IV-ChB; median FBD-estimates, **Table SR3-8**).

**Table SR3-8 | Genetic distinctness and divergence estimates for species-level plastid types (SLPT) detected so far in *F. engleriana*.** Given is the number of differing SNPs ($\Delta_{SNP}$, based on **SupplGenetics.xlsx**, sheet *PlstmDissim*, see also **Fig. SR2-4**) to the respective sister tip (in case of subtrees: tip with the shortest root-tip distance in the ML tree in **Fig. SR2-2**) and the age (in Ma; median FBD- and ND-estimates) of the according node. $\Delta_{SNP}$ and divergence for the sister SLPT IV-CrC (*F. crenata*) and -JaG (*F. japonica*), the genetically closest *F. crenata* and *F. japonica* SLPTs within Lineage IV, are given for comparison.

| SLPT, individual, node | Sister tip (SLPT, individual) | $\Delta_{SNP}$ | Age [Ma] |
|---|---|---|---|
| MRCA of Lineage IV | | Max. | 19 /22 |
| MRCA of Lin. IV core clade | | Max. | 8.6 /12.1 |
| IV-EnA: en34\|China | IV-CrE: cr03\|Kyushu | 98 | 8.6[a] /9.4 |
| IV-EnB: enGB\|KX852398 | IV-ChB: enGB\|MT762293[b] | 60 | 5.5 /7.2 |
| IV-ChB: enGB\|MT762293 | IV-ChB: lu37\|China | 15 | 1.0 /1.7 |
| IV-CrC \| IV-JaG | IV-CrC: cr06\|Kanto; IV-JaG: jaGB\|MT762294 | 101 | 9.5 /11.5 |

[a] Not sister tips in the MCC tree optimised via the FBD-model; equals the MRCA of Lin. IV core clade.
[b] Not sisters; the first diverging branch within IV-ChB subtree are the two *F. longipetiolata* plastomes.

An ancient remnant of once widespread continental North-East Asian (and northern Asian) beeches are clearly the Lineage III plastomes that survived in the beeches of the Korean island Ulleungdo in the Sea of Japan, *F. multinervis*, and a *F. japonica* population just on the other side of the Sea of Japan (cf. **Fig. SR2-6**). Their composition places them between the East Asian Lineage IV and Western Eurasian Lineage V plastomes, in perfect fit with their modern-day distribution area in the centre of the Oligocene and Miocene beech biodiversity hotspot (Denk & Grimm 2009; this study). According to a very conservative estimate, using only a single root age constraint, i.e. largely uninformed by the fossil record, Lineage III beeches were diverged by the late Eocene (> 38 Ma; cf. **Fig. SR3-2**). Their lack of genetic drift points to a much larger population size in the past; their phylogenetic affinities and endemism make it unlikely that they represent original plastomes of the subgenus-*Englerianae*-FCA. Within the extant species of *Fagus* subgenus *Englerianae, F. multinervis* is the earliest diverged one (**Table SR3-9**) but it cannot have been evolved on Ulleungdo as this volcanic island is much too young with the main volcano-building stage at Ulleung-do starting after 2.7 my (Song et al. 2006). Our age estimates place the geographic fragmentation of Lineage III SLPT III-Mu between north-western mainland (Korean) and south-eastern island (Japan) populations in the Pliocene, slightly younger than the estimates for the split between *F. multinervis* and *F. engleriana-japonica* (**Table SR3-9**).[24] *Fagus multinervis* is not a typical endemic island species but rather a leftover population of earlier radiations, a sort of 'genetic fossil', which has

[24] Higher estimates obtained with High- | Low-latitude scenario are due to the particularities of the according gene sample and probably represent earlier speciation events retained in the plastome of the modern-day species in the form of intra-specific, intra-genomic diversity.

conserved the plastid signatures of the once widespread high-latitude beeches of continental North-East Asia and adjacent regions picked up by its ancestors. Being morphologically indistinguishable from *F. engleriana* (Shen 1992; Denk 2003), it may further represent the archetype of the *F. japonica*-related introgressor that escaped south in wake of the Pleistocene glaciations and became the modern-day *F. engleriana*.

**Table SR3-9 | Divergence age estimates (FBD/ND medians, rounded to full myrs) for *F. multinervis,* Lineage III plastomes (SLPT III-Mu), and divergence between *F. japonica* and *F. engleriana*.** Values in brackets give the estimates for the phylogenetically equivalent node (MRCA; see footnotes for details).

| Node | Topology<br>Atl \| Pac | High \| Low | NW \| OW | Plastid |
|---|---|---|---|---|
| *F. multinervis* stem age | 6 /6 | 19 /9 | 7 /5 | = Lin. III crown age |
| Lineage III stem age[a] | – (61/:≥34) | – (68/:≥34) | – (73/:≥34) | 35 /32 |
| Lineage III crown age[b] | – (6/6) | – (19/9) | – (7/5) | 4 /3 |
| *F. engleriana* \| *F. japonica* | 3 /3 | 3 /3 | 5–3[c] /3 | – (44/37) |

[a] In nuclear dated trees, the topological equivalent would be the stem age of *F.* subg. *Englerianae*, i.e. the age of the MRCA of all extant spp. (bracketed values); ND-estimates are set to ≥ 34 Ma for the nuclear topologies and ≥ 28 Ma for the plastid topology by according age priors.
[b] Split between the island mother population, *F. multinervis*, and its *F. japonica* sister population across the Sea of Japan. The topological equivalent in the nuclear dated trees would be the MRCA of *F. multinervis* and *F. japonica*, i.e. the MRCA of extant species of *F.* subg. *Englerianae,* which equals the *F. multinervis* stem age (bracketed values).
[c] Depending on the treatment of the Kraskino palaeoflora fossils; does not matter for the other two topologies.

### 4 Palaeogeographic framework for beech evolution and differentiation

In contrast to many top-down biogeographic papers on extratropical northern hemispheric tree genera or other organismal groups, which predominately rely on modern-day geographic-political concepts for identifying biogeographic regions and reconstructing ancestral areas, we here consider geographic regions (**Fig. SR4-1**) as representing natural units, characterized by their vegetation, topography and dominant climate, in space *and* time. These areas either have provided or still provide suitable habitats for beech within its primary ecological-climatic niche (cf. Maycock 1994, Peters 1997). As Earth's climate, biosphere and lithosphere are evolving and changing, the position and size of these areas have shifted through time. In the following, we describe the main biogeographic regions that had or have been playing and important role in the evolution of beech across the Northern Hemisphere.

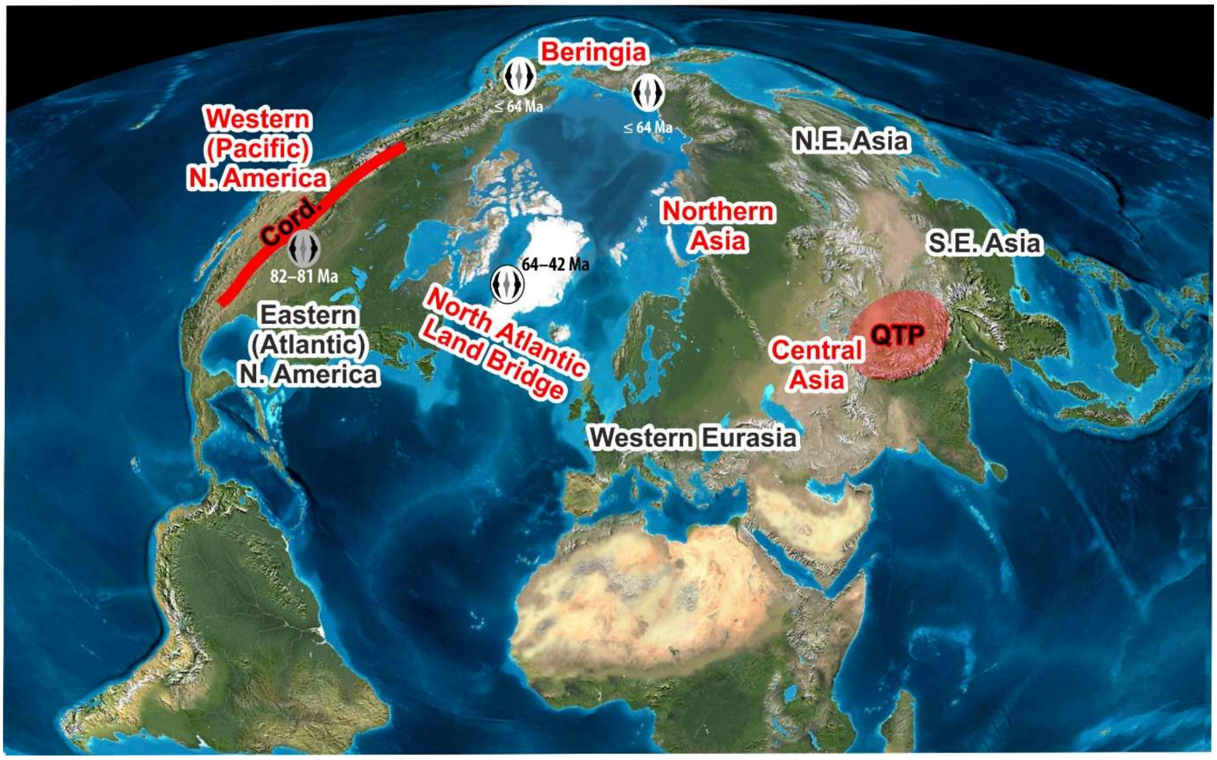


**Figure SR4-1 | Main biogeographic areas for beech during the Cenozoic.** Shown is a 50°N centred projection of today's Earth (provided by R. Blakey for Denk & Grimm 2009). Extinct areas are shown in red font. The localities (sites) with the earliest (Late Cretaceaous–Paleocene) fossil records of ancient (grey; oldest known member of the lineage leading to *Fagus*; Grímsson et al. 2016) and modern (white) beech (potential last common ancestors) are indicated by pollen signatures. N. E. Asia = coastal North-East Asia; S. E. Asia = montane southern East Asia. Major continental barriers: Cord. = Cordilleran main chains. QTP = Qinghai-Tibetan Plateau.

**Origins of modern beech**—Oldest fossils of beech are dispersed Paleocene pollen from high-Arctic palaeolatitudes. With a distinctly temperate area, they were climatically not unlike (Korasidis et al. 2022) the modern-day climax niche of the genus (Grimm & Denk 2012, and literature cited therein): fully temperate, perhumid, oceanic climates (Köppen-Geiger zones *Cfb/Dfb*; Köppen-Trewartha zones *Do/Dc*). Thus, a High-Arctic origin (impossible to reconstruct using top-down biogeographic methods: Jiang et al. 2021) can be assumed as well as a wide distribution of the Modern-Beech-LCA(s), the last common ancestor(s) of today's beeches (**Fig. SR4-2**). With respect to the spread of the ultimate stem group of modern-day beech comprising the Modern-Beech-LCAs, the deep split seen in the plastomes between (eastern) North American and Eurasian beeches is unsurprising. The Modern-Beech-LCAs and the first emerging species lineages of modern beeches may have been nucleome-wise still homogenous but plastid-wise already polymorphic. Given the palaeotopographic situation (the North Pole being largely covered by sea), these primordial beeches would have showed a starting west-east differentiation in their slowly evolving plastomes. Irrespective of later speciation events and secondary contact, the eastern North American beeches would have the highest likelihood to retain the original eastern (Atlantic) plastome lineage, i.e. Lineage I, while any western (Pacific) beech, would have an already different plastome (Lineages II–V).

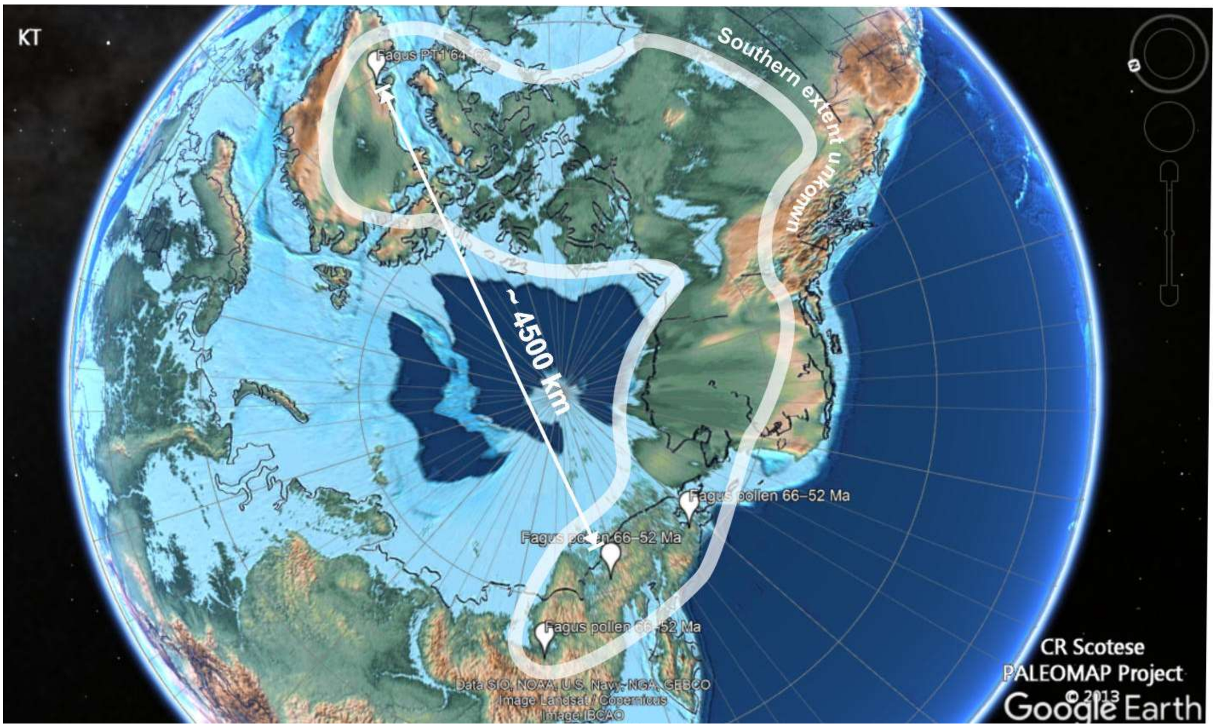


**Figure SR4-2 | Screenshot of earliest record of beech mapped on a palaeoglobe representing the Earth at the Cretaceous-Paleogene (KT) boundary** (Scotese 2014a). GoogleEarth kmz files including all fossils considered in this study mapped on the next-older palaeoglobe layer (Scotese 2014a,b) are included in the ODA.

**NAT: Northern North Atlantic**—The continental pieces (Canadian Continental Shield, Greenland Continental Shield) and volcanic islands (modern-day Iceland, Faroes, British Igneous Province) forming the North Atlantic Land bridge (Denk, Grímsson et al. 2011) are possibly one of the cradles of beech (**Fig. SR4-2**) but also a corridor for later introgression from (northern/eastern) North America into Western Eurasia. Until the onset of the Pliocene, this area enjoyed an oceanic, mild subtropical to temperate climate. Including an old craton (continental landmass: Greenland Continental Shield), the topography would have been steep enough to support an altitudinal vegetation gradient during warmer and colder phases. The earliest record of beech consists of pollen indistinguishable from modern-day beech pollen as part of the extremely diverse Agatdalen plant assemblage, which combines distinctly subtropical, temperate and boreal elements (Grímsson et al. 2016a, 2016b). The pollen-producing beeches would have grown in the (cold-)temperate to boreal altitudinal belt above a rich lowland subtropical forest. These NAT beeches would have always been in contact with those of lower latitude (lowland to montane, to mid-altitude) 'Eastern North America', with the Canadian Subarctic plains being episodically populated by all Atlantic beeches; the according forest biome is documented from Axel Heiberg Island (McIver and Basinger 1999). The Gulf Stream, which is part of the Global Conveyor Belt, perpetually transports heat into this area, buffering and delaying global cooling episodes (Denk et al. 2013). The last known NAT beech is †*F. gussonii*, a morphologically ambiguous taxon and possible vector for introgression of North American/NAT gene variants into the gene pool of Western Eurasian beeches (Cardoni et al. 2022; this study). This morphotype can be found in Iceland at 9–8 Ma, but earlier and contemporary records are from across the Mediterranean region (**Fig. SR4-3**).

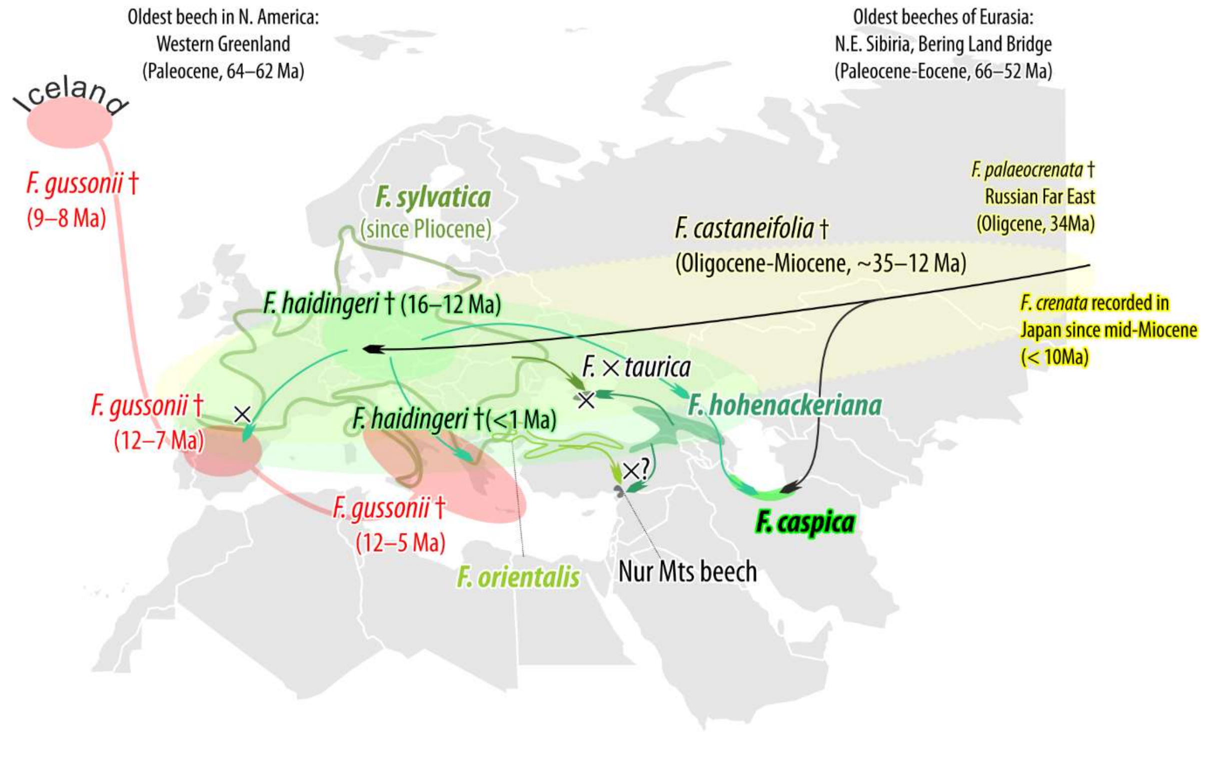


**Figure SR4-3 | History and modern-day distribution of beeches in Western Eurasia (modified from Schulze and Grimm 2022).** Yellow-green colours, fossils and modern species of the Western Eurasian clade of *Fagus* subgenus *Fagus*; red, †*F. gussoni*, showing an ancestral, ambiguous morphotype with similarities to both *F. longipetiolata* and *F. sylvatica* (**SupplFossilTable.xlsx**, sheet *main list*; unresolved within *F.* subg. *Fagus* in the phylogenetic reconstructions of Denk & Grimm 2009).

**ENA: low- to high-latitude Eastern (Atlantic) North America**—Today the sole area inhabited by New World beeches. ENA beeches occur from boreal lowlands along the St Lorenz Stream, with relatively long summers and generally mild, ocean-buffered winters with persistent snow cover (boreal *Dfb* climates) into the Appalachian Mts, where they dominate the forests (mostly fully temperate *Cfb*) into lowland, hot subtropical (*Cfa*) littoral forests of Georgia and northern Florida (Maycock 1994). The lack of fossil records from this area is to some degree a taphonomic bias (few Paleogene leaf-bearing localities) but may reflect a true pattern based on the absence of *Fagus* pollen from this area (Frederiksen 1979). Nonetheless, available palaeoclimatic maps (Scotese et al. 2014, 2021) indicate that most of the modern-day area, lacking steep topography in contrast to Western Eurasia and East Asia, would have been hard to colonise during global greenhouse phases dominating the Paleogene. Since north-south barriers are entirely absent, it is likely that during such phases, ENA beeches evaded to high latitudes, thus forming a continuous area and master population with (western) NAT beeches. Likewise, western NAT beech populations could have easily migrated southwards during global ice house situations (e.g. during the Oligocene and Plio-Pleistocene cooling phases); in combination, ENA + NAT would have provided a large potential area for beeches throughout the entire Cenozoic. The dry (continental) interior with the Cordilleran Mountain chains, and related to them, the dry interior, would, on the other hand, have been a persistent barrier to any exchange between Atlantic- and Pacific-bound beeches at low- and mid-latitudes, and high-altitudes unless the global temperate was (much) hotter than today (e.g. during the early to middle Eocene or the Mid-Miocene Climatic Optimum).

**WNA: Western (Pacific) North American coastal highlands**—In this area, characterised by a rugged topography during the entire Cenozoic, beeches are concentrated in the so-called "Okanagan Highlands" of British Columbia and Washington (state) during the Eocene (**Fig. SR4-4**) to the Miocene of British Columbia and the Pacific Northwest (U.S.; e.g. LaMotte 1936; Manchester & Dillhoff 2004). Originally a candidate for the origin of beech (cf. Denk & Grimm 2009), the beeches of the Okanagan Highlands likely represent a dead end of the original Beringian populations that moved south along the mountains superposing the then (hot) subtropical lowlands (Greenwood & Archibald 2005). The main Cordilleran chain, however, with its high (today and in the past alpine) mountain ranges and seasonally dry intra-montane basins as well as the continental, generally dry interior pre-montane plains (beeches are lacking from the transition zone where *Cfa/Dfa* climates approach *BS* climates) would have prevented them from crossing into ENA. The area was probably always limited and highly competitive with other Fagaceae, maples, and various conifers populating the same niche, hence, no beech survived in WNA despite the presence of suitable climate pockets until today. How far beeches reached south along the Pacific-facing Cordilleran foothills is unknown. The relict populations of *F. mexicana* in central Mexico are on the Atlantic side but have only been superficially studied so far: nuclear-genetically (data compiled by Jiang et al. 2021) they seem to be less evolved and homogenised than their northern sisters but according to preliminary plastid data stored in gene banks, they share the Lineage I plastomes. This would indicate that they represent a relict of the first ENA-NAT beeches extending into the low latitudes, as there is no evidence to connect them with the extinct lineage of the beeches of the Pacific side, the WNA beeches, which could be expected to carry a substantially distinct plastome (phylogenetically in between Lineage I and Lineage II plastomes, or close to the latter).

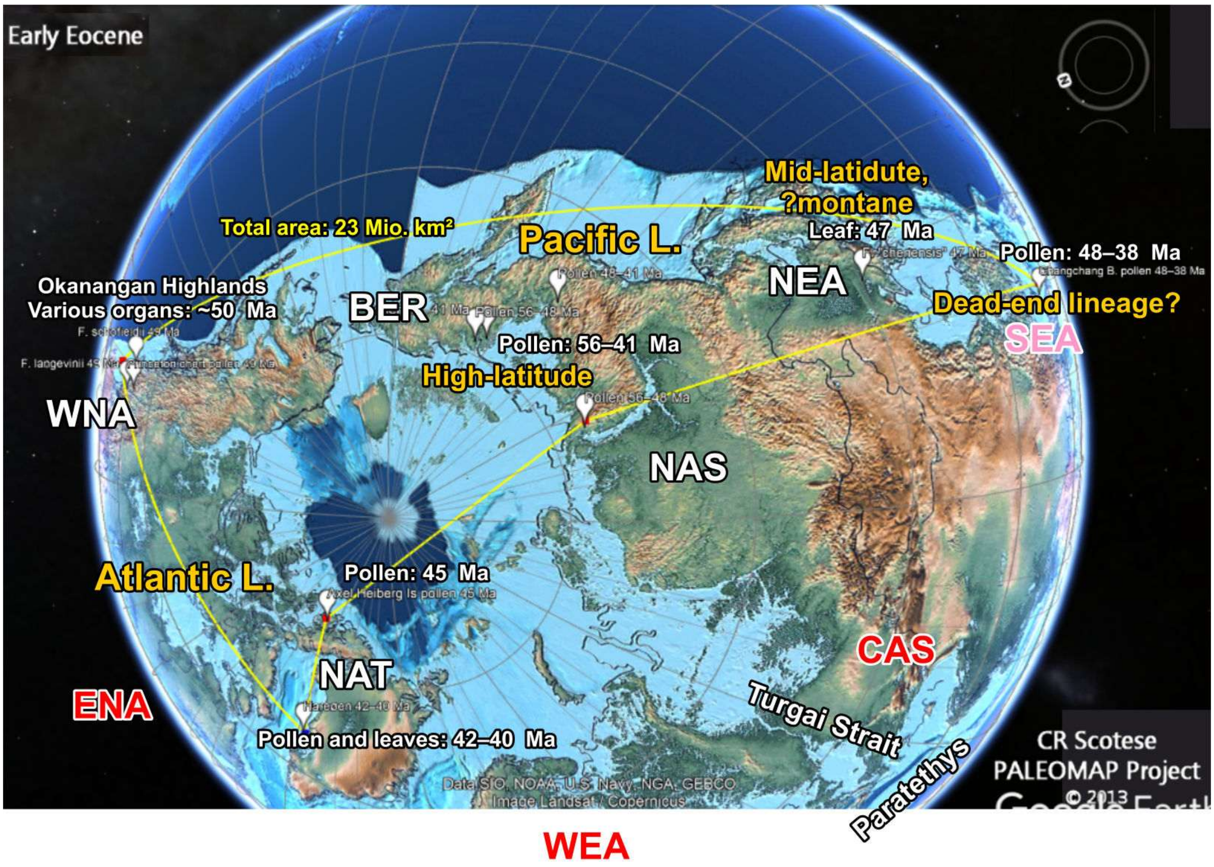


**Figure SR4-4 | Annotated screenshot of the early Eocene record of beech.** Biogeographic areas abbreviated as in the text, areas with fossil beeches in white font, areas with no beech record from this time period in red font. The yellow polygon gives the minimal distribution area. The gap between easternmost northern North Atlantic (NAT) and westernmost northern Asian (NAS) finds may be a sampling artefact (no fossil lagerstätten/limited palaeobotanical research). But note the chains of connecting islands close to the north pole. In contrast, the Eocene of Western Eurasia (WEA) is very well studied. Based on modern-day distribution of main plastid lineages (**section 3.3.4**, the Eocene Hainan beech represents an outlier/dead-end.

**BER: Beringia**—Like the North Atlantic Land Bridge, the Beringian Land Bridge represents a potential cradle (**Fig. SR4-2**) and crossroad for beech (**Fig. SR4-4**). During global greenhouse situations, this corridor could have been repeatedly used by New World as well as Old World beeches for cross-continental exchange (see also Renner et al. 2008 for maples), but also as a refuge for mid-latitude, coastal lowland beeches of (northern) East Asia. The oldest beeches that can be morphologically referred to as *Fagus* subgenus *Englerianae* (†*F. evenensis* Chelebaeva) are from Kamchatka, the western part of BER. This area may have served as a source area for both northern Asian (palaeo-)plains and Central Asian mountain chains during the early Eocene (global greenhouse phase) and the coastal forests of North-East Asia during the late Eocene/Oligocene global cooling (**Fig. SR4-5**).

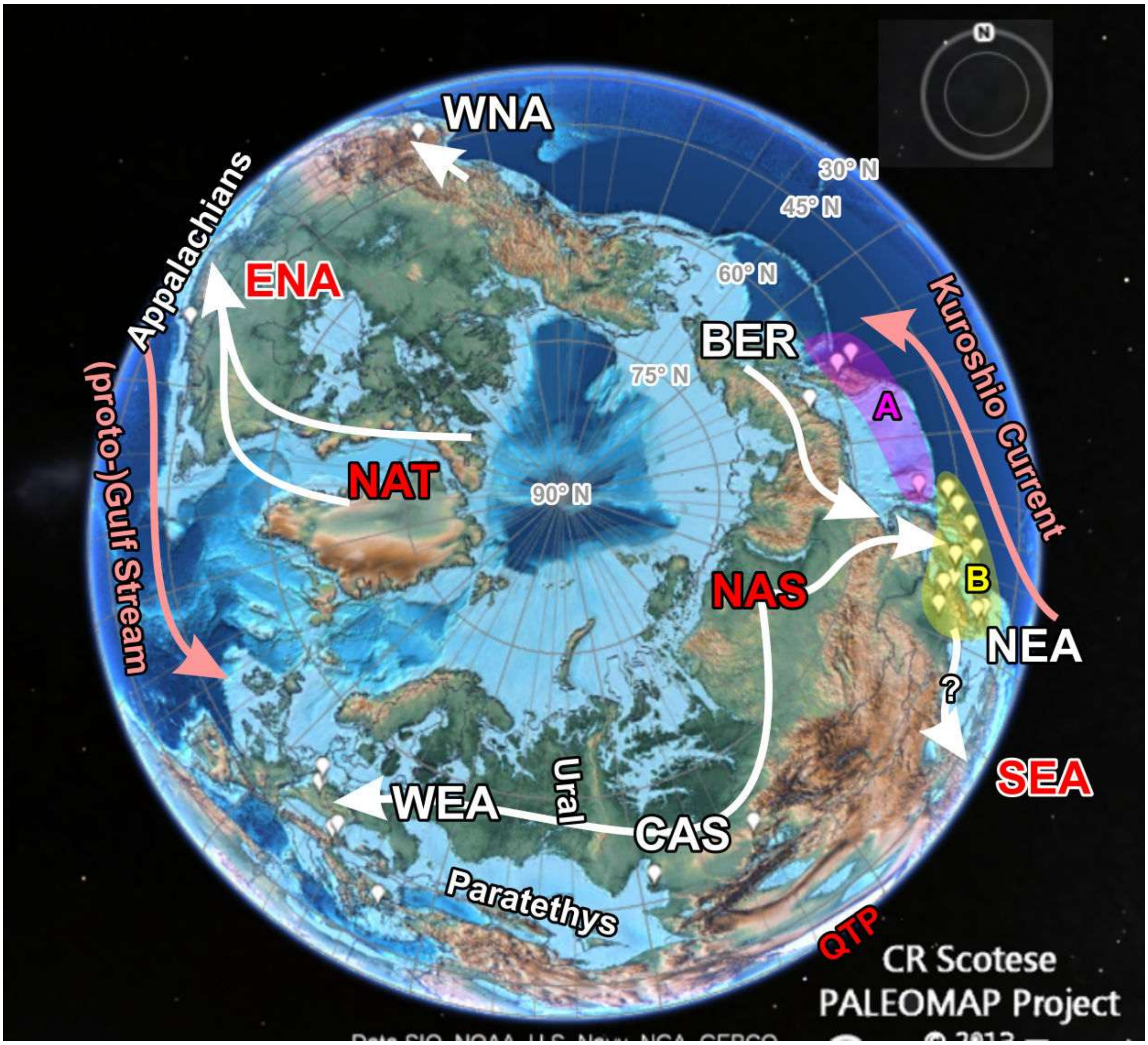


**Figure SR4-5 | Annotated screenshot of the late Eocene and Oligocene record.** Biogeographic abbreviated as in the text, white with, red without reported beech fossils. **A** = area of first morphological *Fagus* subgenus *Englerianae* fossils (fossil-species †*F. evenensis*), **B** = area of highest morphological diversity including *F. longipetiolata*-like morphotypes, morphotypes leading towards *F. crenata*, and extinct morphotypes with affinities to North American fossils and extant species. QTP = centre of the Qinghai-Tibetan Plateau (proto-Himalayas).

**NEA: North-East Asia**—During the Oligocene–Miocene a biodiversity hotspot of beech, with an extremely rich and diverse fossil record (**Figs SR4-5–SR4-7**). Fossils of this area show phenotypes characteristic for both the *F. crenata*(–*F. sylvatica*) and *F. longipetiolata*–*F. hayatae-pashanica* lineages. These morphotypes coexisted with those of the Pacific (BER+WNA)–modern North American (*F. grandifolia-mexicana*; ENA) lineage (Denk & Grimm 2009). North-East Asia was an area where beeches of different provenances (BER—Beringia, NAS—'northern Asia', SEA—'southern East Asia') and evolutionary sources (*F.* subg. *Englerianae*, *F.* subg. *Fagus*, extinct lineages) could have come into secondary contact to exchange genetic material or find refuge. Genetically, this legacy is well-documented in the plastome diversity of the species still thriving in this area: *F. multinervis* and *F. japonica* of *Fagus* subgenus *Englerianae*; *F. crenata* of *F.* subgenus *Fagus*.

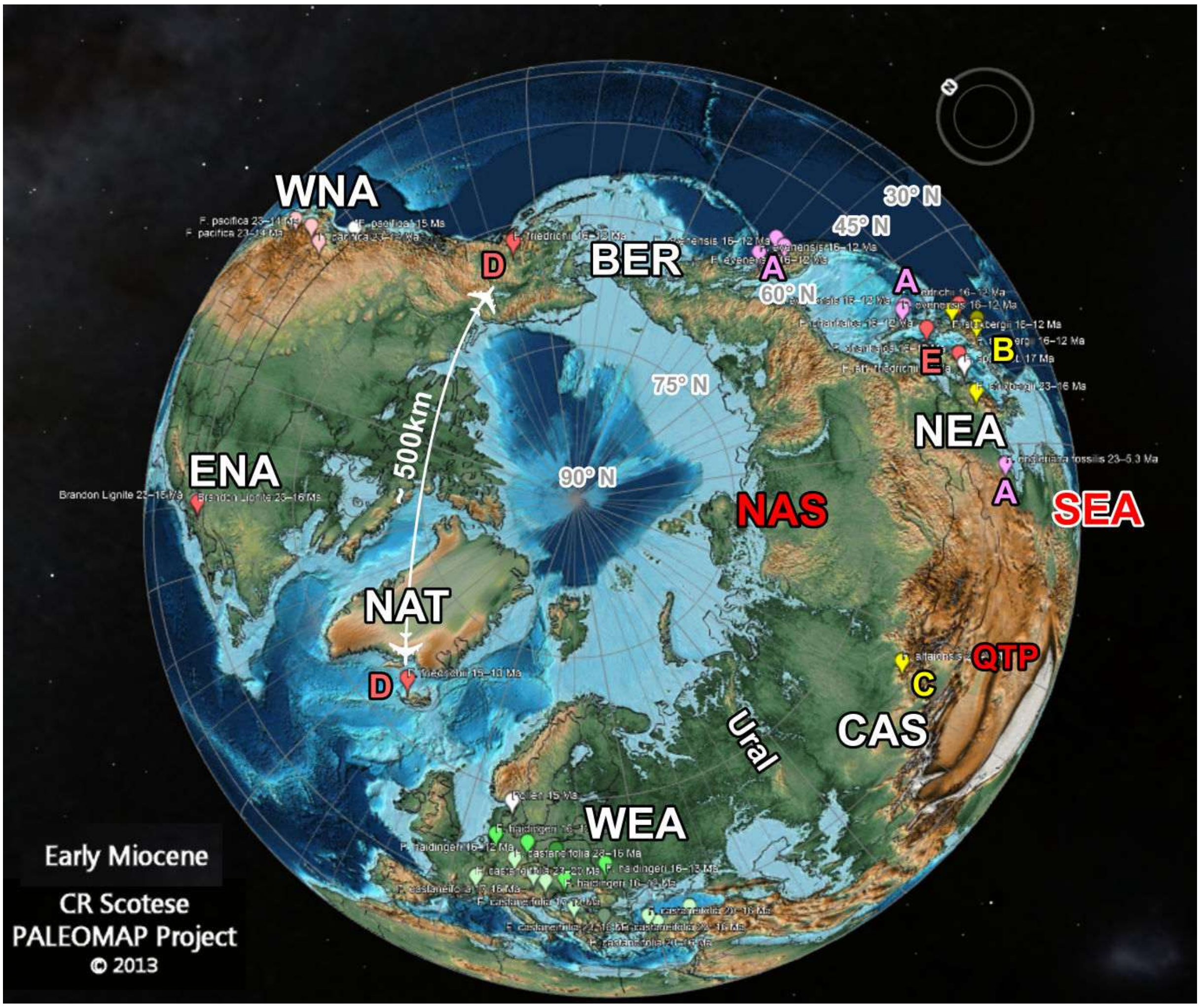


**Figure SR4-6 | Annotated screenshot of the early and mid-Miocene fossil record.** This time period is characterized by a strong phenotypic-palaeogeographic sorting. A = *Fagus* subgenus *Englerianae*, stretching along the Pacific coast lowlands. B = *F. longipetiolata* + *F. hayatae-pashanica* lineage, C = †*F. altaensis*, a fossil precursor of *F. lucida*. D = Widespread †*F. friedrichii*, an early member of the *F. grandifolia-mexicana* lineage. E = NEA fossils with similar morphotypes to *F. grandifolia*-†*F. friedrichii*

**SEA: montane southern East Asia—**This area represents the southward extension of littoral low- to mid-altitude NEA. During most of the Paleogene, the lowlands in this area were covered by (hot) tropical-subtropical moist forests, where beeches would have to compete with better adapted Fagaceae such as *Castanopsis* and evergreen oaks of *Quercus* subgenus *Cerris* (sections *Cyclobalanopsis, Ilex*) and other Fagales such as Engelhardioideae (Juglandaceae); fossil records from this area are notably scarce. Today beeches in this area are strictly confined to the mid- and high altitudes of montane, ± inland areas (Peters 1997) and the mountains of central-northern Taiwan. For example, in the Shennongjia Forest District, the possibly least disturbed natural forest region in central China, beeches of both subgenera (*Fagus engleriana, F. longipetiolata, F. pashanica*) do not go below 1000 m a.s.l., with a mean annual temperature, $T_{\mathrm{ann}}$, of < 12 °C (Zhu and Song 1999, Grimm & Denk 2012). With respect to the modern-day situation, which in many aspects is comparable to the situation in the area during most of the Neogene (hot lowland, rugged topography providing many albeit relatively small niches for temperate-oceanic trees; cf. Scotese 2014b; Scotese et al. 2014, 2021), the fossil record of SEA is probably underestimated. In analogy to western North America and Mexico, while being easily connected with NEA, SEA is otherwise a dead-end for beech migration. The steadily expanding Qinghai-Tibetan Plateau and preceding dry-continental hilly areas would have blocked any expansion into Central Asia during most of the Cenozoic Neogene (**Figs SR4-4**–**SR4-7**). The only possible migration corridor into Western Eurasia would have been a small and stable temperate belt south of the Himalayas: the "Himalayan corridor" (Denk & Grimm 2010). Notably, the most temperate lineage and only deciduous lineage of the exclusively Eurasian *Quercus* subgenus *Cerris*, section *Cerris,* the cork oaks, are absent from the Himalayan corridor as well, despite their pre-adaptation to summer droughts, i.e. climate settings intolerable for beech (cf. Denk et al. 2023). That there are no modern-day beeches in the *Cwa/Cwb* climates of south-western China, home of many Cenozoic relict species (e.g. endemic maple species with primitive morphologies but high-distinct plastomes: Renner et al. 2008; see also Grimm 2022), fits with the complete lack of fossils from this area, and precludes the possibility that any beech used this corridor to move westwards from SEA.

**NAS and CAS: northern and Central Asia—**Despite a patchy fossil record, we can assume that beeches existed in this area during the Paleogene and into the Miocene. During greenhouse phases, the humid-temperate forest belt probably extended from Beringia and North-East Asia via northern Asia into Central Asia bordering the Paratethys (Scotese et al. 2014, 2021). With the uplifting and expansion of the Qinghai-Tibetan Plateau (starting in the Eocene), the final closure of the Turgai Strait (early Oligocene) and the retreat of the Paratethys (Oligocene-

Miocene), both areas became more and more continental and inhospitable to beech. Our fossil data includes only a single, early Eocene NAS record and two from CAS, including the easternmost record of the precursor of the West-Eurasian lineage, †*Fagus castaneifolia* (south-east of the Ural Mts). The decline in total precipitation and shorter summers made these areas hostile for beech between ~15–10 Ma (cf. maps in Scotese et al. 2021). The second record, †*F. altaensis* from the Miocene of the Altay Mts (CAS; **Fig. SR4-6**) with a phenotype that links it to the Chinese *F. lucida* (Denk & Grimm 2009), may represent the last beech of this area. Nevertheless, we cannot exclude the possibility that local beeches persisted in areas like the Altay Mts – today with *Dfa, Dfb* climates, i.e. beech-favourable climates – until the Pleistocene cool phases and that there might have been contacts with the direct precursors of *F. caspica* stabilising ancient CAS/NAS remnant genotypes in their gene pool.

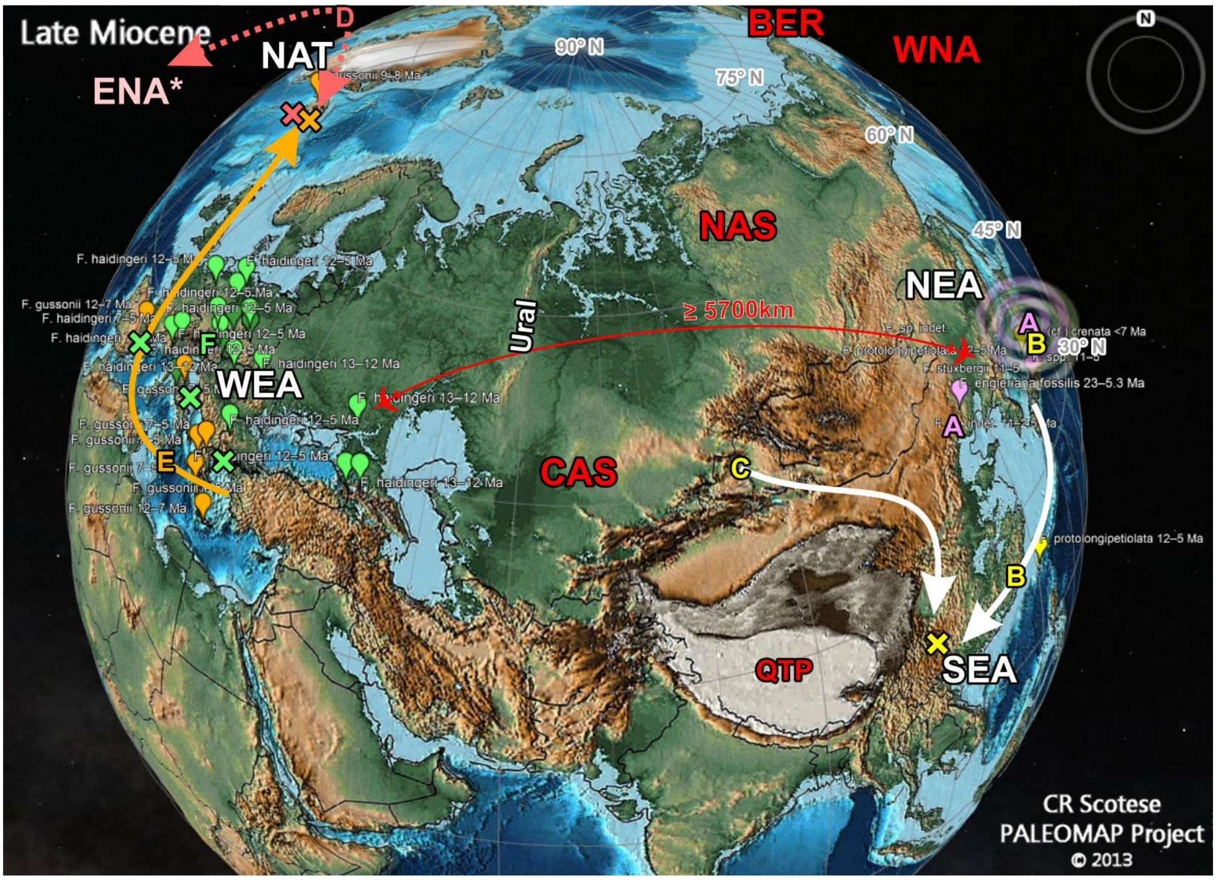


**Figure SR4-7 | Annotated screenshot of the Late Miocene.** **A** = *Fagus* subgenus *Englerianae,* stretching along the Pacific coast lowlands. **B** = *F. longipetiolata + F. hayatae-pashanica* lineage, co-existing in the Japanese archipelago with the precursor(s) of *F. crenata*, **C** = *F. lucida* lineage. **D** = Introgression of North American genes (* there are no Late Miocene lagerstätten in eastern N. America) into the W. Eurasian gene pool via †*F. gussonii* (**E**) into †*F. haidingeri* (**F**), the percursors of the modern-day W. Eurasian species (cf. Denk et al. 2024). × = Likely introgression/hybridisation events.

**WEA: western Eurasia**—The modern-day climate of a substantial portion of Western Eurasia (*Cfa, Cfb*, *Dfb,* perhumid with pronounced summers) is perfect for beech, and this has been the case for the entire Neogene into the Oligocene. The Alpine orogeneses have formed a multitude of west-east striking mountain chains along the south coast of the Paratethys and into Europe conserving the climax niche of beech during greenhouse and icehouse phases; which is reflected by their extremely well and continuous fossil record in Western Eurasia since the Oligocene (Denk 2004, Denk & Grimm 2009; **SupplFossilTable.xlsx**). Even the large-scale glaciations during the Pleistocene apparently did not threaten the beech, noting its quick recovery in the last 7000 years from (putative) relict stands in the Mediterranean region and further east. On the other hand, this particular combination of climate and topography would have provided ample possibilities for shorter-time isolation and speciation, especially with respect to the west-east extension of fossil and contemporary beech forests in this area. WEA transitions directly into NAS and CAS; as long as beeches could thrive in the latter areas, they would have been in contact with the eastern WEA beeches.

From the fossil record (Denk 2004, Denk & Grimm 2009) and (dated or not) molecular phylogenies (plastid or nuclear) and mutation patterns (Denk et al. 2005, Renner et al. 2016, Cardoni et al. 2022, data S5; this study) it is clear that initially WEA was colonised by beeches coming from the east (BER + NEA) via Siberia and the Asian interior north of the Paratethys (NAS + CAS). WEA is probably the latest area colonised by beech. The fossil species †*F. castaneifolia* did only migrate into WEA during the closure of the Turgai Strait (**Fig. SR4-5**), where it gradually evolved into and was replaced by †*F. haidingeri*, the precursor of all modern-day WEA species, during the Miocene (**Fig. SR4-5;** see also **Fig. SR4-3**). By the Late Miocene, any contact and gene flow between the beeches of WEA and any other area must have been severed: NAS and CAS became inhospitable to beech (too cool and/or dry), as well as NAT.

#### Additional related science-trivia and data resources by GWG

The challenging and puzzling ordinary beech – a (hi)story. Blogpost, *Res.I.P.*, posted 16/4/2018 (18 figs). https://researchinpeace.blogspot.com/2018/04/the-challenging-and-puzzling-ordinary.html

Can we depict the evolution of highly conserved genes, such as the ribosomal RNA genes? Blogpost, *Genealogical World of Phylogenetic Networks* (D. Morrison, ed.), posted 18/2/2019 (9 figs). https://phylonetworks.blogspot.com/2019/02/can-we-depict-evolution-of-highly.html

A fully resolved, and perfectly misleading, species tree. Blogpost, *Res.I.P.*, posted 29/9/2021 (17 figs). https://researchinpeace.blogspot.com/2021/09/a-fully-resolved-and-perfectly.html

Monophyletic species. Blogpost, *Res.I.P.*, posted 15/12/2021 (12 figs). https://researchinpeace.blogspot.com/2021/12/monophyletic-species.html

Fagaceae collection. Dataset, *figshare*. https://doi.org/10.6084/m9.figshare.11603547
