## Supplementary material for "Whole chloroplast genomes reveal a complex genetic legacy of lost lineages, past radiations and secondary contacts in the dominant temperate deciduous tree genus *Fagus*": SupplM&M

### Supplementary Materials & Methods

This supplement file details the preparatory and preliminary analyses and methodological approaches.

#### 1 Background: known issues and herein used species concept

The presentation of Worth et al. (2021) shows that, as in other Fagaceae and Fagales (see main text), the plastid genealogy is largely decoupled from speciation processes reflected in morphology (phenotypes) and nuclear-genetic differentiation patterns. In addition, speciation in beech has probably never been dichotomous but involved reticulate processes at several levels (**Fig. SM1-1**).

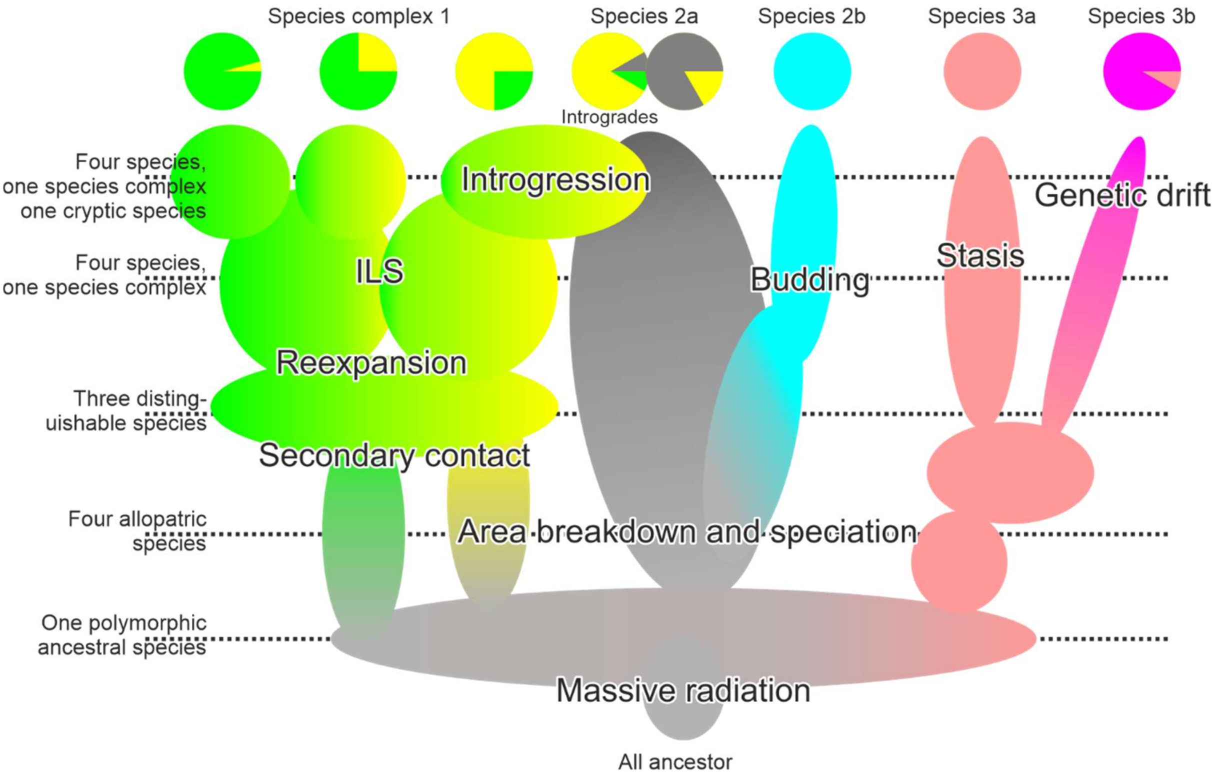


**Figure SM1-1 | Probable speciation and differentiation processes in beech, their effect on the plastid gene pools and identification of nuclear-morphologically distinct species.** Colours indicate the gene pools. Different genetic data sets will cover some (single-copy nuclear gene regions), particular (plastomes → allopatry/sympatry), or most aspects (low- and multi-copy nuclear gene regions) of these processes.

Thus, each phylogenetic tree inference will be incomprehensive on its own, possibly misleading (e.g. proposed species tree and coalescent in Jiang et al. 2022) and has to be carefully evaluated when being interpreted in an evolutionary context (see e.g. Cardoni et al. 2022). Reflecting only the maternal lineage and showing not sorted but high intra-species variability, no plastome-based phylogeny can provide a test to identify quasi-holophyletic species groups in beech (or any other Fagaceae or Fagales). However, being strongly controlled by geography (for details and references see **SupplR&D.pdf**, *section* 2), plastids can provide direct information about whether the common ancestors of e.g. sibling lineages – identified based on nuclear genealogies and differentiation patterns showing high taxonomic correlation – were sympatric (similar or identical plastid signatures) or allopatric (distinct plastid signatures). Thus, they can reflect probable allopatric speciation events predating the manifestation of the modern-day species and species groups and can point towards past reticulation events such as 'ghost' hybridisation and introgression, commonly addressed as "chloroplast capture" in phylogenetic literature although there is no horizontal gene and organelle transfer involved at the cellular level (see e.g. Thyssen et al. 2012, Hertle et al. 2021).

The general correlation of nuclear-genetic signatures with phenotypes, the latter being still main basis for angiosperm taxonomy, allows straightforward identification of empirically defined species (Mallet 1995, 2008) including two pairs of cryptic, i.e. phenotypically indistinguishable, species in East Asia – widespread Chinese *F. engleriana* Seemen ex Diels + narrow-endemic *F. multinervis* Nakai (Ulleungdo Is., Korea); *F. hayatae* Palib. (s.str.) endemic to Taiwan and Central Chinese *F. pashanica* C.C.Yang – and several pseudocryptic, i.e. characterised by morphological clines and so far overlooked morphological traits, species in western Eurasia: (*i*) the chiefly European *F. sylvatica* L. (s.str.), (*ii*) the south-eastern European and north-western Anatolian *F. orientalis* Lipsky (s.str.), (*iii*) *F. hohenackeriana* Palib. (Pontian forests; Caucasus, N. E. Turkey) and (*iv*) the Hyrcanian (N. Iran, S. E. Azerbaidshan) *F. caspica* Denk & G.W.Grimm (Denk et al. 2024). **Table SM1-1** gives the current data coverage situation for, the here recognised, species of *Fagus.*

**Table SM1-1 | Species coverage in this study and preceding molecular-phylogenetic papers on beech.** Numbers give the number of included viz new/gene bank (this study) individuals. D'02/05—Denk et al. (2002, 2005); G'07—Grimm et al. (2007); G&G'08—Göker & Grimm (2008); R'16—Renner et al. (2016); O'16—Oh et al. (2016); J'21—Jiang et al. (2022); C'22—Cardoni et al. (2022; includes an in-depth re-analysis of the data collected by Jiang et al. 2022).

| Subgenus | Species | D'02/05 G'07 G&G'08 | R'16,O'16 | J'21 | C'22 | This study |
|---|---|---|---|---|---|---|
| *Englerianae* | *F. engleriana* | 4 | Yes | 5 | No | 1/1 |
| | *F. japonica* Maxim. | 2 | Yes | 2 | Yes | 15/2 |
| | *F. multinervis* | 1 | Yes | 2 | No | 3/2 |
| *Fagus* | *F. crenata* Blume | 2 | Yes | 3 | Yes | 13/2 |
| | *F. hayatae* | No | No | 2 | No | 2/0 |
| | *F. pashanica* | 3 | Yes | 2 | No | No |
| | *F. longipetiolata* Seemen | 4 | Yes | 4 | No | 1/0 |
| | *F. lucida* Rehder & E.H.Wilson | 2 | Yes | 4 | No | 1/0 |
| | *F. sylvatica* | 4[b] | Yes | 3 | Yes | 6[b]/1 |
| | *F. orientalis*[a] | 2 | Yes | 1 | No | No |
| | *F. hohenackeriana*[a] | 1 | No | No | No | No |
| | *F. caspica*[a] | No | No | No | Yes | 2/0 |
| | *F. grandifolia* | 3[c] | Yes | 4 | No | 4/0 |
| | *F. mexicana* | 1 | No | 3 | No | No |
| | **Data** | ITS (+morph) | (ITS+) LFYi2 | 28 nuclear genes | 5S-IGS | *CRC*+LFYi2 plastomes |

[a] The oriental beeches, traditionally treated as a single species or subspecies of *F. sylvatica* include three morphologically and genetically diagnosable species (Denk et al. 2024): the Western *F. orientalis* s.str.; the central populations of northeastern Turkey and the Caucasus region, *F. hohenackeriana* (see also Shen 1992); while the easternmost populations in the Talysh and Alborz Mts represent a new species, *F. caspica* (formalized in Denk et al. 2024).
[b] Including *F. moesiaca* (K.Malý) Czeczott (ecotype within *F. sylvatica*; cf. Denk 1999a, Denk et al. 2024)
[c] Including an individual of south-eastern "swamp" beech, *F. grandifolia* subsp./var. *caroliniana*.

### 2 Assessment of phylogenetic data of *Fagus* available in gene banks

#### 2.1 *Premise*

To be able to cross-check the newly assembled data, we undertook a complete NCBI GenBank harvest on February 2nd 2021, processed with GBK2FAS (Göker et al. 2009) to extract meta-information linked to each gene bank accession and identify the best- and broadest-sampled DNA regions.

GBK2FAS is available for download at: http://www.goeker.org/mg/clustering/

#### 2.2 Data available

The (full) GenBank flatfile (**All17022021.gb**) includes all genomic DNA/RNA accessions sorted by organism for genus beech. Search string is:

```
txid21024[Organism:exp]
```

Our standard extraction command line was used to read out all meta-information relevant to our study.

```
gbk2fas -l oavtcdbjmsnp -s "|" -t -x -z All17022021.gb > All17022021.csv
```

o – organism name; -x is a correction to clean organism names, i.e. names not using Linnéan binominals.

a – accession number

v – voucher

t – taxonomy tree; not needed in our case (all accessions come from the same genus)

c – geographic provenance (country, state, province, region), especially important when harvesting plastid data (often showing strong geographic filtering in plants, especially wide-spread species)

d – definition (title) line, gene banks include many poorly annotated accessions, where information about sequenced gene regions can only be retrieved from the "note" field or the definition line

b – (first line) of author list

j – (first line) of publication reference: journal, year, pages

m – molecule type; not needed here (pre-filtered for genomic DNA/RNA) other than to eliminate unassigned (patent-related) DNA.

s – source, which is either the organism (species) itself, or its chloroplast or mitochondrion; used to pre-filter the data (note: this is an originally non-curated flatfile field, and may be wrong)

n – sequence length; useful to filter for oligogene accessions (e.g. completely sequenced *trnK/matK,* inverted repeat (plastome) or 35S rDNA and ITS regions (nucleome)

p – product as provided in the flatfile; in conjunction with -z and -s "|", each product will be listed individually in its own column

-t reads out selected (with -l) meta-information into a tab-separated file; -s defines the separator used for the output (here: "|" to facilitate import in Excel®)

#### 2.3 Coverage

The best-sampled genomic region is the ITS region (292 accessions [accs]), comprising the internal transcribed spacer ITS1 and ITS2, the interspersing 5.8S rRNA gene, and end and start of the flanking 18S rRNA (or *SSU* – small subunit) and 25S rRNA (28S; or *LSU* – large subunit) genes of the 35S (45S) rDNA cistron. While providing sufficient discriminate signal to identify species and major species complexes, its multi-copy, potentially multi-loci (Ribeiro et al. 2011) organisation require extensive cloning procedures and makes it very difficult to analyse (Denk et al. 2002, 2005, Grimm et al. 2007).

The only other nuclear-encoded gene region covered for all main species is the *LEAFY* gene, specifically its second intron (LFYi2; 77 accs).

In addition to, at the time of the harvest, 10 completely sequence plastomes (including three reference plastomes for *F. crenata, F. engleriana*, and *F. sylvatica*, good taxonomic coverage is available for five plastid genomic regions, commonly advocated as 'barcodes': (*i*) *rbcL* gene (63 accs); (*ii*) *trnK/matK* region including the *matK* gene and *trnK* intron sequences (total of 142 accs); (*iii*) the *atpB-rbcL* (64 accs) and (*iv*) *trnH-psbA* (42 accs) intergenic spacers (IGS), and (*v*) data covering parts of the trnTLF region, mostly *trnL* intron and *trnL-trnF* IGS, some *trnT-trnL* IGS data (total of 218 accs).

#### 2.4 Harvest and pre-processing of selected sequence data

The following search strings were used to harvest regions with good taxonomic coverage:

txid21024[Organism:exp] AND …

**LFYi2**— … (LEAFY [title] OR LFY [title]); capturing all 77 accessions

***rbcL*** gene—… (rbcL [gene] OR ribulose) NOT matK [gene]; capturing 64 accessions (−1), excluding complete plastome sequences

**trnK/matK** region—… (matK [gene] OR maturase [title] OR trnK [title]) NOT (rbcL [gene] OR matR [gene]); returning all 142 accessions, excluding complete plastome sequences

***trnH-psbA*** spacer—… (trnH-psbA [title] or psbA-trnH [title]); returning all 42 accessions

**trnTLF** region—… txid21024[Organism:exp] AND (trnL OR trnL-trnF OR trnL-F OR tRNA-Leu OR trnT-trnL OR trnT-L) NOT (ndhJ OR rpl32); returning 217 accessions (missing 1)

Searches were downloaded as GenBank flatfiles (*.gb) and transferred into FASTA-formatted files using GBK2FAS with the following batch file:

```
gbk2fas -l oavc -r 4 -s "|" -x 1_tHpA.gb > 1_tHpA.fasta
gbk2fas -l oavc -r 4 -s "|" -x 2_tKmK.gb > 2_tKmK.fasta
gbk2fas -l oavc -r 4 -s "|" -x 3_trnTLF.gb > 3_trnTLF.fasta
gbk2fas -l oavc -r 4 -s "|" -x 4_aBrL.gb > 4_aBrL.fasta
gbk2fas -l oavc -r 4 -s "|" -x 5_rbcL.gb > 5_rbcL.fasta
gbk2fas -l oavc -r 4 -s "|" -x 6_LFYi2.gb > 6_LFYi2.gb.fasta
```

Option -r 4 replaces all non-letter, non-digit characters in the read-out field by underlines.

Broadly sampled plastid regions were extracted from the complete plastomes using the site filtering option of the NCBI GenBank portal (button “Change region shown”), and individually downloaded the sequences in FASTA format. Final files are included in the **Online Data Archive (ODA)**.

#### *2.5 Alignment*

All harvested data were aligned using MAFFT v. 2.273 (Katoh & Standley 2013), and visually inspected using MESQUITE’s bird’s eye view (v. 2.75; Maddison & Maddison 2011). In case of the plastid regions, the complete plastome extracts were added to the FASTA-files before alignment; and added at the start of the files. This, in conjunction with MAFFT’s option --adjustdirection, ensures orientation of sequences as found in the plastome ring and facilitates annotation as well as alignment of non-overlapping fraction-data blocks (such as *trnT-trnL* IGS*, trnL* intron, and *trnL-trnF* IGS accessions). Choice of alignment algorithm was left to MAFFT (**Table SM2-1**) for the plastid regions.

Annotation of gene regions in the plastid alignments follows the *F. sylvatica* reference plastome NC_041437 (≡ MK598696). L-INS-i, being the slowest but also most accurate algorithm, was used to pre-align the harvested LFYi2 data.

**Table SM2-1. Search strategies applied by MAFFT for data harvested from gene banks**

| Dataset | General structure | Chosen alignment algorithm |
|---|---|---|
| *trnH-psbA* | Length-polymorphic and relatively divergent | L-INS-i |
| trnK/matK | Generally length-conserved | FFT-NS-i |
| trnTLF | Relatively conserved, with huge gaps in overlap | FFT-NS-i |
| *atpB-rbcL* | Highly conserved | L-INS-i |
| *rbcL* | Near invariable | L-INS-i |

#### 2.6 Curation of auto-aligned data

The harvest includes partly (very) old accessions prone to sequencing/editing errors (such as singleton 1-nt indels; AATTT instead of AAATT in all other accessions), particularly at the start and end of sequences overlap areas of forward and reverse primers (most non-phylogenomic data in gene banks are based on directly sequenced PCR products). Hence, we trimmed sequence ends; apparent errors and parts with potentially problematic mutation patterns were blanked out (replaced by "?"). In obvious cases such as the AT example previously described (indicated by lowercase letters), mononucleotide repeat numbers were corrected for. Length-polymorphic regions (LPRs) with mononucleotide repeats were left-/right-/centre-aligned to minimise gaps and in cases where the auto-alignment was inconsistent, i.e. same sequence motives aligned differently. As a rule, modification of the alignments were kept to a minimum. Curated alignments are provided as NEXUS format, the original unmodified auto-alignments (MAFFT output) as FASTA files, in the **ODA**.

#### 2.7 Quick-and-dirty guide trees

For nuclear target regions (*CRC,* LFYi2), we ran a full maximum likelihood tree inference and non-parametric bootstrap analysis (Felsenstein 1985) using a stand-alone version of RAxML v. 8.2.10 (Stamatakis 2014) provided as Windows®-executable. RAxML v.8.2.10 used the per-site rate approximation (-GTRCAT; Stamatakis 2006) for fast bootstrapping and topology inference with a final optimisation step under the GTR + Γ substitution model. The number of necessary bootstrap replicates was determined during each run using the extended majority rule (MRE) bootstop criterion (Pattengale et al. 2009). In case of substantial length polymorphism (*CRC,* LFYi2), we run two sets of analysis, one including and one excluding length-polymorphic regions (LPR). In case of the complete tip-set and tip-reduced plastome data (20 individuals covered for both nuclear and plastid data), we performed a RAxML tree inference with 100 BS pseudoreplicates under the GTR+Γ (Rodriguez et al. 1990) and GTR+Γ+I models (-m GTRGAMMA/ GTRGAMMAI) and using three different partitioning schemes: (*i*) unpartitioned; (*ii*) fully partitioned with 389 partitions, whereby each separate intergenic spacer, intron, tRNA region, rRNA region and protein-coding gene (split into 1st, 2nd and 3rd codon positions) was defined as a separate partition; and (*iii*) a "logical" partitioning scheme using six partitions informed by function and mutability (→ **Section 4.1**).

The batch files to re-run analyses are included in the **ODA**. Resulting trees were viewed in DENDROSCOPE3 (Huson & Scornavacca 2012) for initial annotation, exported as (vector-/object-based) PDF or scalable vector graphic (SVG) and graphically enhanced using ADOBE® ILLUSTRATOR and CORELDRAW®. DENDROSCOPE-generated NEXML files are included in the **ODA** as well.

#### 2.8 Comparison with novel nuclear data

During the course of our study of complete plastomes, we generated new *CRC* and LFYi2 sequence data for selected individuals to verify their taxonomic affinity and to assess their fit with the phylogenetic framework of species of *Fagus* (as depicted in ***main-text fig. 1***; **SupplR&D.pdf**, *fig. SR1-1*). Both *CRC* and LFYi2 could be expected to be more divergent than the low-copy nuclear markers assembled by Jiang et al. (2021) and, hence, to add discriminative power to the available nuclear data at and above the species level.

##### 2.8.1 PCR and cloning protocol

For *CRC* and LFYi2 primers iF2 and iR2 amplifications were done in 10µl PCR mixtures contained 0.8µl 2µM F and R primers, 1µl 10xExTaq buffer, 2 µl TBT-PAR, 0.05 µl ExTaq (Takara Bio Inc. Japan) and 1 µl genomic DNA. For LFYi2 primers iF2 and iR4 a Multiplex PCR Assay Kit Ver.2 (Takara) was used following the manufacturers' instructions. The PCR thermocycle for *CRC* followed Oh & Manos (2016) while the PCR thermocycle for LFYi2 iF2 and iR2 primers was as follows: an initial denaturation at 98 °C for 3 min, followed by 10 cycles of 98 °C for 30 s, decreasing the annealing temperature from 60 °C to 55 °C at 0.5 °C/cycle and 60 °C for 30 s, followed by 30 cycles of 98 °C for 30 s, 55 °C for 30 s and 72 °C for 1 min and lastly a final extension at 72 °C for 10 min. For LFYi2 primers iF2 and iR4 the PCR thermocycle consisted of an initial denaturation at 94 °C for 30 s, followed by 40 cycles of 94 °C for 30 s, 60 °C for 15 s, 72 °C for 1 min and finally a 10 min extension at 72 °C. The

TA-cloning method followed Oh et al. (2016) for both *CRC* and LFYi2. Between two and six clones of each sample were sequenced using a BigDye Terminator v1.1 Cycle Sequencing Kit (Thermo Fisher Scientific) with M13R and T7 primers and separated by capillary electrophoresis on an ABI3130 Genetic Analyzer (Life Technologies, Waltham, MA, USA). Due to the long length of *CRC* and the presence of two internal SSR regions three internal primers were used for sequencing in addition to the M13R and T7 primers (**Table SM2-2**). Sequences were checked by eye in GENEIOUS 9.1.5 (Biomatters, Auckland, New Zealand).

**Table SM2-2 | Primer pairs used to amplify nuclear *CRC* and LFYi2 regions.** Due to the long length of the *CRC* region and long SSR regions three internal primer pairs were used for DNA sequencing (Fagus CRC6R-1, Fagus CRC5R-1 and crc3m).

| | Sequence | Reference |
|---|---|---|
| **LFY TA-cloning** | | |
| LEAFY_i_F2 | GTCCAAAACATTGCCAAAGAACGC | This study |
| LEAFY_i_R2 | TTGGCAAACCTAAACACTTGG | This study |
| LEAFY_i_R4 | CACTTGGTTTGTCACCTGTATAC | This study |
| **CRC TA-cloning** | | |
| Fagcrc1 | CATCTTTGYTAYGTCCGYTGCARYTTCTG | Oh & Manos 2008 |
| Fagcrc2 | GGDATGTACYKAGCCCACTATTGCTCAAAAG | Oh & Manos 2008 |
| crc3m | GCGGTTGCTGGACACTGTGAC | Oh & Manos 2008 |
| **LFY Sequencing** | | |
| M13R 5' | CAGGAAACAGCTATGAC | Oh & Manos 2008 |
| T7_5' | GTAATACGACTCACTATAGGG | Oh & Manos 2008 |
| **CRC Sequencing** | | |
| M13R 5' | CAGGAAACAGCTATGAC | Oh & Manos 2008 |
| T7_5' | GTAATACGACTCACTATAGGG | Oh & Manos 2008 |
| Fagus CRC6R-1 | GTTGGCAGCTTTGATACGC | This study |
| Fagus CRC5R-1 | GGGTTCGCTAGATGTTGATG | This study |
| crc3m | GCGGTTGCTGGACACTGTGAC | Oh & Manos 2008 |

##### *2.8.2* CRC data

At the time of the harvest, GenBank included two (direct-PCR) accessions, generated for an all-Fagaceae phylogeny (Oh & Manos 2008). They represent the North American species *F. grandifolia* and *F. mexicana*, a taxon endemic to summer-wet/per-humid montane regions in (north-)eastern Mexico traditionally treated as a subspecies of *F. grandifolia*. The two accessions mainly differ by indels, and three point mutations. We added five cloned sequences (acc. nos OM322853–OM322857) from two *F. grandifolia* individuals, one from Quebec, Canada (#38), the other from Pennsylvania, U.S. (#40). Interestingly, the Pennsylvanian individual matches the sequence of Oh & Manos' (2006) directly sequenced *F. grandifolia*, while the Canadian clones are nearly identical to their Mexican sample (directly sequenced

*F. mexicana*), distinguished by a 6 nt-long insert in the first LPR (TATAGAGA microsatellite).[1]

The newly assembled cloned data (acc. no. OM322828–OM322928; annotated list can be found in **SupplGenetics_ncData.xlsx**, sheet *New LFYi2 and CRC accs*) covers eight more species. No data are available for the Western Eurasian species *F. orientalis* and *F. hohenackeriana* and the Chinese cryptic sister species of *F. hayatae, F. pashanica*.

##### 2.8.3 LFYi2 data

At time of harvest, gene banks included two sets of *LEAFY* intron 2 (LFYi2) sequences by Renner et al. (2016; generated using primers designed for plane trees [*Platanus*, Platanaceae, Proteales]) used for FBD dating of the genus and by Oh, Youm et al. (2016) with a focus on *F. japonica* and *F. multinervis* (the latter uploaded as *F. japonica* var. *multinervis*). Together with the newly generated accessions, a total of 239 LFYi2 sequences are available for all but one species of *Fagus* (according Denk et al. 2024), including Chinese (*F. pashanica*; 1 acc.) and Taiwanese *F. hayatae* s.l. (1 individual, 6 new accs; *F. hayatae* s.str.), representatives of *F. orientalis* (s.str.; Bolu, Turkey, 1 acc.) and *F. caspica* (N. Iran, 1 ind., 6 new accs), and *F.* (×) *moesiaca* specimen cultivated in the Arnold Arboretum of unknown native provenance; local taxonomists have suggested *F.* (×) *moesiaca* might be a hybrid between *F. sylvatica* (s.str.) and *F. orientalis* (but see Denk 1999a, Gömöry et al. 1999, Gömöry and Paule 2010). The only species not covered by LFYi2 data is *F. hohenackeriana*, the Pontic-Caucasian beech. To increase PCR success, new primer pairs were developed (**Table SM2-2**), amplifying nearly the complete putative intron (except for nine invariable bp at the 5' end; see file **full.nex* in ODA). See **SupplGenetics_ncDNA.xlsx**, sheet *New LFYi2 and CRC accs*, for GenBank accession numbers (OM322929– OM323089) and meta-information on newly sequenced cloned data.

[1] This is not as curious as it seems: the Mexican population may be a relict of the first wave of modern *Fagus grandifolia* migrating south, and the Canadian a left-over, both being not yet homogenised by the (sequentially) more derived, dominant eastern U.S. populations representing the most recent radiation. To test this hypothesis, the plastome of the Mexican populations would need to be studied. Population-scale 5S-IGS data (cf. Cardoni et al. 2022; **SupplR&D.pdf**, *fig. SR1-10*) could be beneficial to find further evidence for ongoing (cryptic or pseudocryptic) speciation.

#### 3 Assembly of complete chloroplast genome data

Forty-eight new *Fagus* chloroplast genomes were assembled including 28 from Japan consisting of (*i*) 13 samples of *F. crenata* (*F.* subg. *Fagus*), of which Sanger-based haplotypes for 11 samples have previously been determined (Fujii et al. 2002), and (*ii*) 15 samples of *F. japonica* (*F.* subg. *Englerianae*) of which the Sanger-based chloroplast haplotype have not been determined (**Table SM3-1**; **SupplGenetics_cpDNA.xlsx**, sheet *New and used plastomes*). The whole chloroplast genomes of other species distributed outside Japan were also obtained including (*i*) 14 individuals of *Fagus* subgenus *Fagus*: four *F. grandfolia*, six *F. sylvatica* (incl. one *F.* [×] *moesiaca*), two *F. hayatae* and one each of *F. lucida* and *F. longipetiolata*; and (*ii*) four additional individuals of *F.* subgenus *Englerianae:* three *F. multinervis* and one *F. engleriana*. All samples were from natural beech forests or botanical gardens with known geographic origin except for the *F. lucida* and *F. longipetiolata* individual.

**Table SM3-1 | Source information for all 82 whole chloroplast genomes of *Fagus*** used including 48 samples newly obtained for this study (in bold) and 34 acquired from GenBank.

| Species | Individual | GenBank acc. nos | Provenance | Coordinates |
|---|---|---|---|---|
| *crenata* | cr01\|Chubu | **OR666470** | Mt Amagi, Chubu, Japan | 34.87° N, 138.98° E |
| *crenata* | cr02\|Chubu | **OR807735** | Mt Hakusan, Chubu, Japan | 36.17° N, 136.77° E |
| *crenata* | cr03\|Kyushu | **OR818326** | Mt Seburi, Kyushu, Japan | 33.44° N, 130.37° E |
| *crenata* | cr04\|Kyushu | **OR818324** | Mt Sobo, Kyushu, Japan | 32.83° N, 131.33° E |
| *crenata* | cr05\|Kyushu | **OR818325** | Mt Takakuma, Kyushu, Japan | 31.48° N, 130.8° E |
| *crenata* | cr06\|Kanto | **OR807733** | Hatomachi Pass, Kanto, Japan | 36.88° N, 139.22° E |
| *crenata* | cr07\|Tohoku | **OR807729** | Mt Hayachine, Tohoku, Japan | 39.42° N, 141.5° E |
| *crenata* | cr08\|Tohoku | **OR807731** | Mt Iide, Tohoku, Japan | 37.92° N, 139.68° E |
| *crenata* | cr09\|Shikoku | **OR818328** | Mt Tsurugi, Shikoku, Japan | 33.85° N, 134.1° E |
| *crenata* | cr10\|Shikoku | **OR807734** | Mt Ishizuchi, Shikoku, Japan | 33.77° N, 133.15° E |
| *crenata* | cr11\|Chugoku | **OR818327** | Mt Jyakuchi, Chugoku, Japan | 34.47° N, 132.05° E |
| *crenata* | cr12\|Hokkaido | MH171101 | Daisengendake, Hokkaido, Japan | 41.62° N, 140.13° E |
| *crenata* | cr13\|Tohoku | **OR807730** | Sado Island, skyline, Tohoku, Japan | 38.08° N, 138.32° E |
| *crenata* | cr14\|Kanto | **OR807732** | Hakone, Lake Ashinoko, Kanto, Japan | 35.23° N, 139° E |
| *crenata* | crGB\|MT762292 | MT762292 | Sasari Touge Pass, Kyoto, Japan | 35.28° N, 135.72° E |
| *multinervis* | mu15\|Ulleungdo | **OR826425** | Taeharyong, Ulleungdo, South Korea | 37.49° N, 130.84° E |
| *multinervis* | mu16\|Ulleungdo | **OR826424** | Naesujeon, Ulleungdo, South Korea | 37.51° N, 130.91° E |
| *multinervis* | mu17\|Ulleungdo | **OR826423** | Bongnae Falls, Ulleungdo, South Korea | 37.5° N, 130.89° E |
| *multinervis* | mu18\|Ulleungdo | MK518070 | Bongnae Falls, Ulleungdo, South Korea | 37.5° N, 130.89° E |
| *multinervis* | muGB\|MN894556 | MN894556 | Dodong, Ulleungdo, South Korea | 37.49° N, 130.89° E |
| *multinervis* | muGB\|MN516696 | MN516696 | Unknown, Ulleungdo, South Korea | [no data] |
| *multinervis* | muGB\|MT762296 | MT762296 | Buk-myeon, Ulleungdo, South Korea | 37.52° N, 130.87° E |
| *japonica* | ja19\|Nagano | **OR972976** | Nagiso Dake, Nagano, Japan | 35.58° N, 137.65° E |
| *japonica* | ja20\|Nagano | **OR972975** | Atebi Daira, Nagano, Japan | 35.23° N, 137.67° E |
| *japonica* | ja21\|Tohoku | **OR972962** | Ichinohe, Tohoku, Japan | 40.2° N, 141.29° E |

| **Species (ctd)** | **Individual** | **GenBank acc. nos** | **Provenance** | **Coordinates** |
|---|---|---|---|---|
| *japonica* | ja22\|Tohoku | **OR972963** | MtGoyo, Tohoku, Japan | 39.21° N, 141.76° E |
| *japonica* | ja23\|Tohoku | **OR972964** | MtTani-yama, Tohoku, Japan | 38.14° N, 140.68° E |
| *japonica* | ja24\|Tohoku | **OR972965** | Ogawa, Tohoku, Japan | 36.9° N, 140.63° E |
| *japonica* | ja25\|Kanto | **OR972966** | Mt Takao, Kanto, Japan | 35.62° N, 139.24° E |
| *japonica* | ja26\|Kanto | **OR972967** | Kuni, Kanto, Japan | 36.59° N, 138.64° E |
| *japonica* | ja27\|Kansai | **OR972968** | Ohto, Kansai, Japan | 34.15° N, 135.82° E |
| *japonica* | ja28\|Kansai | **OR972969** | Onzui, Kansai, Japan | 35.24° N, 134.55° E |
| *japonica* | ja29\|Shikoku | **OR972970** | Mt Tsurugi, Shikoku, Japan | 33.89° N, 134.05° E |
| *japonica* | ja30\|Chugoku | **OR972971** | Funo, Chugoku, Japan | 34.86° N, 132.78° E |
| *japonica* | ja31\|Chugoku | **OR972972** | Mt Iigatake, Chugoku, Japan | 34.33° N, 131.74° E |
| *japonica* | ja32\|Kyushu | **OR972973** | Mt Sobo, Kyushu, Japan | 32.84° N, 131.34° E |
| *japonica* | ja33\|Kyushu | **OR972974** | Siiba, Kyushu, Japan | 32.44° N, 131.15° E |
| *japonica* | jaGB\|MT762294 | MT762294 | Sasari Touge Pass, Kyoto, Japan | 35.28° N, 135.72° E |
| *japonica* | jaGB\|MT762295 | MT762295 | Sasari Touge Pass, Kyoto, Japan | 35.28° N, 135.72° E |
| *engleriana* | en34\|Anhui | **OR826426** | Huangshan, Anhui, China | 30.13° N, 118.17° E |
| *engleriana* | enGB\|KX852398 | KX852398 | China, origin indet.; from Wuhan Bot. G. | [no data] |
| *engleriana* | enGB\|MT762293 | MT762293 | China, provenance unclear | [no data] |
| *pashanica* | paGB\|MW846258 | MW846258 | Changtan River, Sichuan, China | 32.67° N, 106.56° E |
| *longipetiolata* | lo36\|China | **OR826427** | China, origin indet.; from Hanzhou Bot. G. | [no data] |
| *longipetiolata* | loGB\|MZ562567 | MZ562567 | China, provenance unclear | [no data] |
| *lucida* | lu37\|China | **OR826428** | China, origin indet.; from Arnold Arboretum | [no data] |
| *lucida* | luGB\|MZ463069 | MZ463069 | cultivated, China, provenance unclear | [no data] |
| *lucida* | luGB\|NC_061574 | NC_061574 | Hengshan, Hunan, China | [no data] |
| *grandifolia* | gr38\|Quebec | **OR900458** | Saint-Hippolyte, Quebec, Canada | 45.99° N, 74° W |
| *grandifolia* | gr39\|Texas | **OR826429** | Martin Dies Jr State Park, Texas, USA | 30.85° N, 94.17° W |
| *grandifolia* | gr40\|Pennsylvania | **OR900459** | Forbes State Forest, Pennsylvania, USA | 40.13° N, 79.17° W |
| *grandifolia* | gr41\|Michigan | **OR900457** | Tahquamenon Falls St. Pk, Michigan, USA | 46.61° N, 85.2° W |
| *hayatae* | ha43\|Southern E. Asia | **OR901330** | Tong Mt, N. Taiwan | 24.51° N, 121.65° E |
| *hayatae* | ha44\|Southern E. Asia | **OR901331** | Tong Mt, N. Taiwan | 24.51° N, 121.65° E |
| *sylvatica* | sy45\|NC Spain | **OR936131** | San Martín de la Virgen del Moncayo, N. C. Spain | 41.8° N, 1.82° W |
| *sylvatica* | sy46\|NE Italy | **OR936130** | Timau, N. E. Italy | 46.36° N, 12.98° E |
| *sylvatica* | sy47\|CW Italy | **OR936133** | Oriolo Romano, C. W. Italy | 42.16° N, 12.18° E |
| *sylvatica* | sy48\|NW Greece | **OR936132** | Voras, N. W. Greece | 40.9° N, 21.88° E |
| *sylvatica* | sy49\|NW Greece | **OR936129** | Paiko, N. W. Greece | 41.06° N, 22.29° E |
| *sylvatica* | sy50\|NE Germany | MK598696 | Gransee/Brandenberg, N. E. Germany | 53° N, 13.14° E |
| *sylvatica* | syGB\|MW531753 | MW531753 | Kellerwald-Edersee Nat. Park, C. Germany | 51.17° N, 8.96° E |
| *sylvatica* | syGB\|MW537046 | MW537046 | Jamy, N. Poland | 53.59° N, 18.94° E |
| *sylvatica* | syGB\|MW566769 | MW566769 | Gdánsk, N. Poland | 54.38° N, 18.52° E |
| *sylvatica* | syGB\|MW566770 | MW566770 | Glorup, Denmark | 55.18° N, 10.68° E |
| *sylvatica* | syGB\|MW566771 | MW566771 | Fôret des Colettes, C. France | 46.18° N, 2.95° E |
| *sylvatica* | syGB\|MW566772 | MW566772 | Limitaciones, N. Spain | 42.82° N, 2.25° W |
| *sylvatica* | syGB\|MW566773 | MW566773 | Bieszczady Nat. Park, S. E. Poland | 49.12° N, 22.58° E |
| *sylvatica* | syGB\|MW566774 | MW566774 | Łopuchówko, W. Poland | 52.58° N, 17.08° E |
| *sylvatica* | syGB\|MW566775 | MW566775 | Ehingen [in Ba.-Wü.], S. Germany | 48.4° N, 9.5° E |
| *sylvatica* | syGB\|MW566776 | MW566776 | Hasbruch, NW Germany | 53.12° N, 8.43° E |
| *sylvatica* | syGB\|MW566777 | MW566777 | Český Krumlov, Czech Republic | 48.85° N, 14.25° E |

| Species (ctd) | Individual | GenBank acc. nos | Provenance | Coordinates |
|---|---|---|---|---|
| *sylvatica* | syGB\|MW566778 | MW566778 | Eisenach, C. Germany | 50.09° N, 10.11° E |
| *sylvatica* | syGB\|MW566779 | MW566779 | Brzeziny, C. Poland | 51.84° N, 19.6° E |
| *sylvatica* | syGB\|MW566780 | MW566780 | Fantanele, C. E. Romania | 46.42° N, 26.47° E |
| *sylvatica* | syGB\|MW566781 | MW566781 | Fläming, N. E. Germany | 52.13° N, 12.58° E |
| *sylvatica* | syGB\|MW566782 | MW566782 | Smolenice, Slovakia | 48.49° N, 17.37° E |
| *sylvatica* | syGB\|MW566783 | MW566783 | Northern Veneto, N. E. Italy | 46.13° N, 12.22° E |
| *sylvatica* | syGB\|MW846258 | MW566784 | Morbach, C. W. Germany | 50.74° N, 6.98° E |
| x *moesiaca* | xm51\|[SE Balkans] | **OR936134** | Prob. S. E. Balkans; from Arnold Arboretum | [no data] |
| *caspica* | or52\|N Iran | **OR900533** | Gorgan, N. Iran | 36.7° N, 54.1° E |
| *caspica* | or53\|N Iran | **OR900534** | Gorgan, N. Iran | 36.68° N, 54.1° E |

DNA extractions were done using a modified CTAB protocol (Doyle, 1990). DNA concentration and quality were assessed by agarose gel electrophoresis and a Qubit 2.0 fluorometer (Life Technologies). DNA was sent to the Beijing Genomic Institute where short-size Truseq DNA libraries were constructed and paired-end sequencing was performed on an Illumina HiSeq2000 Genome Analyser. The assemble of the whole chloroplast genomes was undertaken using default settings in GetOrganelle (Jin et al. 2020). Where samples did not produce a circular genome, assembly was achieved by mapping contigs to the *F. crenata* whole genome (GenBank accession MH171101.2) in Geneious v. 9.0.5.

### 4 Tree and network inferences

#### 4.1 Standard setting for maximum likelihood tree inferences

Phylogenetic trees based on the complete plastome dataset were inferred using maximum likelihood analysis with RAxML v.8.2.10 using the same basic settings as outlined above. For the whole chloroplast genome data, three partition schemes were used: (1) unpartitioned; (2) full-partitioned based on the annotation of *Fagus crenata* chloroplast genome with separate partitions for every intergenic spacer, intron, 1$^{st}$, 2$^{nd}$ and 3$^{rd}$ codon position of each protein-coding gene, tRNAs and rRNAs respectively; and (3) a "logical" partition analysis with five partitions: intergenic spacers (non-coding, neutrally evolving); introns (non-coding, typically low-divergent), 1$^{st}$ + 2$^{nd}$ codon positions (determining amino-acid, conserved); 3$^{rd}$ codon positions (high level of synonymous mutations); tRNA + rRNA genes (structurally constrained, highly conserved). The resulting best-known tree was then rooted in FigTree v1.4.3 (Rambaut

2014) at the node leading to the four *F. grandifolia* samples in case of the plastid data set (see also all-Fagaceae, all-Fagales plastome trees by Yang et al. 2001 and Zhou et al. 2022), and at the subgeneric split in case of the nuclear data sets (cf. Denk et al. 2005, Renner et al. 2016, Simeone et al. 2022, Jiang et al. 2022).

#### 4.2 Individual-based analyses

In the complete plastome data set, each individual is represented by a single, unique complete chloroplast genome. For downstream comparison of nuclear and plastid data, we hence needed to establish inter-individual relationships instead of inter-clone relationships. One quick method to infer phylogenetic relationships between individuals showing substantial intra-individual genotypic variation is the host-associate analysis framework by Göker & Grimm (2008). Individuals are treated as '*hosts*', and their relationships are inferred based on their clone assemblage, as the '*associates*'.

##### 4.2.1 Individual-consensus-based ML trees

The simplest method to transfer a character matrix of *associates* into a character matrix of *hosts* is to compute consensus sequences.[2] Here, we used G2CEF (Göker & Grimm 2008) to transfer the harvested and newly generated cloned data into strict (option -c) and modal (option -v) consensus sequences for each individual, either treating gaps as 5$^{th}$ base (option -g) or not. Gene bank singleton sequences (directly sequenced, not cloned) representing provenances not included in our data were kept by setting the minimum number of *associates* (clones) per *host* (individual) to 1 (option -m 1). Gene bank data of unknown origin was dropped using a dummy *host* ("not_used"), manually deleted from the alignments. The following batch generates individual consensus alignments in extended PHYLIP (*.epf) and (simple) NEXUS (*.nex) format.

[2] The raw nuclear data generated by Jiang et al. (2022) represents *per se* individual-level consensus sequences that masks inter-locus ("low-copy" gene regions) and allelic variation, expressed by polymorphic base calls: 2ISPs (intra-individual site polymorphism; Potts et al. 2014). Lacking a conceptual framework how to deal with 2ISPs, Jiang et al. (2022) decided to phase their data into two arbitrary haplotypes per individual (including obvious chimeras; see supplement to Cardoni et al. 2022), which were then randomly concatenated into two chimeric sequences to represent each individual in their phylogenetic analyses.

```
g2cef -c -m 1 a_CRC.epf a_assoc.txt > a_CRC.c.m1.epf
g2cef -c -g -m 1 a_CRC.epf a_assoc.txt > a_CRC.c.g.m1.epf
g2cef -v -m 1 a_CRC.epf a_assoc.txt > a_CRC.v.m1.epf
g2cef -v -g -m 1 a_CRC.epf a_assoc.txt > a_CRC.v.g.m1.epf

g2cef -s -m 1 a_CRC.epf a_assoc.txt > a_CRC.s.m1.epf
g2cef -c -m 1 a_CRC.epf a_assoc.txt -N > a_CRC.c.m1.nex
g2cef -c -g -m 1 -N a_CRC.epf a_assoc.txt -N > a_CRC.c.g.m1.nex
g2cef -v -m 1 a_CRC.epf a_assoc.txt -N > a_CRC.v.m1.nex
g2cef -v -g -m 1 a_CRC.epf a_assoc.txt -N > a_CRC.v.g.m1.nex

g2cef -c -m 1 b_LFY.epf b_assoc.txt > b_LFY.c.m1.epf
g2cef -c -g -m 1 b_LFY.epf b_assoc.txt > b_LFY.c.g.m1.epf
g2cef -v -m 1 b_LFY.epf b_assoc.txt > b_LFY.v.m1.epf
g2cef -v -g -m 1 b_LFY.epf b_assoc.txt > b_LFY.v.g.m1.epf
g2cef -s -m 1 b_LFY.epf b_assoc.txt > b_LFY.s.m1.epf

g2cef -c -m 1 b_LFY.epf b_assoc.txt -N > b_LFY.c.m1.nex
g2cef -c -g -m 1 -N b_LFY.epf b_assoc.txt -N > b_LFY.c.g.m1.nex
g2cef -v -m 1 b_LFY.epf b_assoc.txt -N > b_LFY.v.m1.nex
g2cef -v -g -m 1 b_LFY.epf b_assoc.txt -N > b_LFY.v.g.m1.nex
```

Individual-based ML trees were inferred from the consensus matrices using the standard 2ISP-semi-aware ("ML-A") 4 x 4 substitution model (cf. Potts et al. 2014).

G2CEF and PBC can be downloaded from: http://www.goeker.org/mg/distance/

##### 4.2.2 Individual-based planar (meta-)phylogenetic networks

Non-treelike evolution will lead to tree-incompatible signal patterns and inevitably triggers tree-branching artefacts. In such cases, planar (meta-)phylogenetic networks such as neighbour-net splits graphs can be more straightforward regarding identification of shared evolutionary trajectories and determination of phyly. For instance, an unambiguous holophylum will be reflected by a neighbourhood defined by a prominent edge bundle, a 'trunk'. A well-developed 'fan', a series of several partly overlapping neighbourhoods, typically reflects common origins (monophyly in the original sense) of various sorts; including (*i*) paraphyly in a strict sense (e.g. budding speciation events, left-over ancestral groups), (*ii*) epi- and periphyly related to multi-level evolutionary reticulation (Wheeler 2014), or (*iii*) holophyly masked by yet incomplete lineage sorting. For instance, populations that are in the process of speciation typically produce neighbour-net fans because of overlapping geographic and ecologic differentiation processes. In trees, they will surface as poorly resolved and supported subtrees. When the used data can *per se* reflect secondary gene flow, either by according intra-individual polymorphism or inter-gene topological conflict (Cardoni et al. 2022), hybridisation and introgression will inflict prominent 'boxes' in the graph: two overlapping neighbourhoods formed by prominent edge bundles (for various real-world examples, see Göker & Grimm 2008 and Potts et al. 2014).

Prominent boxes are not *sufficient* criteria for hybridisation: they can also be the product of (terminal) ancestor-descendant trichotomies. The last common ancestor of two sisters will be ± equally distant to both of them, thus, producing a prominent terminal box. The same hold for a hybrid between siblings showing both parental signatures.

Furthermore, neighbour-nets may give direct information about the primitiveness of tips. Less-evolved tips characterised by ancestral signatures will be placed close to the centre of the graph, while unique (isolated), strongly diverged ones will be most distant but can be directly connected to the graph's centre. A perfect common ancestor is equally similar to all its descendants (cf. Denk & Grimm 2009, fig. 1). On the other hand, if a neighbour-net is relatively tree-like, it shows that the signal in the data used to infer phylogenetic trees is trivial in the sense that the resultant clades are inevitable (being distance-based, neighbour-nets are more vulnerable to long-branch/ edge attraction than ML or Bayesian-inferred trees). In combination with traditional tree inference and branch support analyses, neighbour-nets provide a quick visual assessment of genetic coherence of taxonomic units. Especially also in the case of intra-individual genetic polymorphism.

Using PBC (Göker & Grimm 2008), a pairwise distance matrix of *associates* (here: mostly clones) can be transferred into a pairwise distance matrix of *hosts* (here: individuals; **SupplGenetics_ncDNA.xlsx**, sheet *HostAssoc*). We used two transformations: the "phylogenetic Bray Curtis" transformation (PBC, option -b; Göker & Grimm 2009) and the minimum (MIN) inter-clone distance transformation (option -i). While the former maintains information about intra-*host* variation, the latter focusses on least-distinct *associates* between *hosts*. Here: if two individuals share the same sequence variant, their MIN distance will be 0 irrespective of how different their other clones may be. A PBC distance of 0 on the other hand would mean their clone assemblages are identical, i.e. both individuals have exactly the same set of gene/ sequence variants.

The following batch generates MIN and PBC distance matrices in extended PHYLIP (for import in SPLITSTREE) and NEXUS format (option -N).[3]

```
pbc -b -m 1 a_CRC.unc.dst a_assoc.txt > a_CRC.b.m1.unc.dst
pbc -i -m 1 a_CRC.unc.dst a_assoc.txt > a_CRC.i.m1.unc.dst

pbc -b -m 1 a_CRC.unc.dst a_assoc.txt -N > a_CRC.b.m1.unc.dist
pbc -i -m 1 a_CRC.unc.dst a_assoc.txt -N > a_CRC.i.m1.unc.dist

pbc -b -m 1 b_LFY.unc.dst b_assoc.txt > b_LFY.b.m1.unc.dst
pbc -i -m 1 b_LFY.unc.dst b_assoc.txt > b_LFY.i.m1.unc.dst

pbc -b -m 1 b_LFY.unc.dst b_assoc.txt -N > b_LFY.b.m1.unc.dist
pbc -i -m 1 b_LFY.unc.dst b_assoc.txt -N > b_LFY.i.m1.unc.dist
```

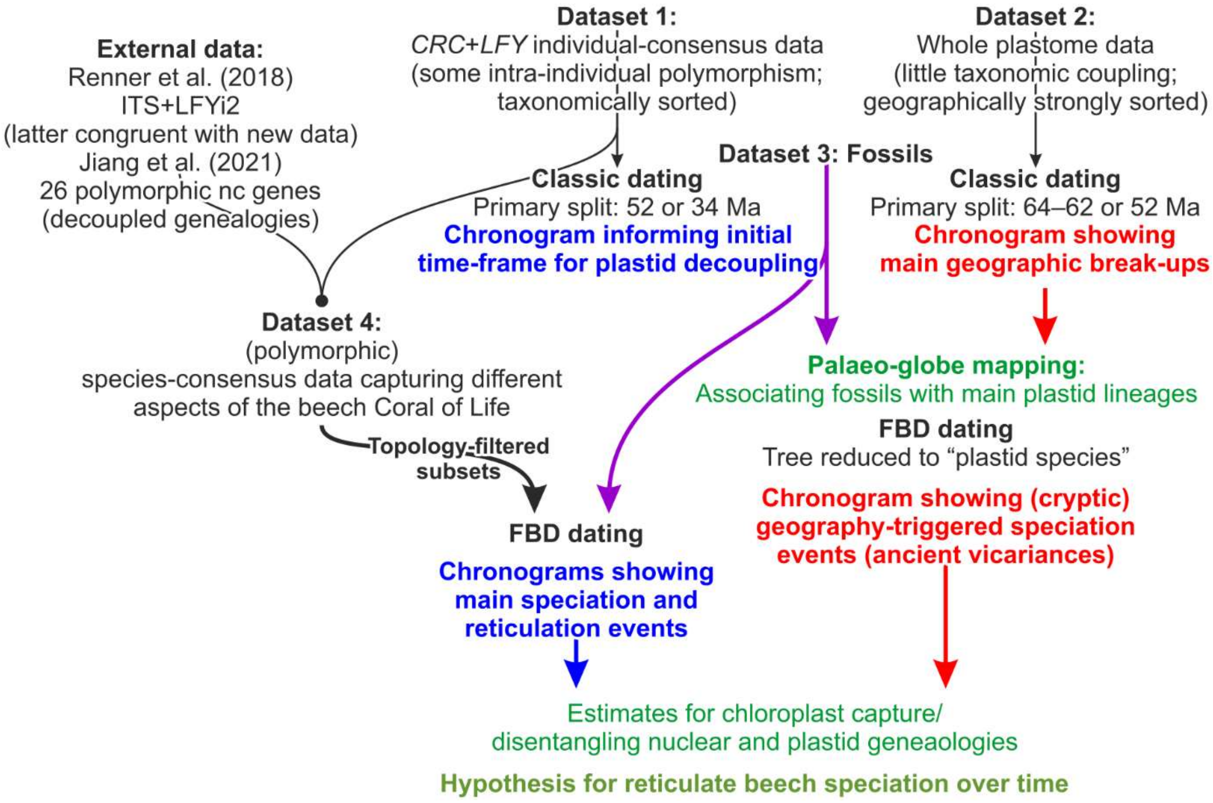


**Figure SM5-1 | Dating experiments flowchart**. Datasets and approaches in black; blue dated nuclear trees; red, dated plastid trees (see **SupplR&D.pfd**, *section 3*). Green, hypotheses and results.

[3] The NEXUS exports are generally compatible with NEXUS-using software (such as e.g. MESQUITE or PAUP*) with the exception of SPLITSTREE4, as it cannot handle certain code lines and additions such as commentary brackets in NEXUS files.

### 5 Molecular dating

We used the protocol shown in **Figure SM5-1** (preceding page) to infer species divergence times (based on nuclear data and topologies) and to correlate them to major phases of plastome divergence, i.e. identify divergence between 'plastid variants' lineages pre-dating or following nuclear-taxonomic speciation events. All dating analyses were performed with BEAST 2 (Bouckaert et al. 2014); according NEXML files are included in the ODA together with the BEAST-generated results files.

Together, the used topological scenarios approximate a comprehensive species network for beech (**Fig. SM5-2**) based on the following general assumptions:

1. **Nuclear data captures primary speciation events**—Nuclear divergences represent speciation events, i.e. the formation of new species that are reproductively isolated to a degree that they undergo subsequent homogenisation involving the accumulation of specific sequence patterns such as SNPs and diagnostic oligonucleotide motifs, and/or genotype stabilisation, the accumulation of private, specific polymorphism, because inbreeding outcompetes outbreeding.
2. **Nuclear data *also* captures secondary reticulation**—Secondary contact of species (hybridisation and introgression) will lead to genetic exchange of evolved, specific nuclear signatures as in the case of *F. longipetiolata* and *F. lucida* and the North American and Western Eurasian beeches. Corresponding divergence estimates (and clades in phylogenetic trees) represent the final isolation of two inter-mixing species. That is, the nuclear data divergences and similarity patterns may reflect both deep, primary and flat, secondary speciation events.
3. **Plastid data captures geographic sorting events—**These include allopatric speciation events in the course of major vicariance and dispersal processes (isolation of haplotypes) *and* intra-species differentiation within a widespread species, defined as a group of populations characterised by shared nuclear genotype(s) and phenotypes (haplotype gradients). Genetic drift in beech plastomes is primarily a function of distance, and only secondarily (Lineage III in *F. multinervis* vs. Lineage IV subtype of *F. hayatae*, both being narrow-endemics with small extant populations) of active population size. Thus, plastid divergences can only be established when the (primordial) mothers, the 'last common mothers' (LCMs), were already geographically isolated to a degree that their prodigy, daughter populations, do not overlap anymore. Two beech trees or maternal lineages growing at the same spot and sharing the same ancestral area, will – irrespectively of their

species – have similar plastomes. In addition, the closer a modern-day beech population (or species) is to the LCM, genetically and geographically, the less evolved will be its plastome.

Associating fossils with edges in a species network, here, represented by branches in the tested topological scenarios (→ **Section 5.1.2**), involves implicit assumptions also about the timing of reticulation. However, we cannot judge from the phenotype whether a fossil showing a clear affinity to a lineage represents a pre- or post-reticulation member of this lineage: leaf and pollen morphologies are generally limited in beeches and even modern-day putative hybrids are very difficult to identify because species that may form hybrid zones (the Western Eurasian species; cf. Kurz et al. 2022, Denk et al. 2024) typically also show an *intra*-species morphological gradient that, furthermore, can be observed in their precursors as well (Denk 1999a,b,c). Moreover, while the incongruence between plastid and nuclear genealogies, and between different nuclear gene markers, can only be explained by large-scale past introgression and hybridisation, all sympatric modern-day species are morphologically distinct from each other in mixed stands, hybrids such as *F.* (×) *okamotoi* Shen (*F. crenata* × *F. japonica*) are anecdotal (discussed in Denk et al. 2024). A fossil with strong affinities to a modern-day species may hence represent an early member of this species, unaffected by secondary gene flow but it also may represent a stabilised, post-reticulation population that already carries, e.g., the plastome of a distant cousin (from a modern-day perspective).

Thus, our dating experiments include an initial dating (**Section 5.1**) using a root age prior but no fossils as additional age priors to relatively gauge plastid and nuclear diversification based only on the signal in the used molecular data sets (three nuclear, one plastid data set). For the final dating experiments (**Section 5.2**), we rely on a combination of traditional node dating with a select and very conservative set of few fossils used as node age priors to establish minimum estimates for divergences and fossilised birth-death dating that can make use of the entirety of the fossil record to inform age distribution of lineages through time (Heath et al. 2014) for maximum divergence estimates.

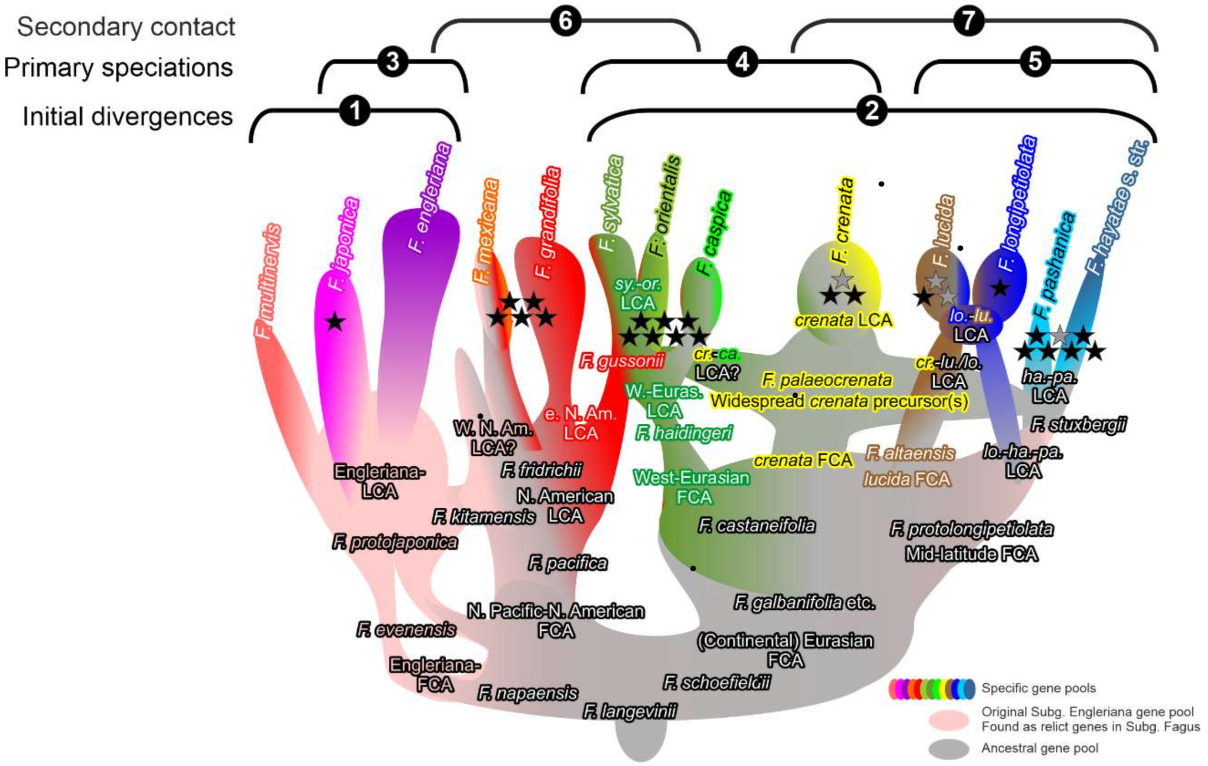


**Figure SM5-2 | Species network used as phylogenetic framework for the dating experiments.** After Cardoni et al. (2022), with fossil taxa added (following Denk & Grimm 2009, this study). The brackets show the topological constraints used on the nuclear data for the purpose of dating different aspects of beech evolution and link them to the according set of fossil species (→ Section 5.1.2): ❶ *Fagus* subgenus *Englerianae*; ❷ Eurasian clade within *F.* subgenus *Fagus*; ❸ *F. multinervis* as first-branching species lineage within *F.* subgenus *Englerianae*; ❹ High-latitude clade ❺ Mid-latitude (or Southern) clade; ❻ 'Euamerican' clade; ❼ East Asian-subgenus-*Fagus* clade. Stars give the number of fossils assigned to each terminal clade (species-group or species-lineage); grey: fossil taxa from the Kraskino palaeoflora (excluded in the 2nd batch of FBD dating experiments; FBD* in **SupplDating.xlsx**, sheet *final dating*). Abbr.: FCA = (putative) first (known) common ancestor; LCA = last common ancestor.

We used and combined four principal datasets in our dating experiments (**Fig. SM5-1**): *(i)* the concatenated nuclear *CRC*-LFYi2 individual (strict) consensus data (gaps treated as missing data), (*ii*) complete chloroplast genome (plastome) data sets pruned to the 20 individuals for which *CRC* and LFYi2 data were available and representing all major plastid haplotypes, *(iii)* a species-level nuclear data set combining data used in earlier studies with our newly assembled *CRC* and LFYi2 data, with the respective gene sets filtered for fit with the three nuclear topological scenarios covering all major aspects of the beech species network (beech Coral of Life; **Fig. SM5-2**), and *(iv)* a set of 100 fossil occurrences to generate meta-calibrated dated trees. For signal testing purposes, we also ran *CRC* and LFYi2 separate inferences, and logically partitioned subsets of the plastome data.

#### 5.1 *Initial node dating of chloroplast against the background of nuclear data*

**Rationale**—Straightforward association of fossil occurrences to plastid lineages is not feasible because plastid evolution is decoupled from speciation processes expressed in morphological differentiation (phenotypes). For instance, the initial subgeneric split, reflecting the putative reciprocal holophyly of both subgenera, is well-substantiated morphologically and using most nuclear gene regions (Denk et al. 2005, Oh et al. 2016, Renner et al. 2016, Jiang et al. 2022, Cardoni et al. 2022; see also **Section 2.8**), but not in chloroplast data (Worth et al. 2021; this study). The members of *Fagus* subgenus *Englerianae* harbour three (Lineage II–IV) of the five main chloroplast lineages, two of which (II, IV) are shared with geographically close members of subgenus *Fagus*. Lineage III, only found in *F. multinervis* and the geographically closest *F. japonica* population (*F.* subg. *Englerianae*), is a potential sister lineage of the West-Eurasian beeches (Lin. V; *F.* subg. *Fagus*). Thus, to be able to link fossil occurrences with plastid lineages, we first inferred a node-dated plastid chronogram using a single root age constraint, and compared it to analogously inferred, equally scaled, nuclear chronograms. We then visually mapped our fossil samples on the inferences. Against the background of the tectonic and climate evolution of the northern hemisphere (Scotese 2014, Scotese et al. 2014), we selected fossils as age constraints for the meta-calibrated chronograms shown in the main-text.

##### 5.1.1 Root age constraint for preliminary node dating experiment

**Traditional root age constraint**—The divergence date for the root in our plastid tree, the crown age of *Fagus* defined by the split between New World and Old World plastid lineages, was set to a minimum of 47.3 Ma and a maximum of 51.5 Ma using a log-normal distribution prior (mean: 48.5 Ma) using the following settings: $\mu = 1.45$, $\sigma = 0.7$ and offset = 47. 52±2 Ma is the (older) age estimate of the early Eocene McAbee Flora (Denk and Dillhoff 2005), a mid- to high-palaeoaltitude, temperate fossil flora including various organs, of †*F. langevinii* Manchester & R.M.Dillhoff (Manchester & Dillhoff 2005). While †*F. langevinii* cannot be assigned to any of the modern-day species' lineages, it represents a modern-type beech by showing all characteristics defining the modern-day genus (Denk et al. 2005, Denk & Grimm 2009, Renner et al. 2016). Older records of beech are limited to dispersed pollen, which may still represent the stem lineage of modern-day genus *Fagus* including but not limited to the last common ancestor(s) of all modern beeches. At and shortly after ~ 50 Ma, *Fagus* foliage was present in western Greenland, high-latitude East Asia, northern North Pacific coastlands ('Beringia', Kamchatka to Alaska) and (montane) western North America (McAbee and other floras associated with the Okanagan Highlands; **SupplFossilTable.xlsx**, sheet *main list*). Thus, it can be argued that †*F. langevinii* reflects beginning crown-group radiation in modern beeches (genus *Fagus*) and gives a minimum age estimate for the root node in our trees.[4] The younger estimate for the McAbee Flora of 49±1 Ma was used to define the offset of the log-normal distribution. The radiometrically dated fossil beech pollen of Agatdalen, western Greenland (Danian, 64–62 Ma; Grímsson et al. 2016) was selected to define the maximum possible age, i.e. the width of the log-normal distribution.

**Alternative older root age constraint for the plastid tree**—Noting the simultaneous occurrence of (modern) *Fagus* pollen in the Paleocene of western Greenland and north-eastern Siberia (Grímsson et al. 2016; this study), and the high geographic correlation of modern-day plastid haplotypes, it is conceivable that the primary plastome split between the eastern (!) North American Lineage I and the Eurasian plastome lineages is a legacy of the (Sub-)Arctic Paleocene range of the first modern beeches (see also **SupplR&D.pdf**, *fig. SR4-2*). The Arctic-Atlantic plastome (Lin. I) was passed on to modern-day *F. grandifolia(-mexicana)*, when its (their) ancestors migrated into eastern North America from Beringia and western North America after the split of modern-day beeches into two subgeneric lineages. *Vice versa*, the Arctic-Pacific plastome (Lin. II to V) was passed on to all Eurasian beeches. Accordingly, we constrained the root MRCA as a lognormal prior with 60 Ma as mean, minimum of 58.8 Ma and maximum of 63.0 Ma using the following settings: $\mu = 1.45$, $\sigma = 0.7$ and offset = 58.5.

**Alternative younger root age constrained for the nuclear tree**—The nuclear divergence between the two subgenera may not reflect beginning crown group radiation but only the formation of the modern-day subgenera as morphologically distinct entities. Notably, fossils that can be unambiguously assigned to one subgenus, i.e. show only affinities to species of the same subgenus, amass in the fossil record at about the same time, around the Eocene-Oligocene boundary at ~34 Ma (**SupplFossilTable.xlsx**). Accordingly, we constrained the root MRCA as a lognormal prior with a minimum and maximum of 30.3 and 34.5 Ma (mean = 31.5) using the following settings: $\mu = 1.45$, $\sigma = 0.7$ and offset = 30.

[4] These fossils are stem fossils of the modern genus *Fagus* because they cannot be assigned to one of the subgenera of (modern) *Fagus* but they are *not* stem fossils of the *Fagus* lineage (monogeneric subfamily Fagoideae) as a whole. Using these Eocene fossils to inform the stem age of the *Fagus* lineage, hence, the minimum age of the MRCA of all Fagaceae, as done in various dating studies of Fagales and Fagaceae (e.g. Sauquet et al. 2014) makes no sense. The earliest fossils that can be assigned to the *Fagus* lineage (Fagoideae) and could inform a stem age of the lineage leading to modern-day *Fagus* and the Fagaceae-MRCA, respectively, are pollen from the Late Cretaceous (82–81 Ma) of North America (Grímsson et al. 2016).

##### 5.1.2 Topological constraints imposed on the combined nuclear data

Since the nuclear data captures two levels of speciation events, we applied three alternative multi-furcating constraints (**Fig. SM5-2**) to enforce the (putative) primary and secondary divergences in addition to the scenario preferred by BEAST 2:

- **Enforcing sequence of speciation events in *Fagus* subgenus *Englerianae***—According to the data of Jiang et al. (2022), *F. multinervis* is sister to *F. japonica* and *F. engleriana* (Cardoni et al. 2022). Also, in our *CRC* and LFYi2 data, we observed mutations unique to *F. multinervis,* while the *F. engleriana* individual falls completely within the variation observed for *F. japonica*. Thus, we constrained *F. multinervis* as the first-diverging lineage within the *F.* subgenus *Englerianae* subtree (files labelled as "…ENG…").
- **Enforcing primary sequence of speciation events in *Fagus* subgenus *Fagus***—Two experiments were conducted enforcing *(a)* an initial divergence between North American (*F. grandifolia-mexicana*) and Eurasian lineages of *Fagus* subgenus *Fagus* (files labelled "…NWOW…"), followed by (*b*) a split into a northern, lowland(?), temperate (nemoral; *F. caspica, F. crenata, F. sylvatica*) and 'southern' montane-subtropical (oreotropical, meridional; *F. hayatae, F. longipetiolata, F. lucida*) lineage (files labelled "…HighLow…"[5]), and subsequent dispersal (†*F. castaneifolia*) of the northern lineage via the 'Northern route' (north of the Paratethys; in analogy to *Quercus* section *Cerris*, Denk et al. 2023) to Western Eurasia followed by vicariance between Western Eurasian beeches and the north-eastern Asian *F. crenata* lineage (not enforced being consequential to either the "NWOW" and "HighLow" constraints). This scenario is in best agreement with non-coding nuclear (ITS, 5S-IGS, LFYi2) and morphological-historical data and inferences (Denk et al. 2005, Grimm et al. 2007, Denk & Grimm 2009, Renner et al. 2016) and most genes compiled by Jiang et al. (2022; as revisited in Cardoni et al. 2022).
- **Unconstrained, reflecting secondary sequence of reticulate speciation events—**The unconstrained dated phylogeny places the East Asian species, 'East Asian-subgenus-*Fagus* clade', as sister to an 'Euamerican clade' comprising the North American and western Eurasian species as preferred by *CRC* (**SupplR&D.pdf,** *section 1*) and some of Jiang et al.'s (2022) nuclear genes (Cardoni et al. 2022; see also **SupplDating.xlsx**, sheet *pre-FBD topology test*).

[5] Applying the "HighLow" constraint, i.e. *F. crenata* as sister to W. Eurasian spp. will automatically result in a topology, where *F. grandifolia-mexicana* is placed as sister to the rest of *Fagus* subgenus *Fagus* because of the attraction between *F. crenata* and the other E. Asian spp.

#### 5.2 Final dating using fossilized birth-death model and traditional node dating

The history of beech represents a case of strongly reticulate evolution, it cannot be modelled by a 'species tree', a multi-species coalescent (MSC), but requires a 'species network', a multi-species coalescent network (MSCN). The only current software to infer a dated, explicitly inferred MSCN is PHYLONET (v. 3; Wen et al. 2018), which has, so far, not the capacity to model the reticulate evolution of beech. First, we lack the necessary amount of high-resolution data providing strong and unambiguous signals (see also Jiang et al. 2022, figs 2 and 3). Second, multiple phases of reticulation encoded in the nuclear genome of modern-day beeches involve too many reticulation events. Using the data at hand, PHYLONET is unable to capture and properly model all reticulation events apparent from the genetic differentiation in groups of nuclear gene regions as depicted in **Figure SM5-2**.

Thus, we chose an indirect approach to implicitly date the beech species network. We used fossilized birth-dating dating (FBD; Heath et al. 2014; as used in Renner et al. 2016) on accordingly filtered gene samples supporting and favouring the same topology (in analogy to the constraints used in **Section 5.1** but without having to enforce phylogenetic splits). In contrast to most applications of FBD in literature that rely on under-sampled fossil records and a subsequent low proportion of fossil sampling (typically set to < 10%), the comprehensive beech fossil record used here may in fact oversample the number of species lineages and speciation events modelled via the Yule birth-death prior. Furthermore, certain phenotypes (fossil morphotypes) may represent stem lineages, and the likelihood for this increases the further we go back in time. Hence, our FBD estimates are considered to represent maximum divergence estimates for major nuclear genetic differentiation events, initial speciation events, most of which will predate the final speciation event that formed the modern-day species. Given the high number of fossils assigned to clades (in total 80 for plastome, 77 for nuclear data; **SupplFossilTable.xlsx**, sheets *cpFBD, ncFBD*; see also **Section 5.2.2** below), our FBD-inferred chronograms can be considered to represent meta-calibrated dated topologies: their estimates are strongly constrained by the temporal distribution of the fossils assigned to each clade.

To establish minimum estimates for final species isolation and last contacts between species lineages, we used traditional node dating with very few node age priors. Traditional node dating is *per se* underestimating, i.e. can only give minimum estimates for divergence ages. In case the fossil priors are close to the speciation event, the estimate must be slightly too young, in case the fossil priors represent young crown fossils of a lineage or its extinct sister lineage, the estimates will be much too young (see e.g. many older dating estimates for the split between *Fagus* and *Quercus* compiled for timetree.org database). Here, we carefully selected the most-conservative, most-straightforward indicators of speciation events and indicator fossils for the last contact between species lineages.

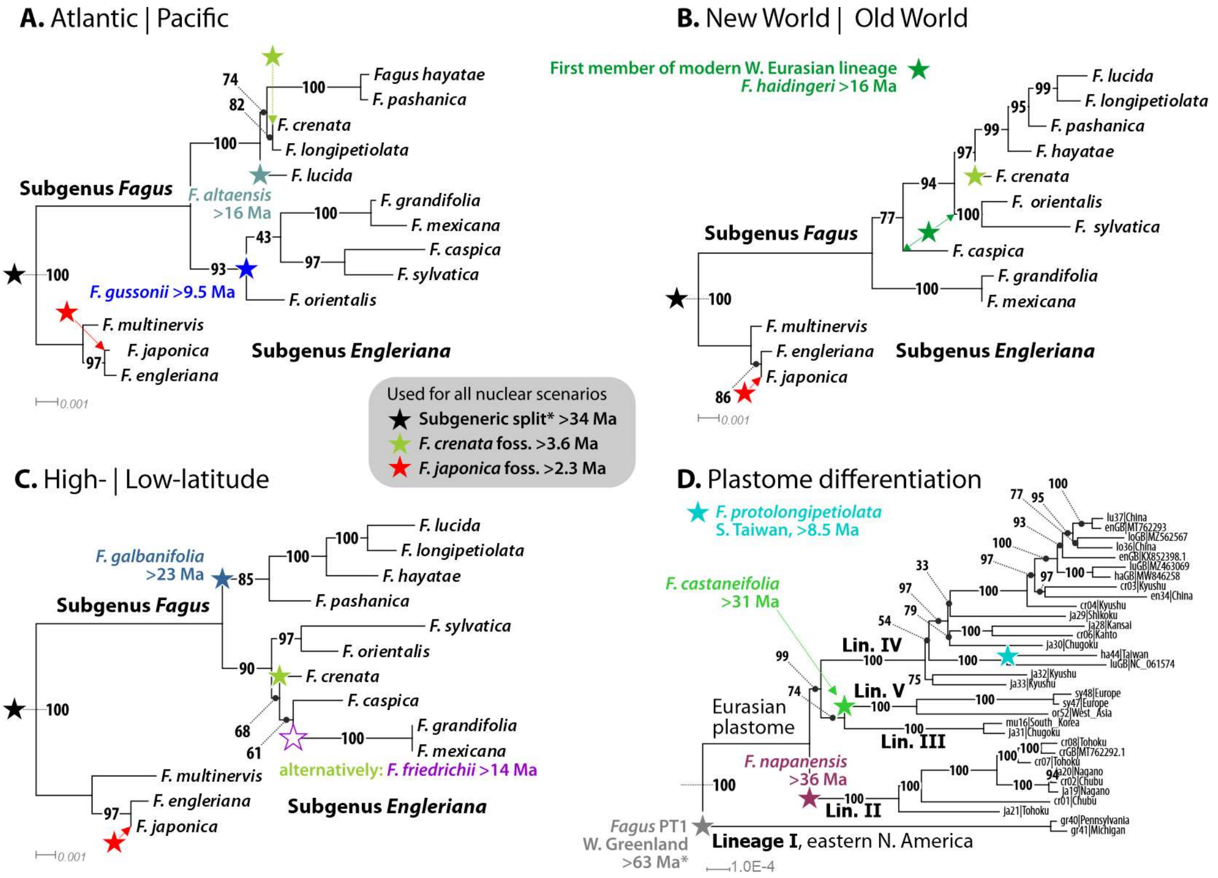


**Figure SM5-3** | Age priors used for establishing minimum estimates for final speciation events and last contacts between species lineages. **A.** Primary split within subgenus *Fagus* between Atlantic- (W. Eurasian, E. North American) and Pacific-bound (E. Asian) species. **B.** Between New (E. North American spp.) and Old World (Eurasian spp.) **C.** Between (originally) high-latitude, circum-arctic and (ancient) mid-latitude, continental-montane E. Asian species. **D.** For the plastome. The shown topologies are unconstrained, i.e. inferred directly from the congruence-filtered gene sample.

##### 5.2.1 Selection of fossil priors for node dating

The fossil priors were selected to match the critical topological aspect of the MSCN reflected in each alternative topological scenario as well as to provide conservative age priors for all major lineages implied by each scenario.

- The **Atlantic | Pacific** scenario (**Fig. SM5-3A**) is supported by 12 of the 31 nuclear loci/ markers. All of these markers indicate a secondary contact between the lineage of the eastern North American species and the precursors of (some of) the West-Eurasian species. Based on its phenotype, place and time, †*F. gussonii* Massolongo of the Mediterranean Miocene of Europe is a natural candidate for a vector species that could have carried North American alleles into the gene pool of †*F. haidingeri* Kováts, the direct precursor of the modern-day West-Eurasian species (Denk 1999c, 2004; Denk & Grimm 2009). Thus, it gives a minimum age for the divergence between the western (eastern N. American) and eastern Atlantic (European) beech populations. As age prior for the Pacific (East Asian) clade, we opted for †*F. altaensis* Kornilova & Rajushkina as the putative precursor exclusive to *F. lucida* (Denk & Grimm 2009), hence, informing this species' stem age.

- Biogeographically, palaeontologically and morphologically obvious, the **New World | Old World** scenario (**Fig. SM5-3B**) is supported by 6 of the 31 nuclear loci/ markers, most importantly the ITS differentiation patterns, with one most distinct ITS lineage, ITS Lineage 2 being exclusive to *F. grandifolia* (no data for *F. mexicana* so far) and another (ITS Lin. 4) shared by all Eurasian species of *Fagus* subgenus *Fagus* (Denk et al. 2005); the sequence similarity observed in ITS Lineage 4 can only be explained by a common ancestor shared by and longer ongoing gene flow between the precursors of the Eurasian species than with the precursors of the North American *F. grandifolia*. Most nuclear gene regions that support a primary divergence between the North American and Eurasian species of subgenus *Fagus* prefer further to place the West-Eurasian species as sister lineage to the East Asian species (see also Cardoni et al. 2022). The formation of the West-Eurasian beech lineage is reflected in the fossil record by †*F. castaneifolia* Unger, a fossil species stretching from north-central Asia to Europe and the Mediterranean, that goes extinct in its most eastern range and is gradually replaced by †*F. haidingeri* in the remainder of its original areal (Denk 2004; Denk & Grimm 2009). Morphologically, †*F. castaneifolia* shows a much stronger resemblance to species further east, as expected for a precursor and stem species of the West-Eurasian beeches, hence, it provides a best-possible age prior for the stem age of the West-Eurasian lineage.
- In the **High- | Low-latitude** scenario (**Fig. SM5-3C**), we used Pleistocene fossils of *F. crenata* to inform stem age of this species (Uemura 1980), i.e. the point in time where it may have been finally disconnected from cross-species, cross-subarctic gene flow with other high-latitude species of *Fagus* subgenus *Fagus*. Older fossils resembling *F. crenata* still have affinities to other species. This scenario implies a primary speciation event between the fully temperate species of (today's) mid- to high-latitudes and a (lower latitude) southern East Asian lineage, the latter passing its genetic legacy only into the

modern-day species of China and Taiwan. Notably, *F. longipetiolata* and *F. pashanica* show a prominent ITS dimorphism: in addition to the widely shared ITS Lineage 4 (found in all Eurasian species of *F.* subg. *Fagus*), they include variants of the substantially distinct ITS Lineage 3. Further signals indicating a deep split between *F. longipetiolata, F. lucida, F. hayatae* and *F. pashanica* and *F. crenata* can be found in five of the 28 nuclear loci compiled by Jiang et al. (2022); this scenario has a correlation in the plastome differentiation (most-distinct, early diverged Lineage IV haplotype found in *F. hayatae* and a *F. lucida* individual, reportedly from Hunan, i.e. south-eastern margin of the species' distribution in China; see **SupplR&D.pdf**, *section 2.3.3*). A key fossil for the establishment of a unique lower latitude lineage that passed its legacy into these four species, being a precursor of some of them and propagated by secondary contact between e.g. the *F. longipetiolata* and *F. lucida* lineages, is †*F. galbanifolia* Guo. As the oldest fossil phenotypically linked to this group (this study) in contrast to coeval species showing a closer similarity to the modern-day *F. crenata*, it represents the best-possible stem age prior for a beech lineage exclusive to low(er) latitudes in continental-montane East Asia.

As outlined before, plastomes are mainly geographically sorted in beech, other Fagaceae and across Fagales (Yang et al. 2021, fig. 1, for Fagales[6]; Zhou et al. 2022, for Fagaceae; Li et al. 2025, for *Quercus* subgenus *Cerris*). Selection of age priors is hence only secondarily informed by their phenotype but primarily by their age and place. The root age, equalling the split between the New World plastomes of Lineage I and the Old World plastomes of Lineages II to V, is constrained by the same western Greenland fossil as in the preliminary node dating (→ **Section 5.1.1**). To constrain the second divergence event, the emergence of the putatively northern Pacific (oceanic) plastome Lineage II, we chose †*F. napaensis* Fotjanova as the oldest fossil from Beringia; notably its also the oldest fossil that shares first features with modern-day *Fagus* subgenus *Engleriana*. We added two tip age priors: (*i*) the oldest †*F. castaneifolia* fossil for the stem age of the unique and exclusive West-Eurasian Lineage V; and (*ii*) the Taiwanese †*F. protolongipetiolata* Huzioka, being the oldest fossil exclusive to the island and probably representing the or a direct precursor of the modern-day *F. hayatae* (**Fig. SM5-3D**).

##### 5.2.2 Set-up for FBD and node dating

As BEAST 2 could not implement the best substitution model for each gene region as determined by MEGA 11 (Tamura 2021), the three datasets were partitioned with the nucleotide substitution model for all partitions set to a GTR+Γ+I substitution model, where between-site

[6] The paper's text doesn't include any reference to or discussion of this result of their analysis.

heterogeneity is modelled via a gamma function (+Γ) and involving a number of invariant sites (+I). The Atlantic | Pacific nuclear dataset consisted of 9618 bp in twelve partitions, the Old World | New World dataset of 4205 bp in six partitions, and the High- | Low-Latitude dataset of 4537 bp in seven partitions (**Table SM5-1**). BEAST analyses used MCMC chains with $10^7$ to $6{*}10^7$ generations sampling every 5000$^{th}$ generation. The first 10% were discarded as burn-in, and multiple MCMC chains were run to check for run performance in TRACER (Rambaut et al. 2018).

**Table SM5-1 | Dimensions and composition of datasets used for dating.**

| Dataset; scenario | Extant taxa (fossil taxa) | Total bp | Locus name[a] |
|---|---|---|---|
| Nuclear; Atlantic \| Pacific | 13 (97) | 9618 | CRC, LFYi2, F202, F289, P12, P14, P21, P28, P52, P54, P72, P97 |
| Nuclear; Old World \| New World; | 13 (96) | 4205 | LFYi2, ITS, F159, P28, P48, P49 |
| Nuclear; High Latitude \| Low Latitude | 13 (96) | 4537 | LFYi2, ITS, F114, F253, P14, P28, P49 |
| Plastome | 33 (100) | 132526 | - |

[a] Locus names are *CRABS CLAW* (CRC), second intron of *LEAFY* gene (LFYi2), ITS region of the nuclear-encoded 35S rDNA (ITS) and other locus names according to Jiang et al. (2022).

The results were visualized with FIGTREE v1.4.4 (Rambaut 2014). A summary of the node dating (ND) and fossilized birth-dating dating (FBD) runs is given in (**Table SM5-2**) including corresponding ESS values.

**Table SM5-2 | ESS values for dating experiments.** Runs marked with asterisks treated the modern-type fossils from the Kraskino palaeoflora as potential stem members of *Fagus* subgenus *Fagus* rather than assigning them to species-level clades.

| | Scenario | Additional topological priors for FBD | Corresponding priors in ND (only extant spp.) | Approach | Chains | ESS-Posterior | ESS-Likelihood | ESS-Prior |
|---|---|---|---|---|---|---|---|---|
| Nuclear species consensi | Atlantic\|Pacific | Euamerican clade, East Asian clade | N. American + W. Eurasian spp., Chinese-Taiwanese spp. | FBD | 50000000 | 482 | 130 | 568 |
| | | | | FBD* | 50000000 | 798 | 726 | 743 |
| | | | | ND | 60000000 | 236 | 459 | 189 |
| | High\|Low | High latitude clade, Southern lineage | *F. crenata* + N. American + W. Eurasian spp. + | FBD | 50000000 | 569 | 385 | 550 |
| | | | | FBD* | 10000000 | 216 | 129 | 154 |
| | | | … (a) Chinese-Taiwanese spp. | ND | 60000000 | 350 | 1437 | 337 |
| | | | … (b) *F. hayatae* + *F. pashanica* | ND | 40000000 | 37 | 34 | 33 |
| | New World\|Old World | Eurasian Fagus, Southern lineage | Eurasian spp. of subg. *Fagus* + | FBD | 50000000 | 861 | 1295 | 815 |
| | | | | FBD* | 10000000 | 139 | 305 | 142 |
| | | | … (a) no additional prior | ND | 60000000 | 242 | 985 | 228 |
| | | | … (b) *F. hayatae* + *F. pashanica* | ND | 40000000 | 243 | 979 | 232 |
| Complete plastomes | (geographically sorted) | | GTR-Option 1 | FBD | 10000000 | 11 | 5 | 120 |
| | | | | ND | 60000000 | 599 | 1960 | 586 |
